# Targeted long-read sequencing resolves complex structural variants and identifies missing disease-causing variants

**DOI:** 10.1101/2020.11.03.365395

**Authors:** Danny E. Miller, Arvis Sulovari, Tianyun Wang, Hailey Loucks, Kendra Hoekzema, Katherine M. Munson, Alexandra P. Lewis, Edith P. Almanza Fuerte, Catherine R. Paschal, Jenny Thies, James T. Bennett, Ian Glass, Katrina M. Dipple, Karynne Patterson, Emily S. Bonkowski, Zoe Nelson, Audrey Squire, Megan Sikes, Erika Beckman, Robin L. Bennett, Dawn Earl, Winston Lee, Rando Allikmets, Seth J. Perlman, Penny Chow, Anne V. Hing, Margaret P. Adam, Angela Sun, Christina Lam, Irene Chang, University of Washington Center for Mendelian Genomics, Tim Cherry, Jessica X. Chong, Michael J. Bamshad, Deborah A. Nickerson, Heather C. Mefford, Dan Doherty, Evan E. Eichler

**Affiliations:** Department of Genome Sciences, University of Washington School of Medicine, Seattle, WA, 98195, USA; Department of Pediatrics, Division of Genetic Medicine, University of Washington and Seattle Children’s Hospital, Seattle, WA, 98105, USA; Department of Laboratories, Seattle Children’s Hospital, Seattle, WA, 98105, USA; Department of Laboratory Medicine and Pathology, University of Washington, Seattle, WA, 98195, USA; Center for Developmental Biology and Regenerative Medicine, Seattle Children’s Research Institute, Seattle, WA, 98101, USA; Brotman Baty Institute for Precision Medicine, Seattle, WA, 98195, USA; Center for Clinical and Translational Research, Seattle Children’s Research Institute, Seattle, WA, 98101, USA; Division of Medical Genetics, Department of Medicine, University of Washington, Seattle, WA, 98195, USA; Department of Genetics and Development, Columbia University, New York, NY 10032, USA; Department of Ophthalmology, Columbia University, New York, NY 10032, USA; Department of Pathology and Cell Biology, Columbia University, New York, NY 10032, USA; Department of Neurology, Seattle Children’s Hospital, University of Washington, Seattle, WA, 98105, USA; Department of Pediatrics, Division of Craniofacial Medicine, University of Washington, Seattle, WA, 98195, USA; Center for Integrative Brain Research, Seattle Children’s Research Institute, Seattle, WA, 98101, USA; Department of Pediatrics, Division of Developmental Medicine, University of Washington and Seattle Children’s Hospital, Seattle, WA, 98105, USA; Howard Hughes Medical Institute, University of Washington, Seattle, WA, 98195, USA

## Abstract

**BACKGROUND:** Despite widespread availability of clinical genetic testing, many individuals with suspected genetic conditions do not have a precise diagnosis. This limits their opportunity to take advantage of state-of-the-art treatments. In such instances, testing sometimes reveals difficult-to-evaluate complex structural differences, candidate variants that do not fully explain the phenotype, single pathogenic variants in recessive disorders, or no variants in specific genes of interest. Thus, there is a need for better tools to identify a precise genetic diagnosis in individuals when conventional testing approaches have been exhausted.

**METHODS:** Targeted long-read sequencing (T-LRS) was performed on 33 individuals using Read Until on the Oxford Nanopore platform. This method allowed us to computationally target up to 100 Mbp of sequence per experiment, resulting in an average of 20x coverage of target regions, a 500% increase over background. We analyzed patient DNA for pathogenic substitutions, structural variants, and methylation differences using a single data source.

**RESULTS:** The effectiveness of T-LRS was validated by detecting all genomic aberrations, including single-nucleotide variants, copy number changes, repeat expansions, and methylation differences, previously identified by prior clinical testing. In 6/7 individuals who had complex structural rearrangements, T-LRS enabled more precise resolution of the mutation, which led, in one case, to a change in clinical management. In nine individuals with suspected Mendelian conditions who lacked a precise genetic diagnosis, T-LRS identified pathogenic or likely pathogenic variants in five and variants of uncertain significance in two others.

**CONCLUSIONS:** T-LRS can accurately predict pathogenic copy number variants and triplet repeat expansions, resolve complex rearrangements, and identify single-nucleotide variants not detected by other technologies, including short-read sequencing. T-LRS represents an efficient and cost-effective strategy to evaluate high-priority candidate genes and regions or to further evaluate complex clinical testing results. The application of T-LRS will likely increase the diagnostic rate of rare disorders.

## INTRODUCTION

Routine use of genetic testing in clinical and research settings has improved diagnostic rates and uncovered the genetic basis for many rare genetic conditions, yet approximately half of individuals with a suspected Mendelian genetic condition remain undiagnosed.^1–4^ Broadly, undiagnosed individuals fall into two main categories: (i) those with a DNA sequence variant or structural difference that does not fully fit their phenotype (i.e., variant of unknown significance), and (ii) those in whom routine clinical evaluation—including exome sequencing—failed to reveal any candidate variants or identified only a single variant for a recessive condition that fits the phenotype. Thus, new tools and technologies that provide a comprehensive and accurate survey of genetic variation have the potential to improve diagnostic rates.

Clinical testing methods such as chromosome microarray and exome sequencing do not provide a complete view of human genetic variation. Structural variants (SVs) such as repeat expansions, insertions, deletions, or rearrangements may account for many of the pathogenic variants that go undetected,^5^ but they are challenging to identify using existing short-read sequencing technology. Long-read sequencing (LRS) technology, which sequences native DNA molecules, can generate reads from 1,000 to over 1 million base pairs in length while also providing information on DNA methylation.^6^ The improved performance of LRS for SV detection has been demonstrated.^7^ However, generating sufficient LRS data for genome-wide analysis remains prohibitively expensive, which makes studies comparing short-read sequencing to long-read sequencing challenging and slows clinical implementation.

Current methods allow LRS of targeted genomic regions (targeted long-read sequencing [T-LRS]) either by PCR enrichment or Cas9-mediated isolation of targets.^8,9^ However, these methods typically remove critical information such as methylation status, take time to design and optimize, and are restricted to a relatively modest number of genomic targets. To overcome these limitations, we implemented a computational method to select and sequence native DNA using Oxford Nanopore Technologies (ONT). This method, known as Read Until, accepts or rejects DNA molecules for sequencing based on set target sequences and can be modified in real time.^10^

We assessed the specificity and sensitivity of T-LRS using Read Until to detect known pathogenic SVs, such as copy number variants (CNVs), repeat expansions, and translocations by sequencing 33 individuals in whom such variants were identified in the course of clinical testing (Table S1). In 6/7 persons who had complex structural rearrangements, T-LRS enabled more-precise resolution of the mutation, which led, in one case, to a change in clinical management. We applied T-LRS to nine persons with a known or suspected autosomal recessive or X-linked Mendelian condition in whom either only one (*n*=7) or no pathogenic variants (*n*=2) were found by standard clinical testing and identified likely causal variants in seven of nine. Our results demonstrate the potential added value of T-LRS as a clinical test to efficiently and cost-effectively evaluate patients with complex SVs or to identify causal variants in high-priority candidate genes.

## METHODS

### Selection of individuals

Individuals were identified based on previous clinical or research testing results, which included chromosome microarray, karyotype, clinical exome sequencing, or research whole-genome sequencing. Individuals with complex copy number changes were defined as those with two or more CNVs or one CNV and at least one translocation. Persons with “missing” variants were defined as those in whom clinical testing had identified one pathogenic variant in a gene associated with an autosomal recessive disorder or no variants in a gene associated with an X-linked disorder.

### DNA isolation and library preparation

DNA for sequencing was isolated from blood, saliva, or fibroblasts using standard methods (Table S1). Approximately 1.5 μg of DNA sheared to a target size of 8–12 kbp was used to make sequencing libraries (Supplementary methods).

### Sequencing and selection of target regions

Sequencing was performed using the ONT GridION sequencing platform. We applied the adaptive sequencing mode known as Read Until using the ReadFish software package, which allowed us to dynamically select target regions for sequencing.^10^ In this mode, software analyzes the signal after a DNA molecule enters a pore to determine whether that molecule lies within a specified genomic region of interest. If it does, the pore continues to sequence the molecule; if not, the DNA molecule is ejected from the pore. In cases with complex CNVs, we targeted large genomic regions on either side of the known aberration. For cases in which a single gene was suspected, at least 100 kbp of DNA surrounding the gene was targeted for sequencing (Table S2). In all cases, standard regions were targeted on multiple chromosomes to serve as internal copy number and coverage controls.

### Sequence analyses

FASTQ files were generated using guppy 4.0.11 and aligned to GRCh38 using both minimap2^11^ and NGMLR^12^ with default parameters. Variants were called using Longshot^13^, Clair^14^, and medaka (https://github.com/nanoporetech/medaka). VCF files that combined all variant calls were annotated with variant effect predictor annotations^15^ and CADD scores^16^. Novel intronic variants or those with allele frequencies <2% were annotated using SpliceAI.^17^ Copy number changes and breakpoint transitions were identified using circular binary segmentation.^18^ SVs were identified using both Sniffles^12^ and SVIM^19^. Methylation changes were identified in select samples using Nanopolish.^20^

### Study approval

This study was approved by the institutional review board at the University of Washington under protocols 7064 (University of Washington Repository for Mendelian Disorders), 4125, and 28853. All participants or their legal guardians provided written informed consent.

## RESULTS

T-LRS using the Read Until approach allows for rapid selection and real-time discovery of pathogenic variants from candidate genomic regions.^10^ We applied this method by direct sequencing of DNA from blood, saliva, or cell lines (Table S1) from 37 persons (33 affected individuals and 4 parents) clinically diagnosed with a variety of genetic conditions (Table S1). Among these individuals, 11 had a known (i.e., detected by prior genetic testing) pathogenic CNV or translocation, 6 had known pathogenic repeat expansions, 7 had known complex rearrangements, and 9 had a clinical diagnosis of an X-linked or recessive Mendelian condition but either no known pathogenic variants (*n*=2) or only one known variant out of the expected two (*n*=7).

### T-LRS detects known pathogenic SVs

Eleven individuals were previously found to have a single pathogenic CNV or translocation detected by chromosomal microarray (CMA), karyotype, or short-read sequencing (Table S2). This set includes, for example, frequently observed recurrent deletions associated with autism and developmental delay (chromosomes 15q11.3, 16p11.2, 22q11.1, and 1q21.1). We generated 10–62x coverage of the target regions (1–33 Mbp) using a single flow cell for each individual. This sequencing-based approach identified breakpoints in the expected regions for all 11 persons (Table 1, Figures S1–S10, Table S3). In 4/11 cases, T-LRS provided additional information, including further refinement of the breakpoint region (*n*=3; BK144-03, BK364-03, S046), clarifying a duplication as tandem (BK364-03), and identifying a previously unknown unbalanced translocation (BK506-03). Evaluation of the underlying genic sequence on the normal homologous chromosomes overlapping the deleted segments found no pathogenic or likely pathogenic variants, consistent with a dominant effect of these SVs.

**Table 1.** Targeted long-read sequencing detects structural variants.

|  | Deletions | Duplications | Translocations | Inversions or Rearrangements | Total |
| --- | --- | --- | --- | --- | --- |
| Events identified by clinical testing | 18 | 11 | 7 | 1 | 37 |
| Events identified by clinical testing and missed by T-LRS | 0 | 0 | 0 | 0 | 0 |
| Events newly identified by T-LRS and not reported on clinical testing | 6 | 0 | 13 | 20 | 39 |
| Total | 24 | 11 | 20 | 21 | 76 |

### T-LRS detects known triplet repeat expansions and methylation status

Next, we focused on six persons carrying known repeat expansions associated with spinocerebellar ataxia, Friedreich’s ataxia, fragile X, and Baratela-Scott syndromes. Repeat expansions in the latter two are, in particular, difficult to detect and resolve using standard sequencing because of the length and high GC content of the repeats. Detecting hyper-expansion and methylation typically require time-consuming Southern blotting with methylation-sensitive enzymes to diagnose.^21,22^ We generated a minimum of 8x coverage for all six samples carrying pathogenic expansions in *FMR1*, *FXN*, *ATXN3*, *ATXN8OS*, or *XYLT1*. In each sample, we detected pathogenic repeat-sized alleles, and at least one read spanned the complete expansion, providing a more precise estimate on the allele size. We were also able to determine the exact sequence of the expanded allele (Figure S11–S18, Tables S4–S6). In some instances, especially with DNA from cell lines, the length of the expansion was more variable than anticipated. For example, a cell line heterozygous for an expansion within *FXN* was reported to have predominant alleles at 750 and 1030 repeat units while our sequencing-based estimate identified predominant repeats of 333 and 1049 repeat units. This finding is consistent with previous work showing repeat length instability in cell lines or somatic mosaicism of expanded alleles.^22^

Expansion of a GGC repeat in the 5’ untranslated region (UTR) of *XYLT1* was recently shown to be a common cause of Baratela-Scott syndrome mediated by methylation and transcriptional silencing^21^ (Figure 1A). T-LRS of two affected families from that study allowed us to simultaneously assay repeat length, sequence content, and methylation using a single test (Figure 1B,C). Comparing read length and methylation in each individual revealed that some reads for the premutation haplotype in the proband’s mother (individual 04-02) were methylated, suggesting that some, but not all, of her cells have silenced the expansion. Thus, T-LRS of native DNA molecules provides additional details not available when repeat length and methylation are assayed separately. Interestingly, methylation analysis in the *FMR1*-expanded CGG repeat obtained from a cell line revealed that the disease locus was no longer methylated despite containing an expansion of nearly 400 repeats. This finding is consistent with a recent observation that methylation status of fragile X full-mutation alleles between 200 to 400 is not stably maintained and, if observed in primary material from a patient, may predict a less severe phenotype.^23^

**Figure 1:**
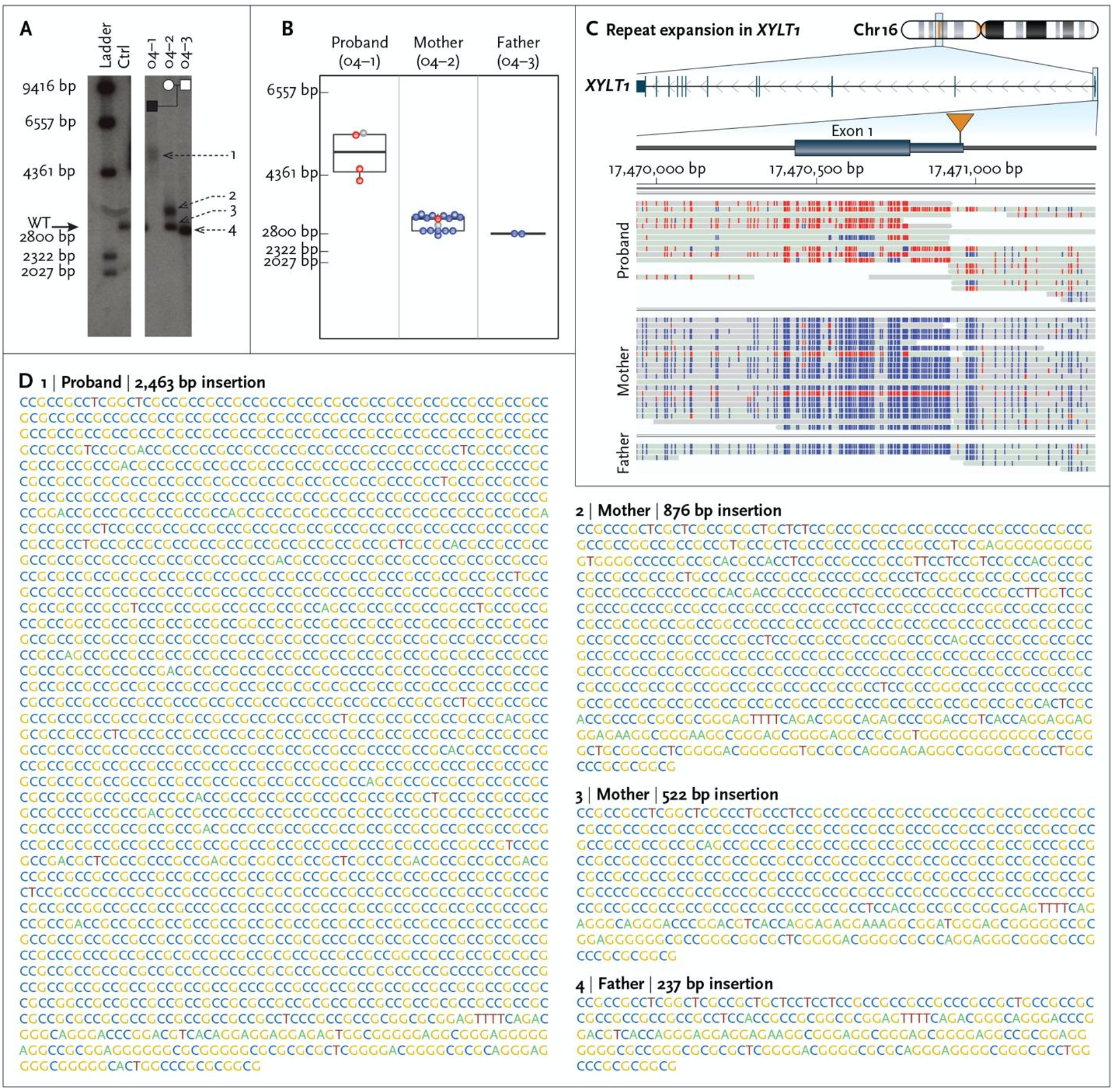
Targeted long-read sequencing simultaneously detects repeat expansion and methylation status. Expansion and methylation of a GGC repeat in the 5’ UTR of *XYLT1* is a common cause of Baratela-Scott syndrome. **A.** Southern blot of family 04 reported by LaCroix and colleagues ^21^ demonstrates that the proband (04-01) carries an expansion (1) of a region defined by two *Kpn*I restriction enzyme sites containing a GGC repeat, the mother (04-02) carries one premutation (2) and one wild-type allele (3), and the father (04-03) carries two wild-type alleles (4). Both panels are from the same Southern blot on day 6 of exposure. **B.** T-LRS of the trio revealed that the length of fragments from single reads spanning both *KpnI* cut sites used in panel A was consistent with the results from the Southern blot. Colored dots in panel B correspond to methylated (red) and non-methylated (blue) reads shown in panel C; gray represents reads where methylation status was not determined. **C.** Expansion of the GGC repeat in the proband results in methylation of the 5’ UTR and exon 1. Two reads in the mother are methylated (red), one of which spans the region between the *KpnI* cut sites and whose length is consistent with a premutation allele as shown in panel B. The second methylated read terminates within the repeat and the length cannot be assayed. **D.** Sequence of the GCC repeat haplotypes as determined by PacBio CLR sequencing is consistent with the Southern blots and the lengths identified using ONT.

### T-LRS enables further characterization of complex structural rearrangements

To assess the added diagnostic value of T-LRS, we selected seven individuals in whom routine clinical testing using CMA or karyotype revealed complex structural changes classified as pathogenic, such as multiple noncontiguous CNVs or rearrangements affecting multiple chromosomes. We hypothesized that T-LRS would identify additional rearrangements or CNVs that would be clinically informative. Samples were sequenced by targeting 15–145 Mbp of genomic sequence generating 7–39x coverage of each target. We identified and refined deletion and duplication breakpoints using a binary segmentation algorithm to delineate transitions in read-depth (Supplementary methods; Figure S1). Our analysis identified all previously reported events, further refining the rearrangements in three of seven individuals: a common duplication (S021), the breakpoints of a focal amplification (S022), and one tandem duplication (S035) (Table 1; Figures S19–S25; Tables S7–S9). In three other individuals, we detected additional CNVs, rearrangements, or translocations of potential clinical relevance, while in the remaining individual (S023), we found no further abnormalities. For example, in patient S014, a CMA identified three noncontiguous deletions of chromosome 6 spanning a 5 Mbp interval. T-LRS of 15 Mbp surrounding the known deletions revealed two additional deletions and an additional rearrangement not associated with a deletion (Figure 2A; Tables S10, S11). Thus, the analysis resolved the structure of the region and identified new candidate genes for further consideration, such as *IPCEF1* and *CNKSR3*. We identified more extensive chromosomal differences in additional affected individuals, such as S020, in whom clinical testing identified multiple deletions and translocations. To evaluate these differences further, we targeted 74 Mbp of sequence and obtained approximately 27x coverage of four target regions using one ONT flow cell (Table S2). However, analysis of these regions indicated rearrangements involving regions outside the targeted area, so a second flow cell was run, targeting 77 Mbp of additional sequence. In total, T-LRS revealed the precise position of 11 translocations, 13 intrachromosomal rearrangements, and 6 deletions that directly impacted 12 genes, 11 of which were not reported by clinical testing. All 30 structural breakpoints were subsequently validated by low-coverage PacBio HiFi whole-genome sequencing (WGS) (Figure 2B; Table S12). Among the 12 genes disrupted by an SV, two may be associated with autosomal dominant arrhythmogenic right ventricular dysplasia (*CTNNA3*) and thoracic aortic aneurysms (*PRKG1*)—two clinical findings (Table S13). As a result, this individual was referred to cardiology for additional evaluation and anticipatory monitoring for dysrhythmias. Similar to patient S020, clinical testing identified multiple SVs in patient S036. T-LRS identified two additional deletions, five rearrangements, and six translocations not previously detected. In total, these events bisected seven genes, only two of which were reported based on prior clinical testing (Tables S14, S15).

**Figure 2.**
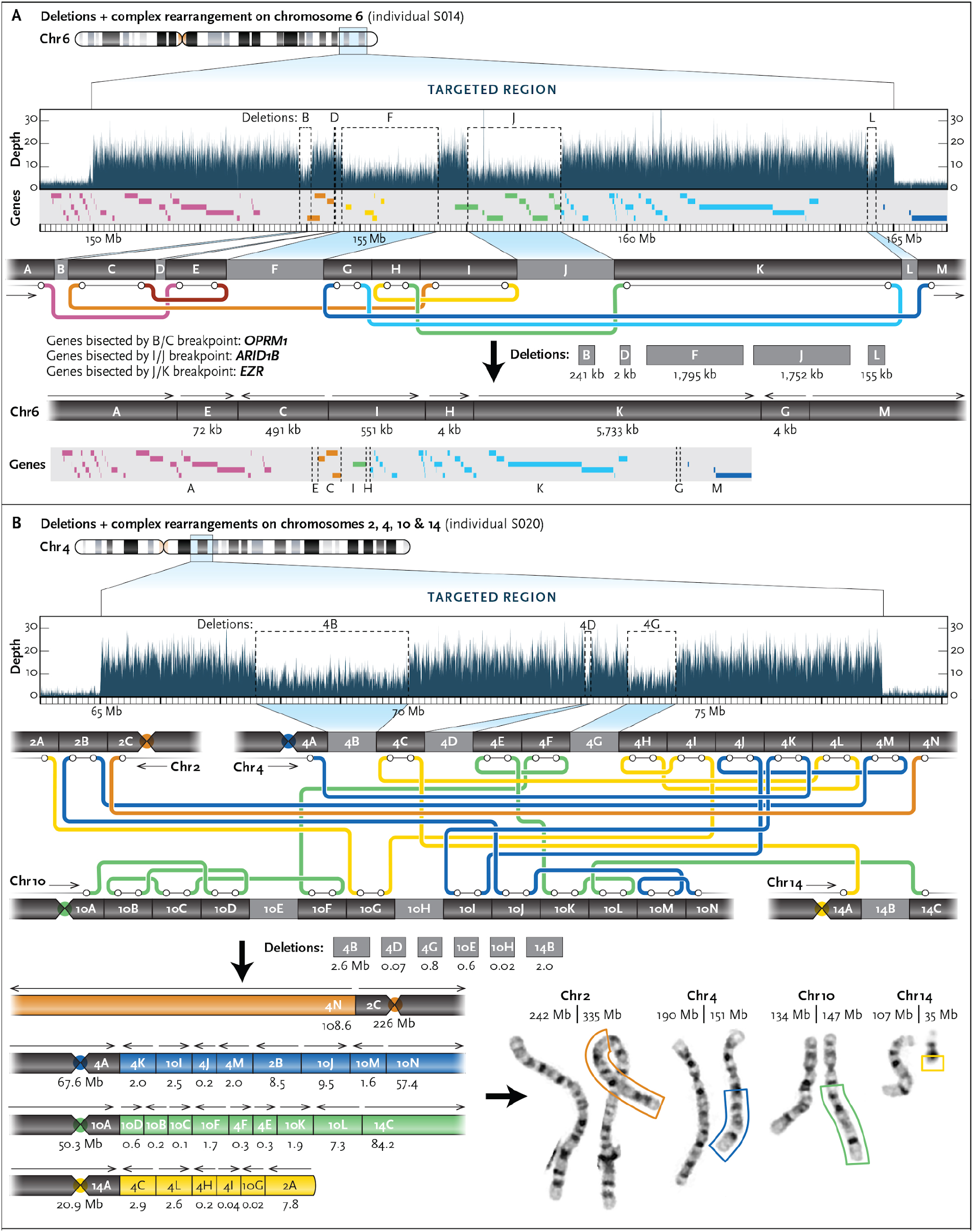
Targeted long-read sequencing identifies additional structural differences not observed by standard clinical testing. **A.** T-LRS of individual S014 revealed two additional deletions and one rearrangement (inversion) not reported by CMA. Reanalysis of the CMA data confirmed deletion L. The “subway” plot shows how each region is connected and allows for reconstruction of the new DNA sequence and gene order in the individual. **B.** Clinical CMA of individual S020 identified three deletions on chromosomes 4 and 14 and the subsequent karyotype revealed a complex translocation involving chromosomes 2, 4, 10, and 14. T-LRS identified 11 translocations, 13 rearrangements, and 6 deletions directly affecting 12 genes. Reconstruction of each derivative chromosome estimates the size of each event, as represented by the boxes surrounding part of the derivative chromosomes on the karyotype and is consistent with expected sizes based on karyotype.

### T-LRS identifies missing variants for recessive and X-linked Mendelian conditions

We performed T-LRS on nine individuals in whom clinical testing or follow-up research studies revealed only a single variant in a gene associated with a recessive condition (*n*=7) or no variants in genes associated with an X-linked condition (*n*=2) (Table 2). Each of these individuals had a strongly suspected clinical diagnosis but the molecular diagnosis was missing or incomplete. Using ACMG criteria,^24^ T-LRS revealed a pathogenic or likely pathogenic variant in four of seven persons with suspected recessive or X-linked conditions, and a variant of uncertain significance (VUS) in one; no second candidate variant was found in two others (S004 and S018) (Figure 3; Table 2; Figures S26–S34; Tables S16–S18). A VUS and a pathogenic variant were identified in the two persons with suspected X-linked conditions. The newly discovered variants included deletions, mobile element insertions, inversions, repeat expansions, and intronic variants predicted to affect splicing. In five of nine patients, we generated the data using a single ONT flow cell (Table S1).

**Table 2.**
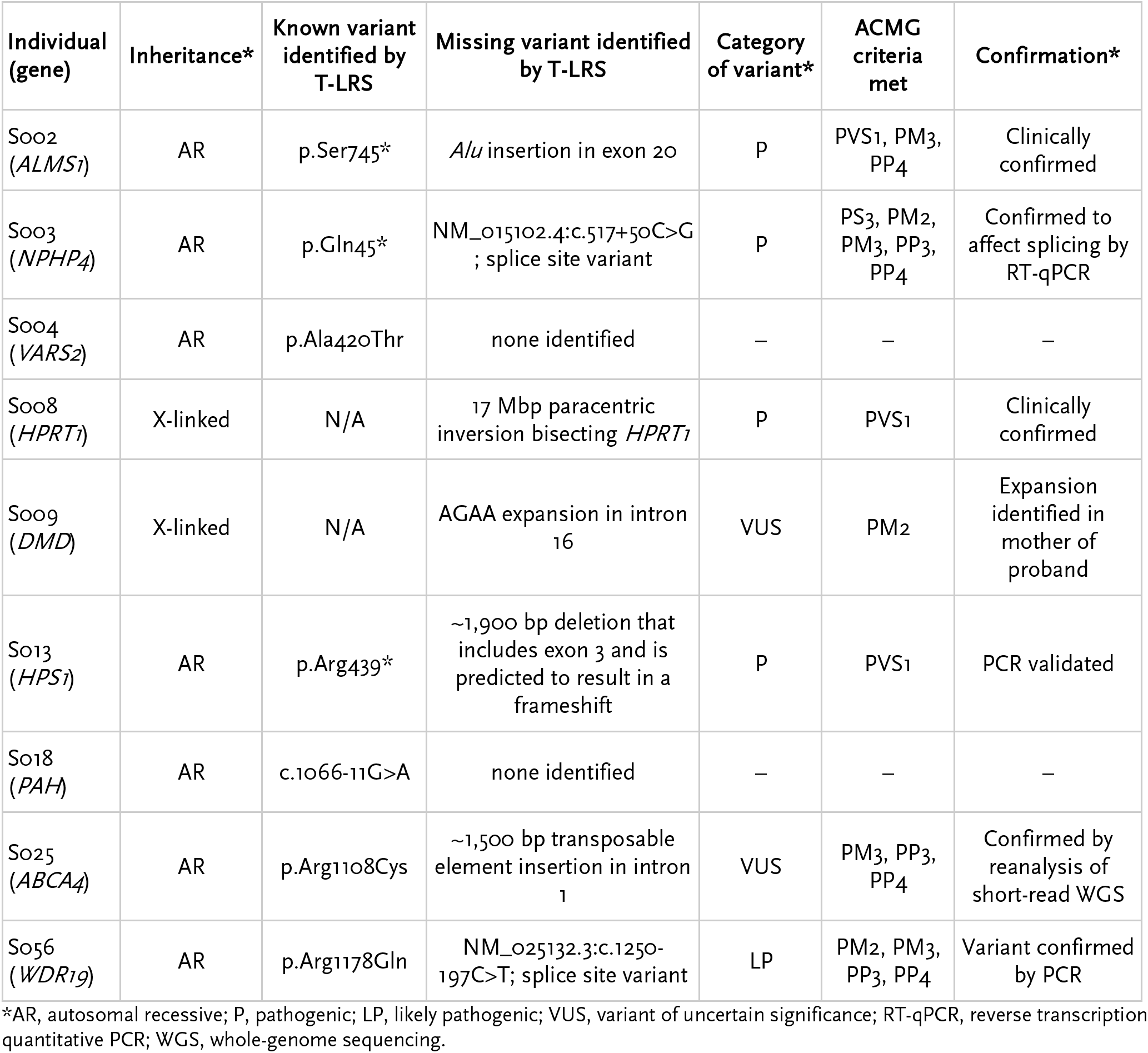
Targeted long-read sequencing identifies missing variants.

**Figure 3.**
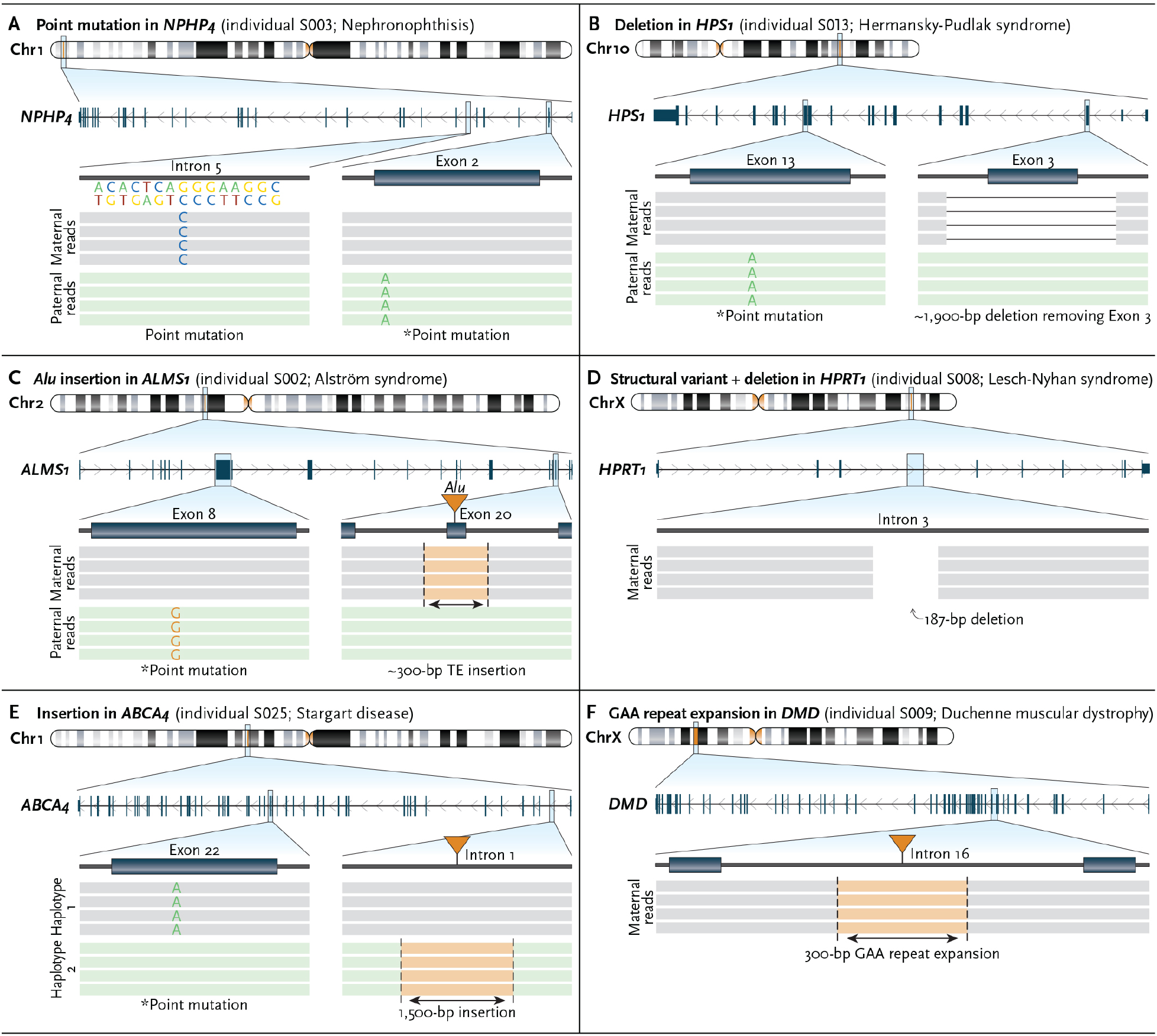
Targeted long-read sequencing identifies variants not detected by clinical testing. Pathogenic, likely pathogenic, or variants of uncertain significance (VUS) identified by T-LRS along with variants identified by prior clinical testing (denoted by an asterisk). **A.** T-LRS detects the known paternally inherited stop-gain as well as a candidate intronic splice acceptor variant. Long-read phasing demonstrates that these variants are in trans. **B.** A 1,900 bp deletion within *HPS1* removes exon 3 and results in a frameshift; phasing revealed that this variant and the previously known paternally inherited stop-gain occur on different haplotypes. PCR of genomic DNA validated the deletion and clinical confirmation is pending. **C.** T-LRS reveals a previously known paternally inherited stop-gain as well as a novel *Alu* insertion in exon 20 of *ALMS1*. Subsequent clinical testing confirmed the *Alu* was pathogenic and maternally inherited. **D.** A 187 bp deletion and 17 Mbp inversion disrupts *HPRT1*. Clinical testing confirmed the presence of an inversion. **E.** Insertion of a 1,500 bp composite retrotransposable element is predicted to create multiple splice acceptor and donor sites and represents a candidate second hit. Linkage disequilibrium phasing suggests the variants are on different haplotypes. **F.** Expansion of an AGAA repeat within *DMD* represents a VUS in an individual with Duchenne muscular dystrophy and a family history lacking a genetic diagnosis.

Sequencing of two individuals with suspected recessive disorders, S003 (Nephronophthisis, *NPHP4*) and S056 (Cranioectodermal dysplasia, *WDR19*), revealed that both carried rare intronic variants predicted to affect splicing located on the opposite haplotype from the known pathogenic variant (Figure 3A). In a fibroblast cell line from S003, we confirmed aberrant splicing by PCR and Sanger sequencing (Figure S27). In S003, we also identified heterozygous intronic GA-rich tandem repeat expansions with both haplotypes fully spanned by at least one long read. Because both expansions are within the range previously observed in controls,^25^ we were able to exclude them as candidate second hits, which would have been challenging to conclude using short reads alone.

In an individual with Hermansky-Pudlak syndrome (S013) and a known paternally inherited stop-gain variant, T-LRS revealed a novel 1,900 bp deletion on the maternal haplotype not identified by CMA or exome sequencing. The deletion spanned all of exon 3, resulting in a frameshift (Figure 3B, Figure S31). In an individual with Alström syndrome and a known paternally inherited stop-gain variant (S002), we identified a novel *Alu* repeat mobile element insertion in exon 20 not identified by clinical exome sequencing, which was confirmed by a clinical laboratory as a pathogenic second hit (Figure 3C, Figure S26). S008 was an individual with biochemically confirmed Lesch-Nyhan syndrome in whom T-LRS identified a 187 bp deletion within intron 3 of *HPRT1*, where evaluation of the flanking reads suggested a 17 Mbp paracentric inversion that was clinically confirmed using FISH (Figure 3D, Figure S29). Research-based WGS and targeted sequencing of *ABCA4* and locus in S025, an individual with Stargardt disease, failed to identify a 1,500 bp composite retrotransposable element insertion consisting of AluJ (SINE) and partial L2a, L2c, L2d2, and L1HS (LINEs) mapping within the first intron of *ABCA4*. We identified the event using both SV callers applied in this study and found that it mapped to a different haplotype than the known pathogenic variant. We categorized this as a VUS; however, consistent with previous work on similar insertions, *in silico* analysis with SpliceAI strongly suggests the insertion results in aberrant splicing of the first exon of *ABCA4* (Figure 3E, Figure S34, Table S18).

Finally, we used T-LRS to evaluate *DMD* in a family with multiple individuals affected by X-linked Duchenne muscular dystrophy lacking a precise genetic diagnosis. T-LRS of *DMD* in the proband (S009) revealed no candidate single-nucleotide variant, while SV analysis identified an intronic 117 AGAA repeat expansion (Figure 3F). The proband’s mother was heterozygous for this variant and the expansion was not found in his unaffected older brother (Figure S30). To determine the frequency of this expansion in a population sample, we analyzed nearly 9,000 short-read genomes^26^ using ExpansionHunter,^27^ identifying 72 individuals with 117 AGAA repeats or longer for an estimated population allele frequency of 0.4%. Remarkably, 71 (98.6%) of the individuals with large alleles are female—an observation inconsistent with Hardy-Weinberg equilibrium (OR = 52, p = 3e-16, Fisher’s exact test). Based on this information, we categorize this expansion as a high-priority VUS for future research investigation.

## DISCUSSION

Here, we show that T-LRS using Read Until on the ONT platform can be used to detect clinically relevant variants such as single-nucleotide variants, CNVs, repeat expansions, and methylation differences. Because target regions are computationally defined for sequencing, this technique is flexible and can be used to interrogate any part of the genome without the need to design specific experimental assays. Use of T-LRS also removes a substantial barrier to widespread clinical use of long-read technology by reducing per-sample costs of sequencing selected genes to a price point comparable to short-read WGS. The immediate potential clinical use of T-LRS includes screening of candidate genes in which existing technologies have failed to provide a precise genetic diagnosis, refinement of isolated or complex structural breakpoints, phasing of known variants, and evaluation of repeat structure.

T-LRS of specific genes in individuals with a clinical diagnosis of a recessive or X-linked condition, in whom a single variant or no candidate variants were identified, revealed a pathogenic variant, likely pathogenic variant, or VUS in more than 75% of instances. The discovery of these missing variants meant an end to the diagnostic odyssey and enabled testing of family members to determine their risk of being a carrier. Standard clinical testing, including WGS, achieves a precise genetic diagnosis in approximately 50% of families.^1–4^ While large-scale, prospective study of varied patient populations will be required to fully assess the advantages of T-LRS over conventional testing strategies, we anticipate use of T-LRS will increase the diagnostic rate for Mendelian conditions. Indeed, given only the small increase in diagnostic rate achieved by using short-read WGS to screen candidate genes or regions identified via exome sequencing or aCGH, T-LRS could be a more sensitive and cost-effective approach.

Clinical evaluation of SVs typically ends after identification of a single pathogenic CNV or a complex series of both CNVs and rearrangements. Here, we demonstrate that among 19 individuals with known or complex SVs, clinical testing identified only 49% (37/76) of the SVs found by T-LRS (Table 1). Additional SVs were recovered in 37% of persons (7/19) and in two persons this information revealed 16 additional genes directly disrupted by an SV. In one individual, the discovery of additional affected genes associated with dysrhythmia and aortic dilation resulted in further clinical evaluation and establishment of a surveillance plan.

Our understanding of the normal SV spectrum is only beginning to emerge from population-based LRS of individuals without a known condition.^7,28,29^ As a result, the pathogenicity of many variants remain uncertain. For example, in case S009 with X-linked Duchenne muscular dystrophy, the intronic AGAA repeat expansion is not only rare in population samples but also found almost exclusively in females. Whether this expansion perturbs the function of *DMD*, perhaps by blocking transcript elongation,^30^ acting as a novel transcription factor binding site,^31^ or inducing cellular death through a process such as RAN translation,^32^ remains to be determined. However, its low prevalence in males makes it a compelling candidate for further evaluation, and if determined to be pathogenic, a potential target for therapeutic intervention.^33^ We anticipate that more widespread application of T-LRS will lead to discovery of many more SVs of unknown significance. Assessment of pathogenicity of these variants will benefit from greater public sharing of SVs (e.g., establishment of a database, development of robust mechanisms for matching, etc.), as has been the case for SNVs and indels discovered by short-read exomes and genomes.^34,35^

We predict eventual implementation of LRS will have a major impact on all aspects of clinical genetic testing, because as a single test LRS has the potential to replace nearly every other genetic test currently offered. For example, in a person suspected to have a Mendelian condition, LRS data could first be used to evaluate sequence variants within a specific gene or genes. If no explanatory variant was found, the same dataset could reflexively be analyzed to interrogate sequence variants in all exons and high-priority noncoding regulatory regions, as well as search genome-wide for SVs and mutated repetitive elements. This testing strategy would replace the often-used stratified approach to testing (i.e., single gene testing, CMA, followed by whole-exome sequencing). Moreover, these steps are computationally applied to the same LRS data, so such a stepwise analysis could be completed in hours or days compared to weeks to months for conventional stratified testing strategies. Overall, this workflow is likely to increase the diagnostic rate, reduce the cost, and shorten the time to diagnosis for families with rare genetic conditions.

## ACKNOWLEDGEMENTS

We are grateful to the individuals and their families who participated in this study. We thank Angela Miller for figure preparation, Tonia Brown for assistance in editing this manuscript, and Sunday Stray and Lemlen Ghile from the laboratory of Mary-Claire King for isolation of DNA from whole blood for the SAGE BK samples. This work was supported by a grant to D.E.M. and E.E.E. from the Brotman Baty Institute for Precision Medicine. This work was also supported, in part, by grants from the US National Institutes of Health (NIH R01 MH101221 to E.E.E.) and the Simons Foundation (SFARI 608045 to E.E.E.). J.T.B is supported by NIH R01 HL130996-05 and BWF CAMS #1014700. R.A. and T.C. are supported by NIH R01 EY29315. Sequencing and analysis were supported in part by the University of Washington Center for Mendelian Genomics (UW-CMG) and funded by NHGRI and NHLBI grants UM1 HG006493 and U24 HG008956. H.L. and D.D. are supported by NIH R01HD100730 and U54HD083091. The content is solely the responsibility of the authors and does not necessarily represent the official views of the National Institutes of Health. E.E.E. is an investigator of the Howard Hughes Medical Institute.

## REFERENCES

1. Lowther C, Valkanas E, Giordano JL, et al. Systematic evaluation of genome sequencing as a first-tier diagnostic test for prenatal and pediatric disorders [Internet]. Cold Spring Harbor Laboratory. 2020 [cited 2020 Nov 1];2020.08.12.248526. Available from: https://www.biorxiv.org/content/10.1101/2020.08.12.248526v1

2. Boycott KM, Rath A, Chong JX, et al. International Cooperation to Enable the Diagnosis of All Rare Genetic Diseases. Am J Hum Genet 2017;100(5):695–705.

3. Frésard L, Montgomery SB. Diagnosing rare diseases after the exome. Cold Spring Harb Mol Case Stud [Internet] 2018;4(6). Available from: http://dx.doi.org/10.1101/mcs.a003392

4. Ewans LJ, Schofield D, Shrestha R, et al. Whole-exome sequencing reanalysis at 12 months boosts diagnosis and is cost-effective when applied early in Mendelian disorders. Genet Med 2018;20(12):1564–74.

5. Eichler EE. Genetic Variation, Comparative Genomics, and the Diagnosis of Disease. N Engl J Med 2019;381(1):64–74.

6. Logsdon GA, Vollger MR, Eichler EE. Long-read human genome sequencing and its applications. Nat Rev Genet 2020;21(10):597–614.

7. Chaisson MJP, Sanders AD, Zhao X, et al. Multi-platform discovery of haplotype-resolved structural variation in human genomes. Nat Commun 2019;10(1):1–16.

8. Gilpatrick T, Lee I, Graham JE, et al. Targeted nanopore sequencing with Cas9-guided adapter ligation. Nat Biotechnol 2020;38(4):433–8.

9. Karamitros T, Magiorkinis G. Multiplexed Targeted Sequencing for Oxford Nanopore MinION: A Detailed Library Preparation Procedure. In: Head SR, Ordoukhanian P, Salomon DR, editors. Next Generation Sequencing: Methods and Protocols. New York, NY: Springer New York; 2018. p. 43–51.

10. Payne A, Holmes N, Clarke T, Munro R, Debebe B, Loose M. Nanopore adaptive sequencing for mixed samples, whole exome capture and targeted panels [Internet]. 2020 [cited 2020 Aug 30];2020.02.03.926956. Available from: https://www.biorxiv.org/content/10.1101/2020.02.03.926956v2

11. Li H. Minimap2: pairwise alignment for nucleotide sequences. Bioinformatics 2018;34(18):3094–100.

12. Sedlazeck FJ, Rescheneder P, Smolka M, et al. Accurate detection of complex structural variations using single-molecule sequencing. Nat Methods 2018;15(6):461–8.

13. Edge P, Bansal V. Longshot enables accurate variant calling in diploid genomes from single-molecule long read sequencing. Nat Commun 2019;10(1):4660.

14. Luo R, Wong C-L, Wong Y-S, et al. Exploring the limit of using a deep neural network on pileup data for germline variant calling. Nature Machine Intelligence 2020;2(4):220–7.

15. McLaren W, Gil L, Hunt SE, et al. The Ensembl Variant Effect Predictor. Genome Biol 2016;17(1):122.

16. Rentzsch P, Witten D, Cooper GM, Shendure J, Kircher M. CADD: predicting the deleteriousness of variants throughout the human genome. Nucleic Acids Res 2019;47(D1):D886–94.

17. Jaganathan K, Kyriazopoulou Panagiotopoulou S, McRae JF, et al. Predicting Splicing from Primary Sequence with Deep Learning. Cell 2019;176(3):535–48.e24.

18. Killick R, Eckley I. changepoint:an R package for changepoint analysis. J Stat Softw 2014;58(3):19.

19. Heller D, Vingron M. SVIM: structural variant identification using mapped long reads. Bioinformatics 2019;35(17):2907–15.

20. Loman NJ, Quick J, Simpson JT. A complete bacterial genome assembled de novo using only nanopore sequencing data. Nat Methods 2015;12(8):733–5.

21. LaCroix AJ, Stabley D, Sahraoui R, et al. GGC Repeat Expansion and Exon 1 Methylation of XYLT1 Is a Common Pathogenic Variant in Baratela-Scott Syndrome. Am J Hum Genet 2019;104(1):35–44.

22. Fu YH, Kuhl DP, Pizzuti A, et al. Variation of the CGG repeat at the fragile X site results in genetic instability: resolution of the Sherman paradox. Cell 1991;67(6):1047–58.

23. Zhou Y, Kumari D, Sciascia N, Usdin K. CGG-repeat dynamics and FMR1 gene silencing in fragile X syndrome stem cells and stem cell-derived neurons. Mol Autism 2016;7:42.

24. Richards S, Aziz N, Bale S, et al. Standards and guidelines for the interpretation of sequence variants: a joint consensus recommendation of the American College of Medical Genetics and Genomics and the Association for Molecular Pathology. Genet Med 2015;17(5):405–24.

25. Sulovari A, Li R, Audano PA, et al. Human-specific tandem repeat expansion and differential gene expression during primate evolution. Proc Natl Acad Sci U S A 2019;116(46):23243–53.

26. Trost B, Engchuan W, Nguyen CM, et al. Genome-wide detection of tandem DNA repeats that are expanded in autism. Nature 2020;586(7827):80–6.

27. Dolzhenko E, Deshpande V, Schlesinger F, et al. ExpansionHunter: a sequence-graph-based tool to analyze variation in short tandem repeat regions. Bioinformatics 2019;35(22):4754–6.

28. Audano PA, Sulovari A, Graves-Lindsay TA, et al. Characterizing the Major Structural Variant Alleles of the Human Genome. Cell 2019;176(3):663–75.e19.

29. Beyter D, Ingimundardottir H, Eggertsson HP, et al. Long read sequencing of 1,817 Icelanders provides insight into the role of structural variants in human disease [Internet]. 2019 [cited 2020 Oct 15];848366. Available from: https://www.biorxiv.org/content/10.1101/848366v1

30. Punga T, Bühler M. Long intronic GAA repeats causing Friedreich ataxia impede transcription elongation. EMBO Mol Med 2010;2(4):120–9.

31. Bourque G, Leong B, Vega VB, et al. Evolution of the mammalian transcription factor binding repertoire via transposable elements. Genome Res 2008;18(11):1752–62.

32. Zu T, Gibbens B, Doty NS, et al. Non-ATG-initiated translation directed by microsatellite expansions. Proc Natl Acad Sci U S A 2011;108(1):260–5.

33. Levin AA. Treating Disease at the RNA Level with Oligonucleotides. N Engl J Med 2019;380(1):57–70.

34. Philippakis AA, Azzariti DR, Beltran S, et al. The Matchmaker Exchange: a platform for rare disease gene discovery. Hum Mutat 2015;36(10):915–21.

35. Karczewski KJ, Francioli LC, Tiao G, et al. The mutational constraint spectrum quantified from variation in 141,456 humans. Nature 2020;581(7809):434–43.

