## Supplementary Appendix for "Targeted long-read sequencing resolves complex structural variants and identifies missing disease-causing variants"

**Supplementary Methods** 5
- DNA isolation 5
- Library preparation, DNA sequencing, and target selection 5
- Basecalling, alignment, and candidate variant identification 5
- Depth of coverage calculations within target regions and genome-wide 6
- Generation of coverage plots 6
- Calculation of average read length within and outside of targeted regions 6
- Calculation of repeat lengths 6
- Refinement of copy number variant breakpoints using binary segmentation 6
- Southern blot of Baratela-Scott family 04 in Figure 1 7
- Estimating the size of reads spanning the KpnI cut sites in Baratela-Scott family 04 7
- PacBio CLR sequencing of Baratela-Scott family 04 7
- Confirmation of splice variant in patient S004 (NPHP4) 7
- Estimate of the size of the AGAA repeat within DMD in patient S009 8
- PCR of deletion breakpoints in patient S013 8
- HiFi sequencing of patient S020 and analysis for additional rearrangement breakpoints 8
- Phasing of patient S025 by linkage disequilibrium 9
- PCR of intronic variant in patient S056 9
- Data and software availability 9

**Supplementary Text** 10
- Clinical summary and prior testing for simple structural variant cases 10
- Clinical summary and prior testing for repeat expansion cases 10
- Clinical summary and prior testing for complex structural variant cases 10
- Clinical summary and prior testing for missing variant cases 11

**Supplementary Figures** 13
- Figure S1: Binary segmentation figures. 13
- Figure S2. BK144-03, known 22q13.3 deletion. 24
- Figure S3. BK180-03, known 15q11-13 duplication. 27
- Figure S4. BK294-03, known 22q11.2 duplication. 28
- Figure S5. BK364-03, known 1p36.11 duplication. 29
- Figure S6. BK397-101, known 16p11.2 deletion. 32
- Figure S7. BK430-103, known 16p11.2 duplication. 32
- Figure S8. BK482-101, known 1q21.1 duplication. 33
- Figure S9. BK487-101, known 1q21 deletion. 33
- Figure S10. BK506-03, known 5p15.33 deletion. 34
- Figure S11. S016, known tandem duplication within CTNND2. 37
- Figure S12. S046, known unbalanced translocation between chromosomes 4 and 15. 39

Figure S13. S011 (ATXN3 and ATXN8OS), evaluation of repeat length in patient sample. 41
Figure S14. S039 (FMR1), evaluation of repeat length and methylation status. 42
Figure S15. S040 (FXN), known repeat expansion. 43
Figure S16. S041 (FXN), known repeat expansion. 44
Figure S17. 04-01, 04-02, and 04-03 (XYLT1), evaluation of repeat length in family 04 from LaCroix et al. 2019. 45
Figure S18. 06-01, 06-02, and 06-03 (XYLT1), evaluation of repeat length in family 06 from LaCroix et al. 2019. 47
Figure S19. S014, three noncontiguous deletions of chromosome 6 identified by CMA. 48
Figure S20. S020, individual with three deletions identified on array and multiple rearrangements on karyotype. 56
Figure S21. S021, mosaic loss of 8p and mosaic gain of 8q. 86
Figure S22. S022, focal amplification of 4q with adjacent region of homozygosity, duplication of 15q11.2. 89
Figure S23. S023, mosaic ring 18 present in 40% of cells. 93
Figure S24. S035, duplications of 8q24 and 16p13.11 identified by clinical testing. 94
Figure S25. S036, individual with multiple rearrangements and translocations of chromosomes 5, 6, 10, and 18. 96
Figure S26. S002 (ALMS1), IGV views of known inherited stop variant and Alu insertion. 115
Figure S27. S003 (NPHP4), IGV views of inherited stop, splice variant, and data showing variant does affect splicing. 116
Figure S28. S004 (VARS2), IGV view of known inherited variant. 120
Figure S29. S008 (HPRT1), view of inversion and FISH results. 121
Figure S30. S009 (DMD), IGV view of AGAA expansion and frequency in SSC samples. 122
Figure S31. S013 (HPS1), IGV view of inherited variant and deletion identified by LRS. 124
Figure S32. S018 (PAH), known inherited splice variant identified, no second variant found. 126
Figure S33. S025 (ABCA4), the previously known variant and a 1,500 bp insertion can be phased into different haplotypes. 127
Figure S34. S056 (WDR19), IGV views of inherited and splice variants identified by LRS. 131

**Supplementary Tables** 132
Table S1: Sample summary, DNA source, flow cells and libraries used per sample. 132
Table S2: Per sample sequencing targets, coverage, and average read length. 133
Table S3: Overview of individuals with a single structural variant. 134
Table S4: Overview of individuals with repeat expansions, including expected and observed repeat expansion sizes. 135
Table S5: Per-read details of individuals with ATXN3, ATXN8OS, FMR1, and FXN repeat expansions. 136
Table S6: Per-read details of XYLT1 repeat expansions. 137
Table S7: Summary of individuals with complex SVs, including previously known events and new events identified by T-LRS. 138
Table S8: Known and new events observed for individuals with known complex SVs. 138
Table S9: Details of focal amplification of 4q in patient S022. 139

Table S10: Rearrangement numbers and sizes for patient S014. 139
Table S11: Genes impacted by S014 breakpoints. 139
Table S12: Rearrangement numbers and sizes for patient S020. 140
Table S13: Genes impacted by S020 breakpoints. 141
Table S14: Rearrangement numbers and sizes for patient S036. 142
Table S15: Genes impacted by S036 breakpoints. 142
Table S16: Summary of individuals with missing variants. 143
Table S17: Reads used to calculate length of AGAA motif in patient S009. 144
Table S18: Predicted strength of the canonical splice donor site at the Exon 1–Intron 1 boundary of ABCA4 (NM_000350) and alternative sites introduced by the ~1,500 bp insertion in patient S025. 145

**Supplementary Appendix References** 146

### Authors and affiliations

Danny E. Miller, MD, PhD[1,2,*], Arvis Sulovari, PhD[1,&], Tianyun Wang, PhD[1,&], Hailey Loucks, BS[2], Kendra Hoekzema, MS[1], Katherine M. Munson, BA[1], Alexandra P. Lewis, BS[1], Edith P. Almanza Fuerte, BS[2], Catherine R. Paschal, PhD[3,4], Jenny Thies, MS, CGC[2], James T. Bennett, MD, PhD[2,3,5,6], Ian Glass, MB ChB, MD[2], Katrina M. Dipple, MD, PhD[2,6,7], Karynne Patterson, BS[1], Emily S. Bonkowski, MS, CGC[2], Zoe Nelson, MS, CGC[2], Audrey Squire, MS, CGC[2], Megan Sikes, MS, CGC[2], Erika Beckman, MS, CGC[2], Robin L. Bennett, MS, CGC[8], Dawn Earl, ARNP[2], Winston Lee, MA[9,10], Rando Allikmets, PhD[10,11], Seth J. Perlman, MD[12], Penny Chow, MS, CGC[13], Anne V. Hing, MD[13], Margaret P. Adam, MD[2], Angela Sun, MD[2], Christina Lam, MD[2,8,14], Irene Chang, MD[2], University of Washington Center for Mendelian Genomics, Tim Cherry, PhD[5], Jessica X. Chong, PhD[2,6], Michael J. Bamshad, MD[1,2,6], Deborah A. Nickerson, PhD[1,6], Heather C. Mefford, MD, PhD[2,6], Dan Doherty, MD, PhD[2,6,15], Evan E. Eichler, PhD[1,6,16,*]

[1] Department of Genome Sciences, University of Washington School of Medicine, Seattle, WA, 98195, USA
[2] Department of Pediatrics, Division of Genetic Medicine, University of Washington and Seattle Children's Hospital, Seattle, WA, 98105, USA
[3] Department of Laboratories, Seattle Children's Hospital, Seattle, WA, 98105, USA
[4] Department of Laboratory Medicine and Pathology, University of Washington, Seattle, WA, 98195, USA
[5] Center for Developmental Biology and Regenerative Medicine, Seattle Children's Research Institute, Seattle, WA, 98101, USA
[6] Brotman Baty Institute for Precision Medicine, Seattle, WA, 98195, USA
[7] Center for Clinical and Translational Research, Seattle Children's Research Institute, Seattle, WA, 98101, USA
[8] Division of Medical Genetics, Department of Medicine, University of Washington, Seattle, WA, 98195, USA
[9] Department of Genetics and Development, Columbia University, New York, NY 10032, USA
[10] Department of Ophthalmology, Columbia University, New York, NY 10032, USA
[11] Department of Pathology and Cell Biology, Columbia University, New York, NY 10032, USA
[12] Department of Neurology, Seattle Children's Hospital, University of Washington, Seattle, WA, 98105, USA
[13] Department of Pediatrics, Division of Craniofacial Medicine, University of Washington, Seattle, WA, 98195, USA
[14] Center for Integrative Brain Research, Seattle Children's Research Institute, Seattle, WA, 98101, USA
[15] Department of Pediatrics, Division of Developmental Medicine, University of Washington and Seattle Children's Hospital, Seattle, WA, 98105, USA
[16] Howard Hughes Medical Institute, University of Washington, Seattle, WA, 98195, USA

[&] Both authors contributed equally (second authors), *Corresponding authors

Evan E. Eichler, Ph.D.
Department of Genome Sciences
University of Washington School of Medicine
3720 15th Ave NE, S413A
Box 355065
Seattle, WA 98195-5065


Danny E. Miller, M.D., Ph.D.
Department of Pediatrics, Division of Genetic Medicine
Seattle Children's Hospital
4800 Sand Point Way NE, OC.7.830
Seattle, WA 98105-0371


### Supplementary Methods

#### DNA isolation

DNA was isolated from blood, saliva, or fibroblasts using one of the following methods (Table S1). Saliva was collected in an Oragene DNA collection kit (DNA Genotek) and 2-4 mL of saliva plus stabilizing fluid was incubated for 1-16 hours in a water bath at 50°C. 40 uL per 1 mL of Oragene purifier (prepIT-L2P) was added and mixed well, followed by a 10-minute incubation on ice. The protein precipitate was pelleted by centrifugation (10 minutes, 3500 rpm) and the clear supernatant was transferred to a new tube. An equal volume of 100% EtOH was added to precipitate DNA, which was then pelleted and washed twice with 70% ethanol. DNA was rehydrated with Elution Buffer and left for 2 days to resuspend. Further cleanup with a 0.45x bead wash was required for some saliva samples.

DNA was extracted from whole blood by separating white blood cells (WBCs) using the Puregene Red Blood Cell Lysis solution (QIAGEN). Four cycles of washing were done where three volumes of red blood cell (RBC) lysis solution was mixed with the sample then pelleted. After the final wash WBCs were suspended in cell lysis solution (QIAGEN) and incubated with RNase A at 37°C for 40 min. Protein Precipitation Solution (QIAGEN) was added at 0.33x and mixed well. After a 10 min incubation on ice, the precipitate was pelleted and the supernatant was transferred to a new tube. DNA was precipitated with equal volumes of isopropanol and then washed three times with 70% ethanol. DNA was resuspended in DNA Hydration Solution (QIAGEN) for at least 2 days prior to use. DNA was extracted from fibroblasts or cell lines using the Gentra Puregene Cell Kit (QIAGEN) following the manufacturer's instructions.

#### Library preparation, DNA sequencing, and target selection

Extracted DNA was quantified and sheared to a target fragment size of 8-12 kbp using a Covaris g-TUBE. Approximately 1.5 ug of sheared DNA was used to make sequencing libraries using the Oxford Nanopore Ligation Sequencing Kit (SQK-LSK109) following the manufacturer's instructions, except that for each library the short fragment buffer (SFB) was used during cleanup, and all elutions were done for 10 minutes at 37°C. All 15 ul of each library was loaded onto a release 9.4.1 flow cell for sequencing on an Oxford Nanopore GridION. Target regions were enriched using Read Until (Payne et al. 2020). Read Until was run with guppy 3.4.5 and configured to use the dna_r9.4.1_450bp_fast model with min_chunks = 0 and max_chunks = 12. The sequencing_MIN106_DNA file was modified to set break_reads_after_seconds = 0.4. For each experiment, at least 100 kbp and up to several megabases on either side of the gene or region of interest were targeted (Table S2). Sequencing experiments were run for up to 72 hours and, in some cases, a second DNA library was loaded onto the same flow cell after washing at approximately 24 hours into a sequencing experiment in order to increase output (Table S1).

#### Basecalling, alignment, and candidate variant identification

FASTQ files were generated using guppy 4.0.11 and aligned to GRCh38 using both minimap2 (version 2.17-r941) (Li 2018) and NGMLR (version 0.2.7) (Sedlazeck et al. 2018) with default parameters. Single-nucleotide variants (SNVs) were called using Longshot (Edge and Bansal 2019) from both minimap2 and NGMLR alignments and both SNVs and insertion/deletion (indel) variants were called using clair (Luo et al. 2020) and medaka (version 1.0.3) (https://github.com/nanoporetech/medaka) from both minimap2 and NGMLR alignments. VCF files that combined all variant calls were annotated with global allele frequency, variant effect predictions (McLaren et al. 2016) from gnomAD v2 liftover files (Karczewski et al. 2020), and CADD version 1.6 scores (Rentzsch et al. 2019) using vcfAnno (Pedersen, Layer, and Quinlan 2016). Novel intronic SNVs or those with allele frequencies <2% were passed to SpliceAI (Jaganathan et al. 2019) for analysis. Variants for analysis were filtered based on allele frequency, CADD score, and SpliceAI prediction. Variants were phased using Longshot. Structural variants (SVs) were identified using both Sniffles (Sedlazeck et al. 2018) and SVIM (Heller and Vingron 2019) on both minimap2 and NGMLR alignments. Only those SVs supported by four or more reads within the regions targeted for sequencing were analyzed. For cases in which CpG methylation was assayed, methylation changes were identified within target regions using Nanopolish (Loman, Quick, and Simpson 2015) and BAM files were subsequently converted for visual analysis using Nanopore methylation utilities (Lee et al. 2018).

#### Depth of coverage calculations within target regions and genome-wide

Coverage of target regions and genome-wide coverage was calculated using SAMtools depth with the -a and -Q 0 flag, which calculates coverage using reads with quality scores of 0 or above (Li et al. 2009).

#### Generation of coverage plots

Data for coverage plots was generated using SAMtools depth with the -a and -Q 0 flags, a custom script then calculated the average coverage in 1 kbp nonoverlapping windows. Plots were generated using average coverage in karyoploteR (Gel and Serra 2017).

#### Calculation of average read length within and outside of targeted regions

Average read length both genome-wide and within target regions (Table S2) was calculated using a custom script. Briefly, the average length of all reads in all FASTQ files from a sample was used to calculate the genome-wide average read length. To calculate the average length of reads within target regions, SAMtools was used to isolate reads that mapped to the target region. Read IDs were then extracted and the length of the read in the FASTQ file was calculated. Each read ID was counted once. Because two flow cells with two different target regions were run for samples S020 and S036, the genome-wide read length was calculated using reads separated by experiment.

#### Calculation of repeat lengths

To estimate the size of repeats we analyzed regions within *FMR1*, *ATXN3*, and *ATXN8OS* identified as tandem repeats by Tandem Repeats Finder (Benson 1999) using sensitive parameter settings to maximize tandem repeat discovery despite potential sequence errors in the ONT reads: trf dna_sequence.fa 2 7 7 80 10 20 50 -h -d. For *FXN* we defined the repeat window as the region of the reference genome containing the GAA repeat and for *XYLT1* we used the position given in (LaCroix et al. 2019). All targets can be found in Table S4. We used a reference-guided approach to estimating the size of the repeat length. Prior to analysis we re-aligned reads to GRCh38 (without alternative contigs) using minimap2 with the -r 50000, -end-bonus 10000, and --no-end-flt options to optimize the number of reads that spanned the repeats. This reduced the number of reads split by the aligner (Figures S13–S18). A custom script was then used to identify reads that spanned the target region plus a variable number of repeats that depended on the quality of the alignment (given in Table S4). For each read, the CIGAR string was then parsed to determine the length of the sequence that spanned the interval and the length of the additional sequence analyzed was subtracted from the length to get the estimated repeat size. The supplemental alignment of read b0a508ce-069d-43ac-865e-7b7cd900eb70 in sample 04-02 was manually removed leaving 16 reads remaining for that sample. Repeats were then grouped by their length and the average was calculated (Tables S5, S6).

#### Refinement of copy number variant breakpoints using binary segmentation

We used sequence depth information from the ONT reads to refine the CNV breakpoints. Specifically, we processed the read-depth information through a binary segmentation, implemented in the R package changepoint (Killick and Eckley 2014). The function cpt.meanvar() was used, which considers both mean and variance of sequencing depth to identify the transition points in the data (i.e., point of sudden increase or decrease in depth). The Bayesian Information Criterion was used to identify the best fit for the optimal regions of distinct depth profiles. This approach helped us refine coordinates for deletions and duplications. All analyses were done using R.3.6.1 and the scripts used for breakpoint refinement are publicly available on GitHub (see data sharing statement below). Results are in Figure S1.

#### Southern blot of Baratela-Scott family 04 in Figure 1

Southern blot was performed using standard methods as previously described (LaCroix et al. 2019). DNA was digested with KpnI restriction enzyme (New England Biolabs), followed by electrophoresis (0.8% agarose), overnight capillary transfer of the separated DNA fragments via charged nylon membrane (GE Amersham), and crosslinking by exposure to ultraviolet light. The probe (chr16:17,563,659-17,564,191, GRCh37/hg19) was prepared by PCR amplification, cloned into a plasmid, labeled with p32-alpha-dCTP (MegaPrime), and hybridized to the membrane at 6°C overnight. The membrane was washed two times for 15 min each time in 2x SSC, 0.1% SDS and once with 0.2x SSC, 1% SDS at 6°C. Probes were exposed to film for 6 days at -80°C before development.

#### Estimating the size of reads spanning the *KpnI* cut sites in Baratela-Scott family 04

The number of base pairs between *KpnI* cut sites in Baratela-Scott family 04 (Figure 1; Tables S4, S6) was estimated by first determining the genomic position of both KpnI sites by computationally digesting 5 kbp of reference genome using restriction analyzer (http://www.molbiotools.com/restrictionanalyzer.html). This resulted in a 2,589 bp fragment that aligned to chr16:17468735-17471324 using BLAT (GRCh38 coordinates). A custom script was then used to extract reads from the minimap2 assembly that spanned a 500 or 50 bp interval around the repeat expansion site (Table S4) and counted the total number of nucleotides within that interval by parsing the CIGAR string. All reads spanned the complete interval between the two KpnI sites. The length of the read in the targeted interval was then reduced by the additional target space (either 500 or 50 bp) and 2,589 bases were added to this value, which represented the difference between the length of the interval within the KpnI cut sites and the 1 bp interval that was targeted for counting.

#### PacBio CLR sequencing of Baratela-Scott family 04

PacBio CLR libraries were generated according to manufacturer's instructions and as described in (Chaisson et al. 2019) with some modifications. Briefly, high-molecular-weight DNA was sheared using Megaruptor (Diagenode) using the 50 kbp setting. After adapter ligation with the SMRTbell Express Template Prep Kit, samples were size-selected on a BluePippin instrument using a high-pass cutoff of 35 kbp or 40 kbp resulting in average library sizes (measured with FEMTO Pulse) of 61 and 72 kbp, respectively. Each library was loaded on three SMRT Cell 1Ms on the Sequel platform using v3 chemistry with 10-hour movie times. Final data yield was 32 Gbp Reads of Insert (ROI) (10x coverage) for 38-2 and 38 Gbp ROI (12x coverage) for 38-4, with mean subread lengths of 23 kbp and N50 subread read lengths of 40 kbp.

#### Confirmation of splice variant in patient S004 (*NPHP4*)

To validate that the splice variant in S004 indeed affected splicing, we assayed for a 50 bp insertion between *NPHP4* exons 5 and 6 with PCR of cDNA from fibroblasts using two primer pairs. The first pair flanks the exon junction (forward: CTCCTGCACCCGCTTCTC and reverse: GGATTCTCCATGAGCTGGAA), the second pair uses the same reverse primer but the forward primer (CAGCACTCACTGCTCTCGTG) falls within the expected 50 bp insertion of intron 5-6. RNA was extracted using the Aurum Total RNA Kit (Bio-Rad) with a spin-mediated protocol. cDNA was synthesized using iScript cDNA Synthesis Kit (Bio-Rad). PCR was performed on the cDNA using 15 µl 2x Failsafe

PCR Buffer J (2X) (Lucigen), 5.6 µl of water, 2.5 µl of 10 µM forward and reverse primer mix, 2 µl of 50 ng/µl template, and 0.4 µl Platinum Taq (5 U/µl) per reaction. A touchdown cycling protocol was used: the first 10 cycles had a variable annealing temperature from 65-56°C, and the next 25 cycles had an annealing temperature of 55°C, for a total of 35 cycles. Extension time was 30 seconds per cycle. Bands were then excised and extracted using Monarch DNA gel extraction Kit (New England Biolabs). For the first primer pair, two bands were seen which were extracted for separate sequencing. Due to low yield after gel extraction, the extracted bands were run again using the same PCR protocol and primers, then un-purified PCR products or column-purified products (Monarch PCR & DNA Cleanup Kit, New England Biolabs) were submitted to Genewiz with the reverse primer for Sanger sequencing (Figure S28D).

For validation of segregation in S004, PCR was performed using the above protocol with same temperature and extension time on DNA samples from the proband and parents. Proband DNA was extracted using Gentra Puregene Cell kit (QIAGEN) from fibroblasts, and for parents the prepIT-L2P reagent (DNA Genotek Inc.) was used to extract from saliva. Primers used were forward (TTGAGAACCACTGCTCCAGA) and reverse (ACGAAACATCTGCCAAAACC). Un-purified PCR products or column-purified PCR products were submitted to Genewiz for Sanger sequencing using the forward primer, and the splice variant was confirmed to be maternally inherited.

#### Estimate of the size of the AGAA repeat within *DMD* in patient S009

Reads spanning the interval chrX:32,554,300-32,555,300 were isolated and used to estimate the number of AGAA repeats using Tandem Repeats Finder within *DMD*. Nine reads were used to estimate the number of AGAA repeats within the interval; three reads were excluded because the repeat in those reads contained a mix of AGAA and TGTT repeats (Table S17).

#### PCR of deletion breakpoints in patient S013

The ~1,900 bp deletion breakpoint in sample S013 was validated by PCR. Briefly, 12.5 µl of Roche FastStart PCR Master mix was combined with 10.5 µl of water, 1 µl of genomic DNA, and 1 µl each of forward (CCCCTTAGAGCAGAAAGGGAC) and reverse (TCATTACCTGACACCCGCAC) primers. PCR was run at an annealing temperature of 55°C for a total of 35 cycles with an extension time of 2 minutes.

#### HiFi sequencing of patient S020 and analysis for additional rearrangement breakpoints

A PacBio HiFi library was generated as in (Wenger et al. 2019) with the following modifications: high-molecular-weight DNA was sheared using g-TUBE (Covaris) to a mode size of 26 kbp. After adapter ligation with the SMRTbell Express Template Prep Kit 2.0 and removal of imperfect SMRTbells with the Enzyme Clean Up Kit, the library was size-fractionated on a SageELF platform (Sage Science) using the 1-18 kbp protocol and the fraction's size was measured on a FEMTO Pulse instrument (Agilent) and quantified with the Qubit dsDNA HS (High Sensitivity) Assay Kit (ThermoFisher). A fraction with a roughly 22 kbp average size was sequenced on one SMRT Cell 8M on a Sequel II instrument (PacBio) using version 2.0 bind and sequencing chemistry, with 4-hour pre-extension and 30-hour movie time. CCS analysis was performed through SMRT Link v9.0 with default settings (3 full passes, estimated quality 0.99) except the maximum read-length cutoff was extended to 100 kbp. Final data yield was 12.2 Gbp of sequence (~4X coverage) with an average length of 21.6 kbp and median estimated quality (Phred scaled) of Q28. Reads were aligned to GRCh38 and SVs were detected as described in (Audano et al. 2019). We searched for genome-wide translocations or rearrangements missed by T-LRS by filtering out BND variants overlapping a segmental duplication, near a reference gap, or near a contig end. Variants that passed this filter were visually evaluated with IGV and none identified were missed by T-LRS.

### Phasing of patient S025 by linkage disequilibrium

Using physical phasing information from the ONT reads that span the 1,450 bp insertion, we determined that a nearby SNV (rs2184339, T>C) had its alternative allele, C, on the same haplotype as reads with the insertion. Using the 1000 Genomes Project SNV genotypes, we calculated linkage disequilibrium between rs2184339 and the missense mutation rs61750120 (G>A) using the R^2 and D' statistics (Machiela and Chanock 2015). Among 2,504 total unrelated individuals representing 26 world populations (1000 Genomes Project Consortium et al. 2015), we observed no haplotypes containing both the C and A alleles (D'=0, and R^2=0.0002) (Figure S33). These observations suggest that the insertion allele within intron 1 of *ABCA4* and the missense allele in exon 22 reside on different haplotypes.

### PCR of intronic variant in patient S056

Sanger sequencing was performed to validate the *WDR19* intronic variant. Parental DNA was extracted from saliva using the prepIT-L2P reagent (DNA Genotek Inc.). The same PCR protocol described above for S004 was used, with the forward (CTCCTCCCCATCACCTTTC) and reverse (ACATCCTTGCTTCCTGACCA) primers. The forward primer was used for Sanger sequencing (Genewiz).

### Data and software availability

Scripts are available on GitHub at https://github.com/danrdanny/targetedLongReadSequencing and https://github.com/asulovar/ONT_binary_segmentation. Sequencing data for all cases are pending submission to dbGaP with accession numbers forthcoming. Please contact corresponding author D.E.M. for further information.

### Supplementary Text

#### Clinical summary and prior testing for simple structural variant cases

All nine cases beginning with "BK" are from the Study of Autism Genetics Exploration (SAGE) collection, five cases of which have been previously described (Guo et al. 2019) and four (BK397-101, BK430-103, BK482-101, BK487-101) are samples collected after the publication.

Patient S016 has a known duplication in *CTNND2* confirmed by clinical mate-pair sequencing to be tandem (Miller, Squire, and Bennett 2020).

Patient S046 presented with developmental delay and was found to have an unbalanced translocation between chromosomes 4 and 15. A single-nucleotide polymorphism (SNP) array identified a copy state of 1 at 4q35.1q35.2 and a copy state of 3 at 15q26.1q26.3.

#### Clinical summary and prior testing for repeat expansion cases

Patient S011 was an individual with clinically confirmed expansions in *ATXN3* (74 and 28 repeats) and *ATXN8OS* (80 and 25 repeats). Patient 04-01 (proband), 04-02 (mother), 04-03 (father), 06-01 (proband), 06-02 (mother), and 06-03 (father) are described in (LaCroix et al. 2019).

#### Clinical summary and prior testing for complex structural variant cases

Patient S014 presented prenatally with agenesis of the corpus callosum and ventriculomegaly. At birth the child was noted to have mild dysmorphic features and pelviectasis without hydroureter. A SNP array identified a complex pathogenic heterozygous deletion within 6q25.2 to 6q25.3, which included *ARID1B* and perhaps explained most of his clinical findings.

Patient S020 presented with hypotonia, developmental delay, epilepsy, and dysmorphic features. A SNP array revealed deletions of 4q13.2q13.3, 4q13.3, and 14q11.2. Karyotyping identified rearrangements among chromosomes 2, 14, 10, and 4 that involved the deleted regions and also 2p23, 2p25, 10q21.2, 10q21.1, and 10q22.3.

Patient S021 presented with developmental delay and was found by microarray and karyotype to be mosaic for complex changes to chromosome 8 that included loss of 8p23.2–pter (copy number 1), mosaic loss of a segment within 8p23.2–8p23.1 (copy number 1–2), and mosaic gain of 8q22.1–qter (copy number 2–3). Karyotype identified one cell line with a derivative chromosome 8 that consists of a terminal duplication of 8q, from 8q22.1 to qter, and a terminal deletion of 8p, from 8p23.2 to pter. The duplicated region of 8q is present on distal 8p, such that 8q22.1 to qter is present at both ends of the derivative chromosome 8. The other cell line has a terminal deletion of 8p, from 8p23.1 to pter.

Patient S022 presented with hemihypertrophy of unclear etiology. A standard Beckwith-Wiedemann workup including *CDKN1C* sequencing and deletion/duplication analysis was unremarkable. A SNP array identified a focal amplification of 4q with the copy state reported as more than 4 with an adjacent 27.5 Mbp region of homozygosity on 4q. A SNP array also identified a duplication of 15q11.2. Metaphase and interphase FISH confirmed that the focal amplification of 4q was more than 4.

Patient S023 presented with developmental delay, epilepsy, and hypothyroidism. A SNP array identified a mosaic region of 18p11.32 and 18q21.31q23 with copy number between 1 and 2. Evaluation of chromosomes revealed that this patient was mosaic for a ring chromosome 18 in ~40% of cells.

Patient S035 presented with developmental delay and had duplications of 8q24.3 and 16p13.11.

Patient S036 underwent genetic testing for expressive language delay, microcephaly, and mild dysmorphic features. A SNP array revealed four noncontiguous deletions of chromosome 10 at 10p12.2p12.1, 10p11.21, and 10q21.1 (two deletions in this interval). Karyotype revealed a translocation between chromosomes 6 and 18 at 6q22.2 and 18p11.2 as well as a pericentric inversion of chromosome 10 at 10p11.2q11.2.

#### Clinical summary and prior testing for missing variant cases

Patient S002 presented with early-onset obesity, type 2 diabetes, cone-rod dystrophy, and sensorineural hearing loss. Clinical testing by SNP array was unremarkable and exome sequencing identified a single paternally inherited stop-gain variant in *ALMS1*, the gene associated with Alström syndrome, a recessive disorder that fit the phenotype well (Paisey et al. 2003). Subsequent exon-level array revealed no deletions or duplications in the gene.

Patient S003 was an individual with renal failure, retinal degeneration, and essential tremor. A SNP array was unremarkable, trio exome revealed a single variant in *NPHP4*, and deletion/duplication analysis of NPHP4 was unremarkable. Clinical RNA testing was sent, which was interpreted as indeterminate.

Patient S004 presented with agenesis of the corpus callosum, microcephaly, poor growth, lactic acidosis, and global developmental delay. The SNP array was unremarkable, trio exome sequencing revealed a single pathogenic variant in *VARS2*, and deletion/duplication analysis of the gene was unremarkable.

Patient S008 was an individual with biochemically confirmed Lesch-Nyhan syndrome. Clinical testing included karyotype and sequencing of exons on an exome backbone, both of which were unremarkable.

Patient S009 presented with concern for Duchenne muscular dystrophy due to a family history of the disease. The proband's maternal uncle passed away at age 29 from the disease without a molecular diagnosis. Molecular testing in the proband included SNP array, targeted exon sequencing, and deletion/duplication analysis, all of which were unremarkable. Analysis of muscle biopsy by immunohistochemistry revealed staining indicative of a dystrophinopathy, but dystrophin 1 antibody staining (rod domain) was more than classically seen in Duchenne type dystrophy.

Patient S013 presented with oculocutaneous albinism and platelet dysfunction, suggesting Hermansky-Pudlak syndrome, but SNP array and exome sequencing only identified a single paternally inherited stop in *HPS1*, one of several genes associated with this recessive disorder (Huizing et al. 2000).

Patient S018 presented with elevated phenylalanine consistent with phenylketonuria. Panel testing revealed a single pathogenic variant in *PAH*.

Patient S025 presented in their early twenties with disease characteristics in the macula of both eyes consistent with recessive Stargardt disease. No other systemic issues or family history was reported. Research sequencing revealed a single inherited pathogenic variant and research whole-genome short-read sequencing failed to identify a second hit. After identification of a 1,500 bp insertion in the first intron of *ABCA4* with LRS visual reanalysis of the short-read data confirmed an 11 bp target site duplication at the same position.

Patient S056 presented with end-stage renal disease related to nephronophthisis, rhizomelic/metaphyseal skeletal dysplasia, retinal dystrophy, developmental delays, hyperparathyroidism, and hepatic fibrosis. A ciliopathy gene panel revealed a single heterozygous pathogenic variant in *WDR19* and deletion/duplication analysis of the gene was unremarkable.

### Supplementary Figures

#### Figure S1: Binary segmentation figures.

To complement existing SV callers and independently guide the refinement of CNV breakpoints, we applied the binary segmentation method to the sequencing depth profiles of each CNV region. The coordinates of each region were further refined through visual validation. Due to the sensitivity of the binary segmentation method to duplicated genomic sequences, 6 of the 26 regions could not have their breakpoints refined by this method; these were refined using a combination of existing SV callers and visual refinement of sequencing depth profile.

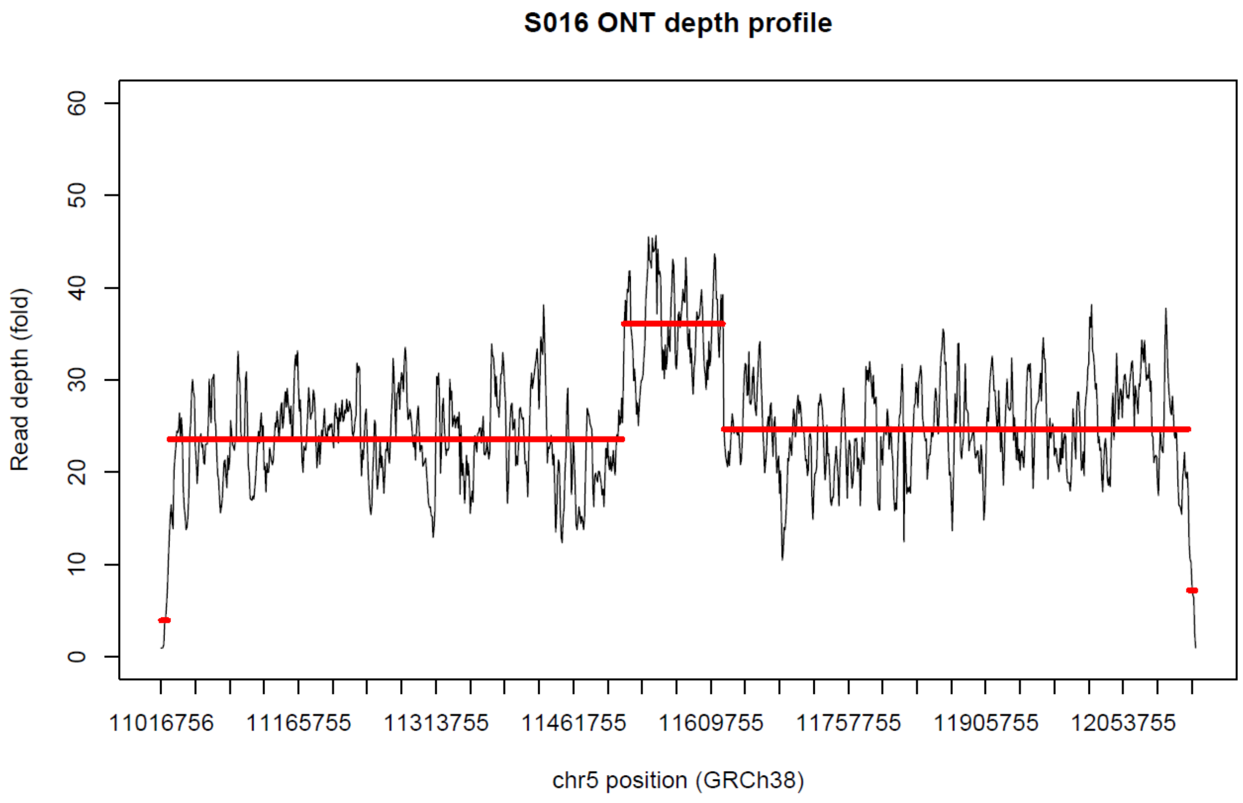

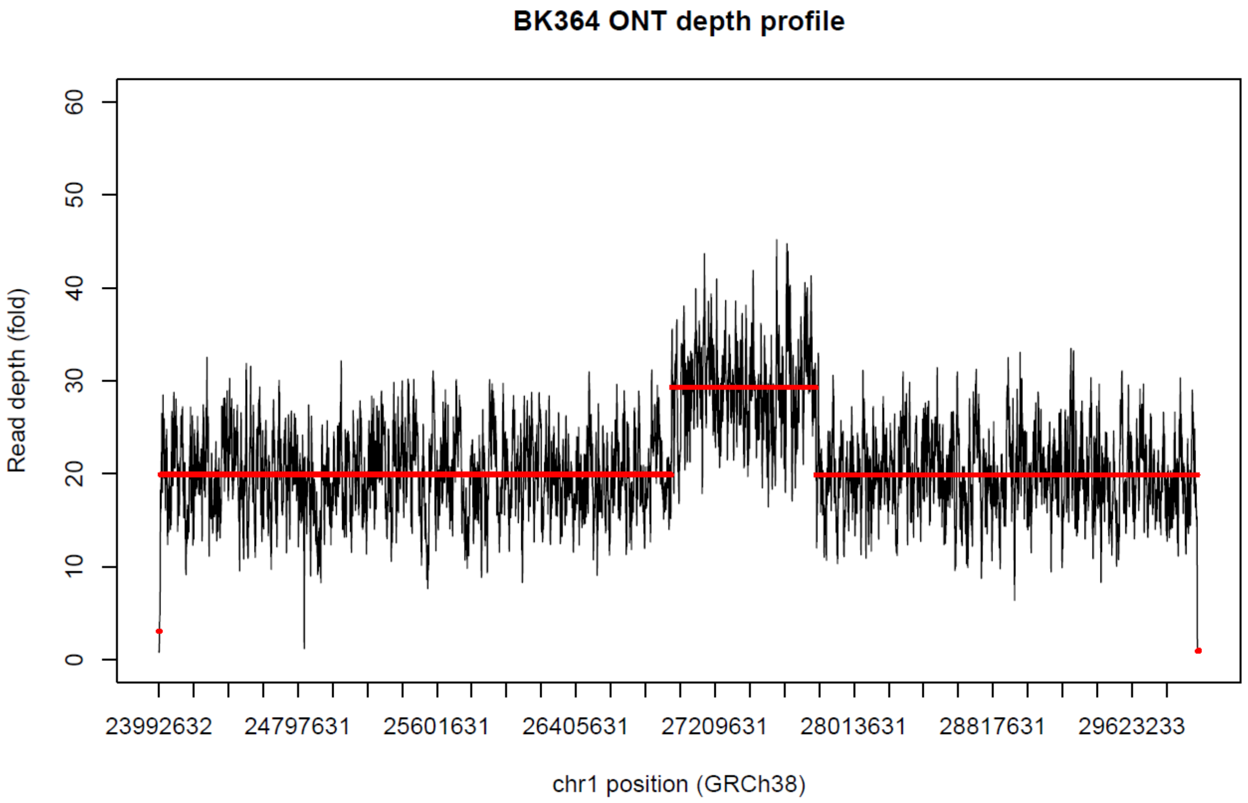
BK364 ONT depth profile
Read depth (fold)
0
10
20
30
40
50
60
23992632
24797631
25601631
26405631
27209631
28013631
28817631
29623233
chr1 position (GRCh38)

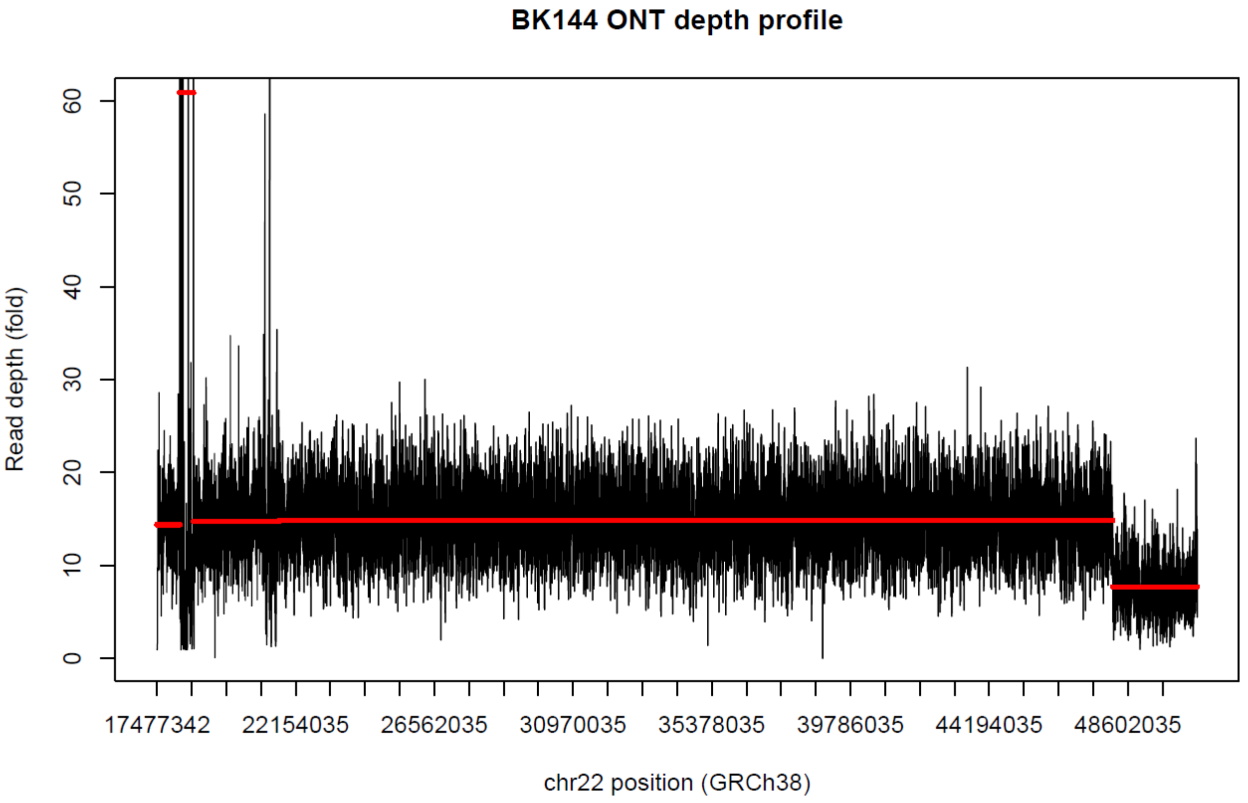
BK144 ONT depth profile
Read depth (fold)
0
10
20
30
40
50
60
17477342
22154035
26562035
30970035
35378035
39786035
44194035
48602035
chr22 position (GRCh38)

**BK397 ONT depth profile**

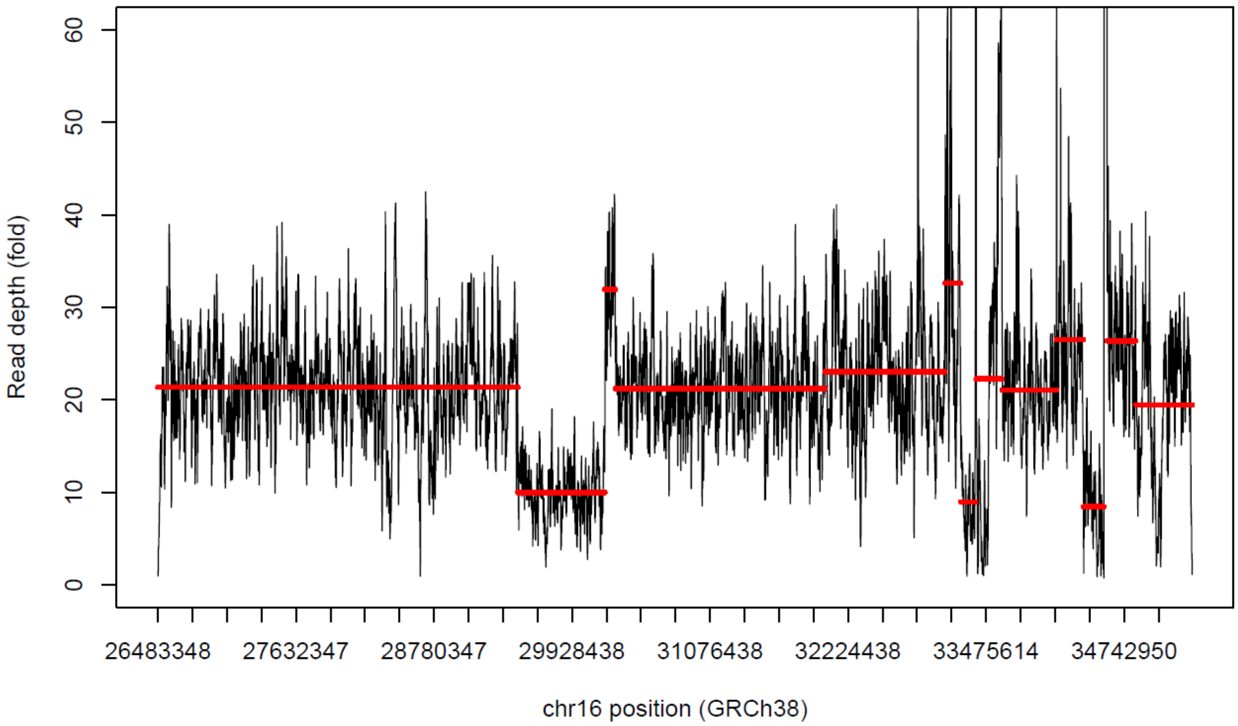

**BK430 ONT depth profile**

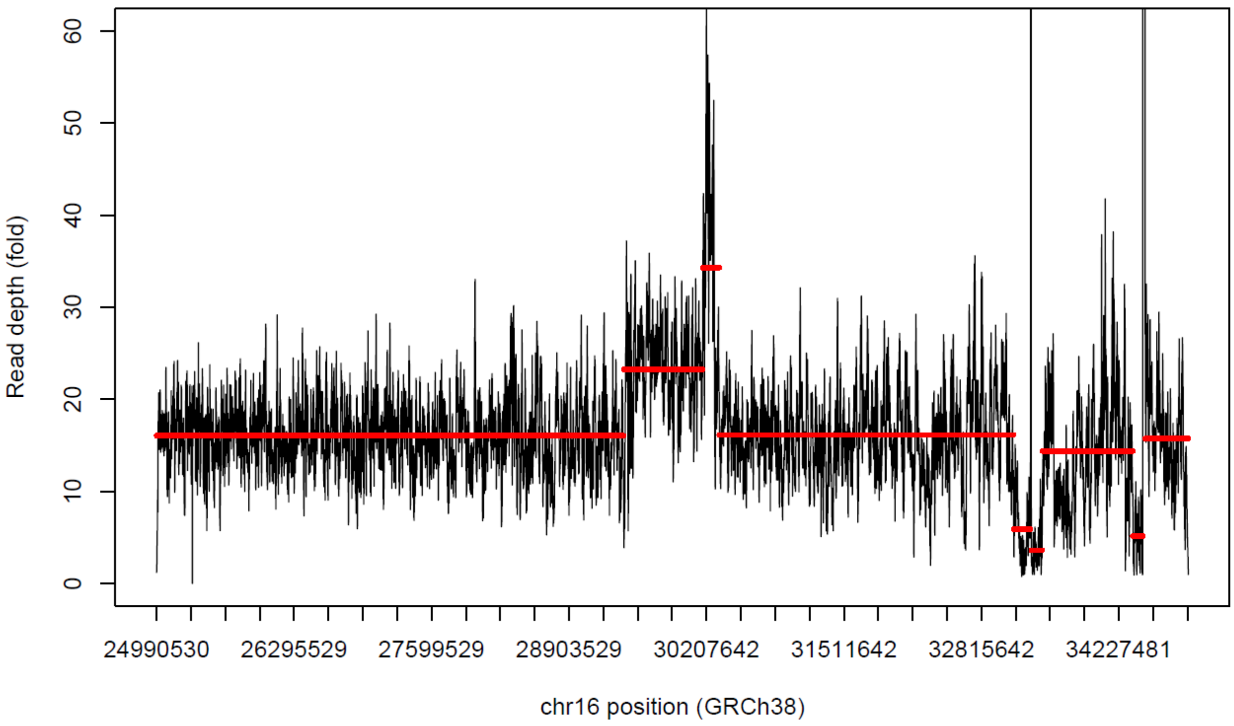

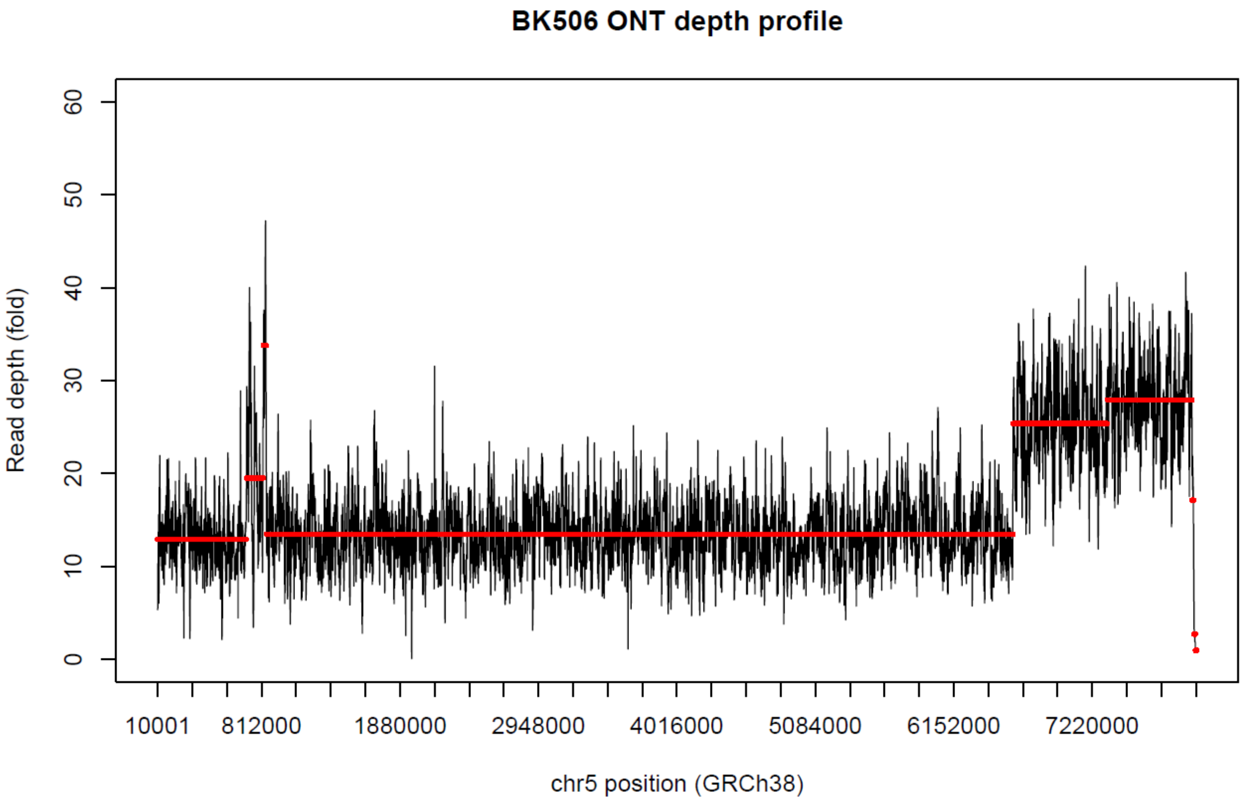
BK506 ONT depth profile
Read depth (fold)
0
10
20
30
40
50
60
10001
812000
1880000
2948000
4016000
5084000
6152000
7220000
chr5 position (GRCh38)

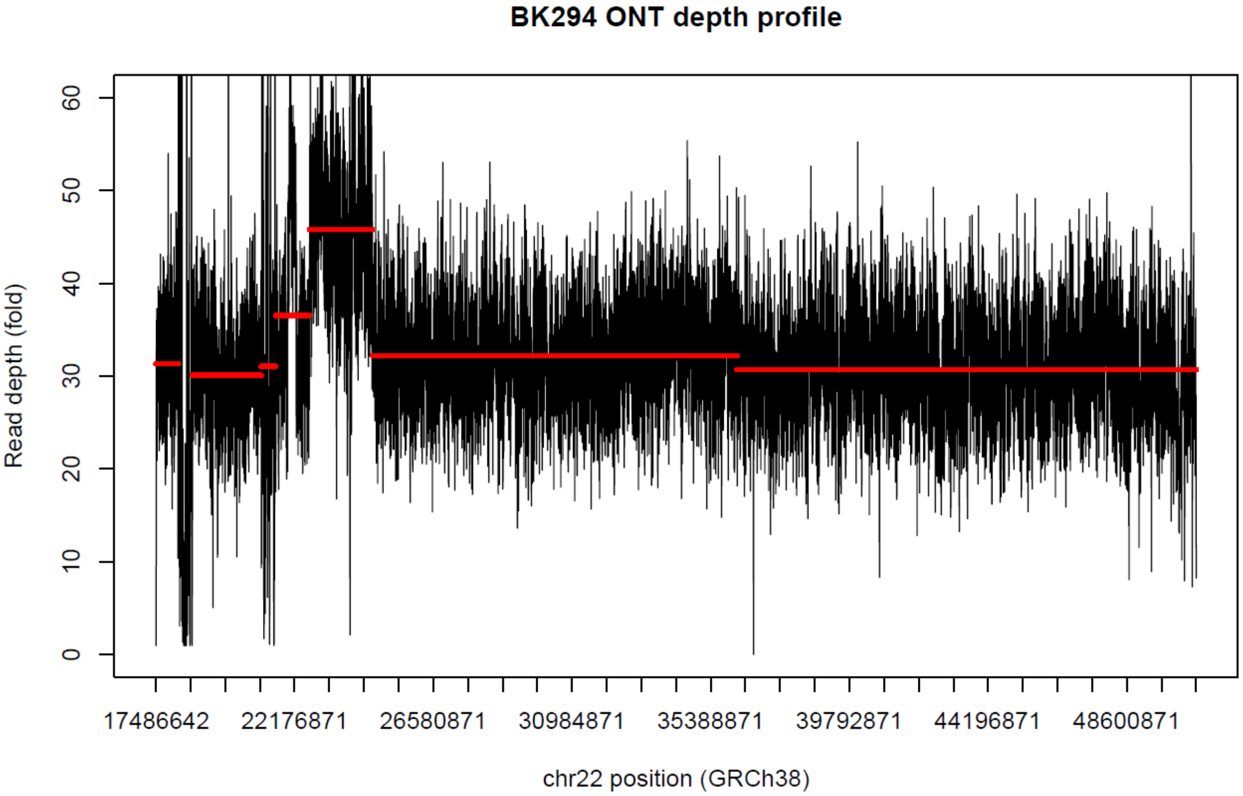
BK294 ONT depth profile
Read depth (fold)
0
10
20
30
40
50
60
17486642
22176871
26580871
30984871
35388871
39792871
44196871
48600871
chr22 position (GRCh38)

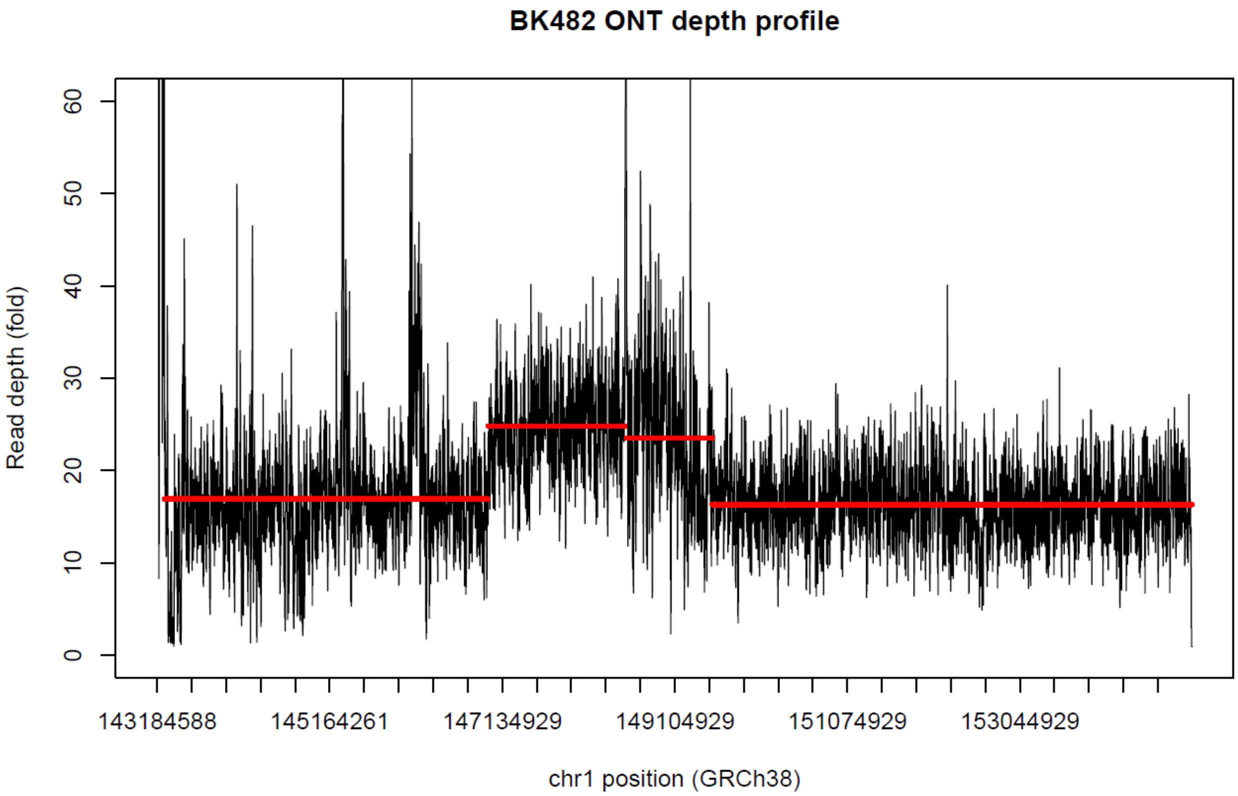
BK482 ONT depth profile
Read depth (fold)
0
10
20
30
40
50
60
143184588
145164261
147134929
149104929
151074929
153044929
chr1 position (GRCh38)

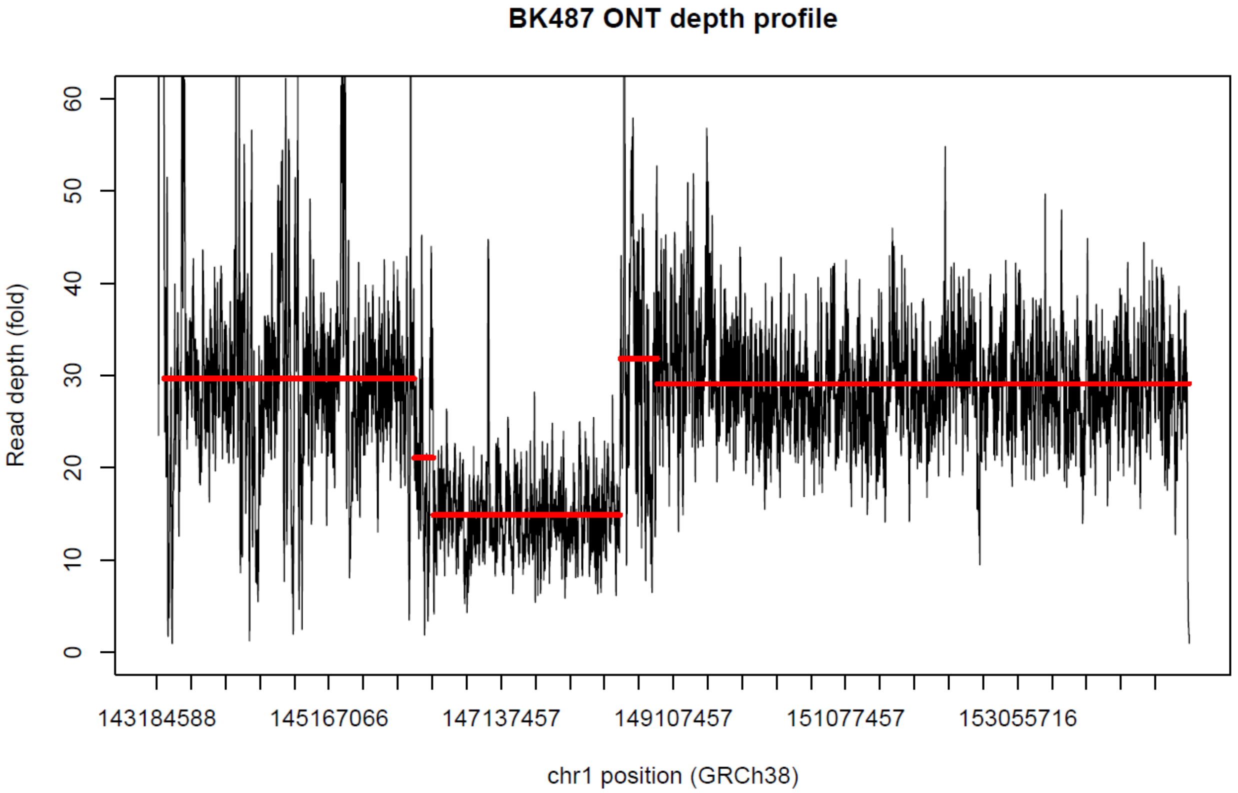
BK487 ONT depth profile
Read depth (fold)
0
10
20
30
40
50
60
143184588
145167066
147137457
149107457
151077457
153055716
chr1 position (GRCh38)

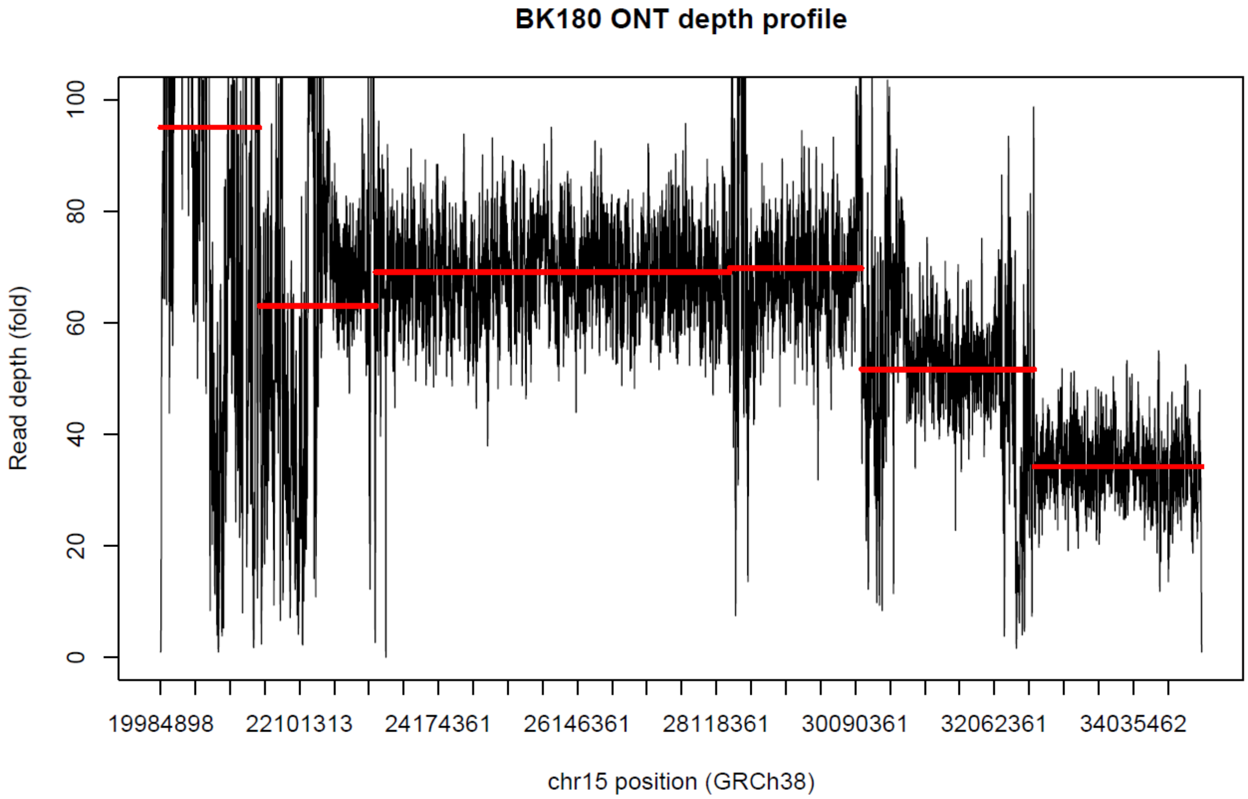
BK180 ONT depth profile
Read depth (fold)
0
20
40
60
80
100
19984898
22101313
24174361
26146361
28118361
30090361
32062361
34035462
chr15 position (GRCh38)

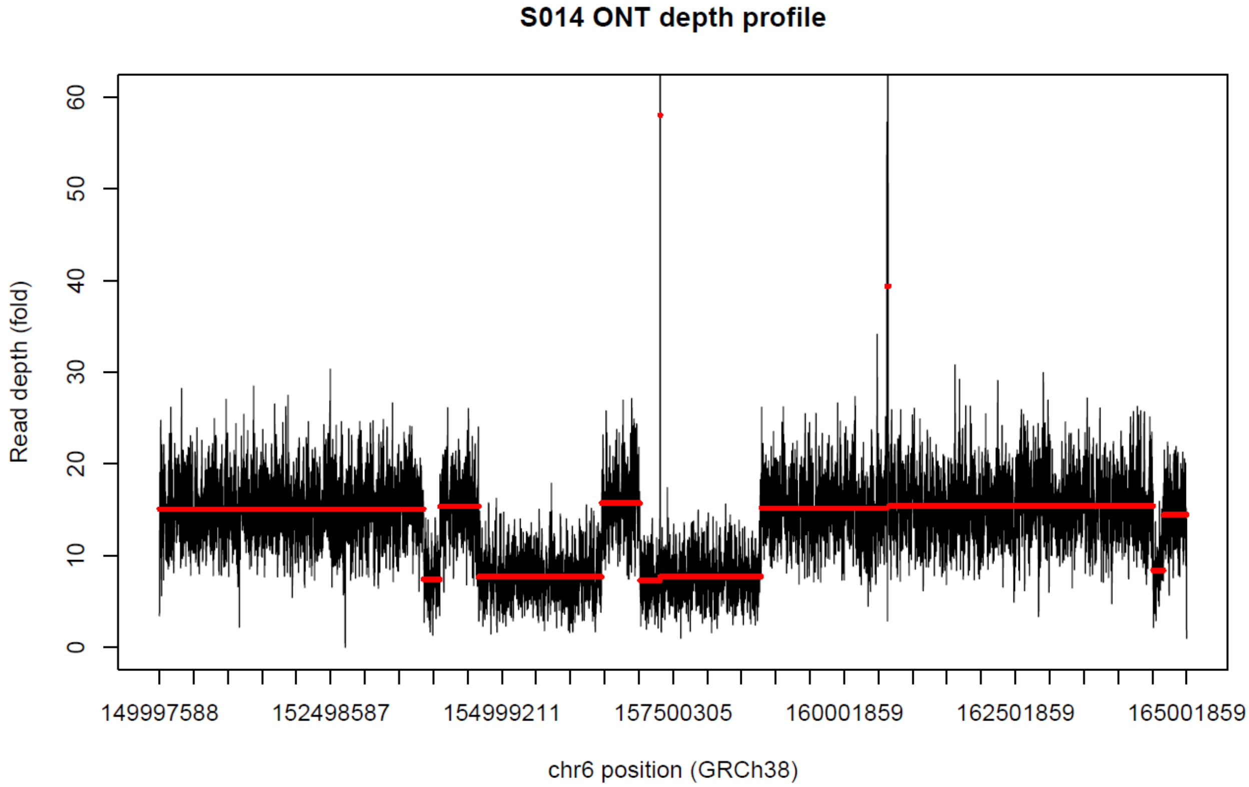
S014 ONT depth profile
Read depth (fold)
0
10
20
30
40
50
60
149997588
152498587
154999211
157500305
160001859
162501859
165001859
chr6 position (GRCh38)

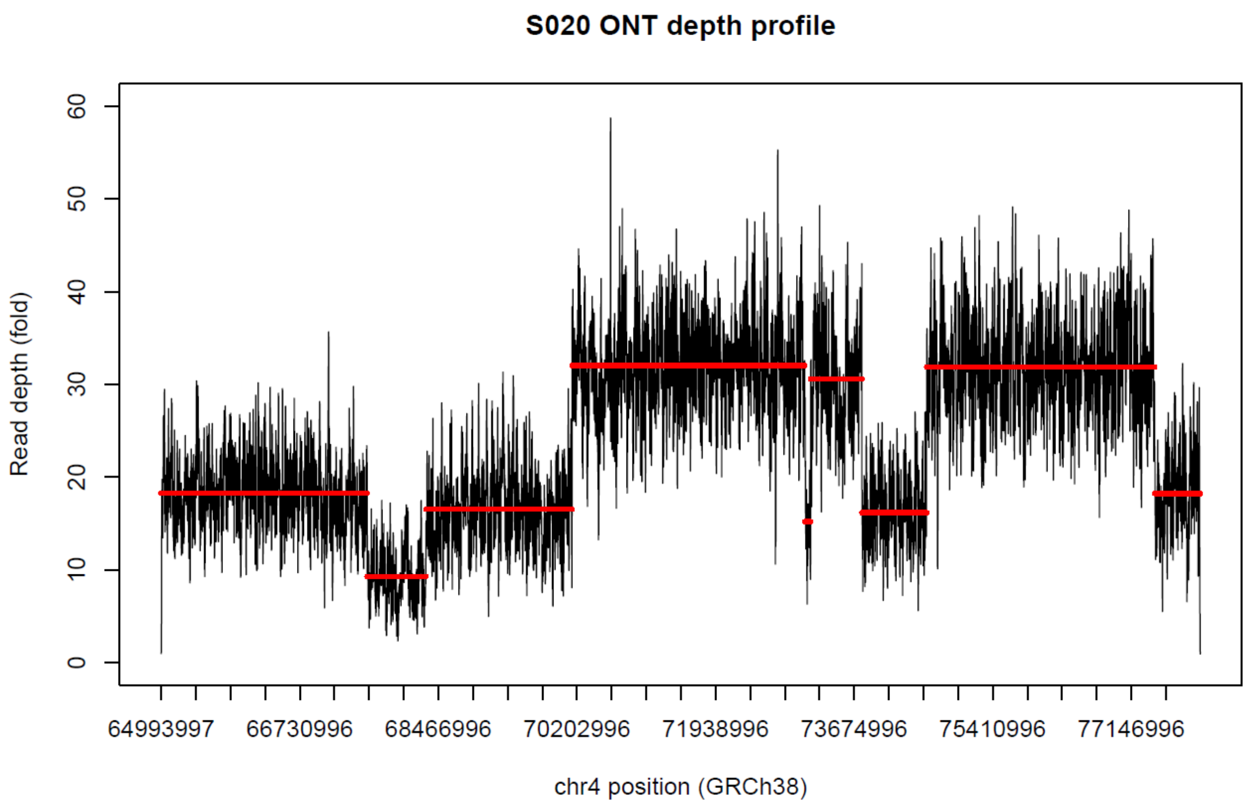

S020 ONT depth profile
Read depth (fold)
0
10
20
30
40
50
60
64993997
66730996
68466996
70202996
71938996
73674996
75410996
77146996
chr4 position (GRCh38)

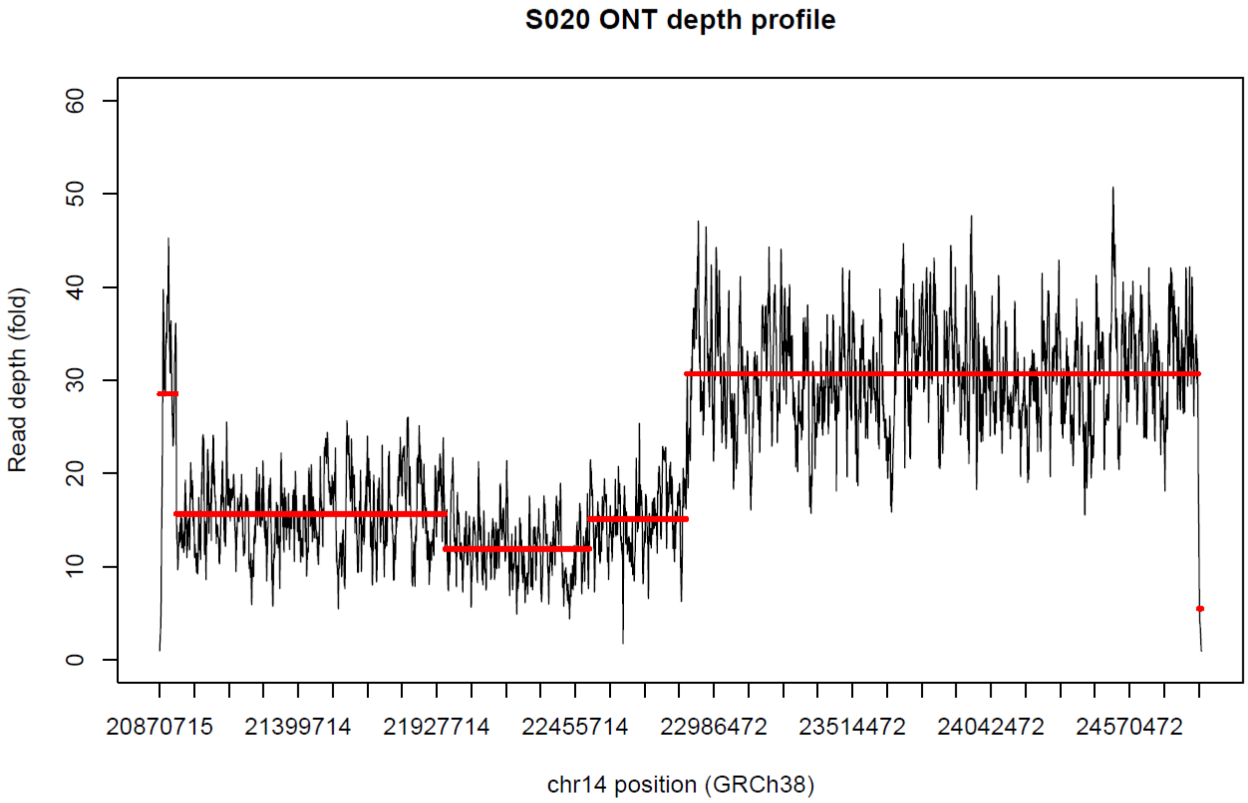

S020 ONT depth profile
Read depth (fold)
0
10
20
30
40
50
60
20870715
21399714
21927714
22455714
22986472
23514472
24042472
24570472
chr14 position (GRCh38)

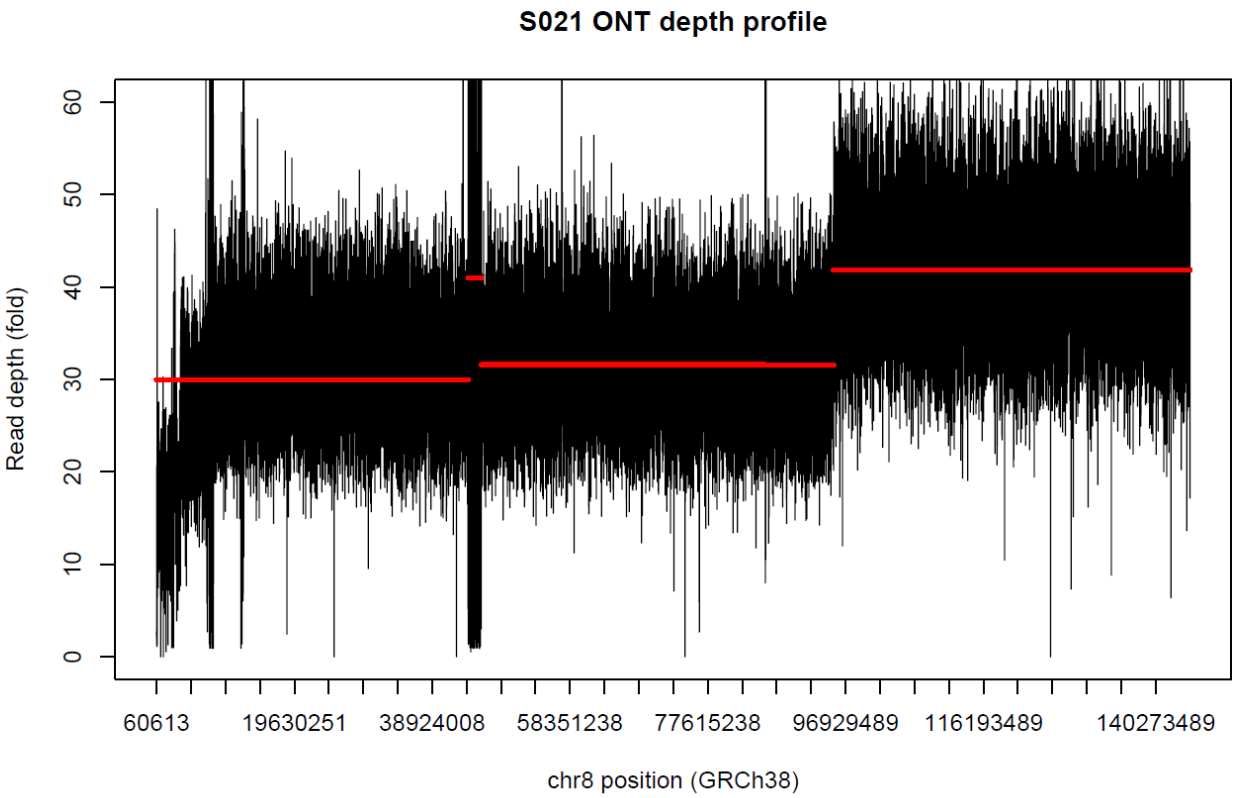
S021 ONT depth profile
Read depth (fold)
0
10
20
30
40
50
60
60613
19630251
38924008
58351238
77615238
96929489
116193489
140273489
chr8 position (GRCh38)

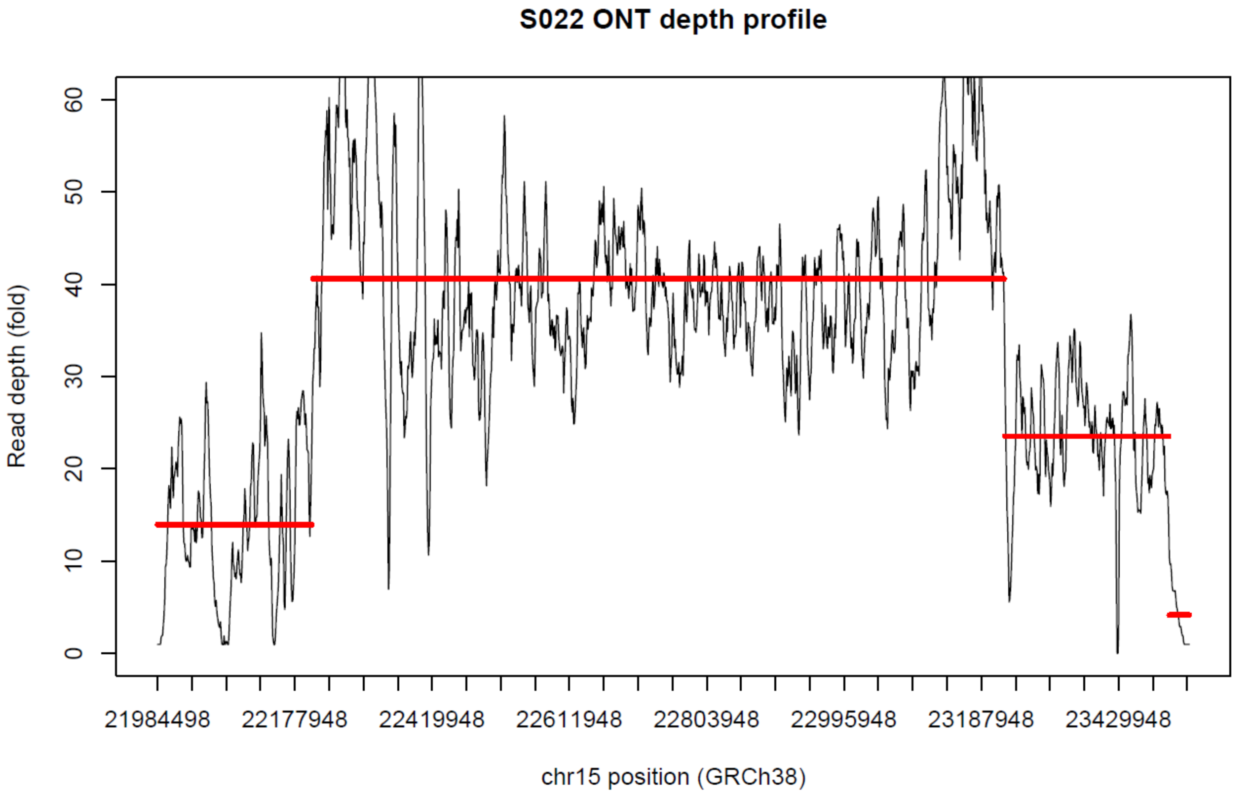
S022 ONT depth profile
Read depth (fold)
0
10
20
30
40
50
60
21984498
22177948
22419948
22611948
22803948
22995948
23187948
23429948
chr15 position (GRCh38)

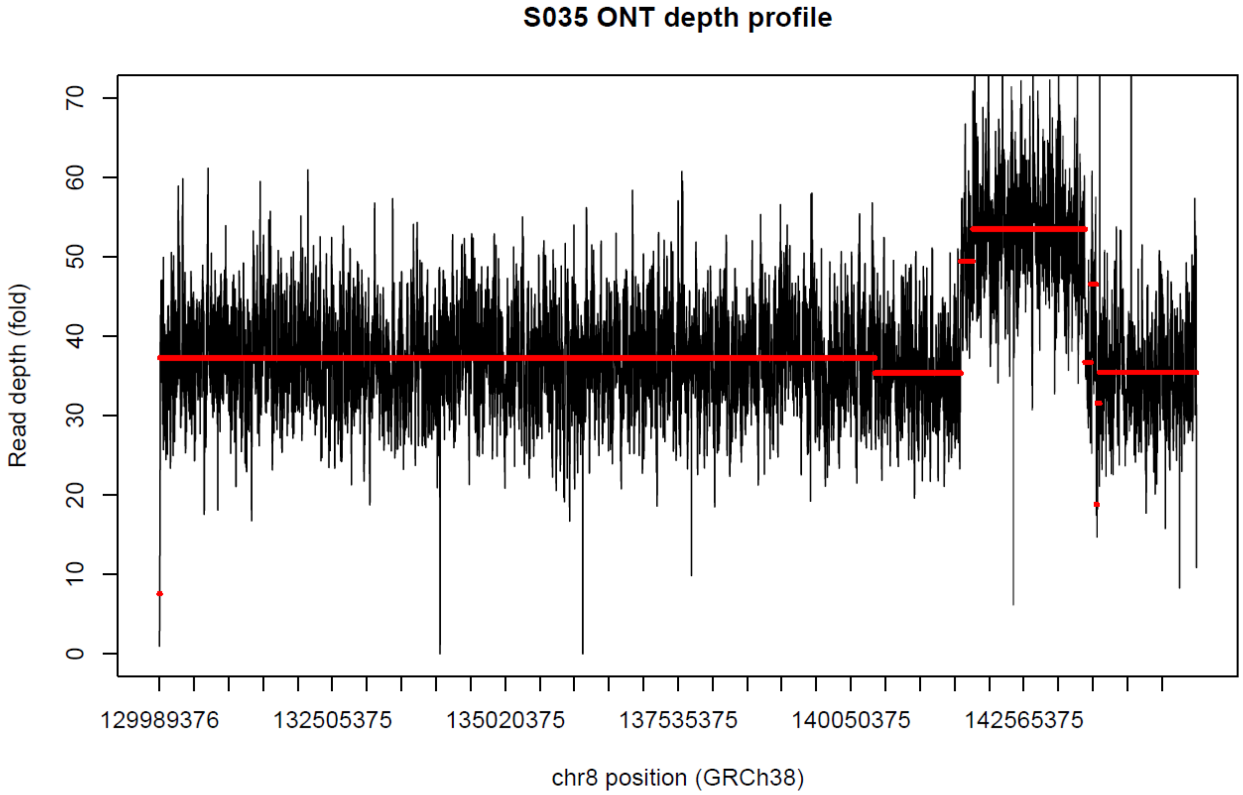
S035 ONT depth profile
Read depth (fold)
0
10
20
30
40
50
60
70
129989376
132505375
135020375
137535375
140050375
142565375
chr8 position (GRCh38)

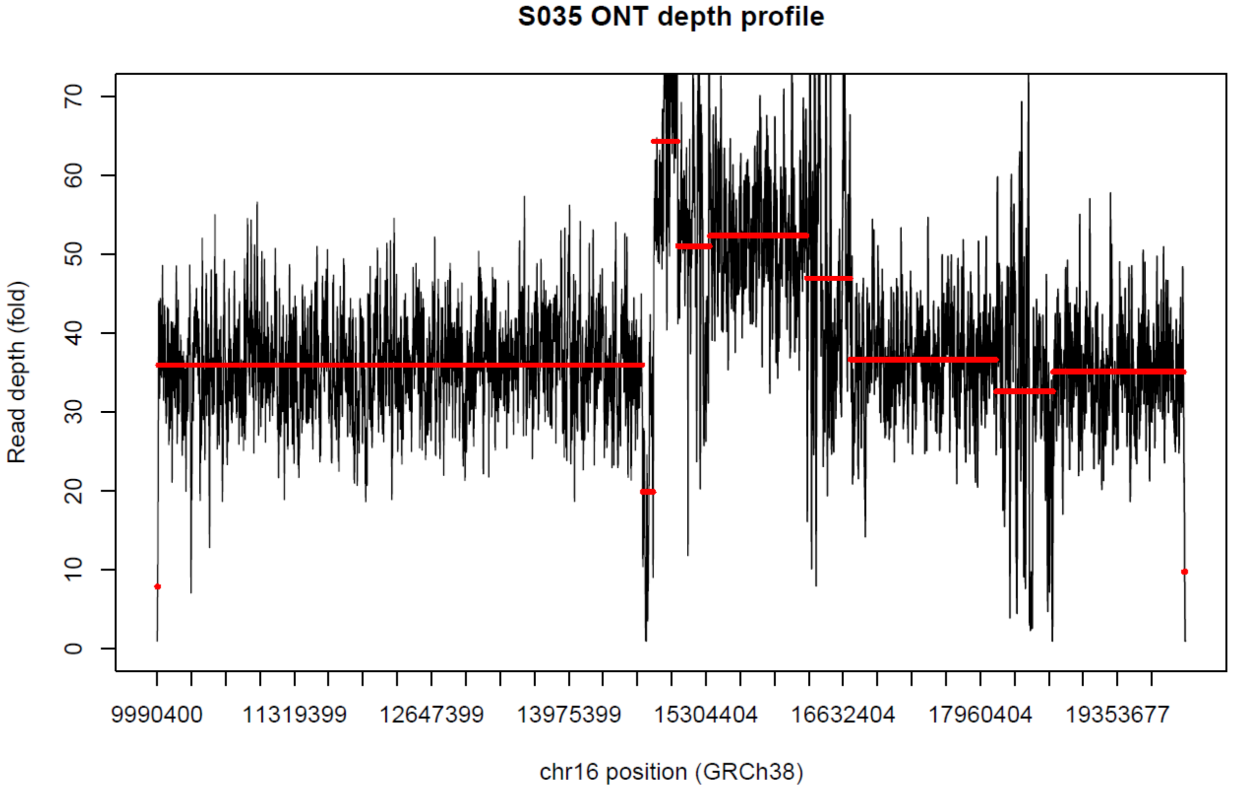
S035 ONT depth profile
Read depth (fold)
0
10
20
30
40
50
60
70
9990400
11319399
12647399
13975399
15304404
16632404
17960404
19353677
chr16 position (GRCh38)

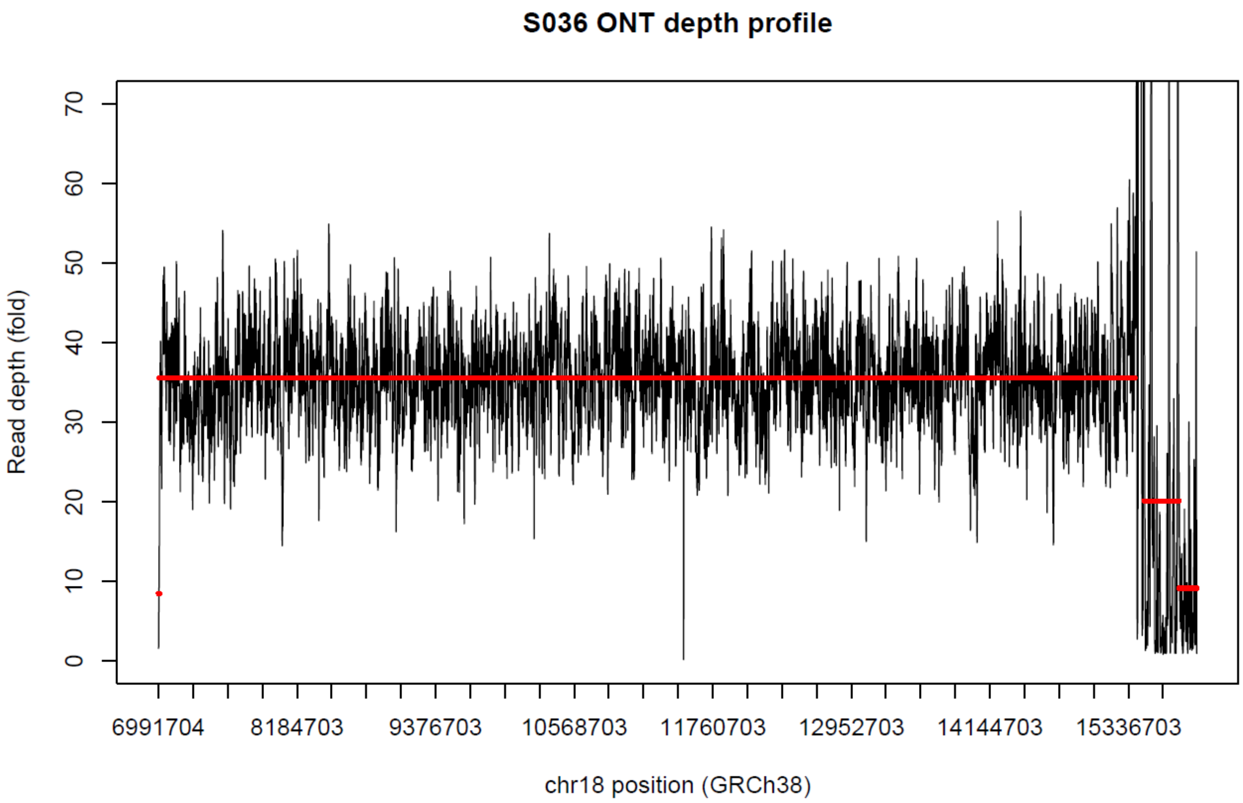
S036 ONT depth profile
Read depth (fold)
0
10
20
30
40
50
60
70
6991704
8184703
9376703
10568703
11760703
12952703
14144703
15336703
chr18 position (GRCh38)

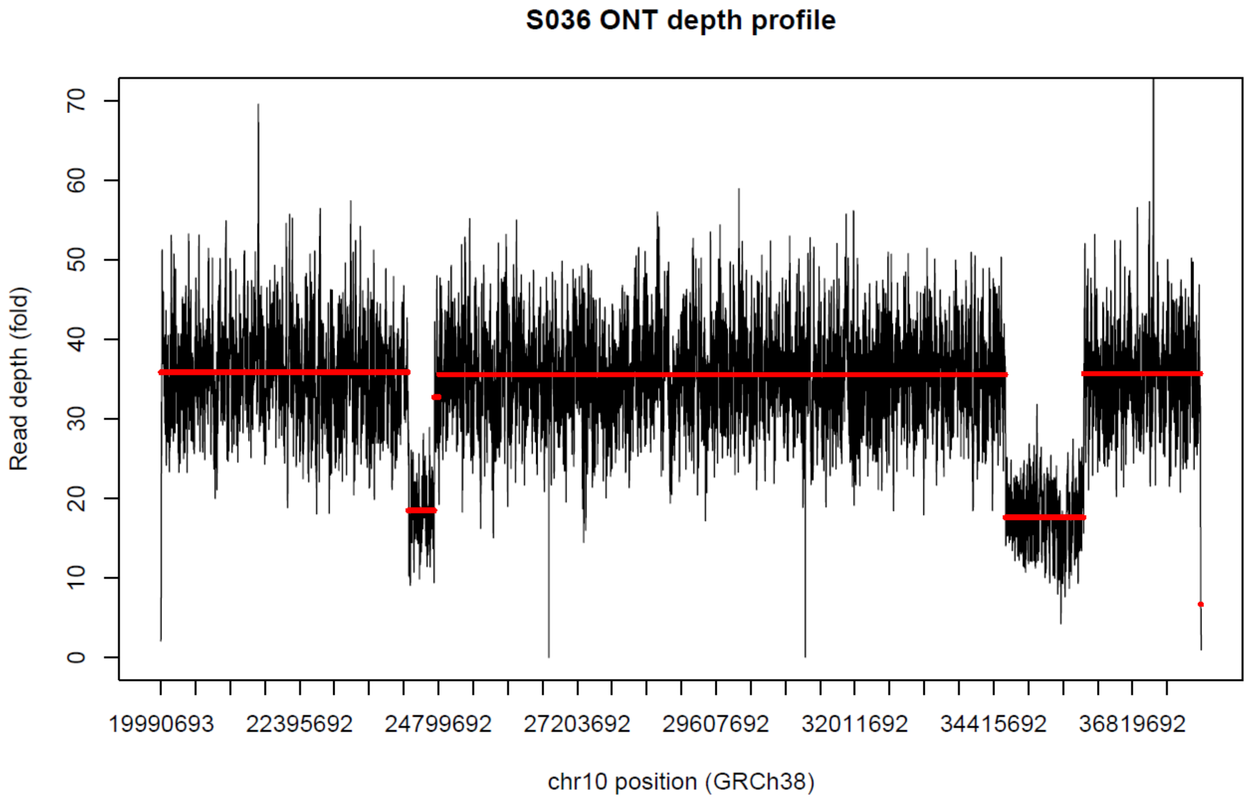
S036 ONT depth profile
Read depth (fold)
0
10
20
30
40
50
60
70
19990693
22395692
24799692
27203692
29607692
32011692
34415692
36819692
chr10 position (GRCh38)

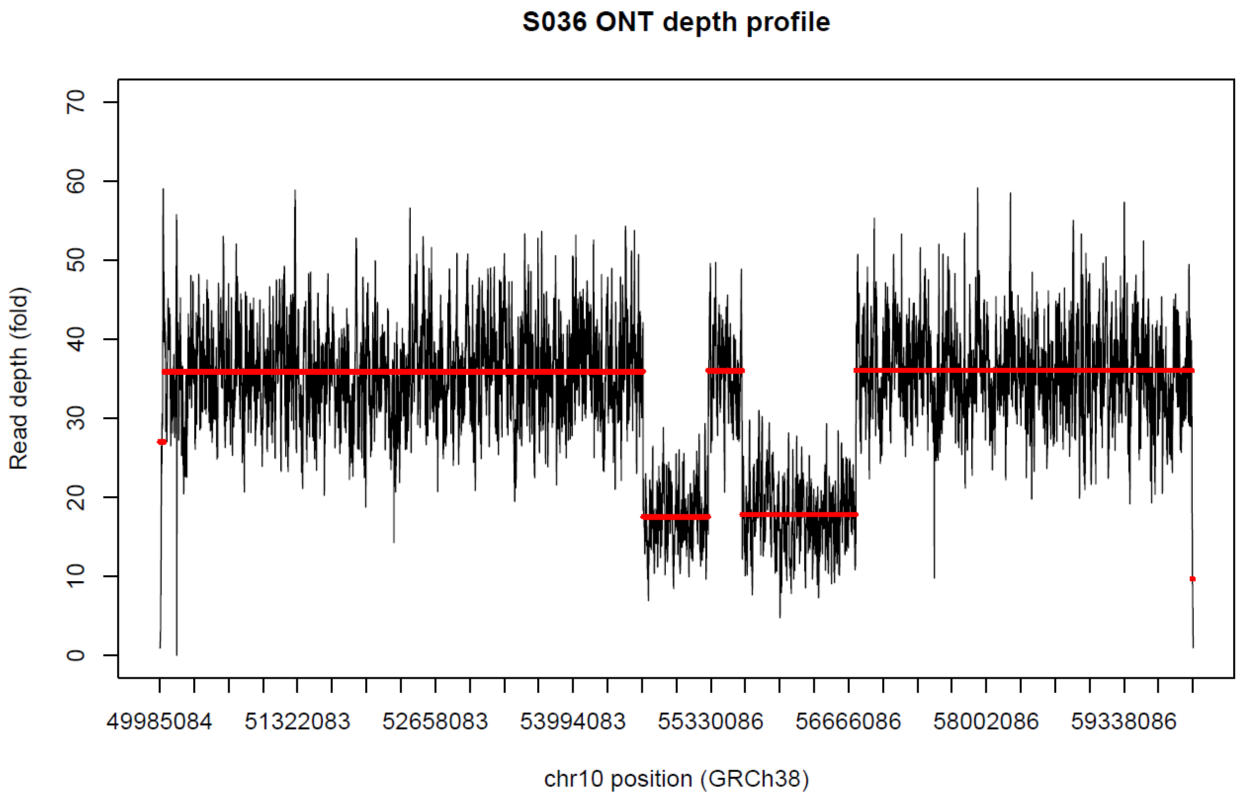
S036 ONT depth profile
Read depth (fold)
0
10
20
30
40
50
60
70
49985084
51322083
52658083
53994083
55330086
56666086
58002086
59338086
chr10 position (GRCh38)

**Figure S2. BK144-03, known 22q13.3 deletion.**

**A.** Coverage for sample. **B.** Centromere proximal end of deletion, long-read BAM file is top track, short-read BAM is bottom track. Long reads that span the deletion breakpoint are highlighted with color; IGV view is chr22:48,119,958-48,120,228. **C.** Telomere proximal end of deletion, long-read BAM file is the top track while short-read file is the bottom track. Long reads that span the deletion are colored as in (B); IGV view is chr22:50,757,177-50,757,279. **D.** IGV view of only long reads at the end of chromosome 22; view is chr22:50,739,987-50,818,468.

A:

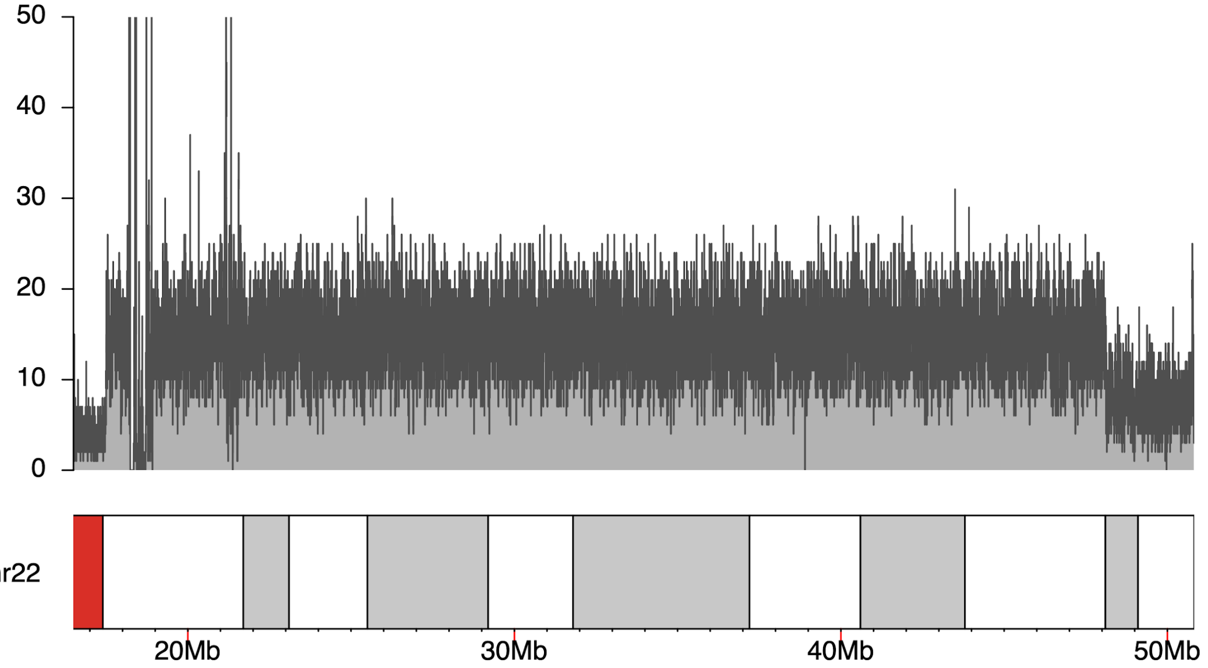

B:

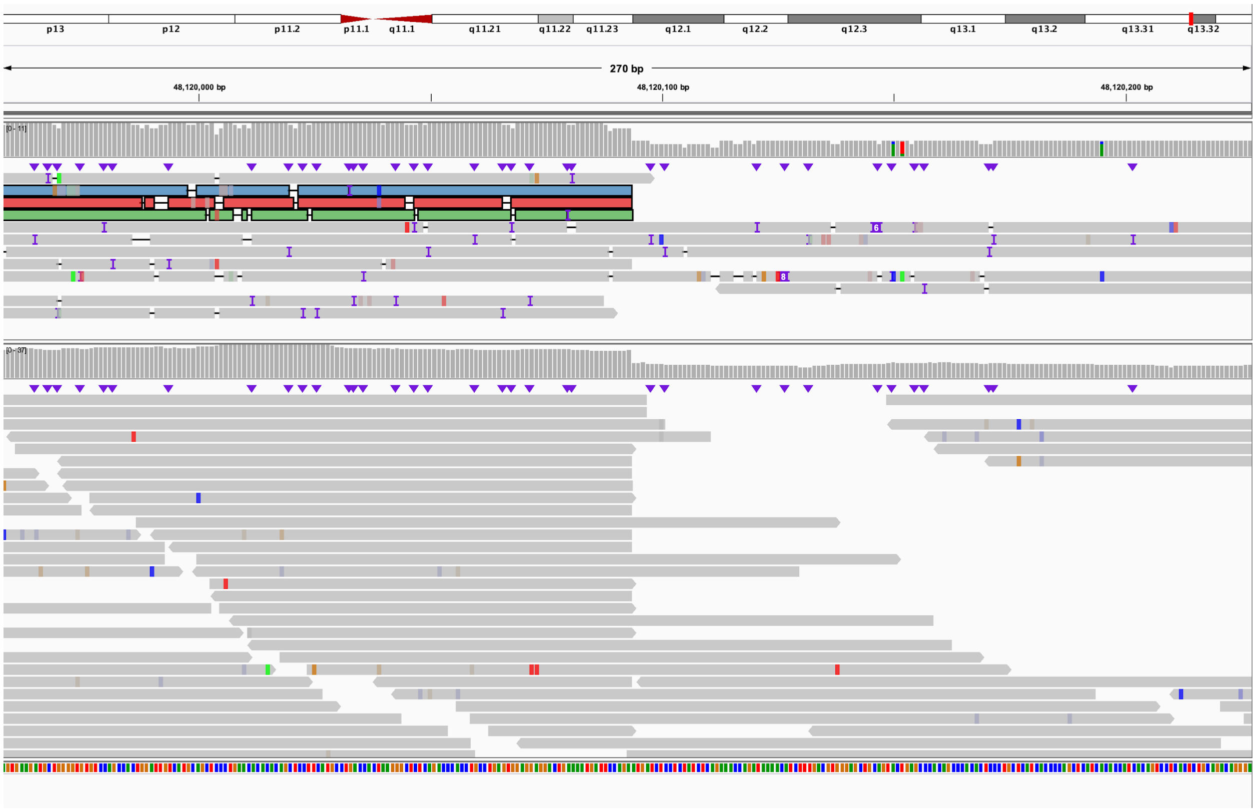
p13
p12
p11.2
p11.1
q11.1
q11.21
q11.22
q11.23
q12.1
q12.2
q12.3
q13.1
q13.2
q13.31
q13.32
270 bp
48,120,000 bp
48,120,100 bp
48,120,200 bp

C:

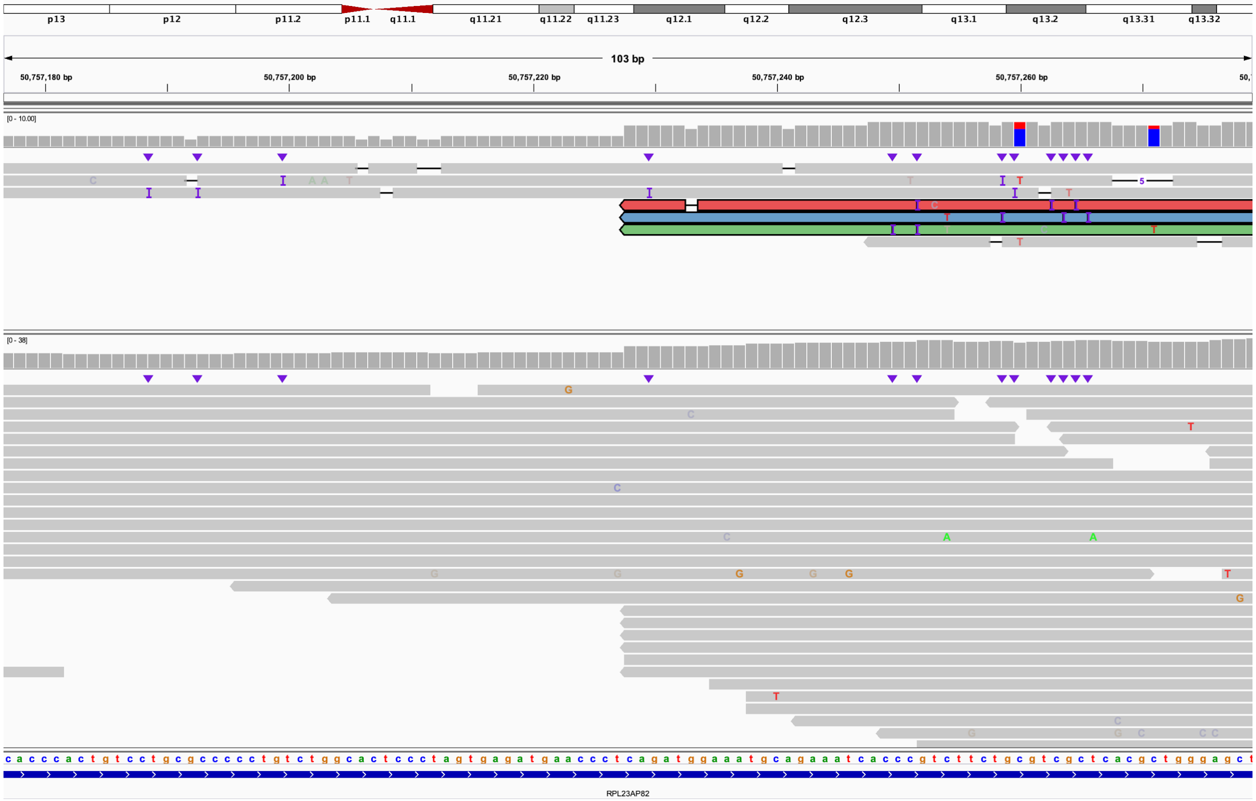
p13
p12
p11.2
p11.1
q11.1
q11.21
q11.22
q11.23
q12.1
q12.2
q12.3
q13.1
q13.2
q13.31
q13.32
103 bp
50,757,180 bp
50,757,200 bp
50,757,220 bp
50,757,240 bp
50,757,260 bp
[0 - 10.00]
[0 - 38]
RPL23AP82

D:

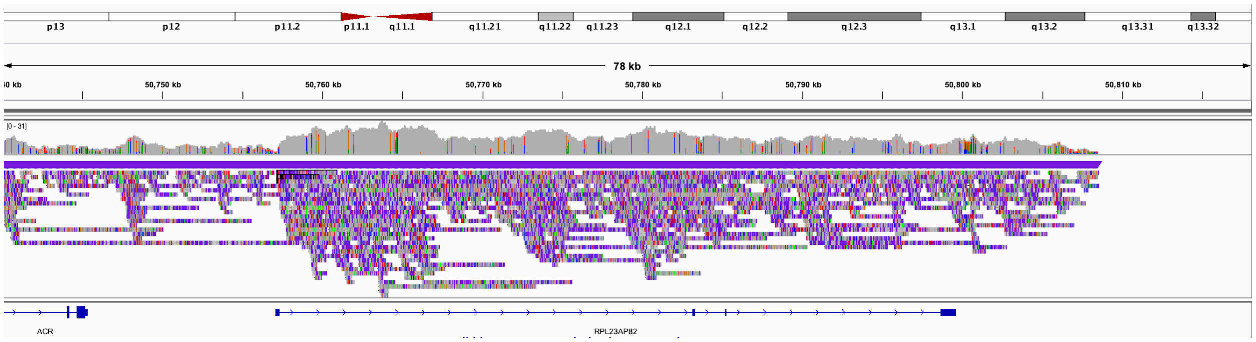
p13
p12
p11.2
p11.1
q11.1
q11.21
q11.22
q11.23
q12.1
q12.2
q12.3
q13.1
q13.2
q13.31
q13.32
78 kb
50,750 kb
50,760 kb
50,770 kb
50,780 kb
50,790 kb
50,800 kb
50,810 kb
[0 - 31]
ACR
RPL23AP82

**Figure S3. BK180-03, known 15q11-13 duplication.**

Coverage of target region in 15q. No definitive duplication breakpoint was found using targeted long reads.

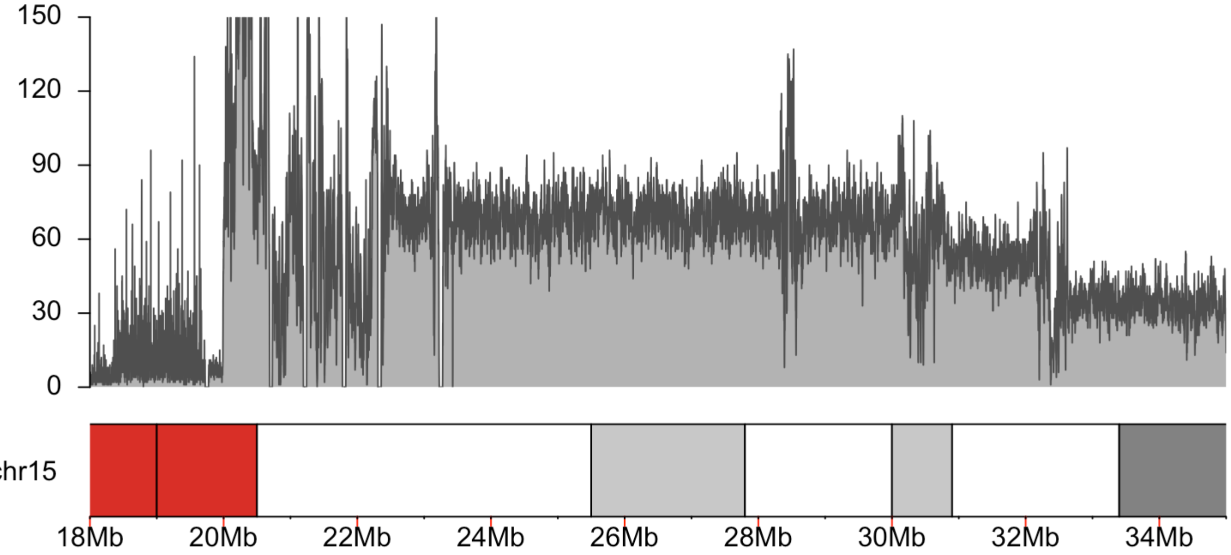

**Figure S4. BK294-03, known 22q11.2 duplication.**

Coverage of chromosome 22 target region. No definitive duplication breakpoint was found using targeted long reads.

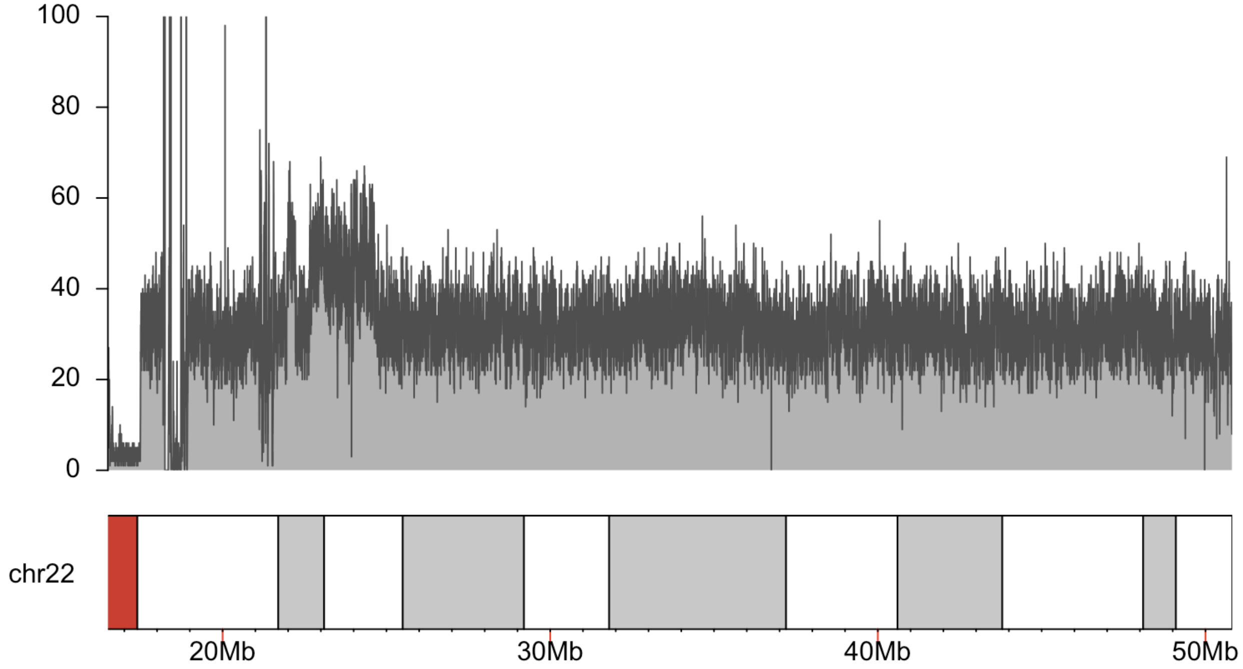

**Figure S5. BK364-03, known 1p36.11 duplication.**

**A.** Coverage of target region demonstrating the presence of a duplication. **B.** IGV view of 3' end, or centromere-proximal end of the duplication, reads that define the deletion breakpoint are represented by colors. IGV view is of chr1:27,792,056-27,792,256. **C.** Reads from B are split and align to the 5' end, or telomere-proximal end of the duplication. The region contains several repetitive elements, as shown in the bottom track. IGV view is of chr1:26,956,878-26,959,178. **D.** Fragments of four reads seen in B align incorrectly to a region outside of the duplicated region that includes a SINE/Alu and low-complexity region (bottom track); these fragments are notably shorter than those in C. IGV view is of chr1:26,895,202-26,895,689.

A:

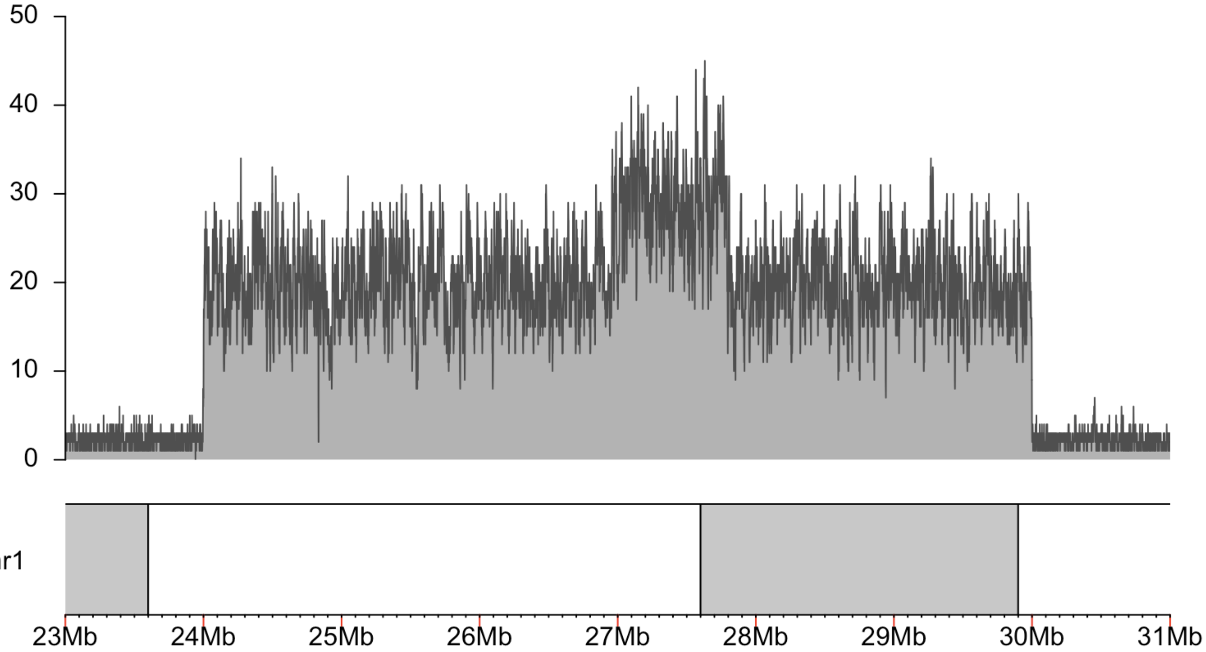

B:

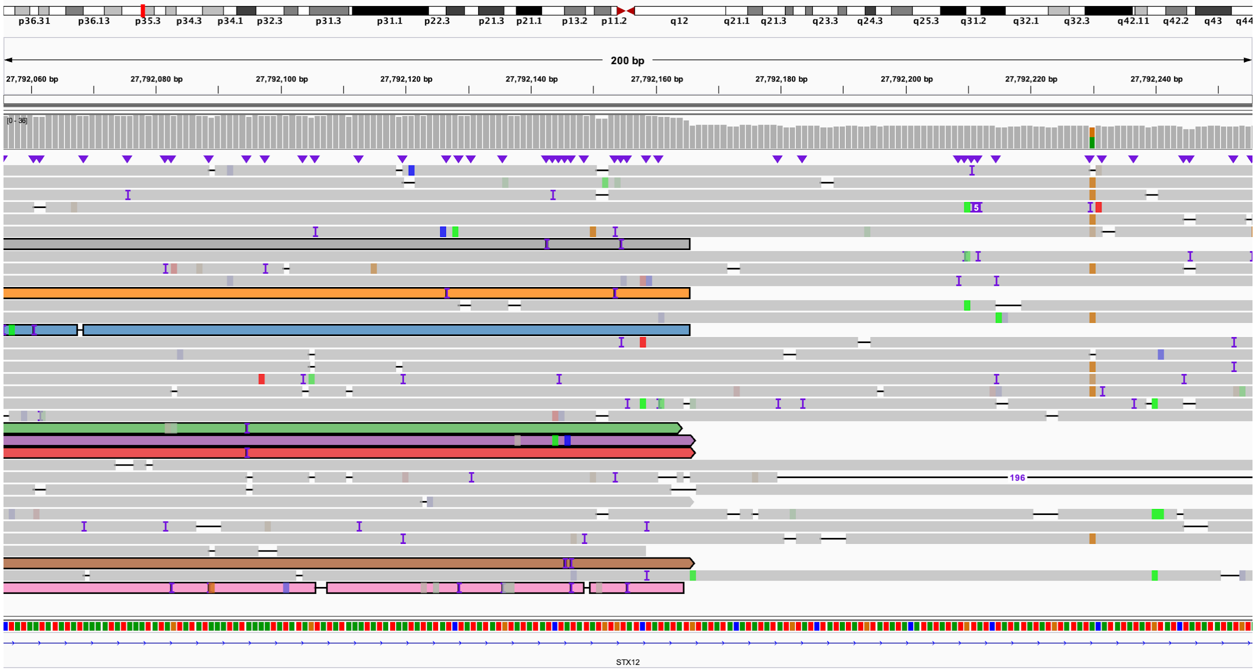
p36.31 p36.13 p35.3 p34.3 p34.1 p32.3 p31.3 p31.1 p22.3 p21.3 p21.1 p13.2 p11.2 q12 q21.1 q21.3 q23.3 q24.3 q25.3 q31.2 q32.1 q32.3 q42.11 q42.2 q43 q44
200 bp
27,792,060 bp
27,792,080 bp
27,792,100 bp
27,792,120 bp
27,792,140 bp
27,792,160 bp
27,792,180 bp
27,792,200 bp
27,792,220 bp
27,792,240 bp
196
STX12

C:

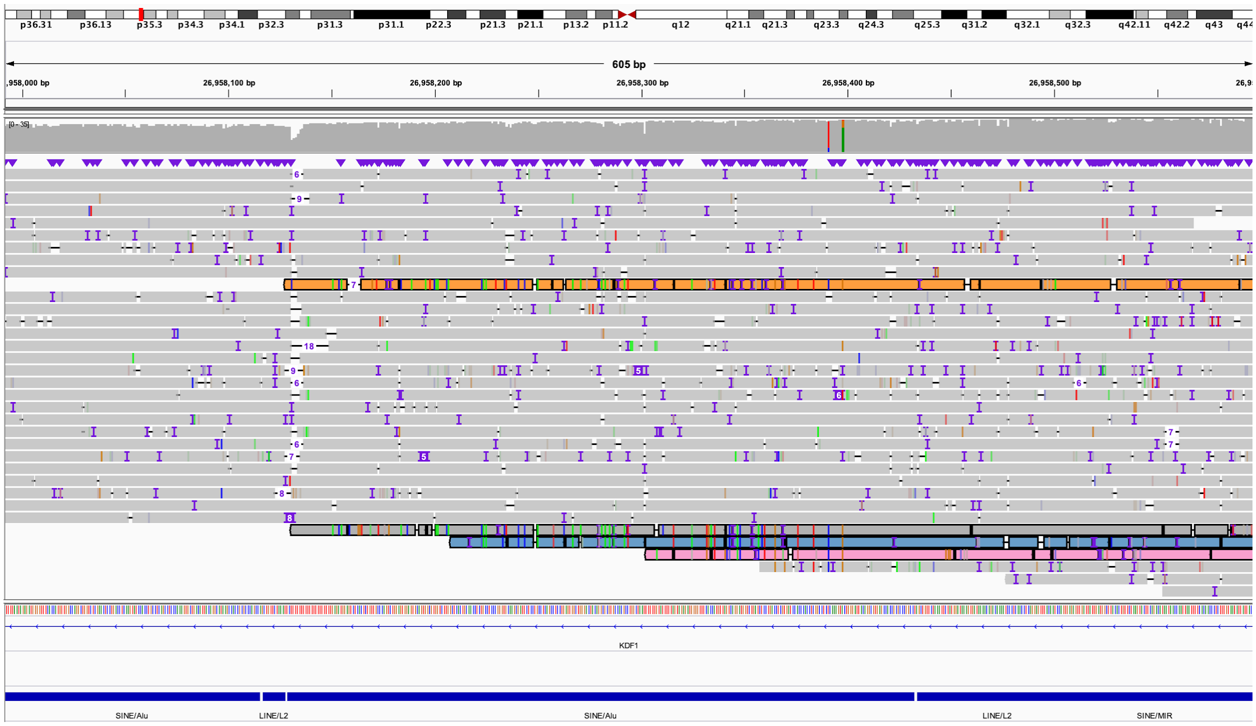
p36.31 p36.13 p35.3 p34.3 p34.1 p32.3 p31.3 p31.1 p22.3 p21.3 p21.1 p13.2 p11.2 q12 q21.1 q21.3 q23.3 q24.3 q25.3 q31.2 q32.1 q32.3 q42.11 q42.2 q43 q44
605 bp
26,958,100 bp
26,958,200 bp
26,958,300 bp
26,958,400 bp
26,958,500 bp
KDF1
SINE/Alu
LINE/L2
SINE/Alu
LINE/L2
SINE/MIR

D:

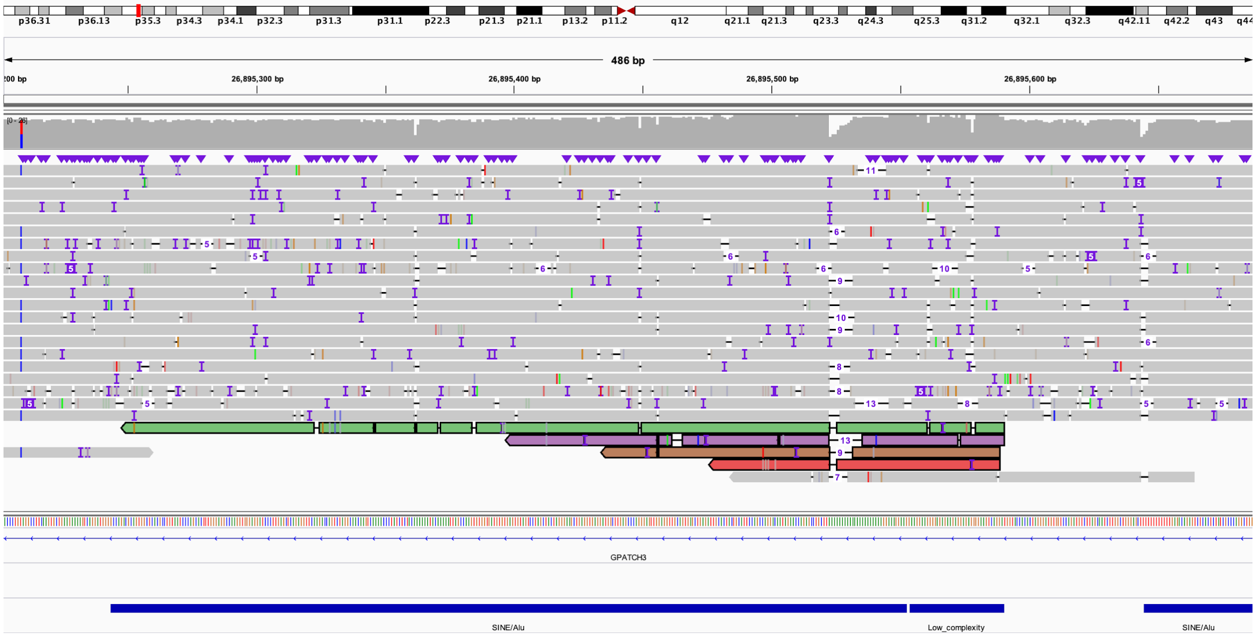
p36.31 p36.13 p35.3 p34.3 p34.1 p32.3 p31.3 p31.1 p22.3 p21.3 p21.1 p13.2 p11.2 q12 q21.1 q21.3 q23.3 q24.3 q25.3 q31.2 q32.1 q32.3 q42.11 q42.2 q43
486 bp
26,895,300 bp
26,895,400 bp
26,895,500 bp
26,895,600 bp
GPATCH3
SINE/Alu
Low_complexity
SINE/Alu

**Figure S6. BK397-101, known 16p11.2 deletion.**

Coverage of chromosome 16 target region. No definitive deletion breakpoint was found using targeted long reads.

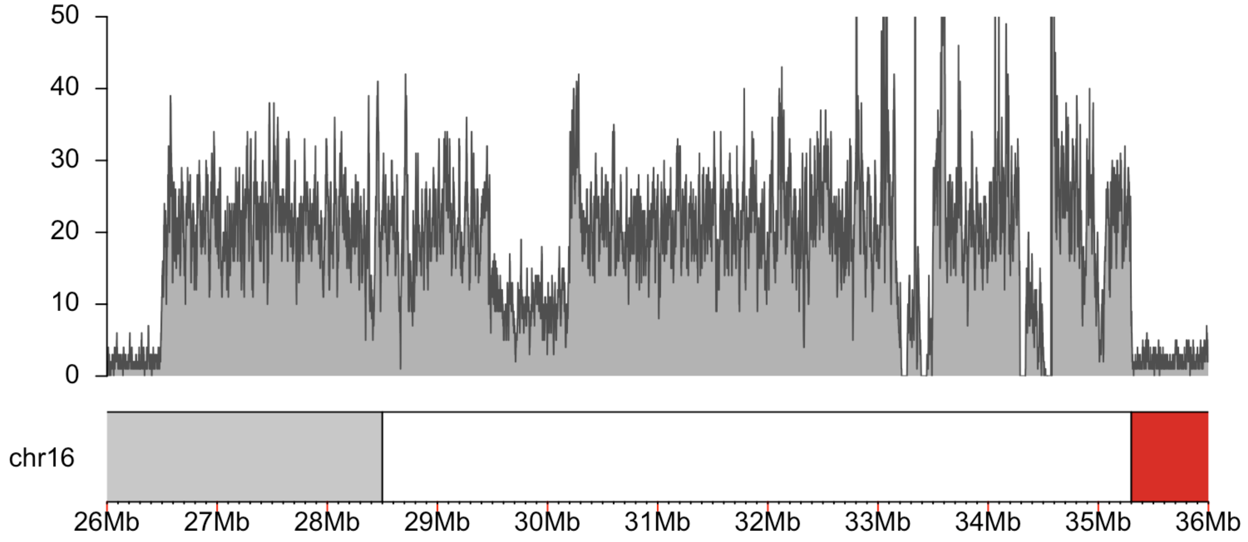

**Figure S7. BK430-103, known 16p11.2 duplication.**

Coverage of chromosome 16 target region. No definitive duplication breakpoint was found using targeted long reads.

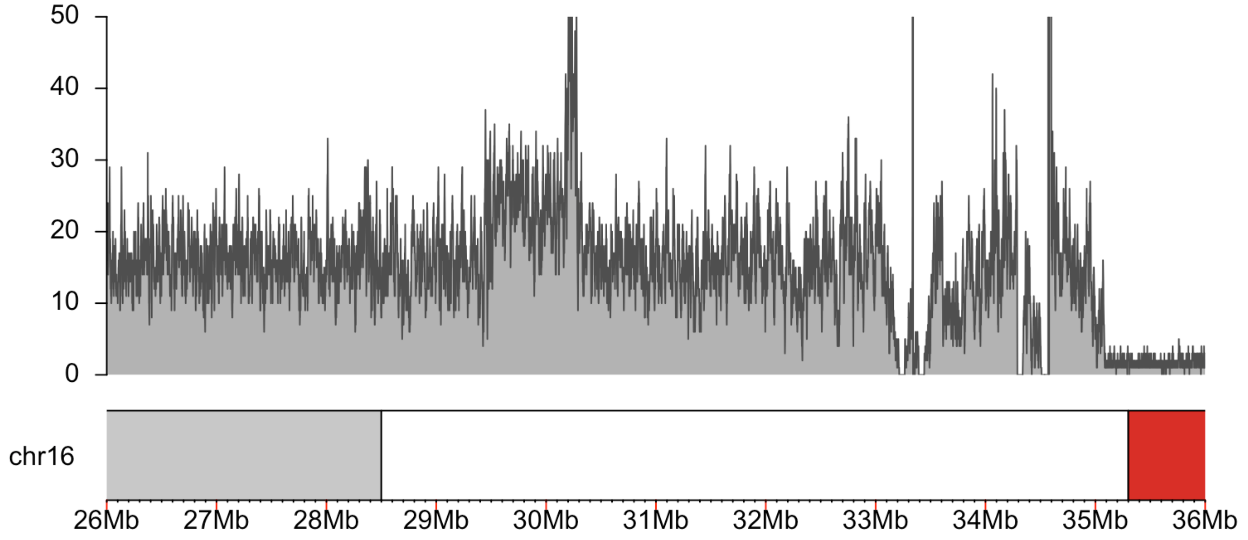

**Figure S8. BK482-101, known 1q21.1 duplication.**

Coverage of chromosome 1 target region. No definitive duplication breakpoint was found using targeted long reads.

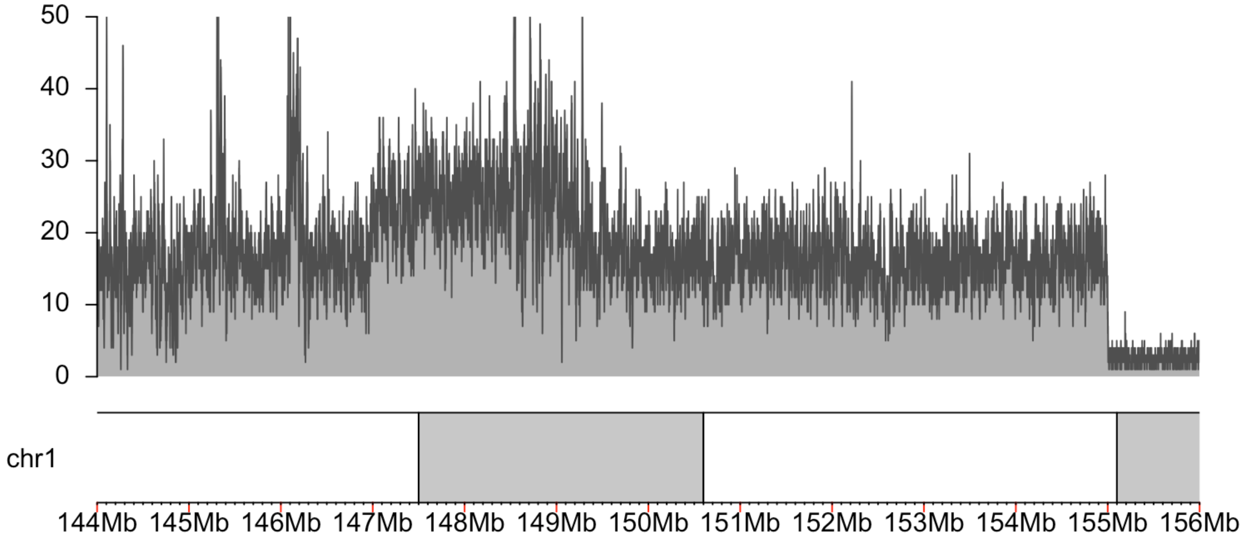

**Figure S9. BK487-101, known 1q21 deletion.**

Coverage of chromosome 1 target region. No definitive deletion breakpoint was found using targeted long reads.

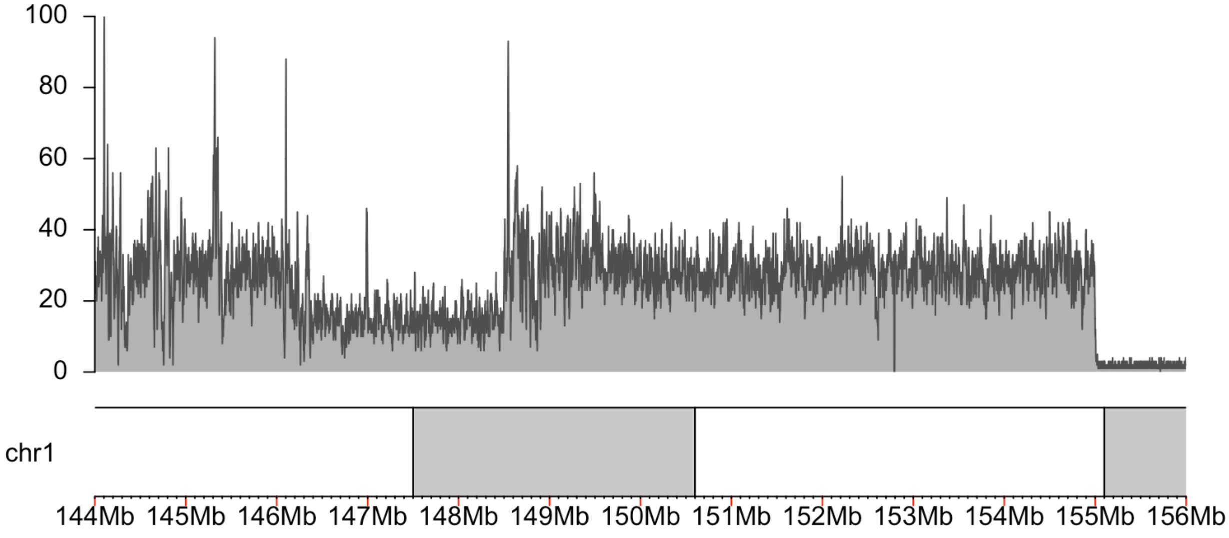

**Figure S10. BK506-03, known 5p15.33 deletion.**

**A.** Coverage of chromosome 5 target region. **B.** Breakpoint of the chromosome 5 deletion in IGV: long-read data is top track; short-read data is bottom track. Colors in top track correspond to colors of reads in C. View is of chr5:6,605,898-6,606,003. **C.** View of both long-read (top track) and short-read (bottom track) data from BK506-03 showing the centromere-proximal sequence is not deleted, but the telomere-proximal sequence is present in three copies. Long reads in the top track can be matched with those from B by their color. IGV view is chr12:132,877,990-132,886,995. **D.** View of short-read sequencing data at the chromosome 5 breakpoint from the parents of BK506-03 (BK506-01 and BK506-02). IGV view is chr5:6,605,898-6,606,003. **E.** View of short-read sequencing data at the chromosome 12 breakpoint from the parents of BK506-03. IGV view is chr12:132,877,990-132,886,995.

A:

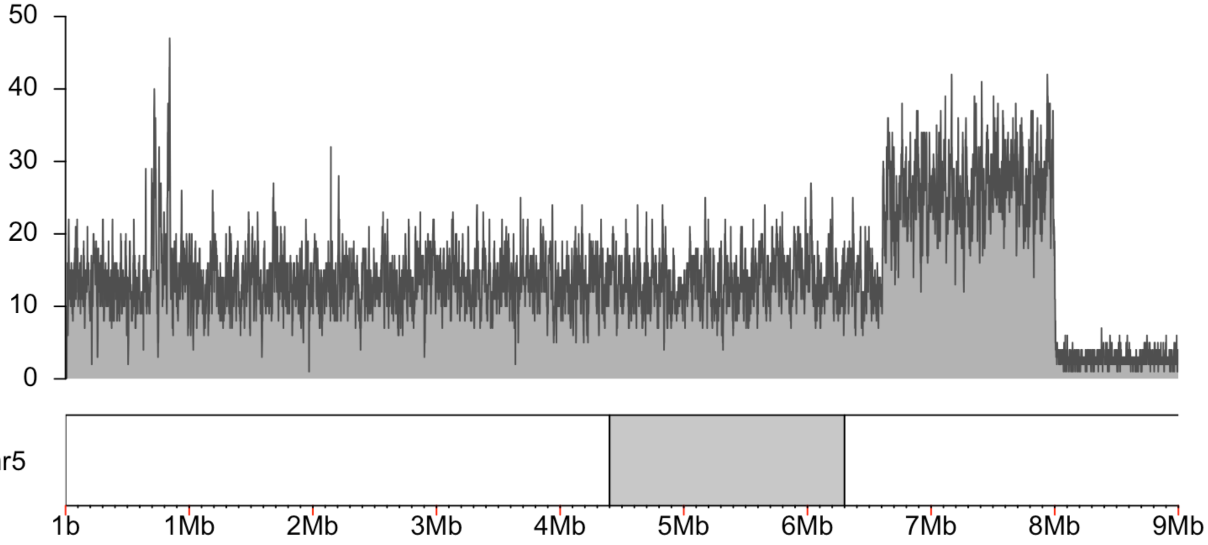

B:

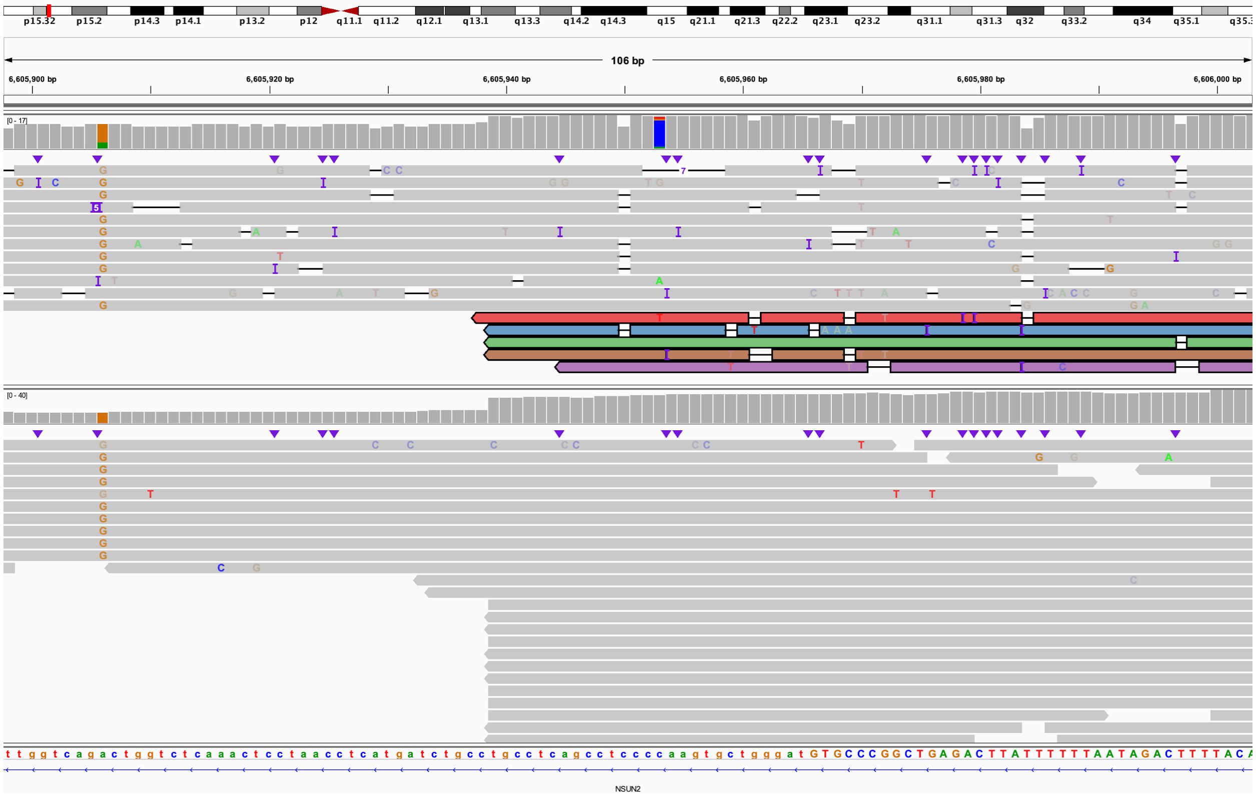
p15.32 p15.2 p14.3 p14.1 p13.2 p12 q11.1 q11.2 q12.1 q13.1 q13.3 q14.2 q14.3 q15 q21.1 q21.3 q22.2 q23.1 q23.2 q31.1 q31.3 q32 q33.2 q34 q35.1
106 bp
6,605,900 bp
6,605,920 bp
6,605,940 bp
6,605,960 bp
6,605,980 bp
6,606,000 bp
[0 - 17]
[0 - 40]
NSUN2

C:

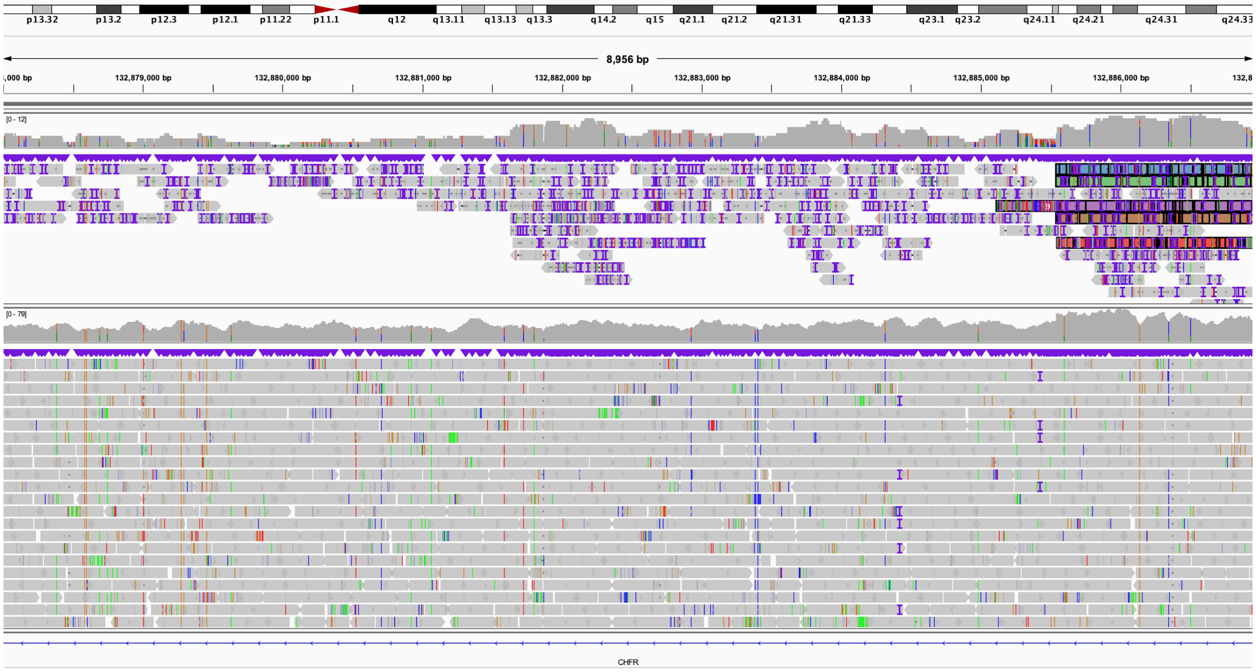
p13.32 p13.2 p12.3 p12.1 p11.22 p11.1 q12 q13.11 q13.13 q13.3 q14.2 q15 q21.1 q21.2 q21.31 q21.33 q23.1 q23.2 q24.11 q24.21 q24.31 q24.33
8,956 bp
132,879,000 bp
132,880,000 bp
132,881,000 bp
132,882,000 bp
132,883,000 bp
132,884,000 bp
132,885,000 bp
132,886,000 bp
[0 - 12]
[0 - 79]
CHFR

D:

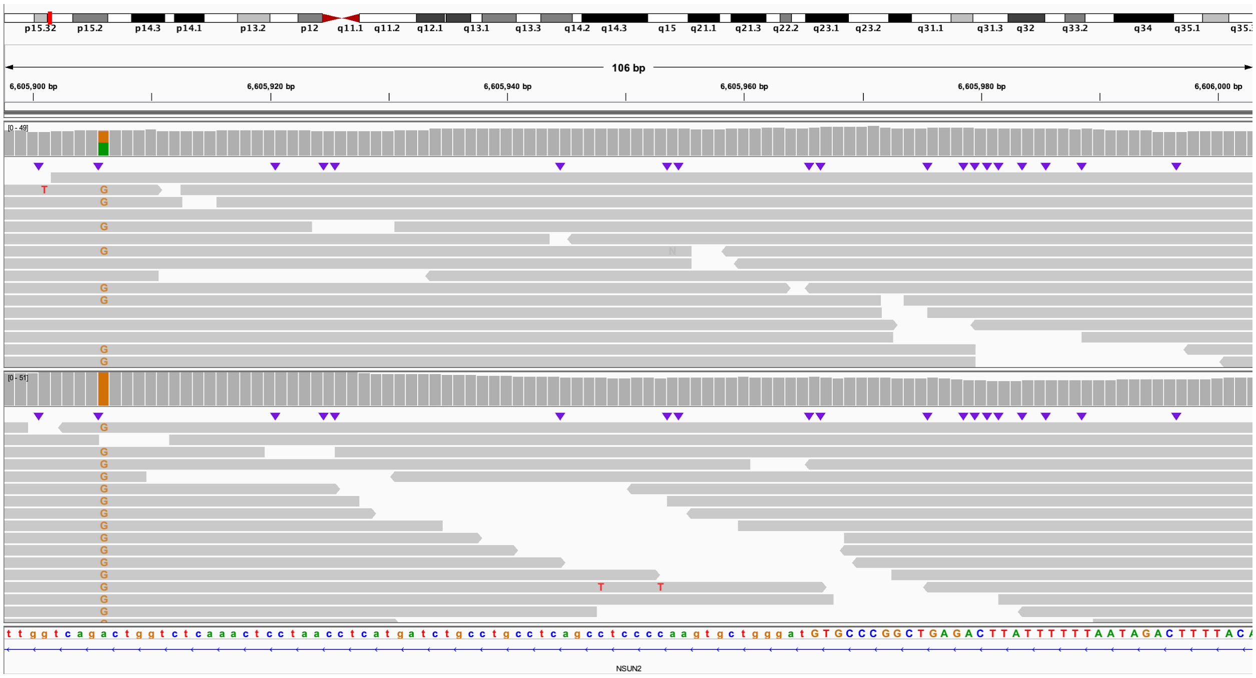
106 bp
6,605,900 bp
6,605,920 bp
6,605,940 bp
6,605,960 bp
6,605,980 bp
6,606,000 bp
[0 - 49]
[0 - 51]
NSUN2

E:

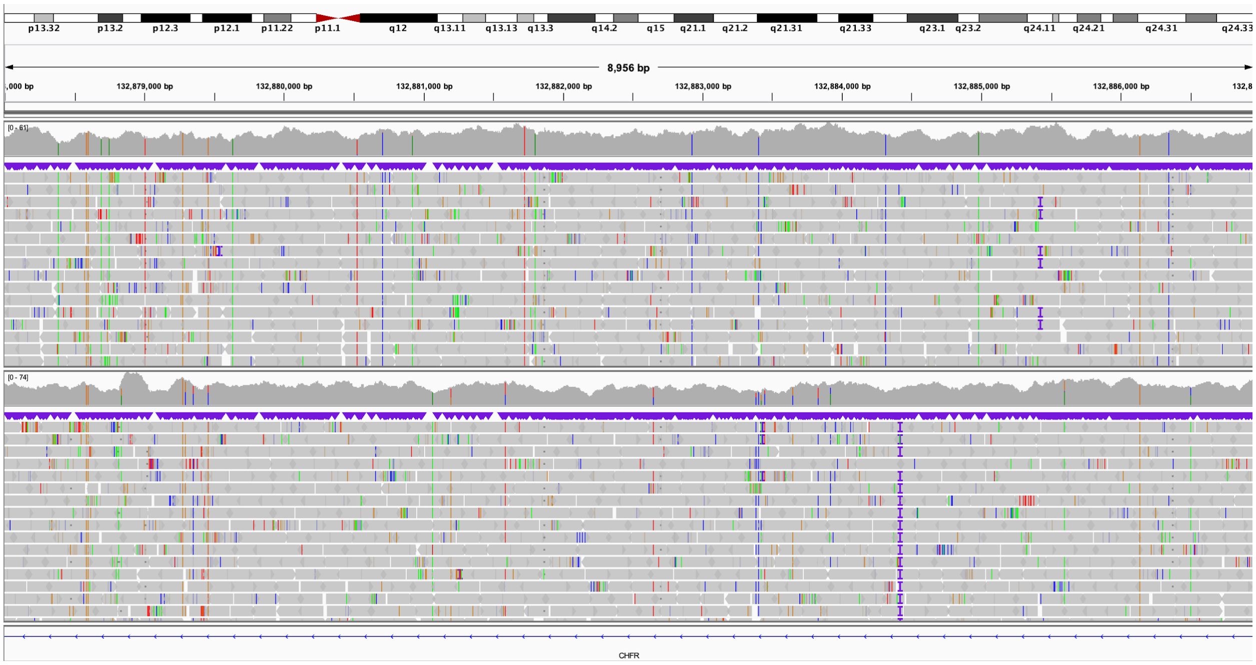
8,956 bp
132,879,000 bp
132,880,000 bp
132,881,000 bp
132,882,000 bp
132,883,000 bp
132,884,000 bp
132,885,000 bp
132,886,000 bp
[0 - 61]
[0 - 74]
CHFR

**Figure S11. S016, known tandem duplication within *CTNND2*.**

**A.** Coverage of duplication within *CTNND2*, gene body of *CTNND2* is represented by the black bar below the coverage graph. **B.** Reads in the telomere-proximal end of the duplication, colors correspond to reads in C. The orientation of reads in B and C confirm that it is a tandem duplication as previously shown (Miller, Squire, and Bennett 2020). IGV view of chr5:11,515,095-11,515,232. **C.** Reads from the centromere-proximal end of the duplication, colors correspond to reads in B. IGV view is of chr5:11,622,960-11,623,099.

A:

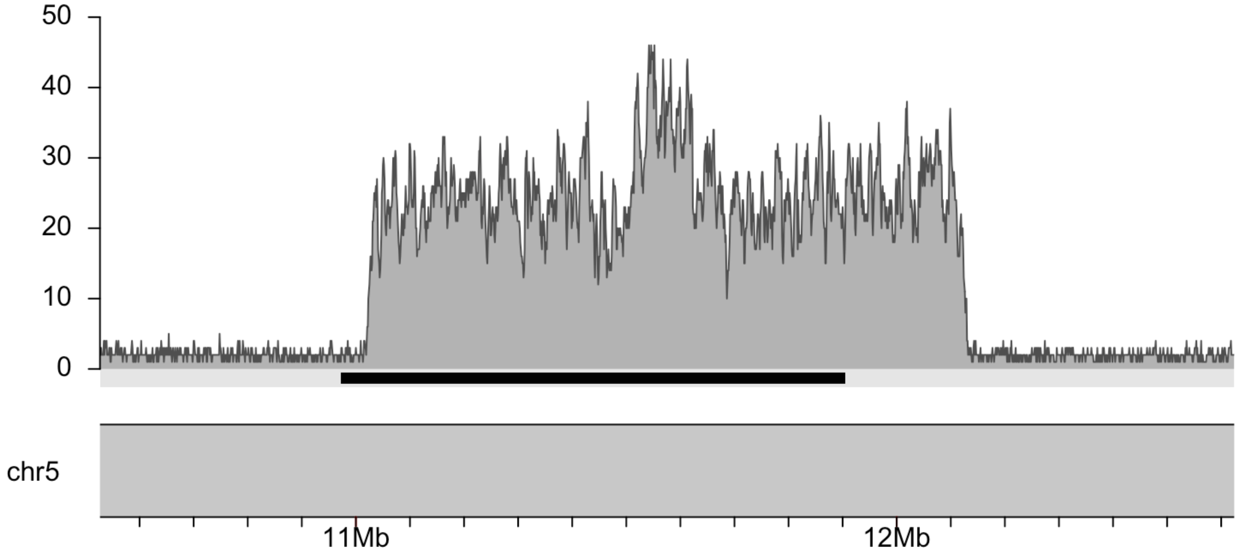

B:

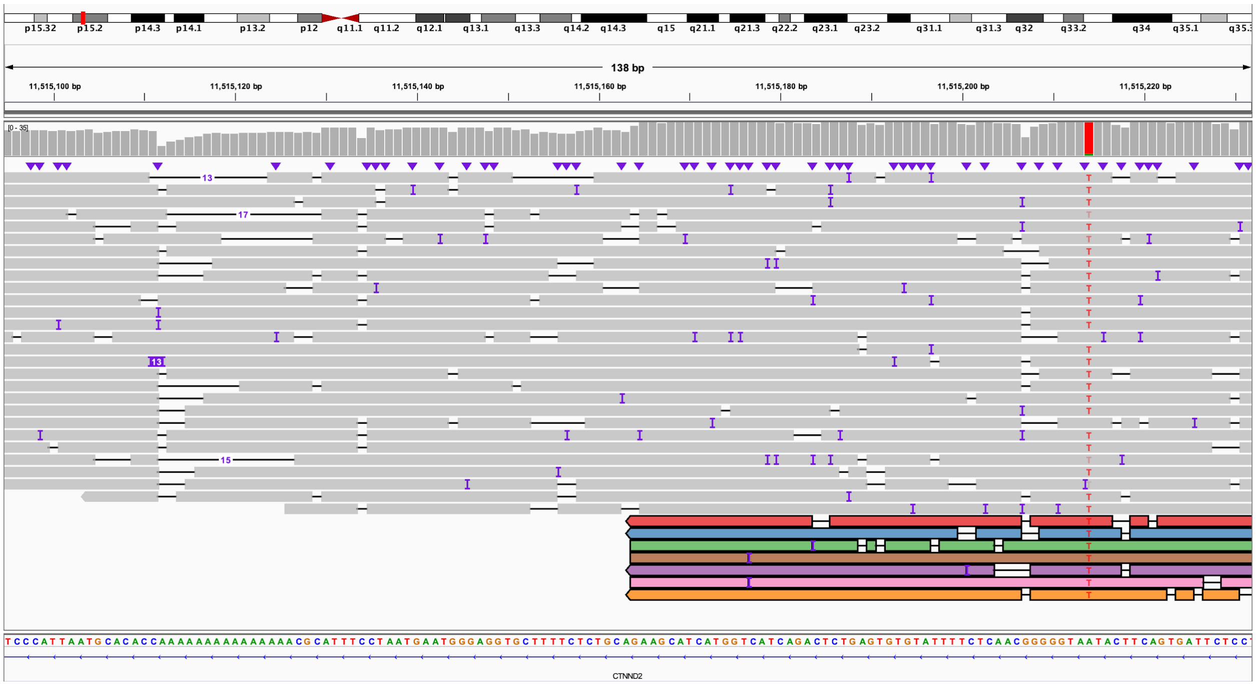

C:

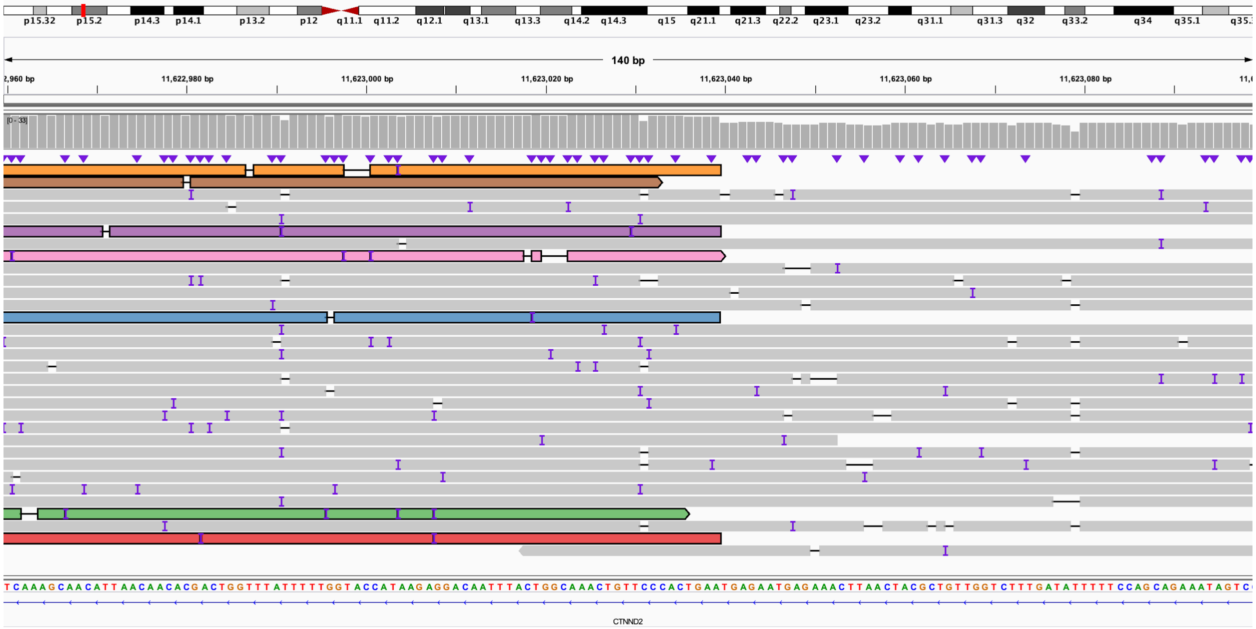

p15.32 p15.2 p14.3 p14.1 p13.2 p12 q11.1 q11.2 q12.1 q13.1 q13.3 q14.2 q14.3 q15 q21.1 q21.3 q22.2 q23.1 q23.2 q31.1 q31.3 q32 q33.2 q34 q35.1
140 bp
11,622,980 bp
11,623,000 bp
11,623,020 bp
11,623,040 bp
11,623,060 bp
11,623,080 bp
CTNND2

**Figure S12. S046, known unbalanced translocation between chromosomes 4 and 15.**

**A.** Coverage of chromosome 4 target region showing a distal deletion. **B.** Coverage of chromosome 15 target region showing a distal gain. **C.** IGV view of reads spanning the translocation breakpoint. Reads are colored to correspond to reads in D. IGV view is of chr4:185,118,270-185,118,370. **D.** View of reads spanning the translocation breakpoint on chromosome 15. IGV view is of chr15:92,684,498-92,684,602.

A:

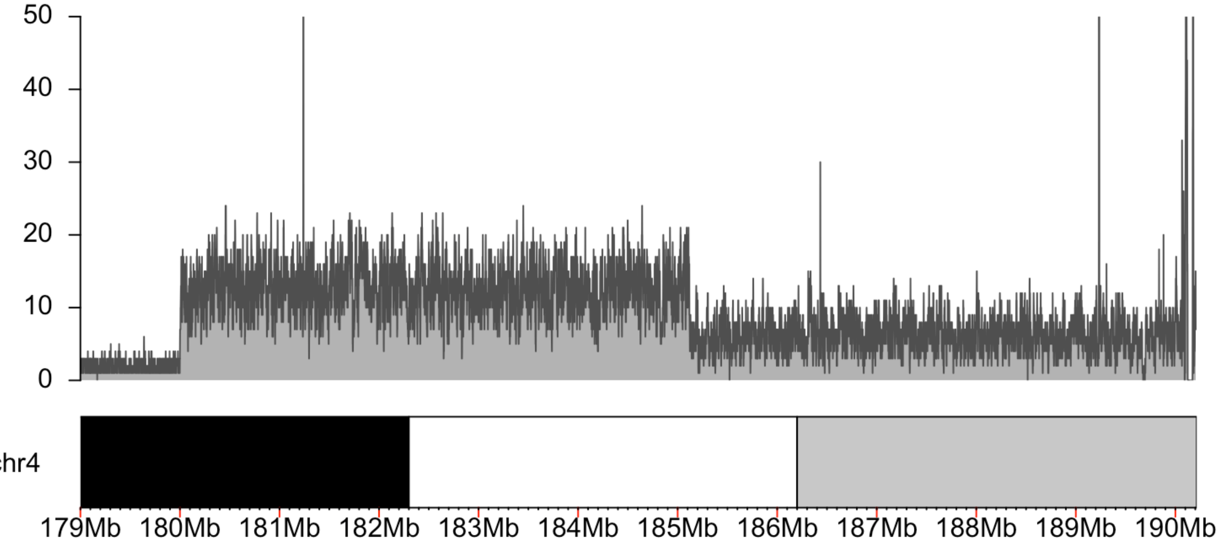

B:

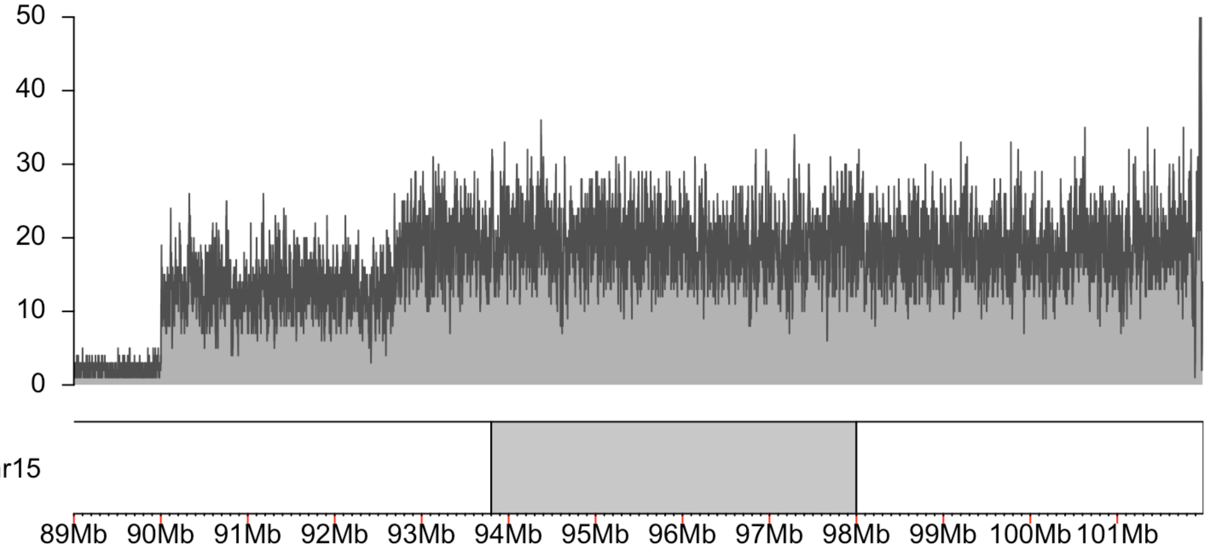

C:

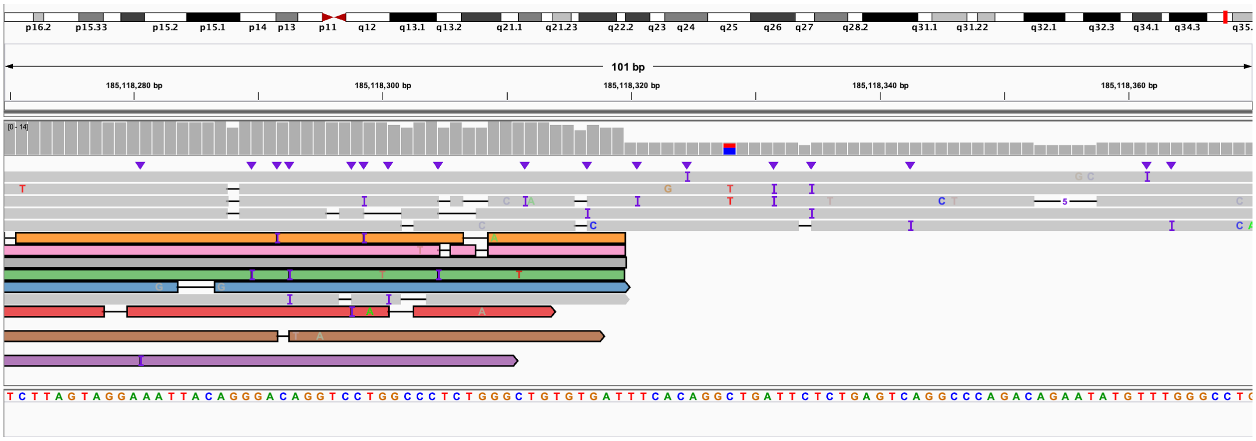
p16.2 p15.33 p15.2 p15.1 p14 p13 p11 q12 q13.1 q13.2 q21.1 q21.23 q22.2 q23 q24 q25 q26 q27 q28.2 q31.1 q31.22 q32.1 q32.3 q34.1 q34.3
101 bp
185,118,280 bp
185,118,300 bp
185,118,320 bp
185,118,340 bp
185,118,360 bp
[0 - 14]

D:

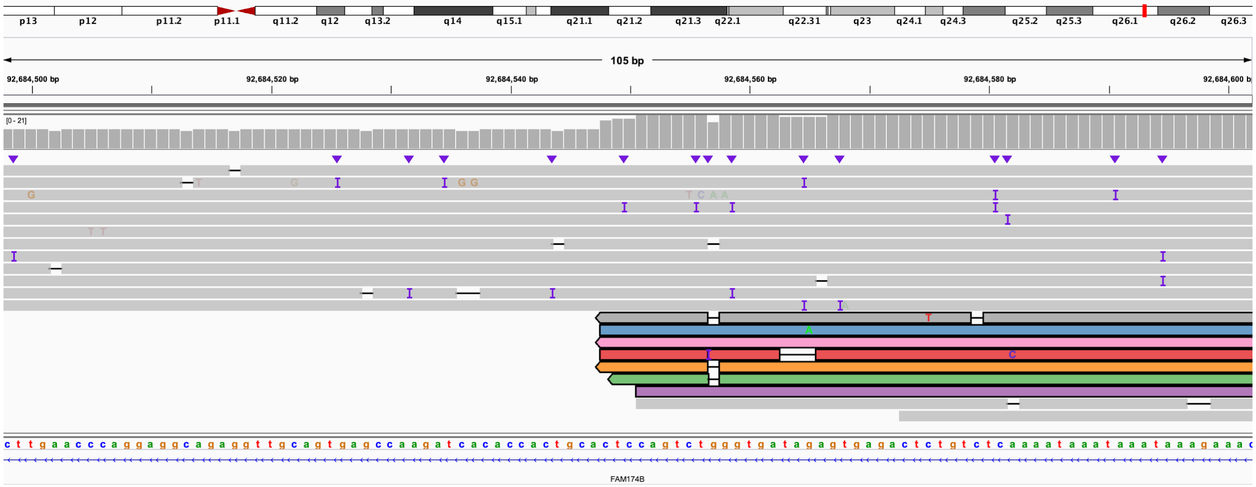
p13 p12 p11.2 p11.1 q11.2 q12 q13.2 q14 q15.1 q21.1 q21.2 q21.3 q22.1 q22.31 q23 q24.1 q24.3 q25.2 q25.3 q26.1 q26.2 q26.3
105 bp
92,684,500 bp
92,684,520 bp
92,684,540 bp
92,684,560 bp
92,684,580 bp
[0 - 21]
FAM174B

**Figure S13. S011 (*ATXN3* and *ATXN8OS*), evaluation of repeat length in patient sample.**

For both A and B, reads aligned to the reference using minimap2 with default parameters is the top panel and reads aligned with modified parameters as outlined in the methods is the bottom panel. Annotation by Tandem Repeats Finder is at the bottom of each screenshot. **A.** T-LRS of an individual known to carry a heterozygous repeat expansion in *ATXN3*. Three reads carry the expansion and span the region. IGV view of chr14:92,070,974-92,071,079. **B.** This individual was also known to carry a heterozygous expansion in *ATXNOS8*. In the minimap2 alignment with default parameters, a single read carries the full expansion and spans the region (top panel), but with modified parameters four reads can be found spanning the interval (bottom panel). IGV window is chr13:70,139,346-70,139,436.

A:

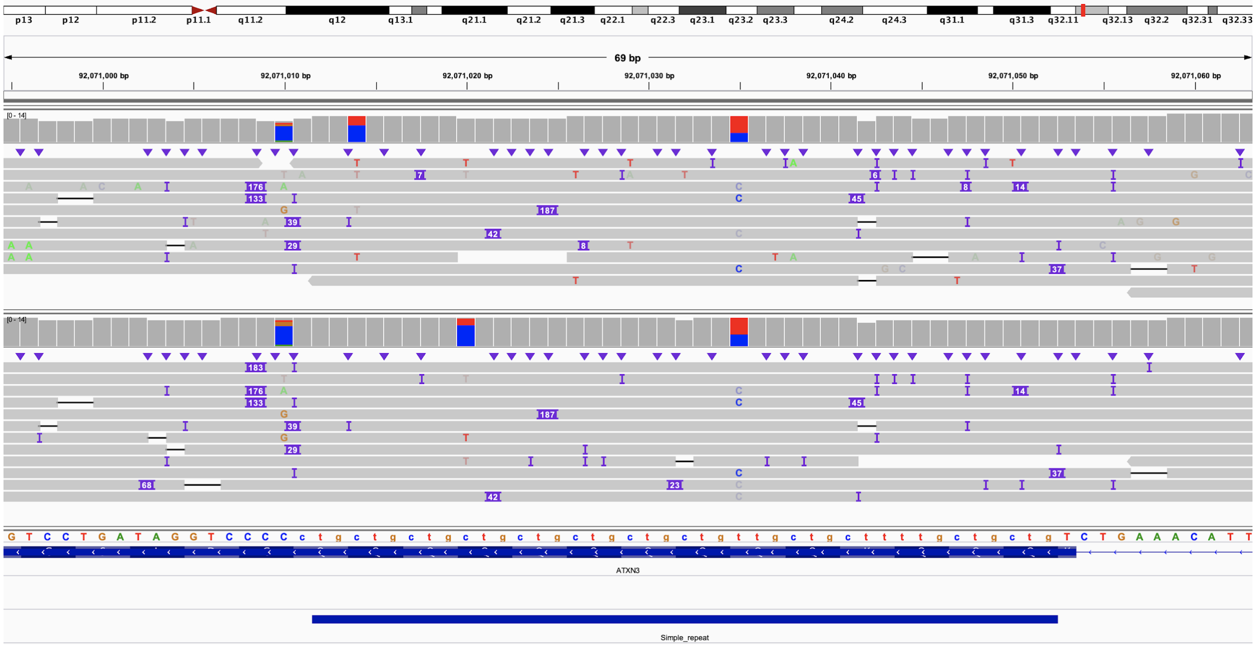

B:

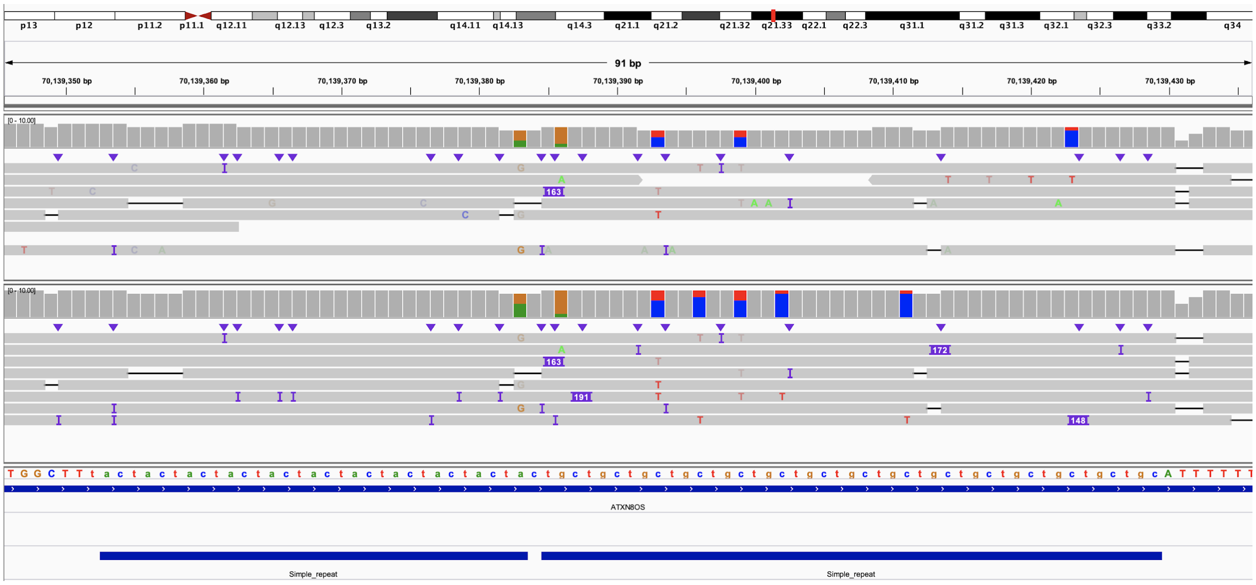

### Figure S14. S039 (*FMR1*), evaluation of repeat length and methylation status.

**A.** Screenshot of CGG repeat expansion in 5' UTR of *FMR1* from a patient known to carry an expansion. DNA was obtained from Coriell (Table S1). Top panel represents minimap2 run with default parameters; bottom panel is minimap2 run with parameters meant to reduce split reads. In the bottom panel, two reads with insert sizes of approximately 1,350 bp can be seen that are not present in the top panel. IGV view is chrX:147,911,949-147,912,149. **B.** Methylation analysis of 5' UTR of *FMR1* reveals that the region around the CGG expansion is no longer methylated. The region shown is larger than in panel A. IGV view is chrX:147,911,502-147,912,719.

A:

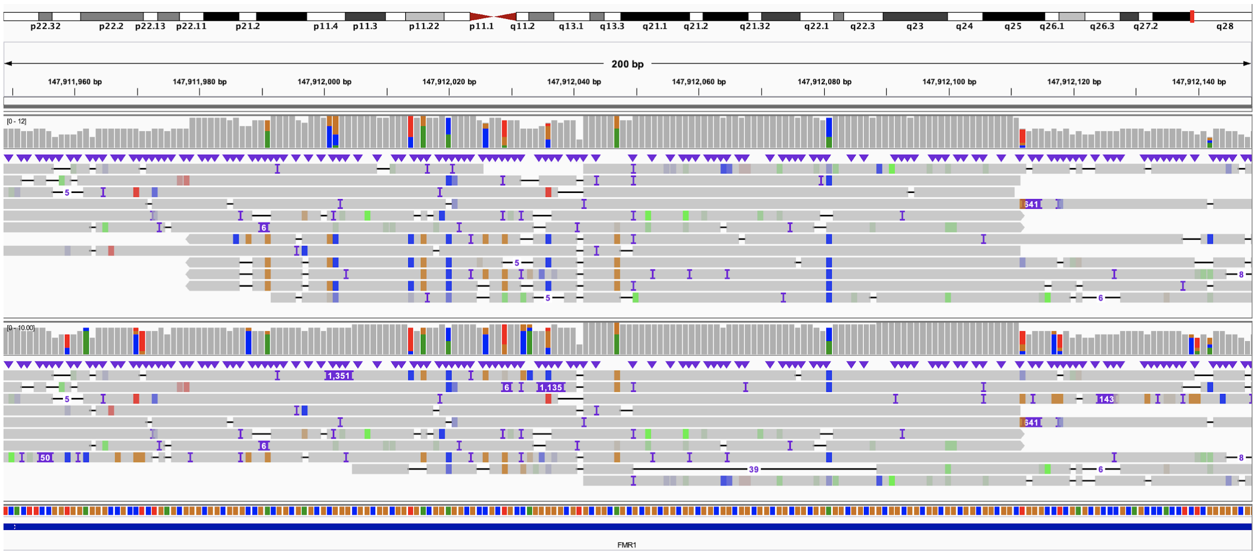

B:

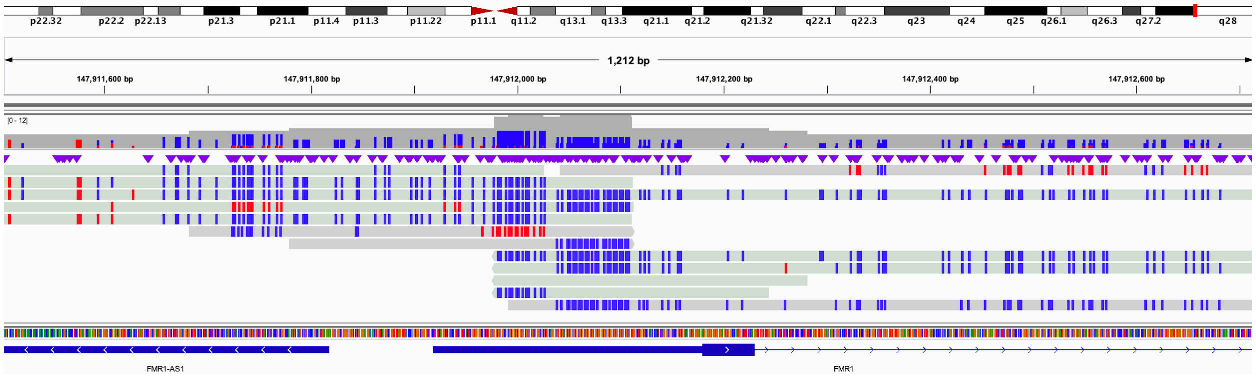

### Figure S15. S040 (*FXN*), known repeat expansion.

IGV view of a sample from an individual with a heterozygous expansion in *FXN*. Sample is from a cell line obtained from Coriell (Table S1). Top panel is reads aligned with minimap2 using default parameters; bottom panel is parameters that reduce the number of split reads. IGV view is from chr9:69,037,240-69,037,319.

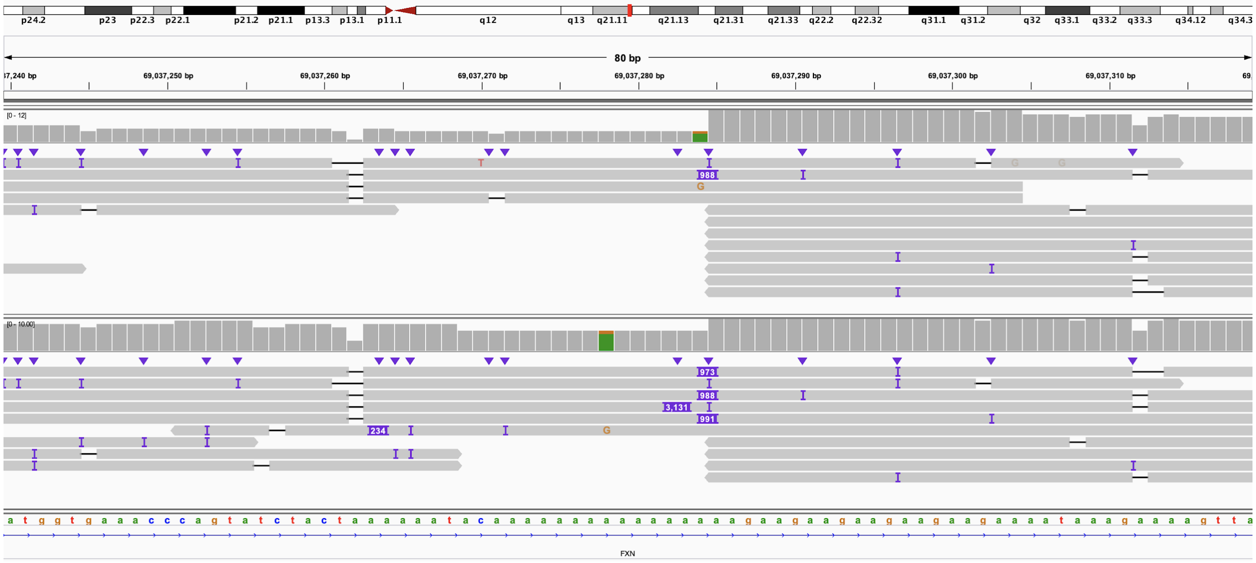

### Figure S16. S041 (*FXN*), known repeat expansion.

IGV view of a case known to be heterozygous for two different repeat expansions in *FXN*; DNA was obtained from Coriell (Table S1). Reads aligned with minimap2 and default parameters are in the top panel; reads aligned with minimap2 and parameters meant to decrease the number of reads broken by the aligner are in the bottom panel. IGV view is from chr9:69,037,243-69,037,334.

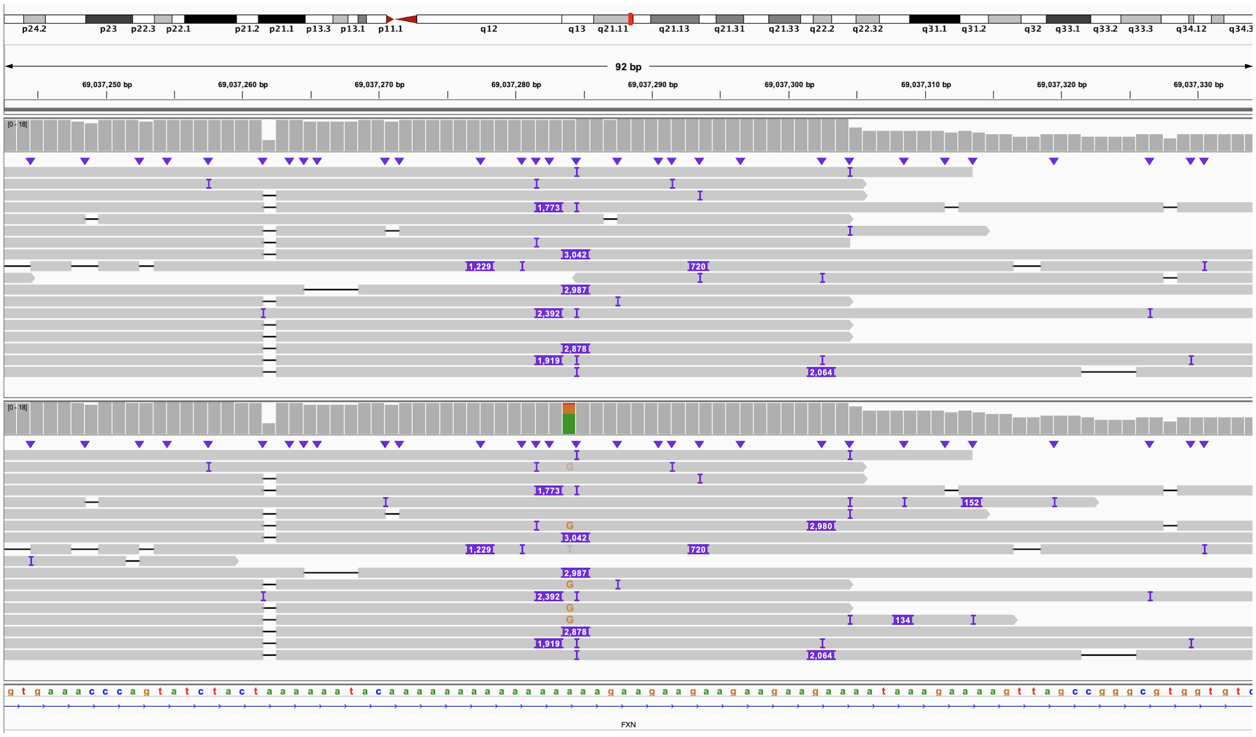

**Figure S17. 04-01, 04-02, and 04-03 (*XYLT1*), evaluation of repeat length in family 04 from LaCroix *et al*. 2019.**

**A.** Screenshot of expansion in 5' UTR of *XYLT1* in family 04 from (LaCroix et al. 2019). The proband (top) inherited a permutation allele from his mother and a *de novo* deletion from his father; the mother (middle) has a wild-type allele and permutation allele, while the father has two wild-type alleles (bottom). **B.** Methylation was called and assigned to each read and shows that all reads in the proband are methylated (red), while most in the mother are unmethylated (blue), and both in the father are unmethylated. IGV view from chr16:17,470,696-17,471,299. B. IGV view from chr16:17,470,288-17,471,705.

A:

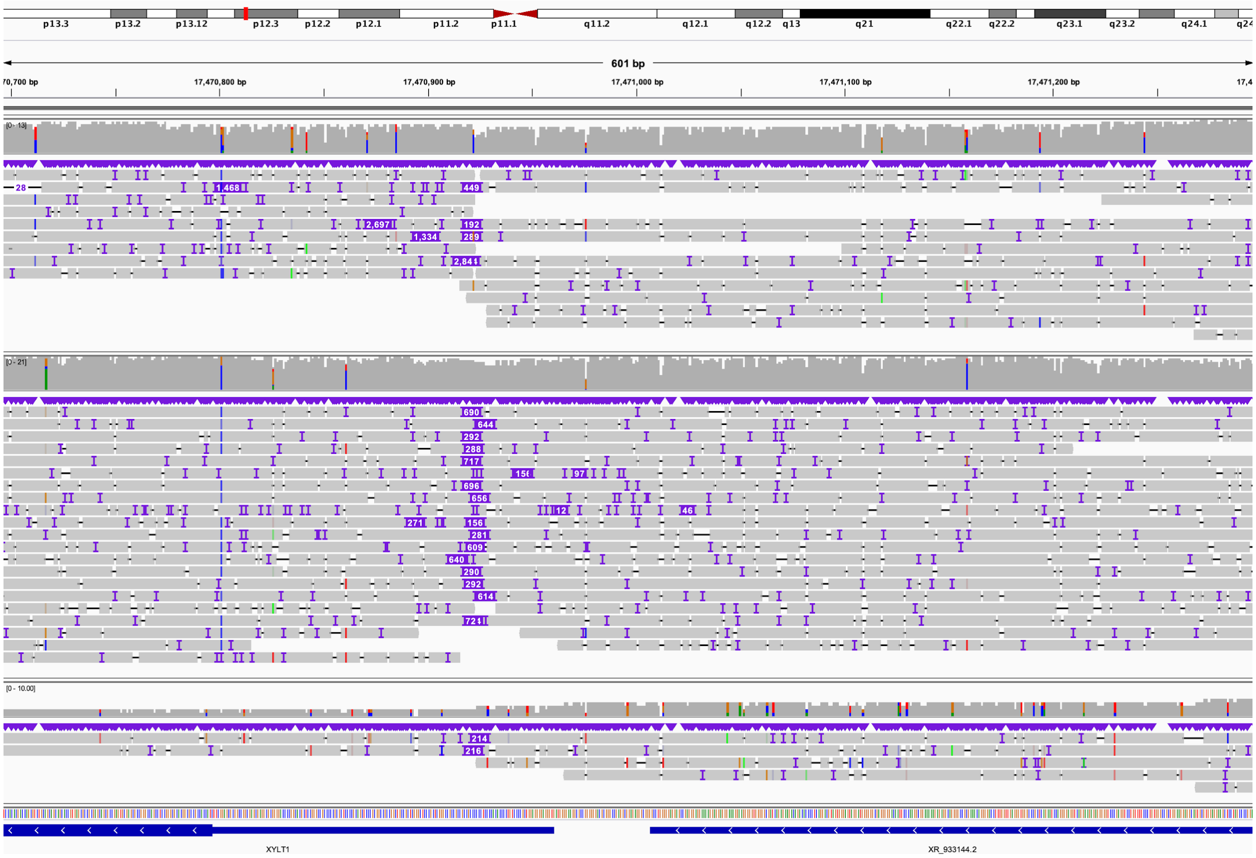

B:

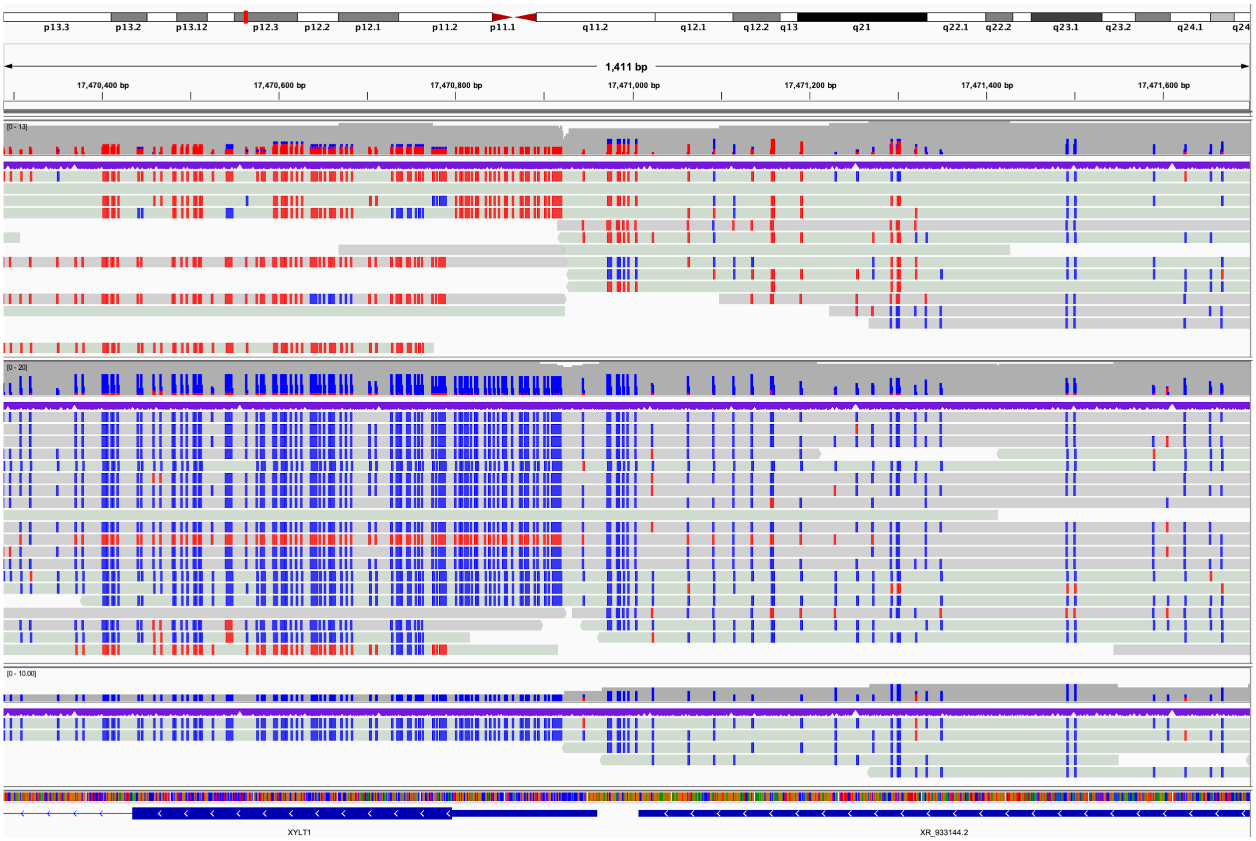
p13.3
p13.2
p13.12
p12.3
p12.2
p12.1
p11.2
p11.1
q11.2
q12.1
q12.2
q13
q21
q22.1
q22.2
q23.1
q23.2
q24.1
q24
1,411 bp
17,470,400 bp
17,470,600 bp
17,470,800 bp
17,471,000 bp
17,471,200 bp
17,471,400 bp
17,471,600 bp
[0 - 13]
[0 - 20]
[0 - 10.00]
XYLT1
XR_933144.2

**Figure S18. 06-01, 06-02, and 06-03 (*XYLT1*), evaluation of repeat length in family 06 from LaCroix *et al*. 2019.**

**A.** IGV view of proband (top), mother (middle), and father (bottom) from family 06 from (LaCroix et al. 2019). Only a single read was recovered spanning the GGC repeat in both the mother and father. IGV view is from chr16:17,470,696-17,471,299. **B.** IGV view of reads converted to show methylation status; order is the same as in A but the view is slightly larger to show methylation of the first exon of *XYLT1* in the proband. IGV view from chr16:17,470,288-17,471,705.

A:

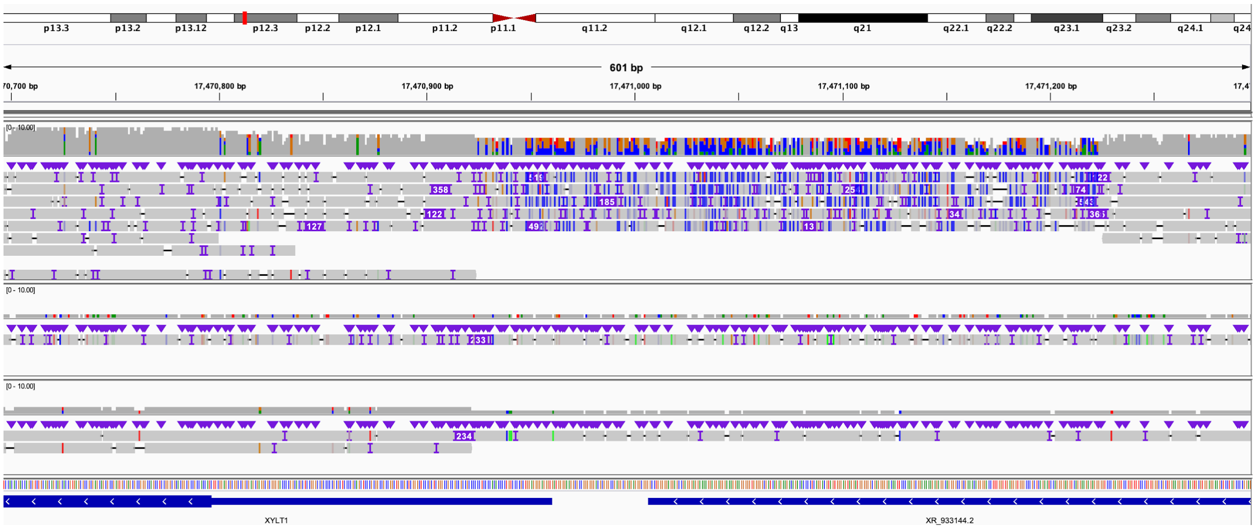

B:

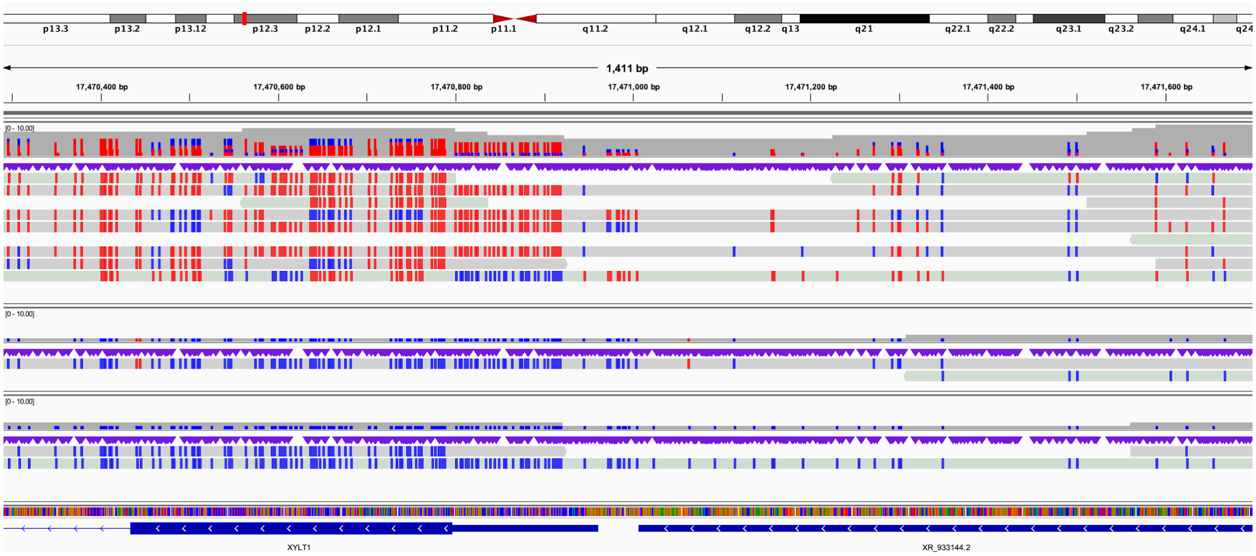

**Figure S19. S014, three noncontiguous deletions of chromosome 6 identified by CMA.**

**A.** Coverage of chromosome 6 target region. **B–H.** Fragments are shown in linked groups as shown in Figure 1A. Colors within groups represent reads that span the breakpoints. Details of any genes bisected and the region the screenshot is from accompany each fragment.

A:

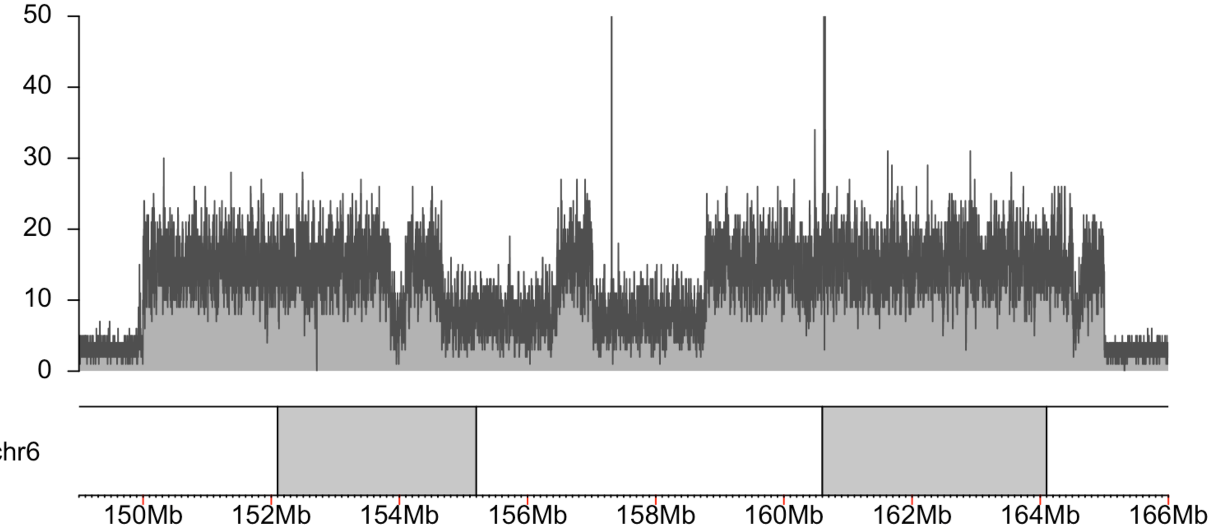

B: 3' end of fragment A (IGV: chr6:153,854,885-153,854,924) connects to 5' end of fragment E (IGV: chr6:154,588,780-154,588,819).

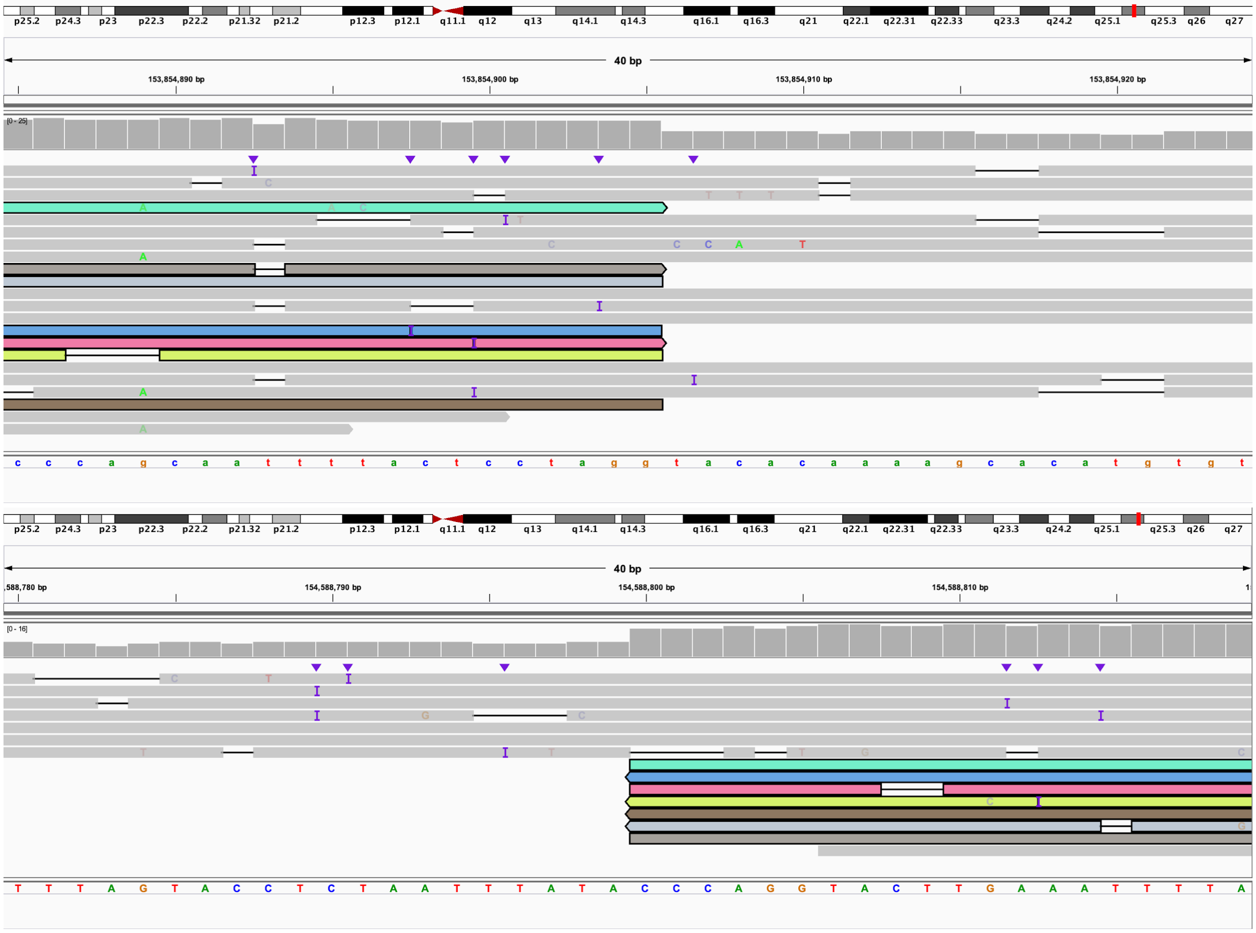

C: 3' end of fragment E (IGV: chr6:154,661,260-154,661,299) connects to 3’ end of fragment C (IGV: chr6:154,587,128-154,587,168).

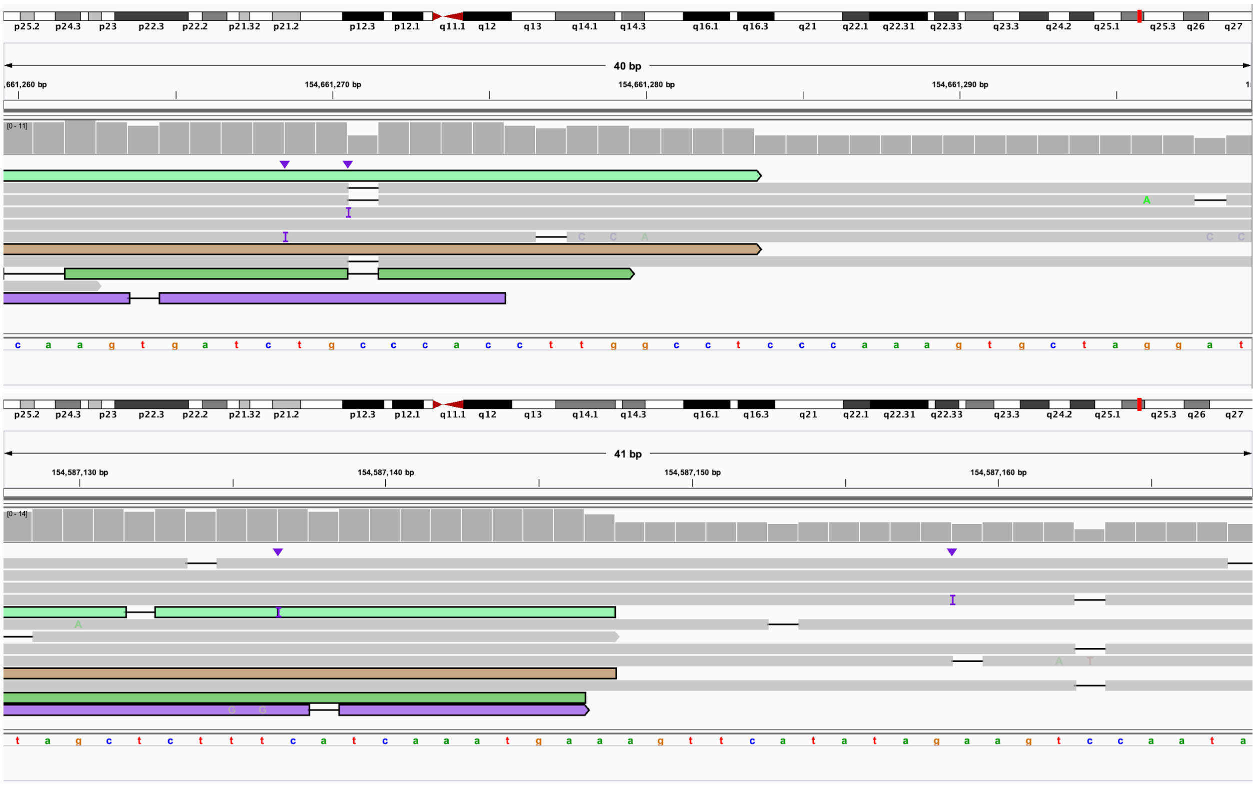

D: 5' end of fragment C bisects *OPRM1* (IGV: chr6:154,095,825-154,095,864) and connects to 5' end of fragment I (IGV: chr6:156,464,482-156,464,522).

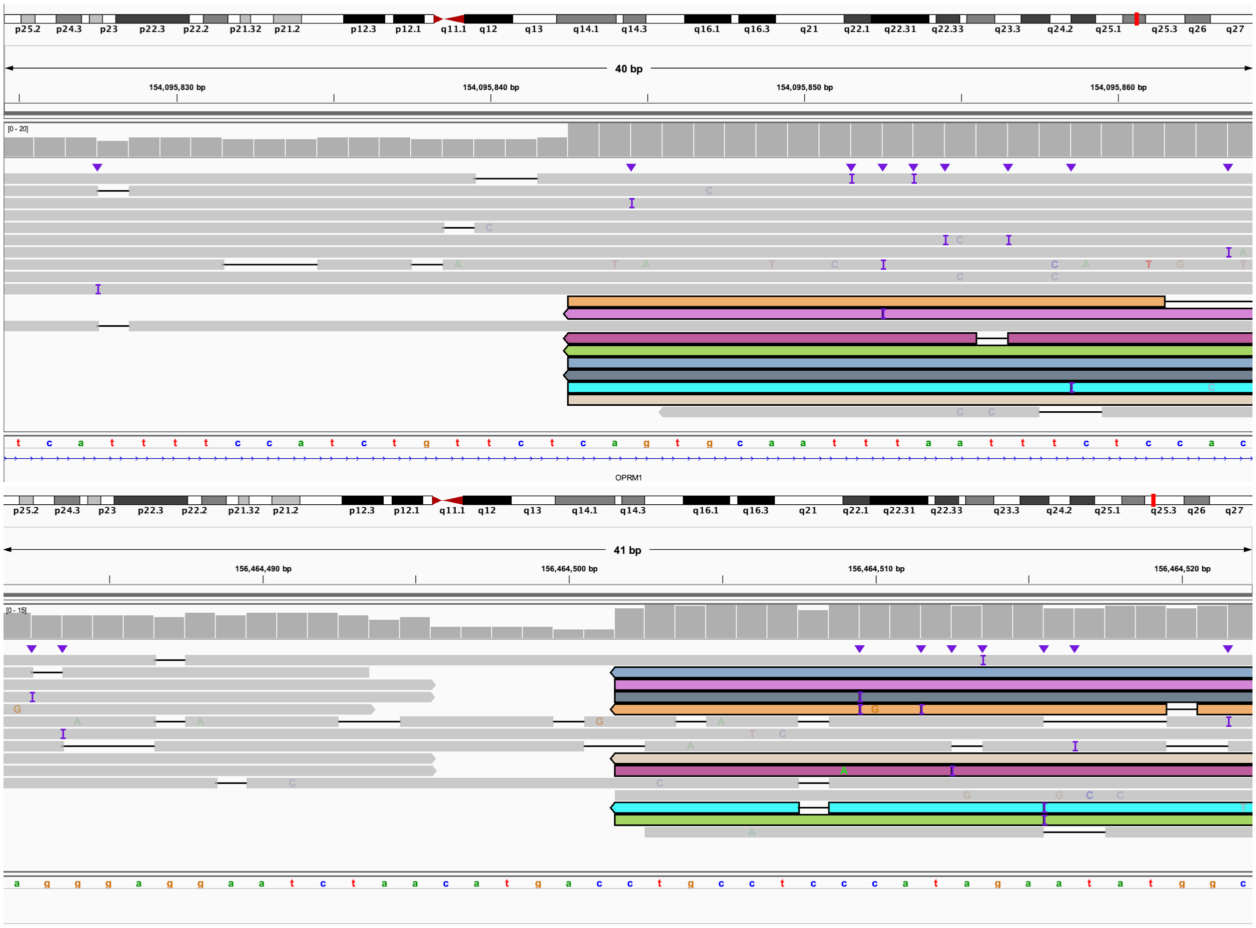

E: 3' end of fragment I bisects *ARID1B* (IGV: chr6:157,015,094-157,015,151) and connects to 5' end of fragment H (IGV: chr6:156,460,095-156,460,134).

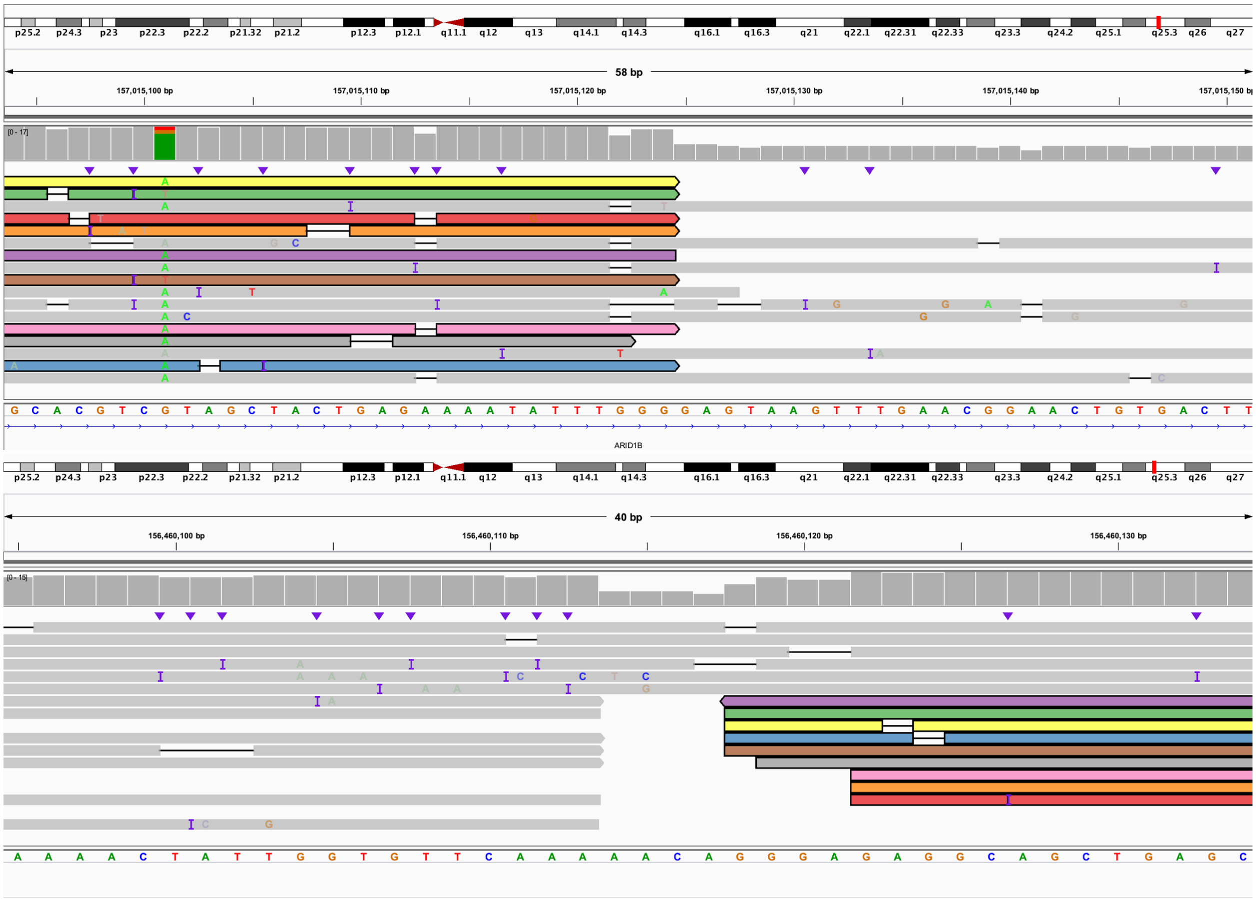

F: 3' end of fragment H (IGV: chr6:156,464,478-156,464,518) connects to 5' end of fragment K and bisects *EZR* (IGV: chr6:158,766,800-158,766,839).

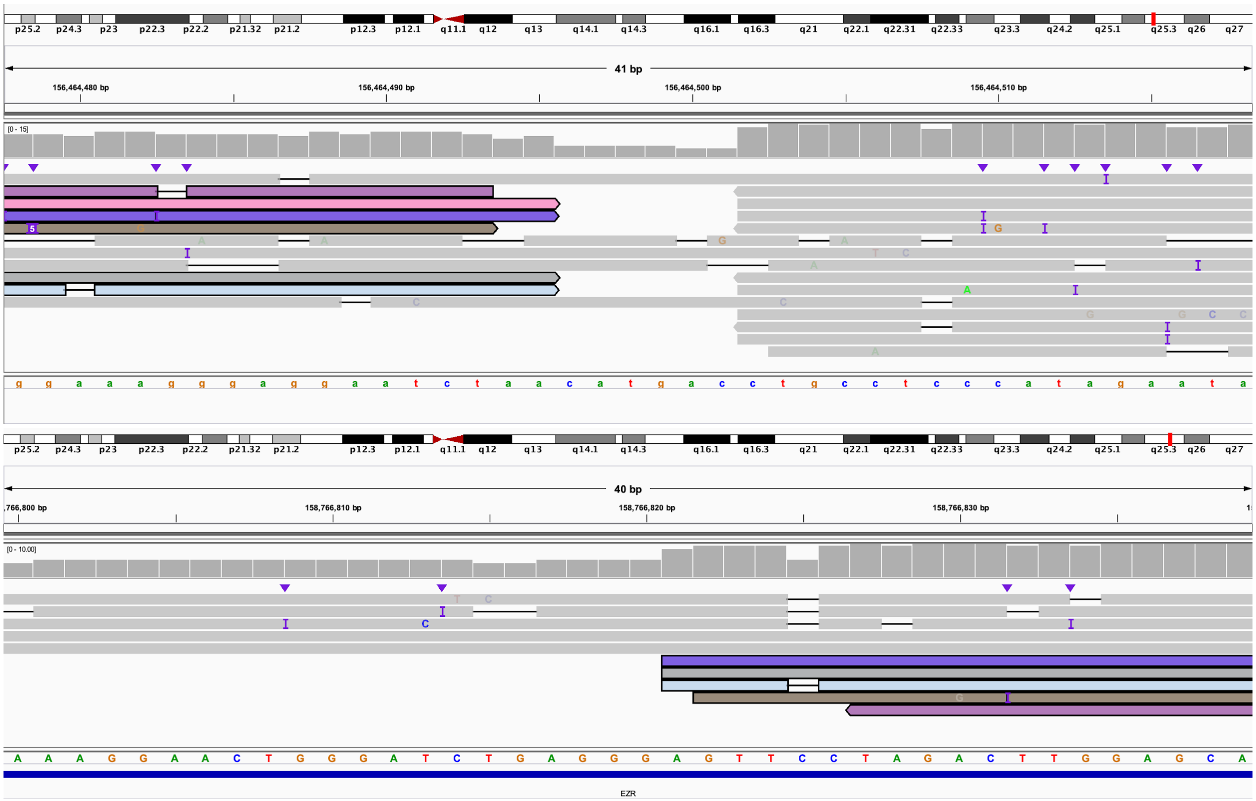

G: 3' end of fragment K bisects *XR_001744460.2* (IGV: chr6:164,500,041-164,500,095) and connects to 3' end of fragment G (IGV: chr6:156,460,095-156,460,134).

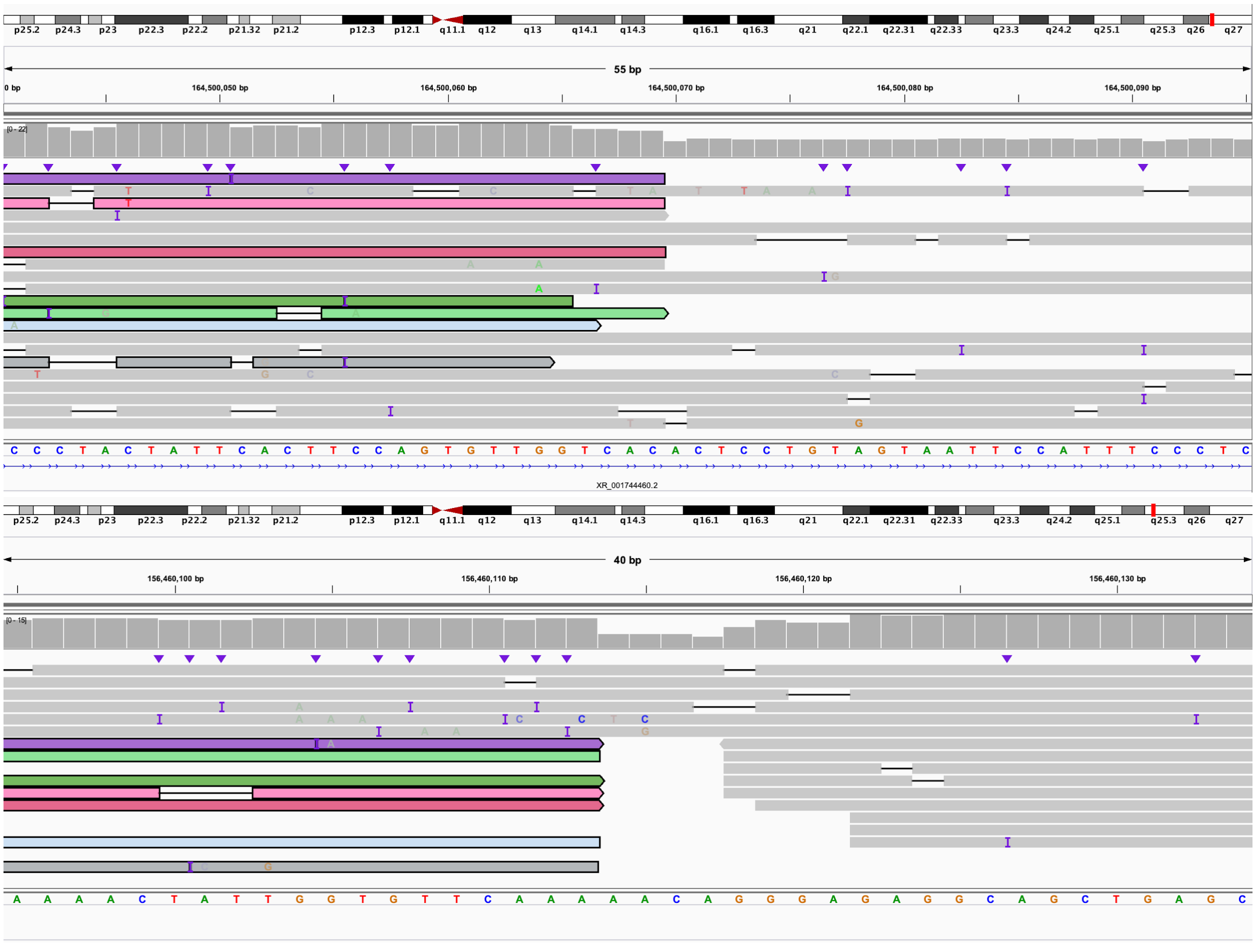

H: 5' end of fragment G (IGV: chr6:156,456,498-156,456,537) connects to 5' end of fragment M (IGV: chr6:164,654,902-164,654,941).

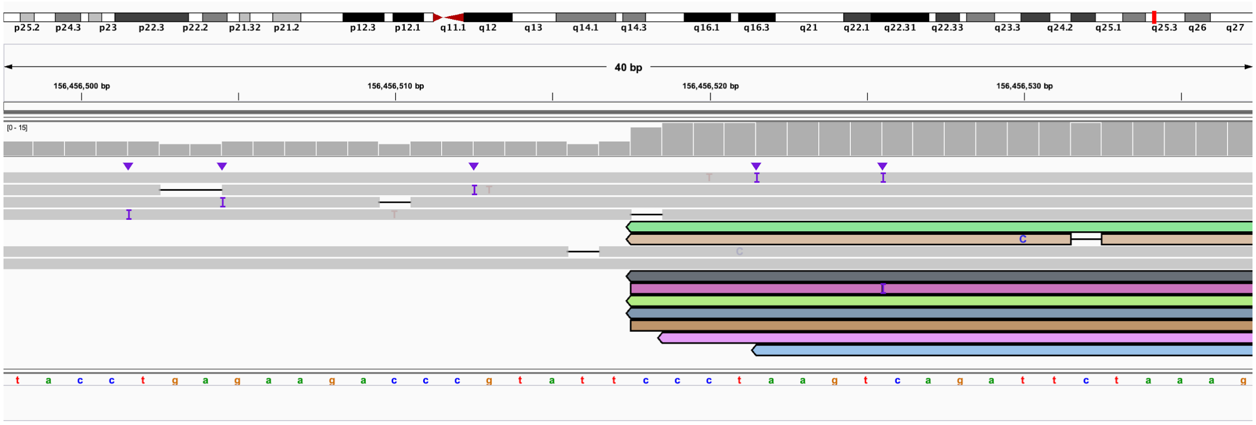

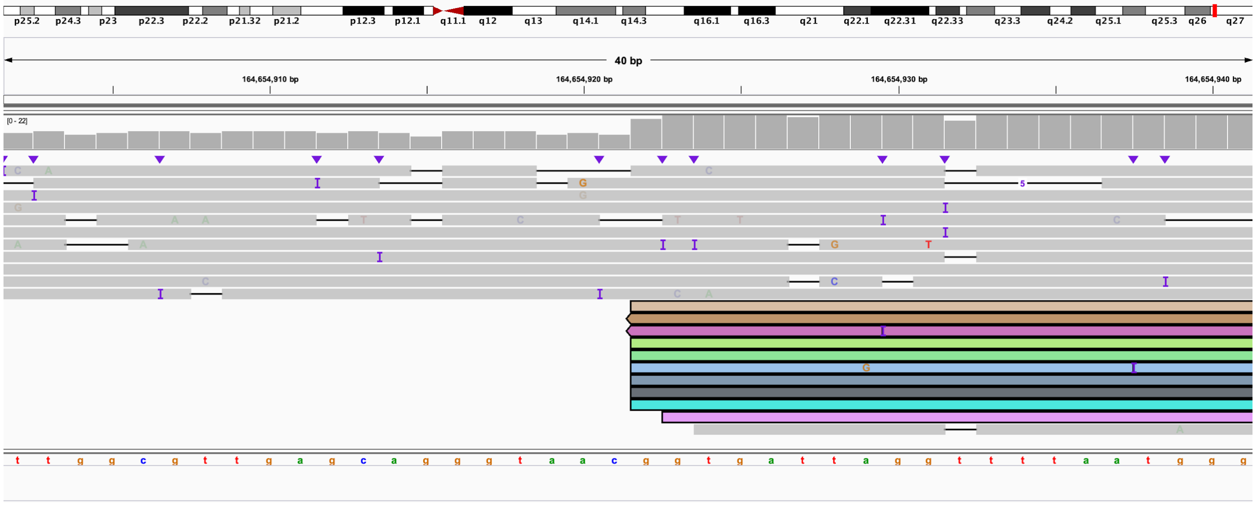

**Figure S20. S020, individual with three deletions identified on array and multiple rearrangements on karyotype.**

**A–H.** Coverage plots from Read Until experiments 1 and 2 as detailed in Table S2. **I.** Karyotype. **J–AG.** IGV images showing rearrangements, inversions, deletions, and translocations. In all panels, Nanopore data is the top track and PacBio HiFi data is the bottom track. Colors are used to show which reads are linked between the two regions of the genome. J represents derivative chromosome 2, K–R represent derivative chromosome 4, S–AA represent derivative chromosome 10, and AB–AG is derivative chromosome 14.

**A:** Coverage of chromosome 2 target region from Read Until experiment 1.

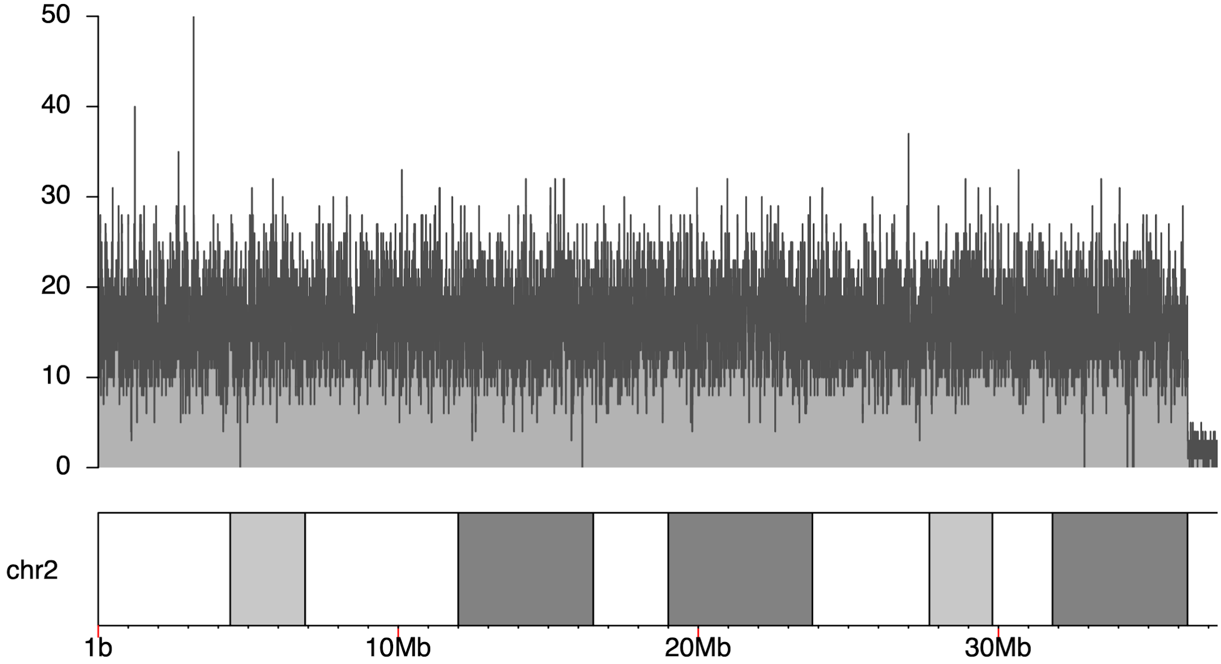

**B:** Coverage of chromosome 4 target region from Read Until experiment 1.

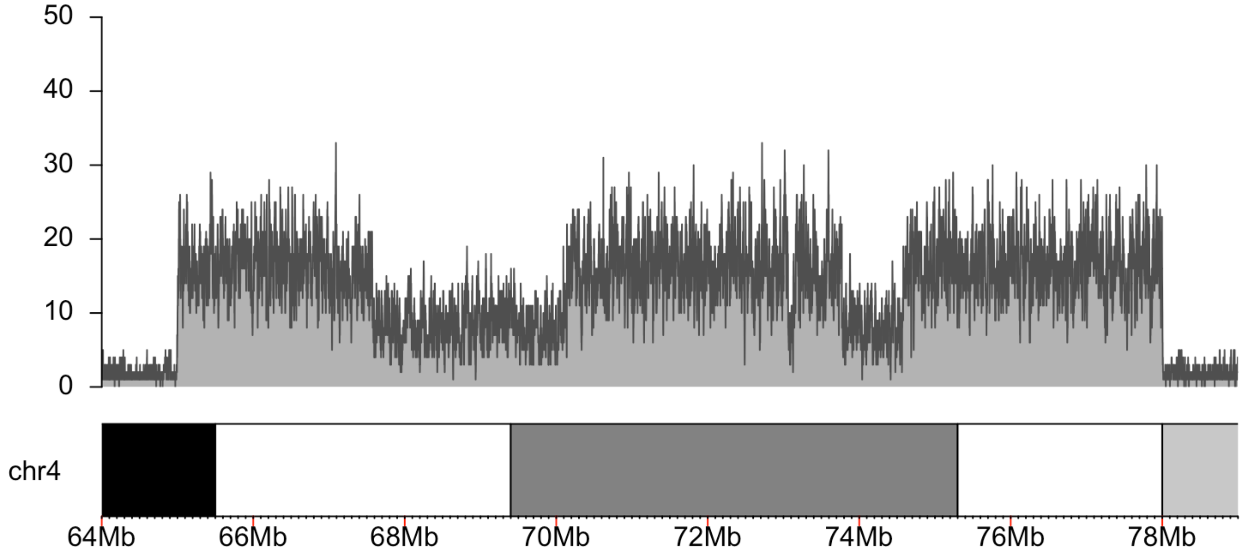

**C:** Coverage of chromosome 10 target region from Read Until experiment 1.

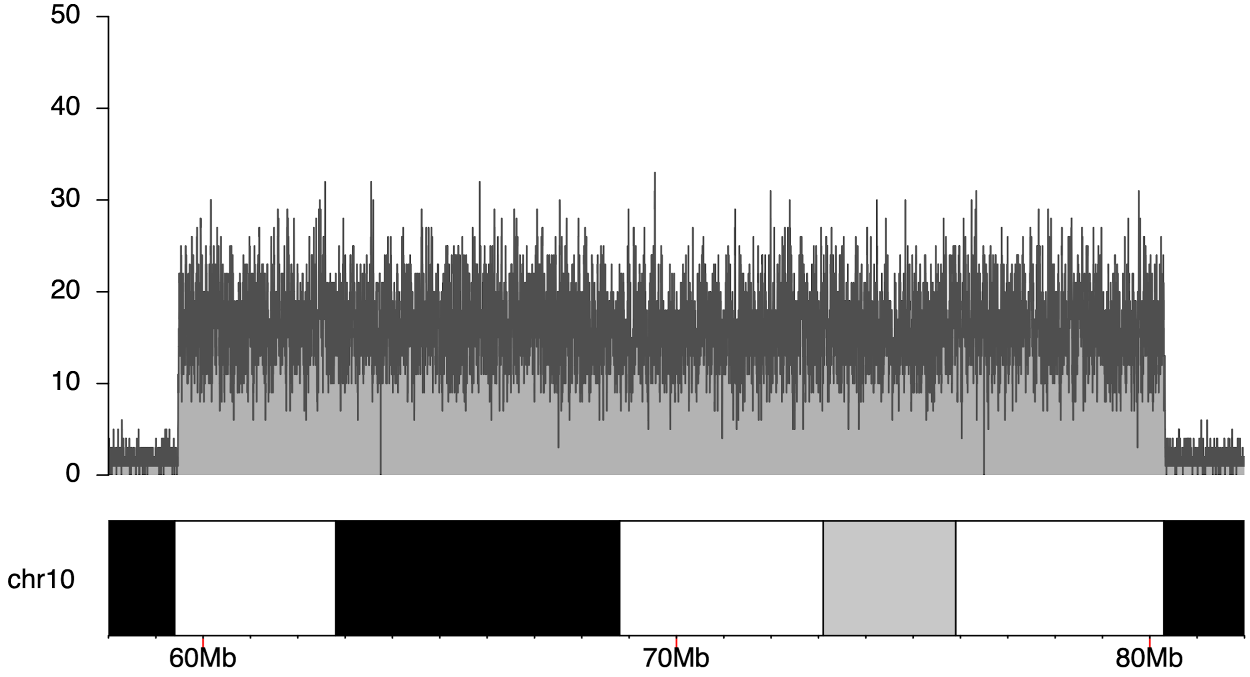

**D:** Coverage of chromosome 14 target region from Read Until experiment 1.

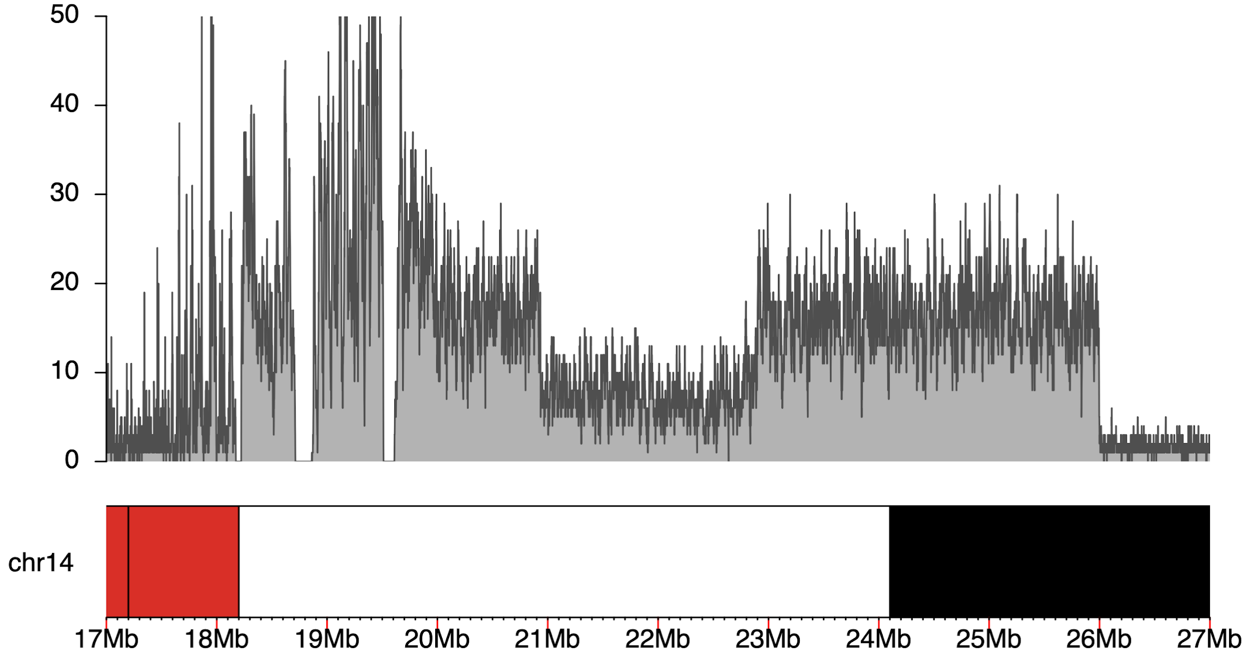

**E:** Coverage of chromosome 10 target region from Read Until experiment 2.

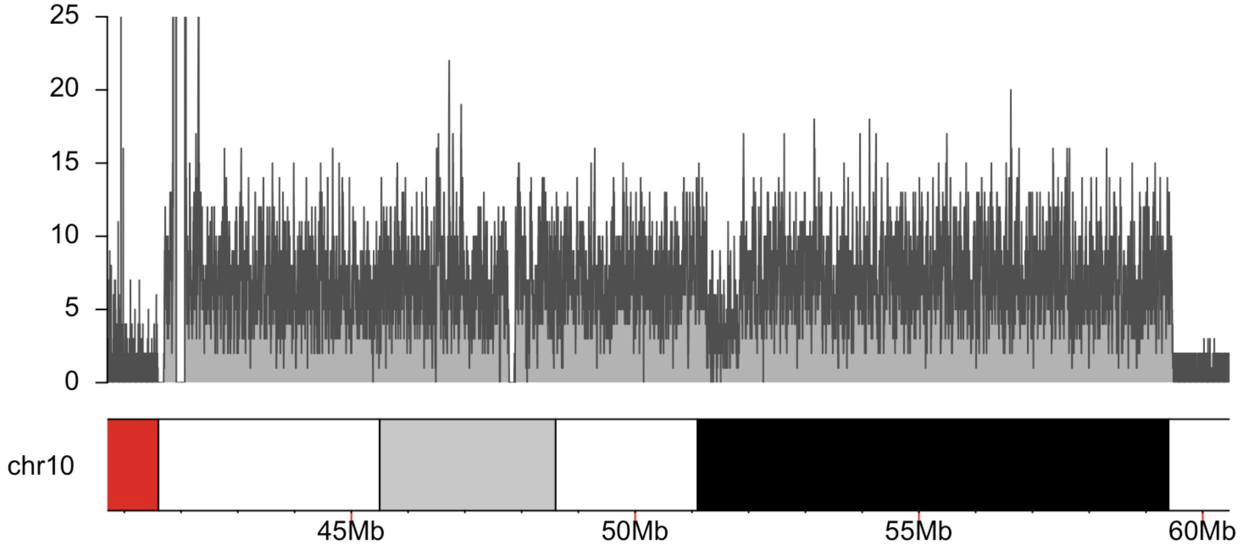

**F:** Coverage of first chromosome 4 target region from Read Until experiment 2.

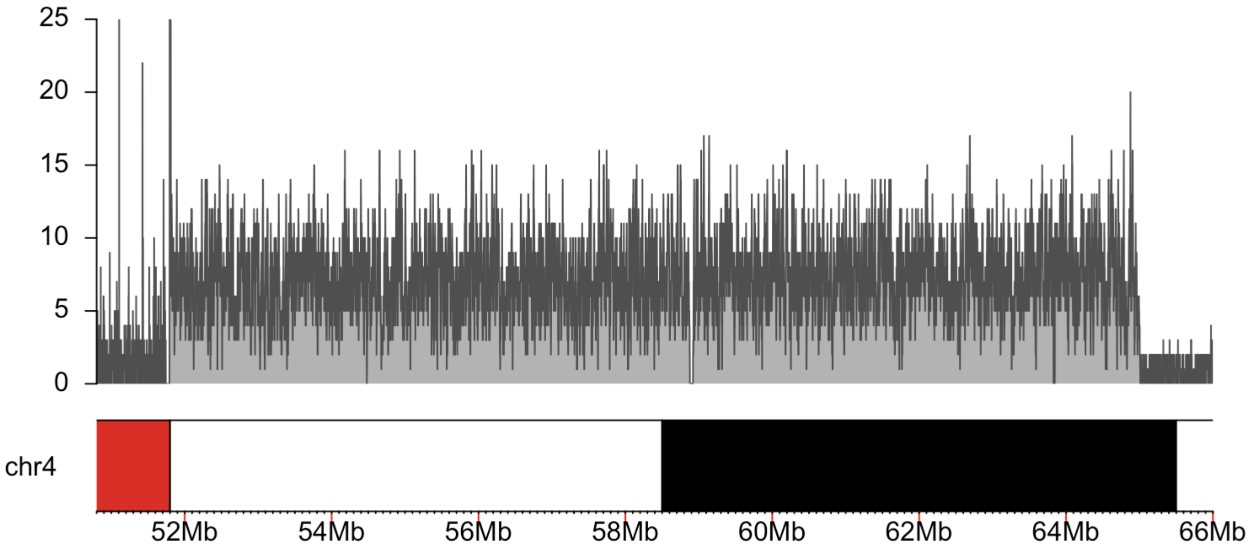

**G:** Coverage of second chromosome 4 target region from Read Until experiment 2.

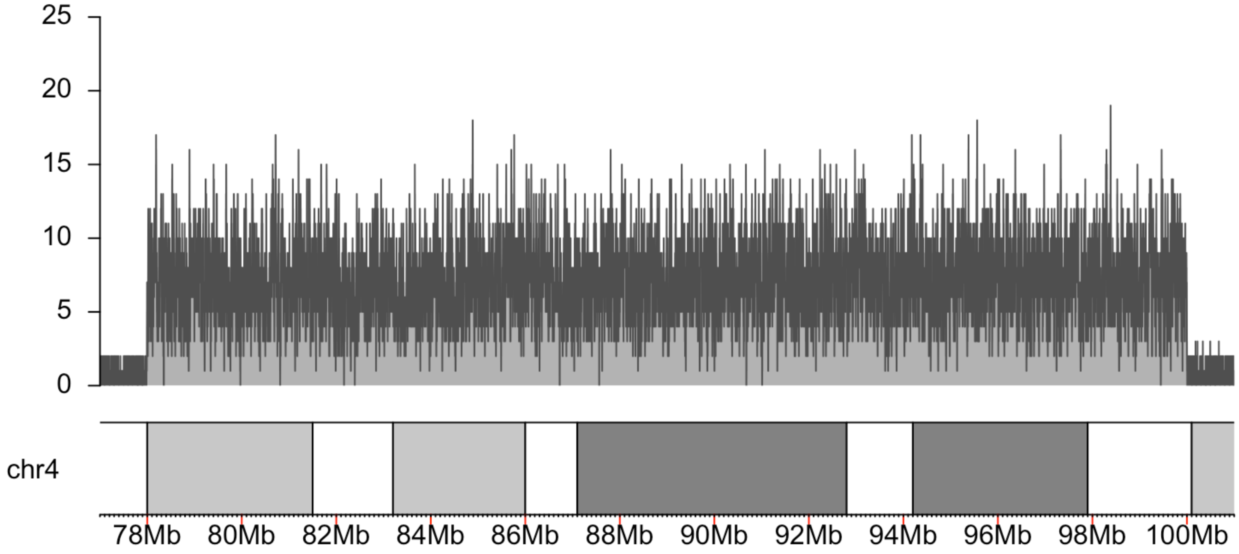

**H:** Coverage of chromosome 14 target region from Read Until experiment 2.

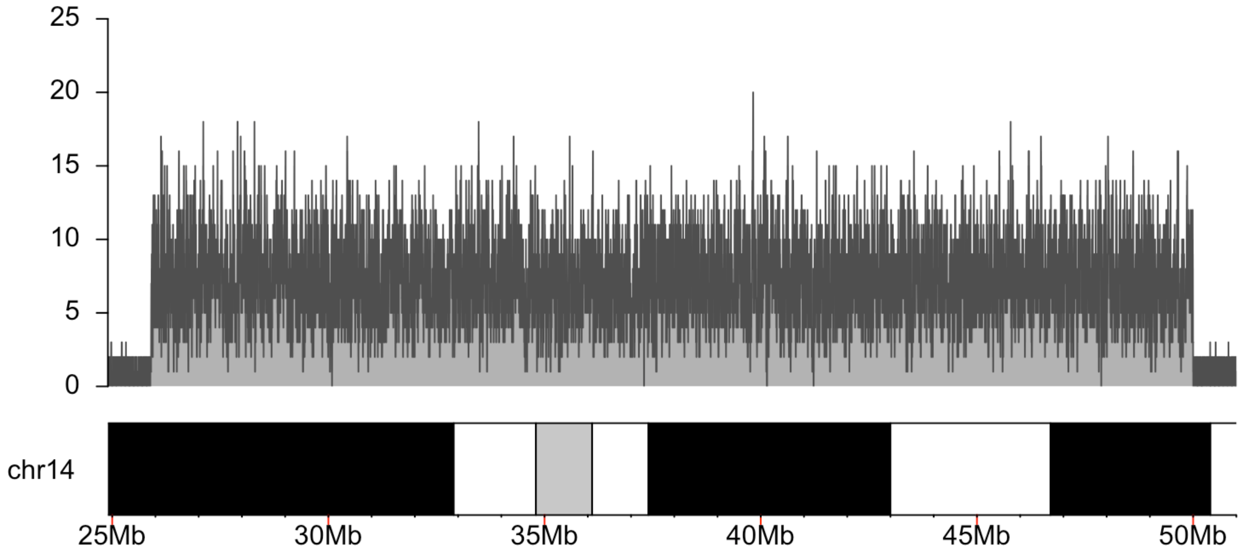

**I:** Karyotype of the case, arrows point to abnormal chromosomes.

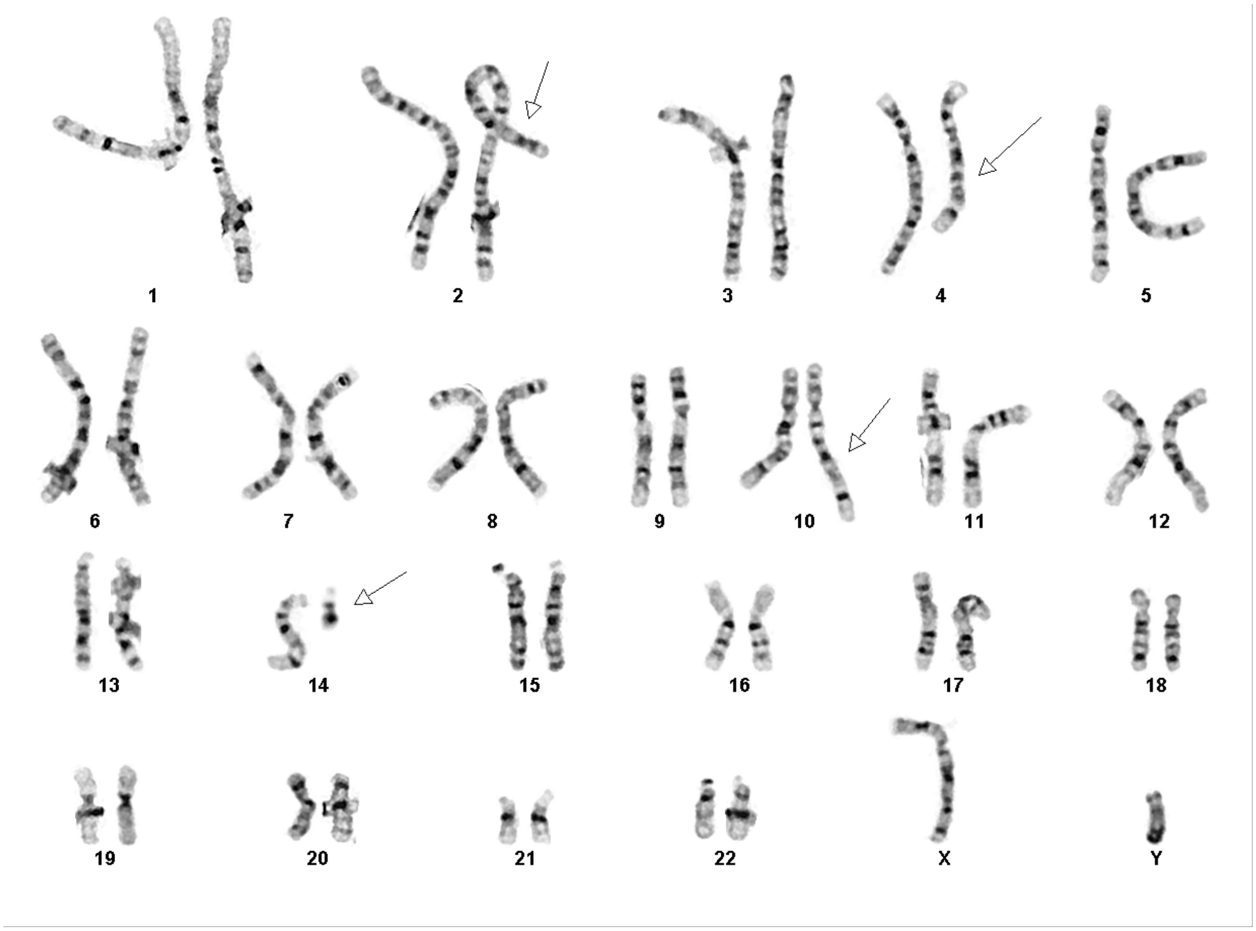

**J.** The 5' end of 2:C (IGV: chr2:16,235,693-16,235,733) is connected to the 5' end of 4:N (IGV: chr4:81,632,551-81,632,590). This is the entirety of derivative chromosome 2.

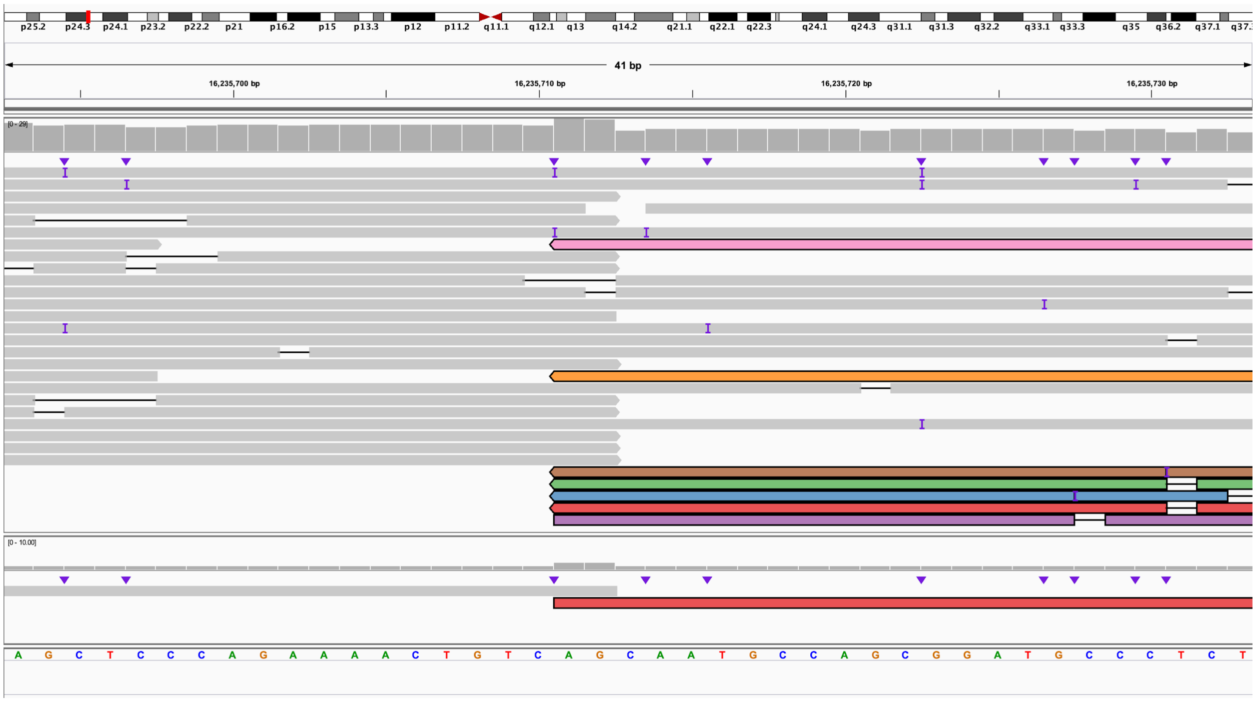

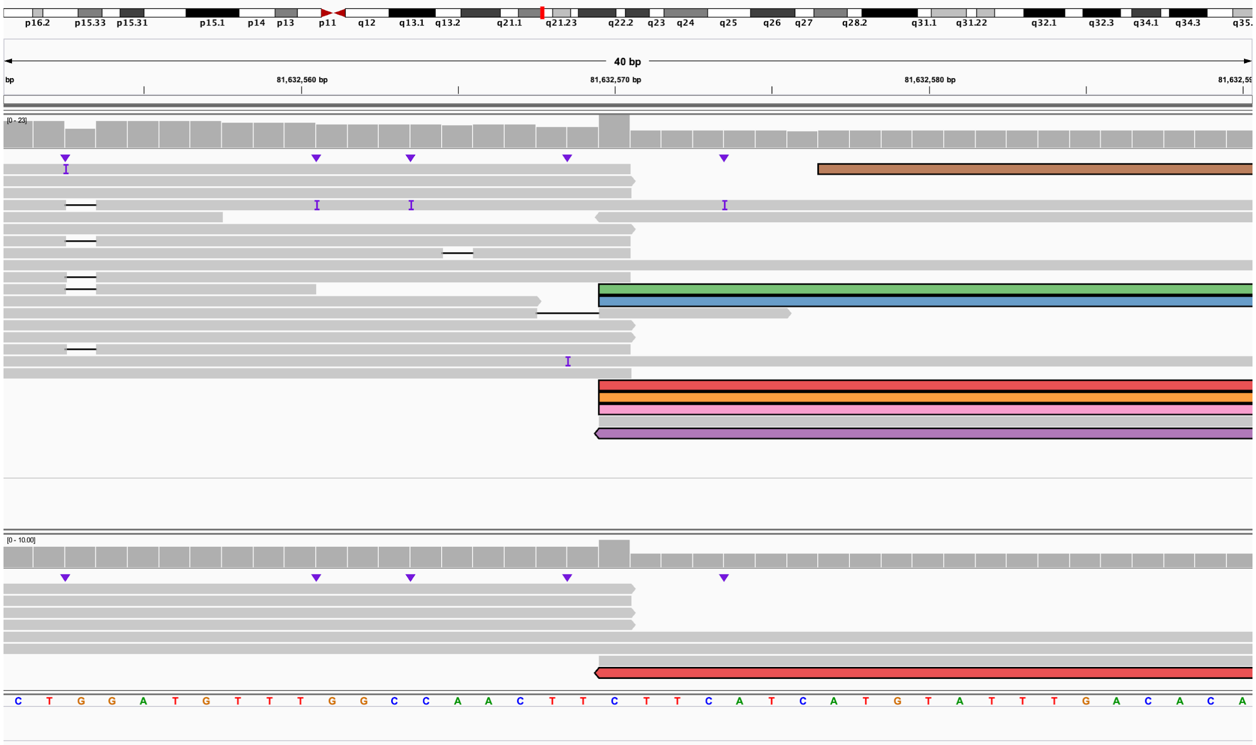

**K.** The 3' end of 4:A bisects *STAP1* (IGV: chr4:67,572,950-67,572,990) and is connected to the 3' end of 4:K and bisects *SEPTIN11* (IGV: chr4:77,012,155-77,012,195). This is the beginning of derivative chromosome 4.

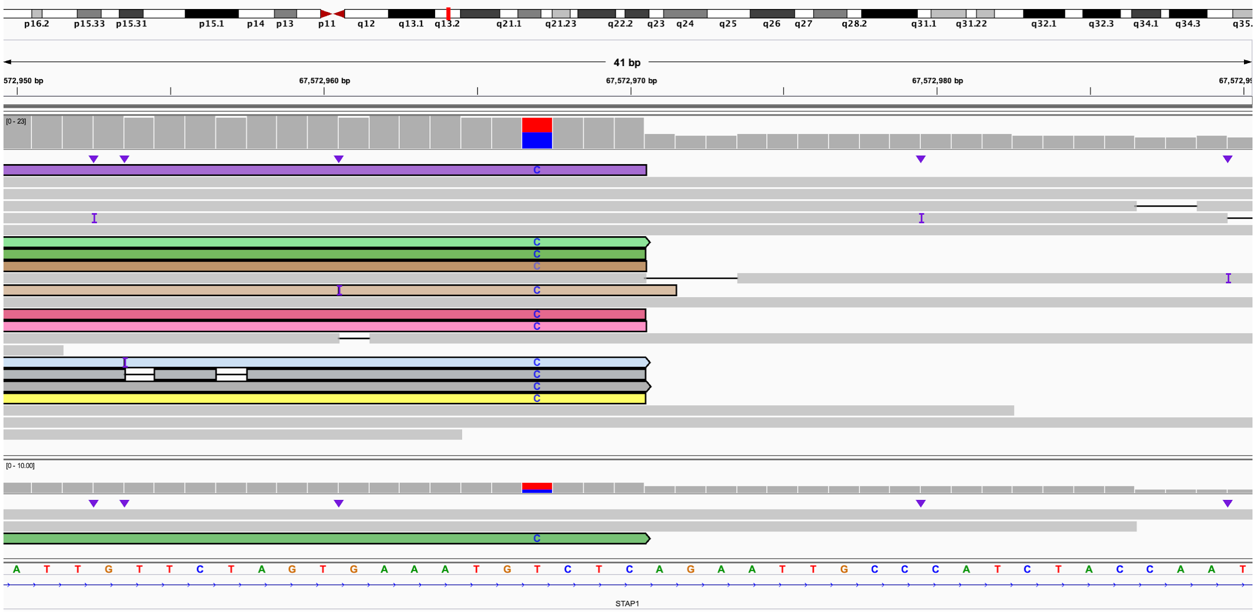

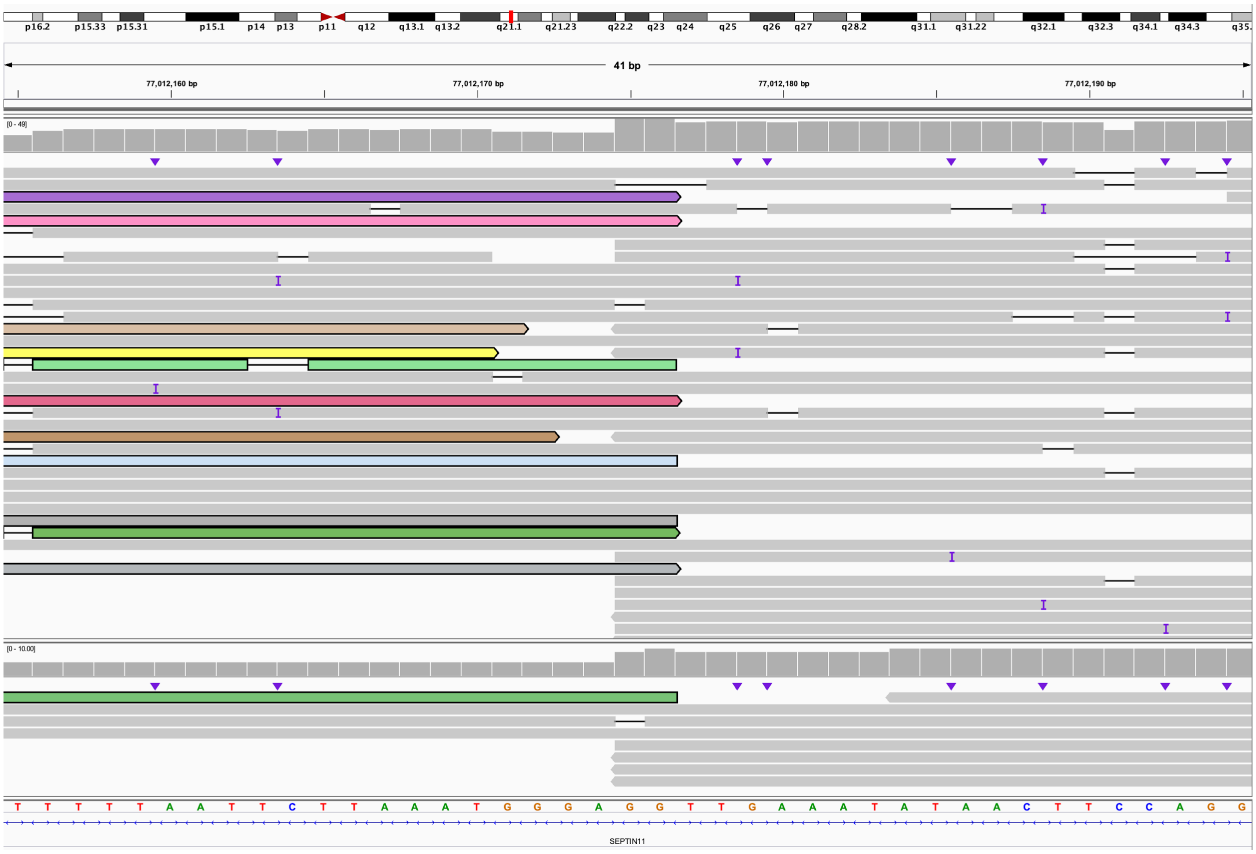

**L.** The 5' end of 4:K bisects *PARM1* (IGV: chr4:75,039,107-75,039,147) and is connected to the 5' end of 10:**I** (IGV: chr10:53,584,415-53,584,454). This is within derivative chromosome 4.

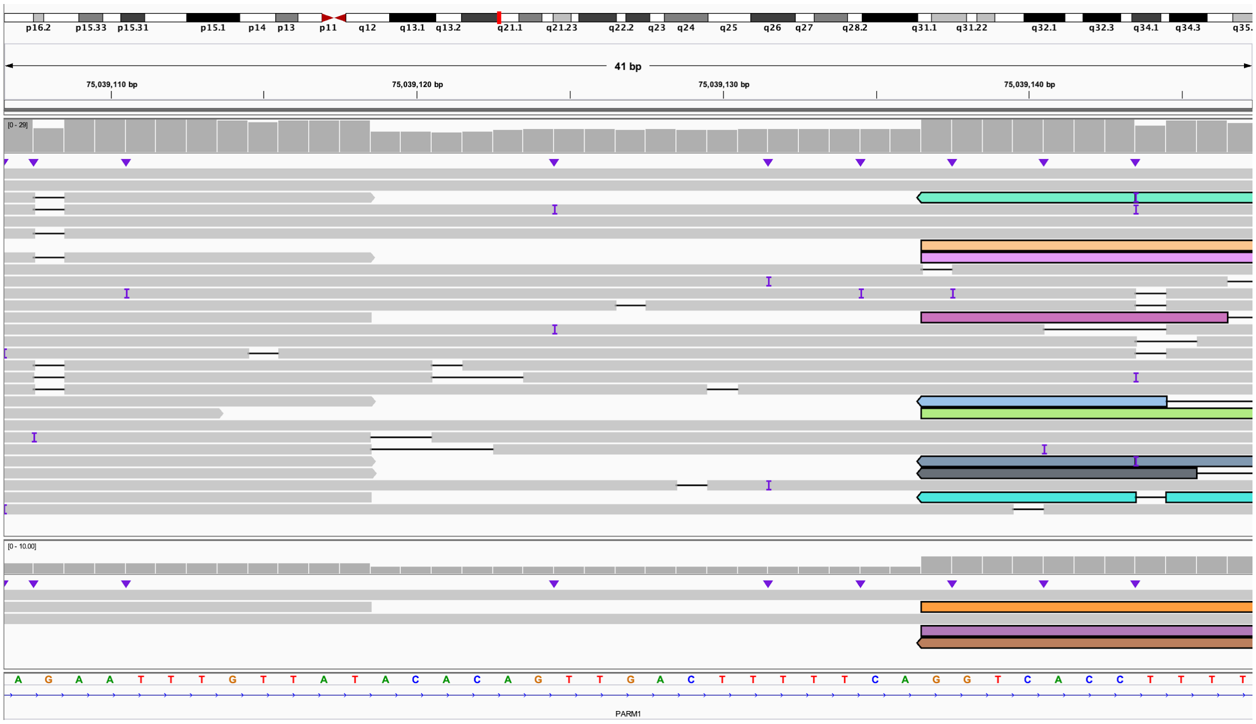

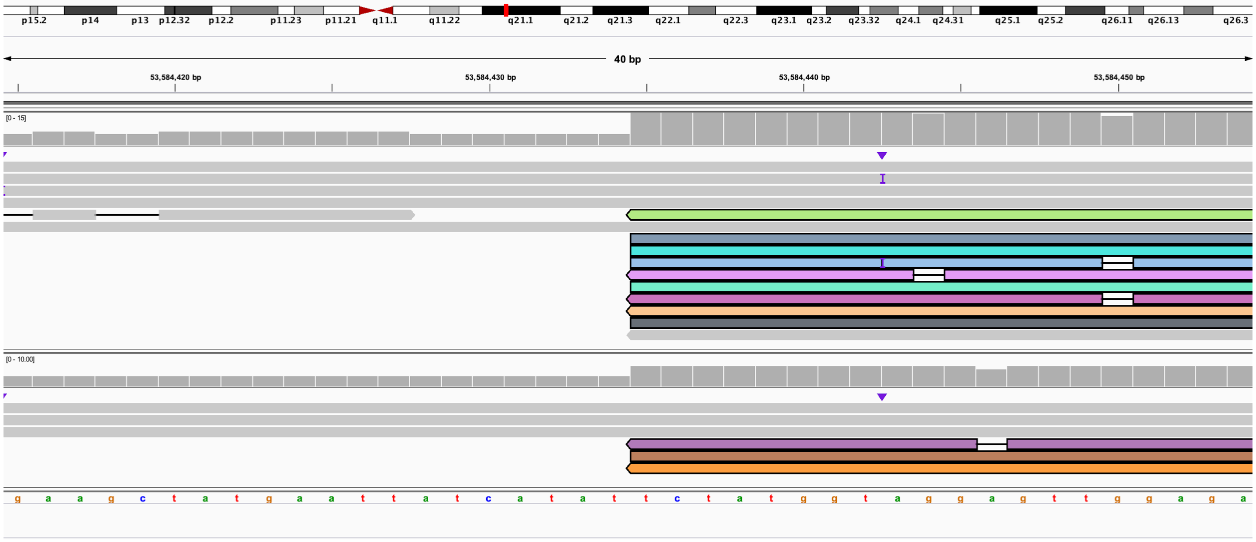

**M.** The 3' end of 10:I (IGV: chr10:56,035,801-56,035,841) is connected to the 3' end of 4:J and bisects *PARM1* (IGV: chr4:75,039,098-75,039,138). This is within derivative chromosome 4.

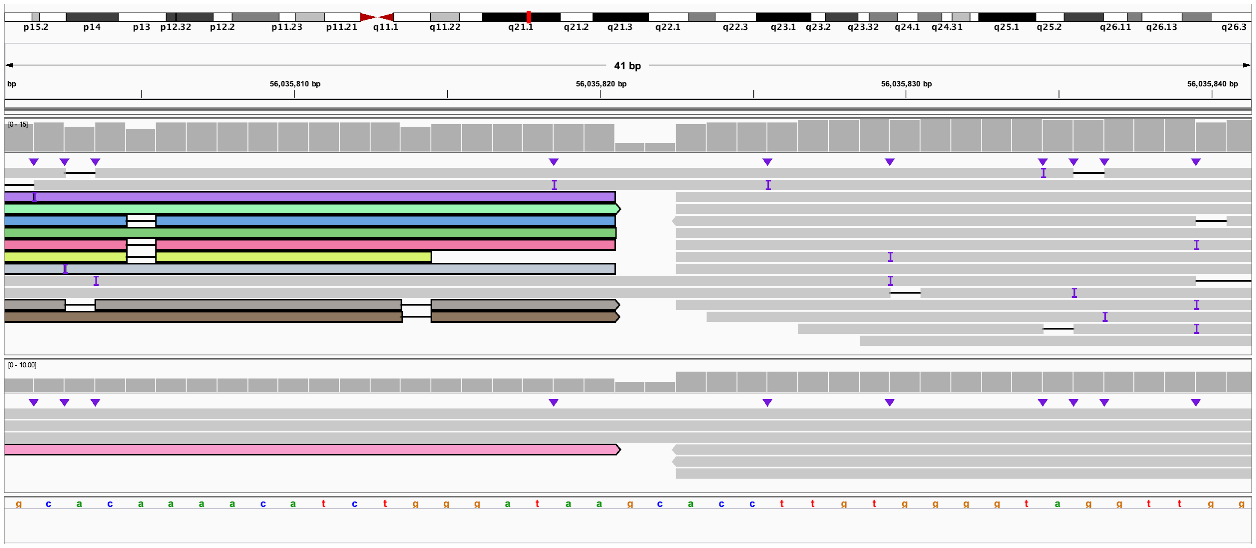

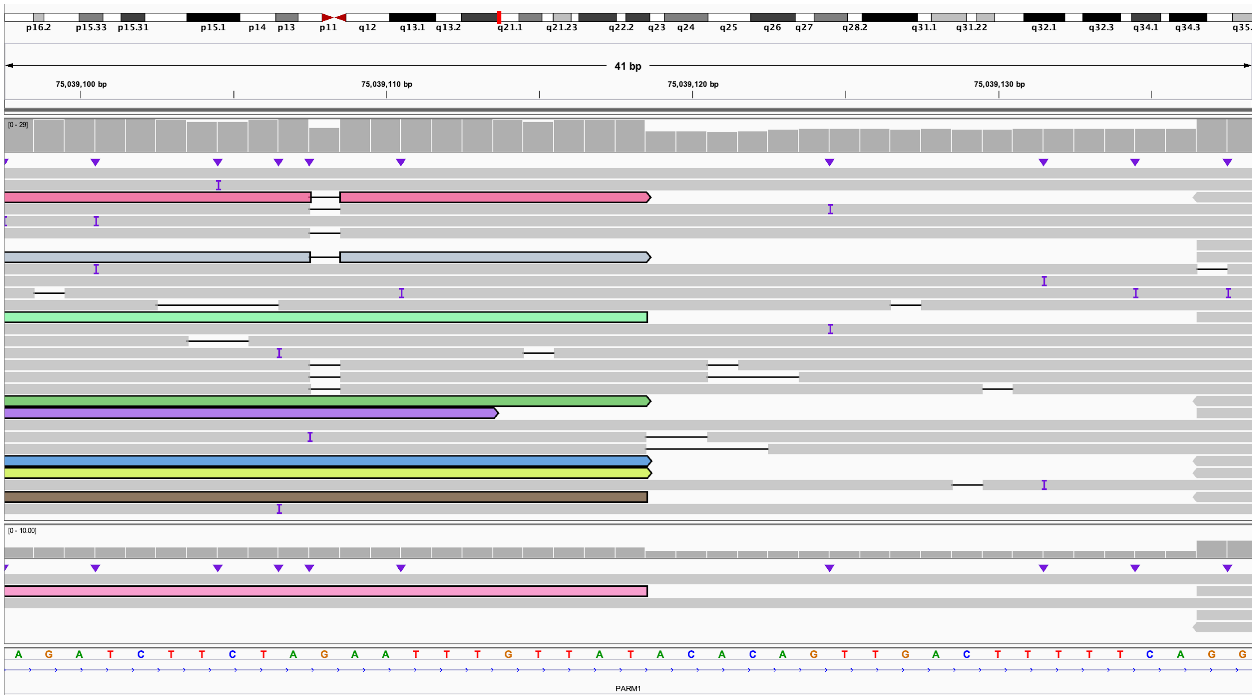

**N.** The 5' end of 4:J bisects *XR_938881.1* (IGV: chr4:74,831,229-74,831,268) and is connected to the 3' end of 4:M (IGV: chr4:81,632,550-81,632,590). This is within derivative chromosome 4.

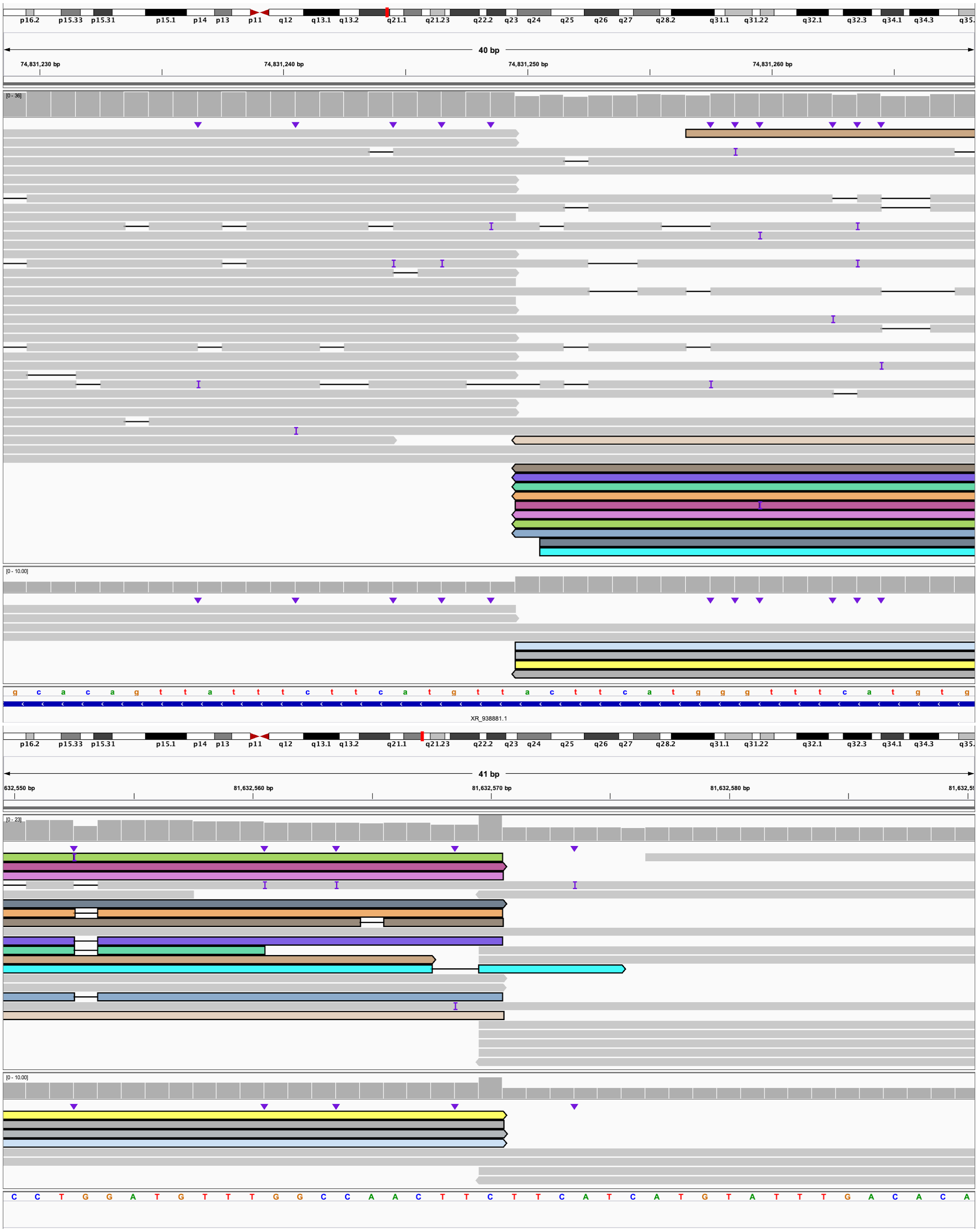

**O.** The 5' end of 4:M (IGV: chr4:79,626,700-79,626,740) is connected to the 3' end of 2:B (IGV: chr2:16,235,691-16,235,730). This is within derivative chromosome 4.

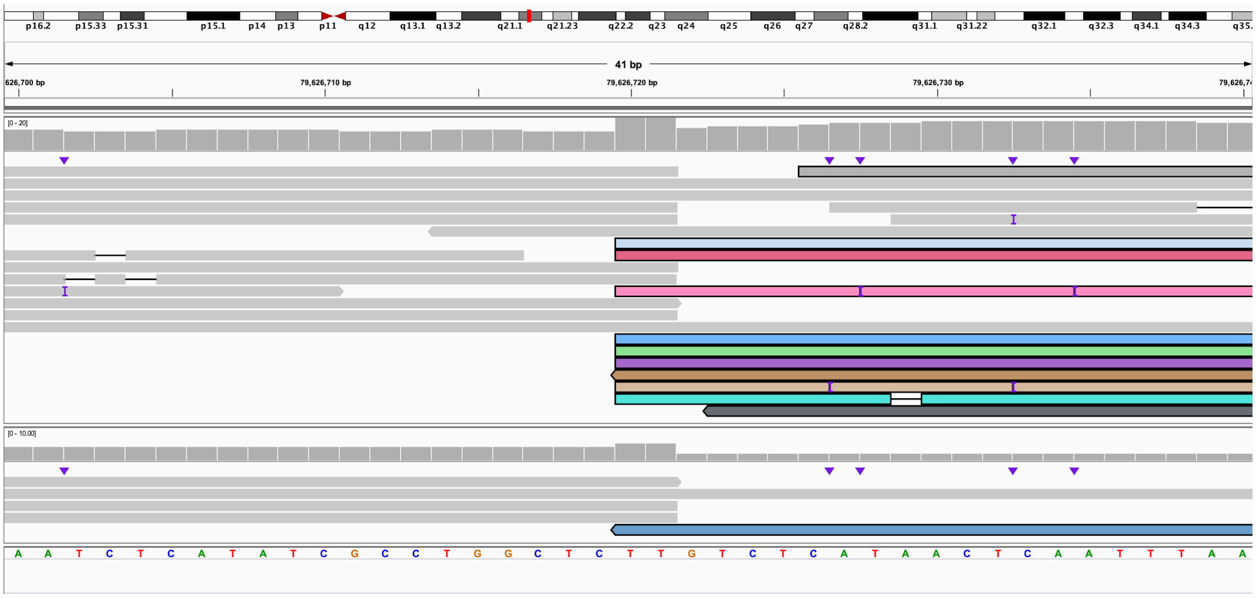

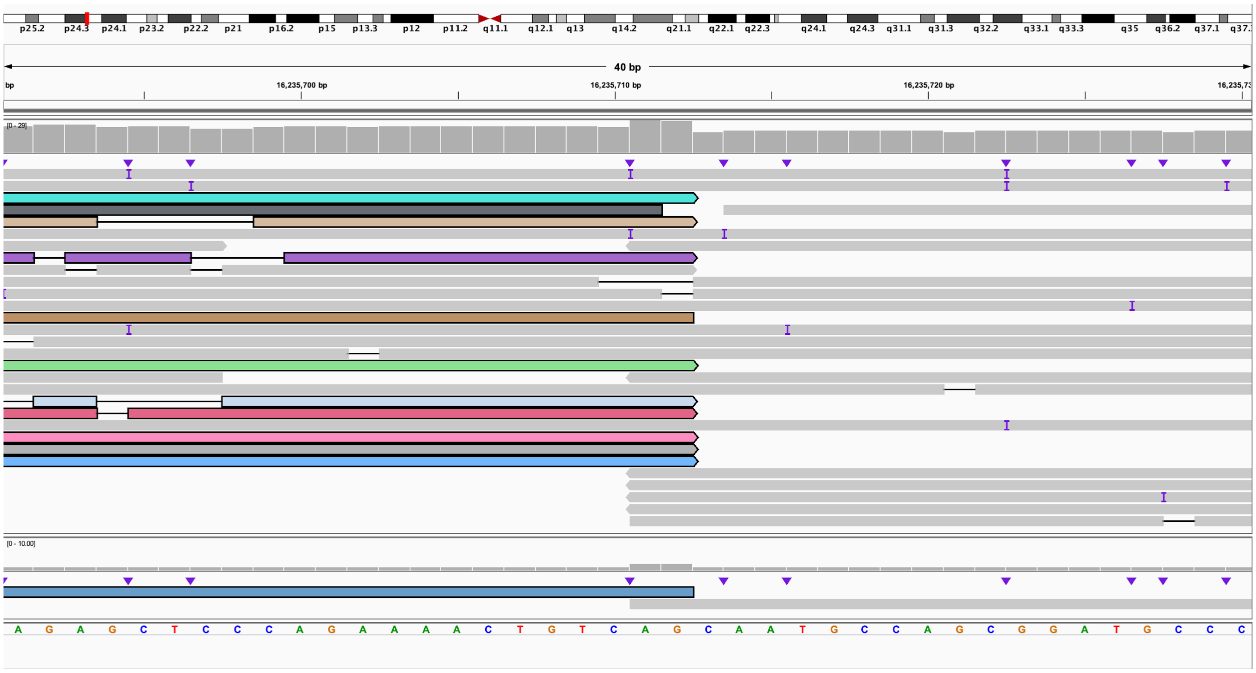

**P.** The 5' end of 2:B (IGV: chr2:7,777,974-7,778,013) is connected to the 5' end of 10:J (IGV: chr10:56,035,802-56,035,841). This is within derivative chromosome 4.

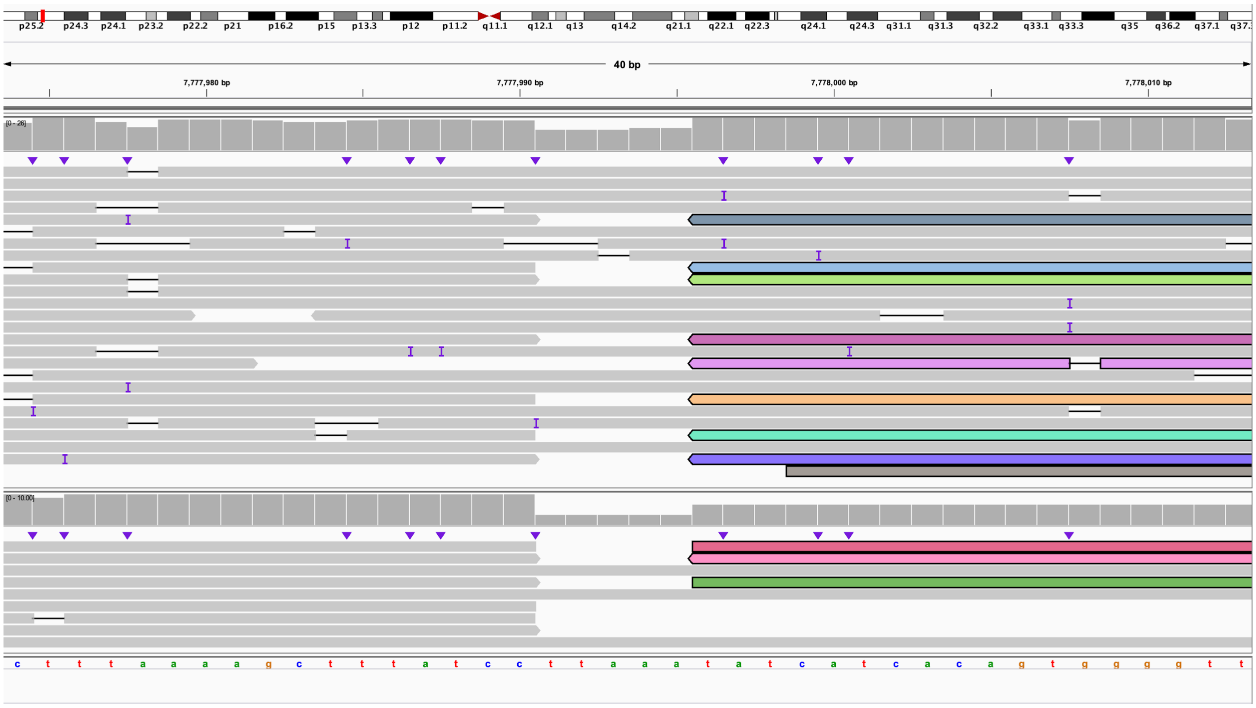

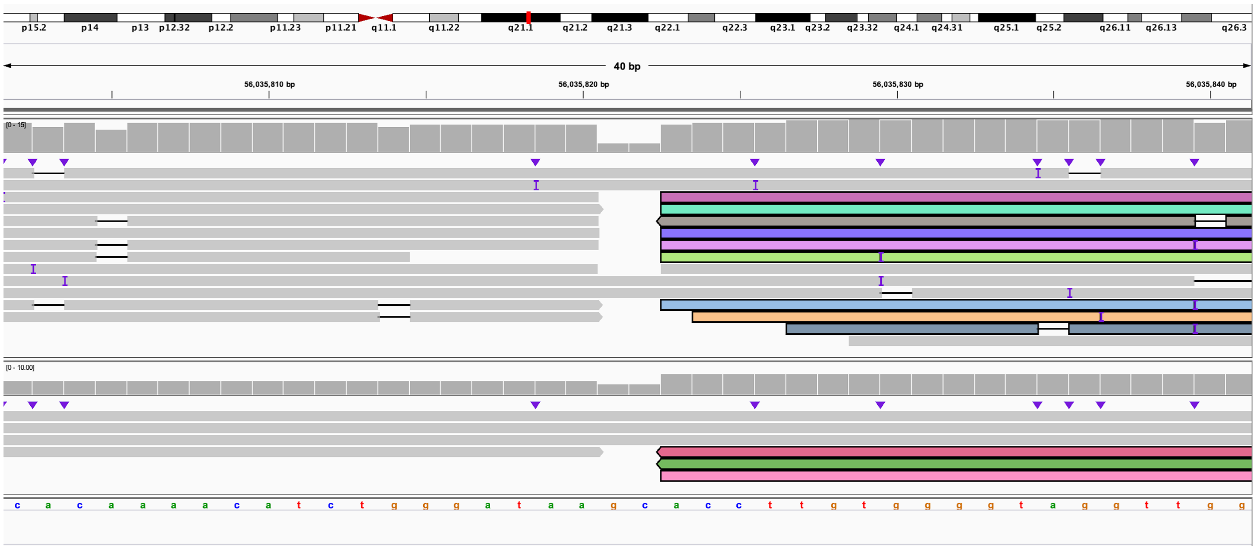

**Q.** The 3' end of 10:J (IGV: chr10:65,519,808-65,519,848) is connected to the 3' end of 10:M and bisects *LRMDA* (IGV: chr10:76,402,362-76,402,402). This is within derivative chromosome 4.

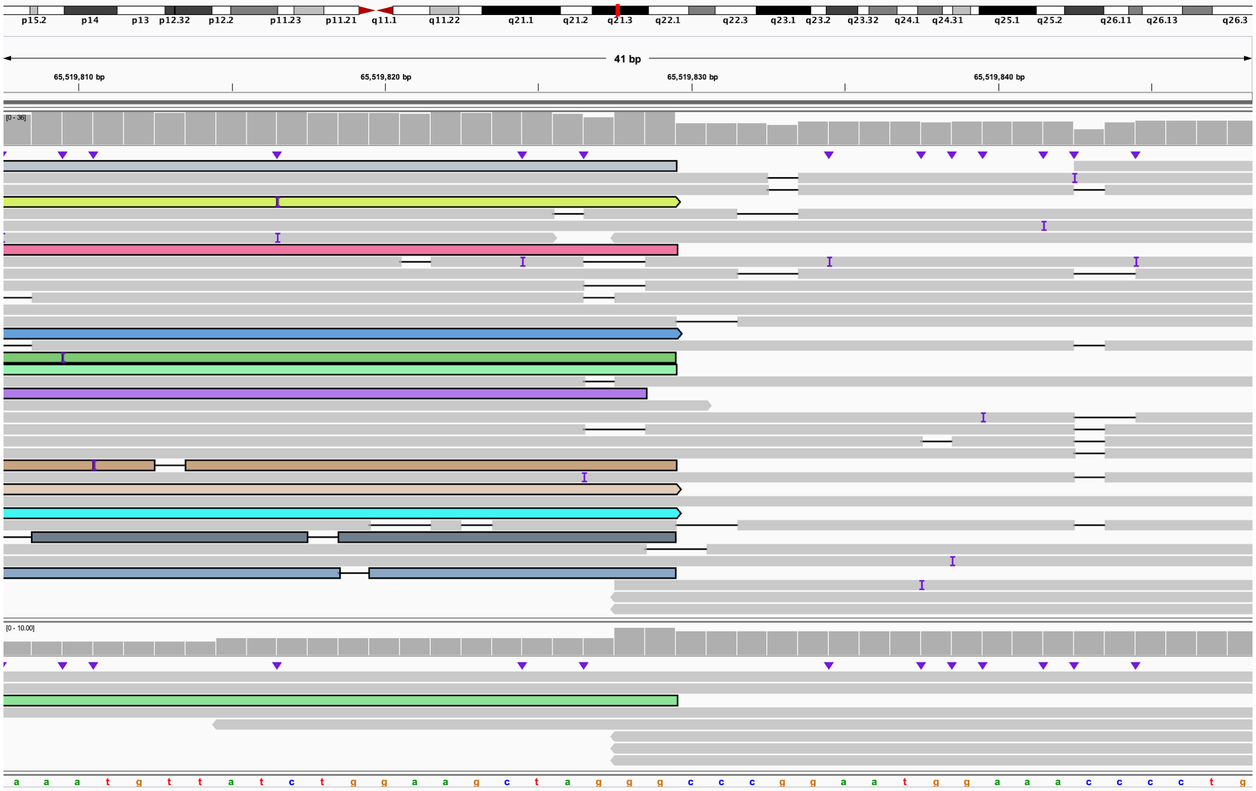

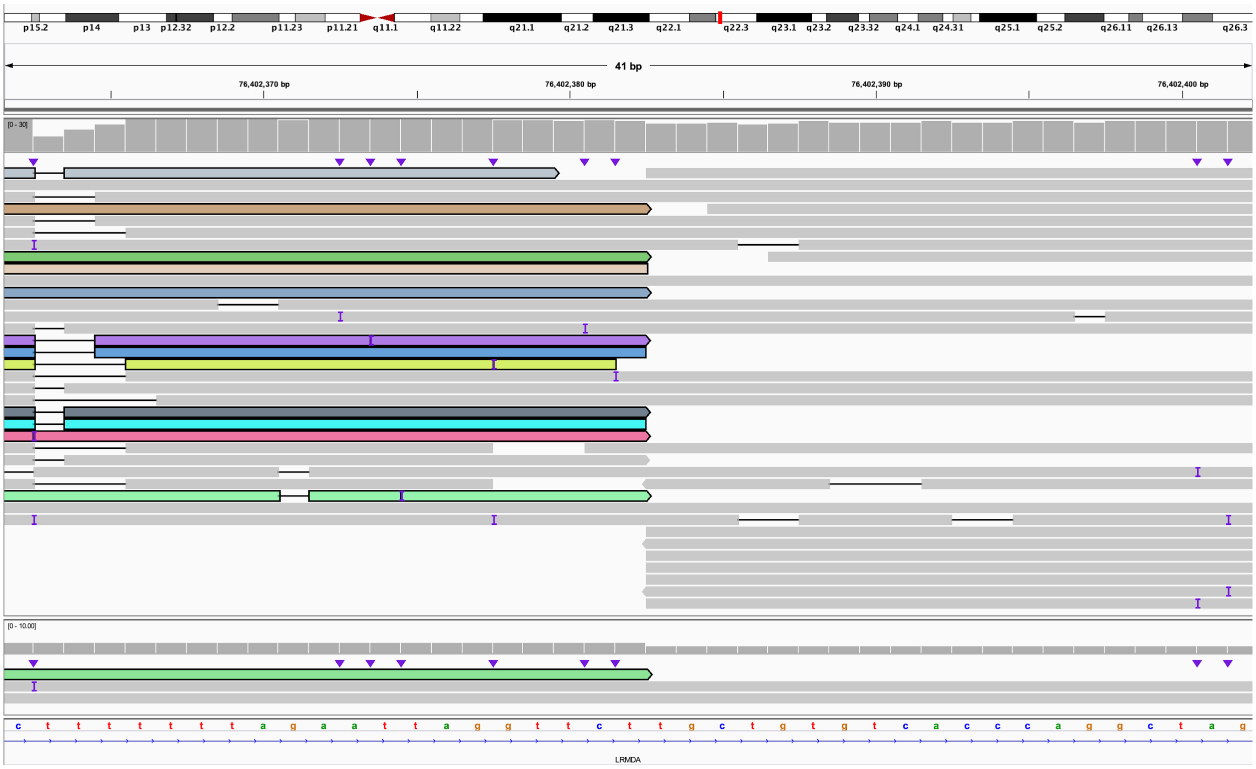

**R.** The 5' end of 10:M (IGV: chr10:74,772,250-74,772,290) is connected to the 5' end of 10:N and bisects *LRMDA* (IGV: chr10:76,402,363-76,402,402). This is the end of derivative chromosome 4.

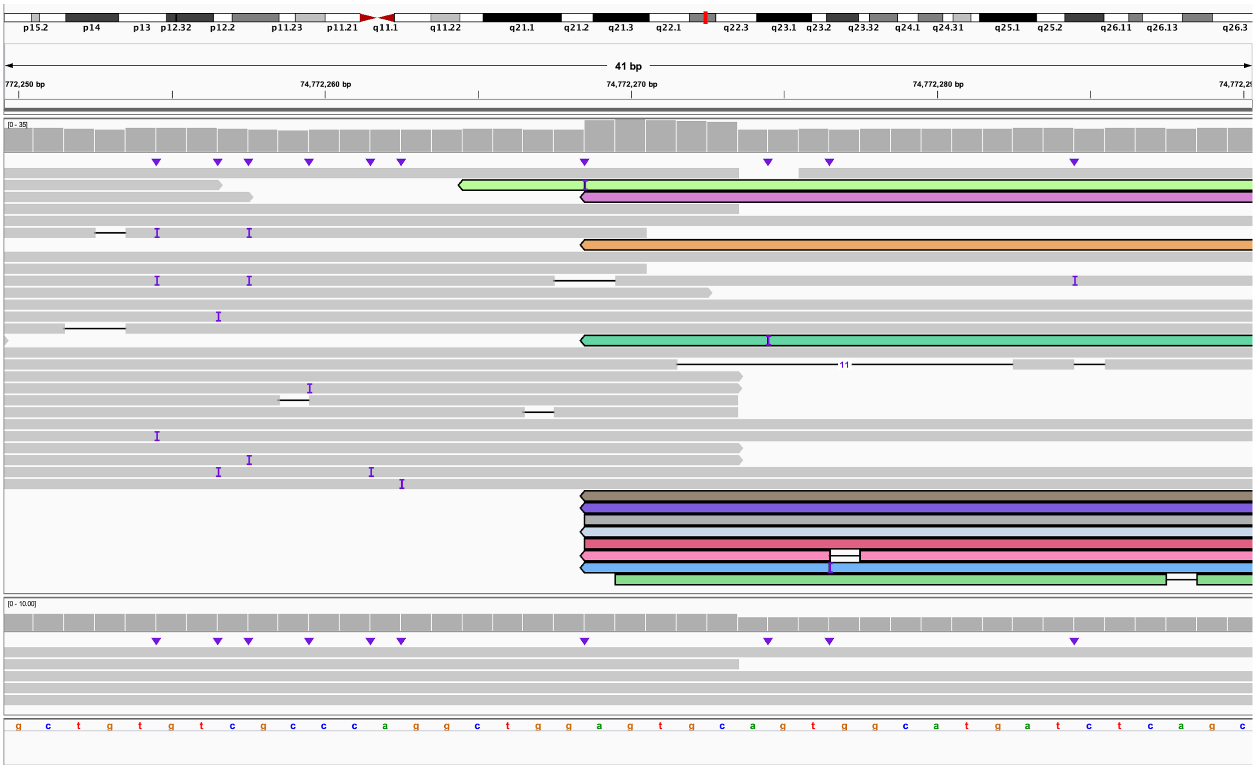

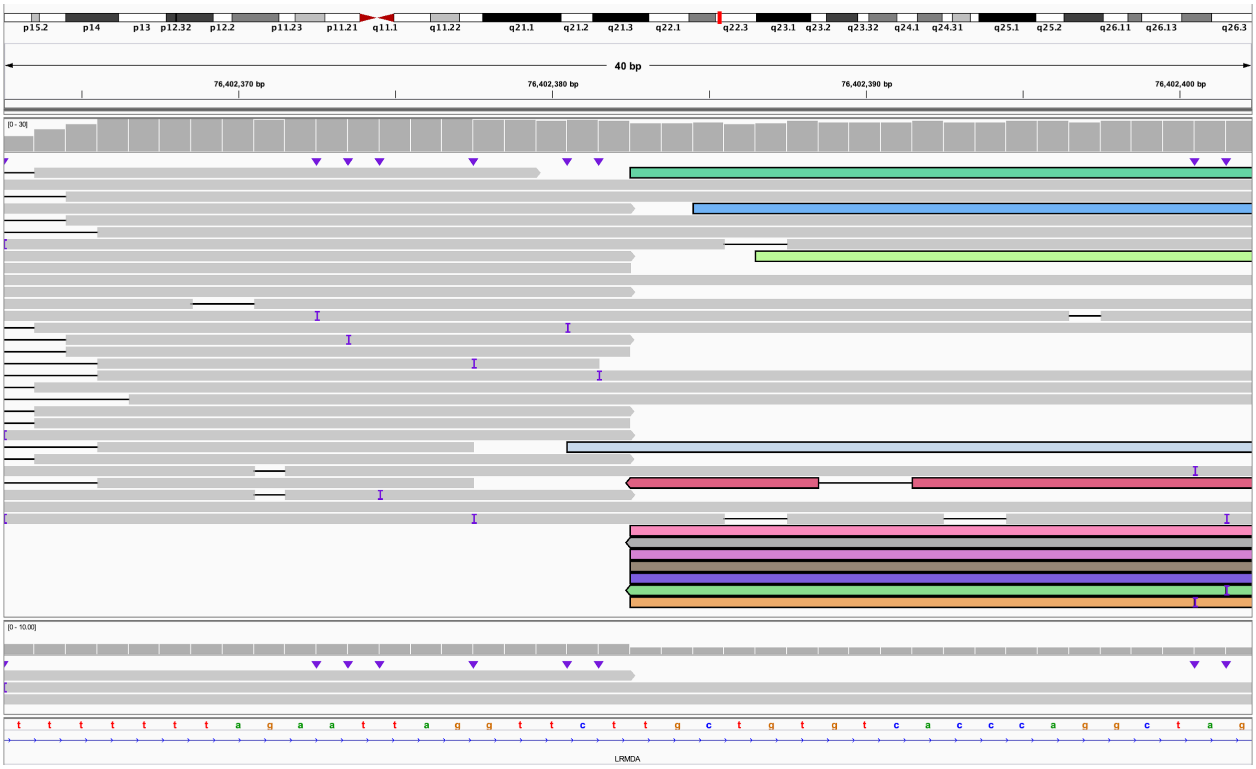

**S.** The 3' end of 10:A bisects *SGMS1* (IGV: chr10:50,382,074-50,382,113) and is connected to the 5' end of 10:D (IGV: chr10:50,694,757-50,694,797). This is the beginning of derivative chromosome 10.

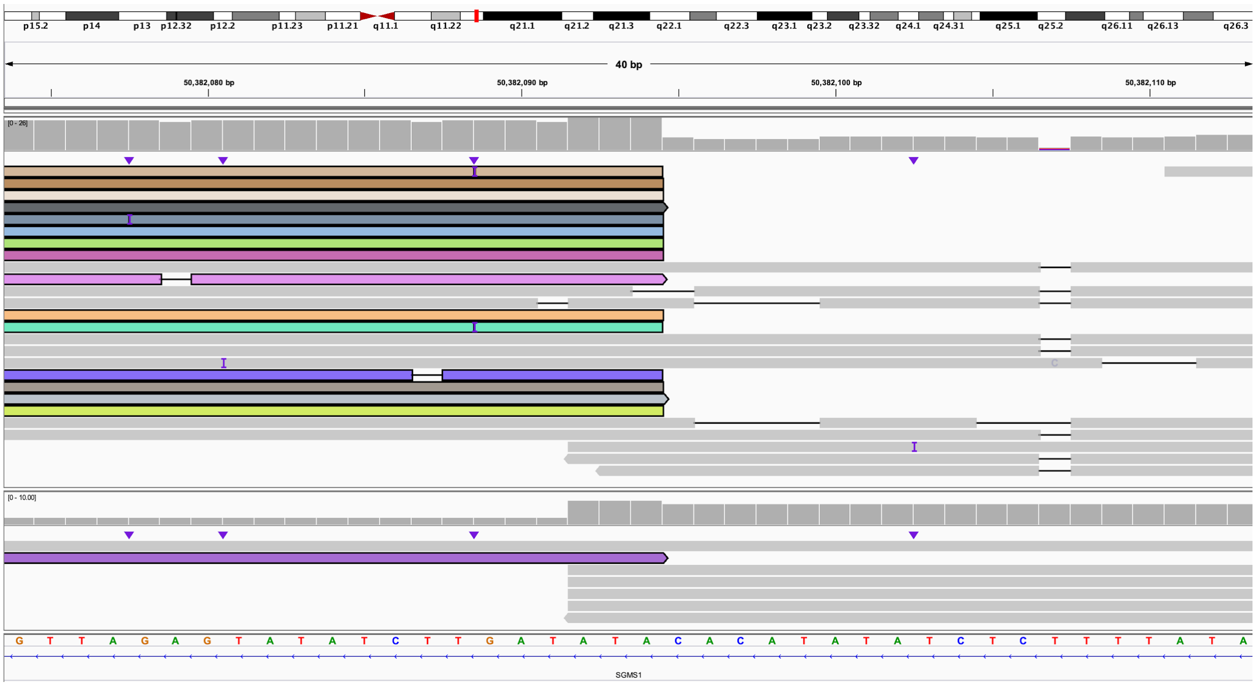

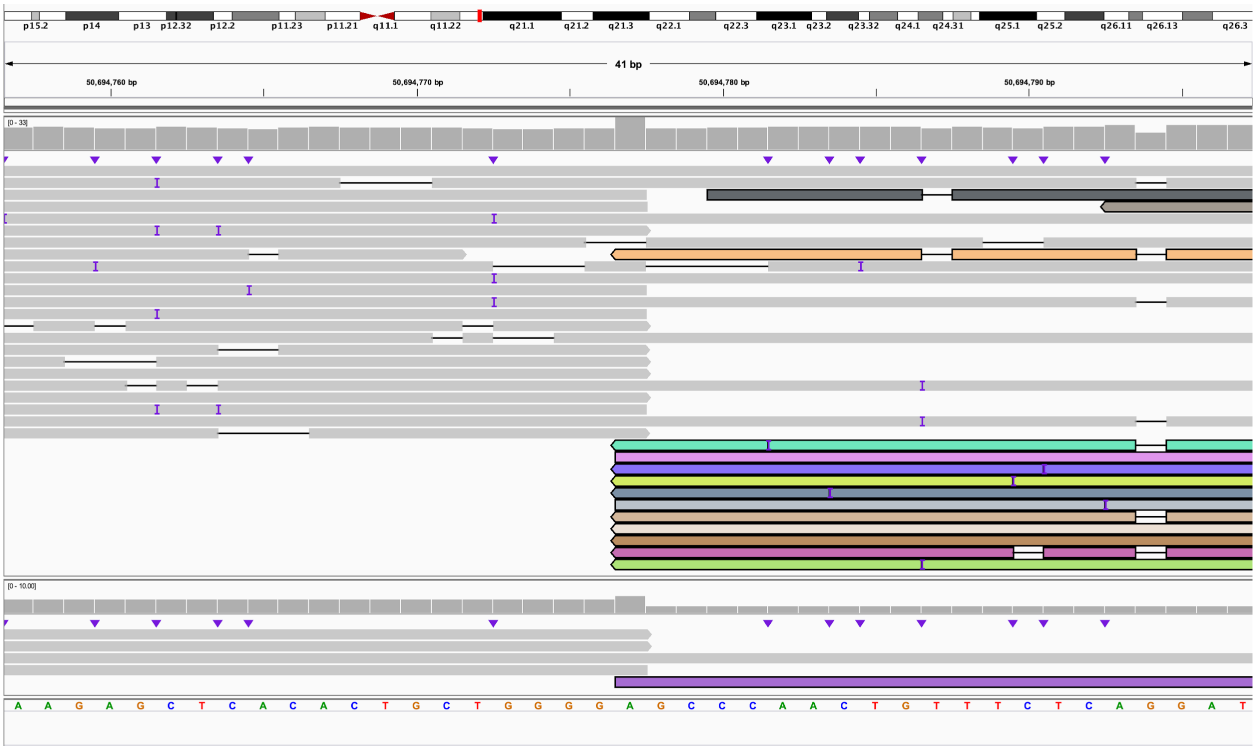

**T.** The 3' end of 10:D bisects *PRKG1* (IGV: chr10:51,260,001-51,260,040) and is connected to the 3' end of 10:B which bisects *SGMS1* (IGV: chr10:50,563,484-50,563,523). This is within derivative chromosome 10.

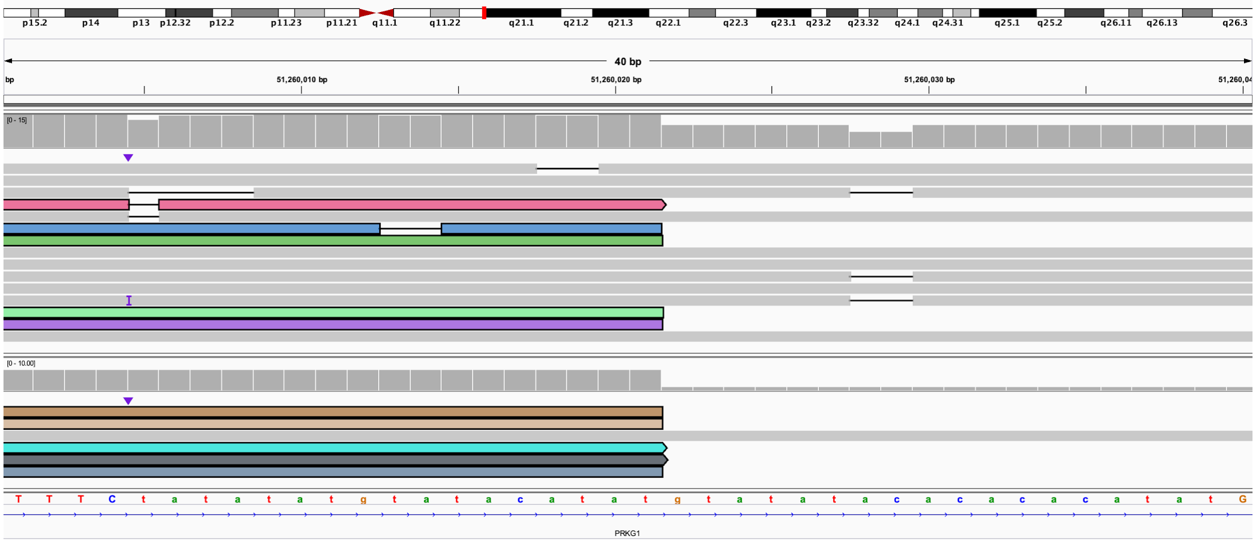

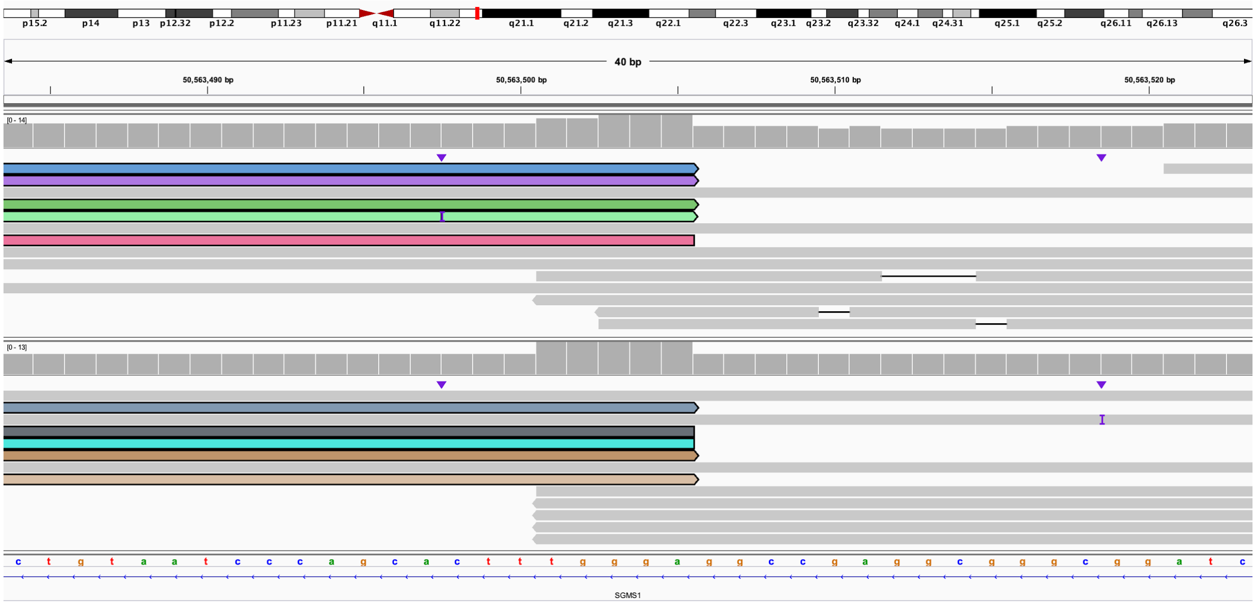

**U.** The 5' end of 10:B bisects *SGMS1* (IGV: chr10:50,382,074-50,382,114) and is connected to the 5' end of 10:C and bisects *SGMS1* (IGV: chr10:50,563,485-50,563,525). This is within derivative chromosome 10.

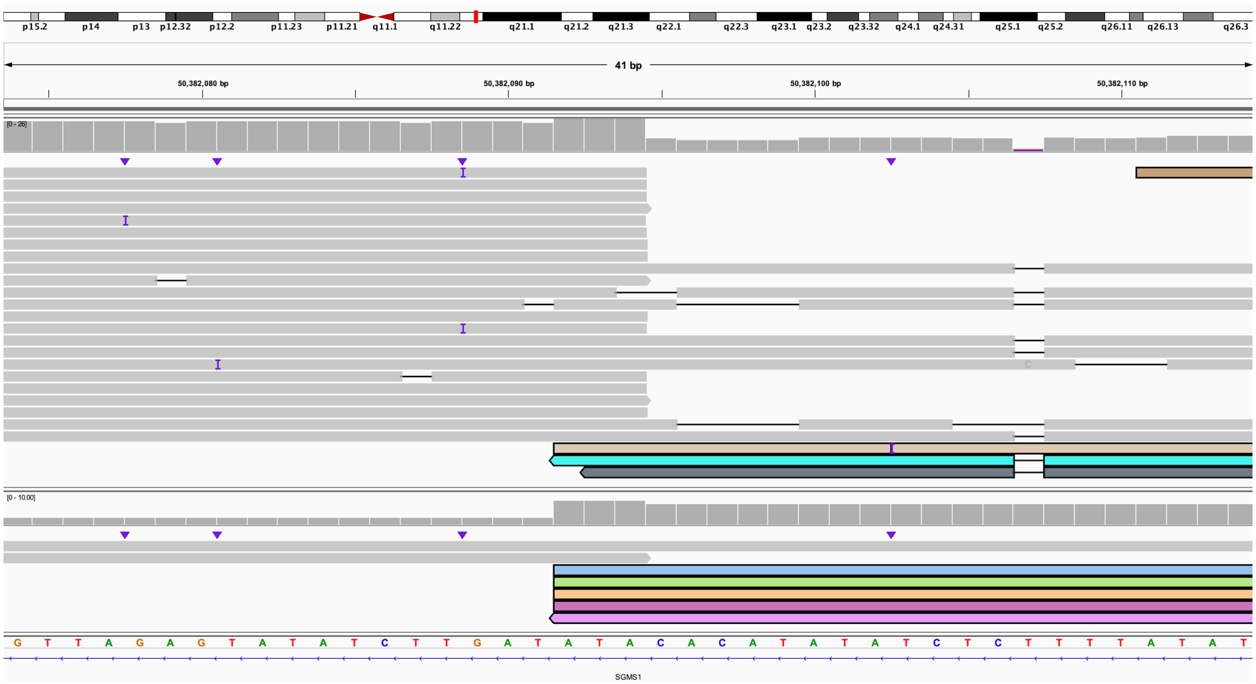

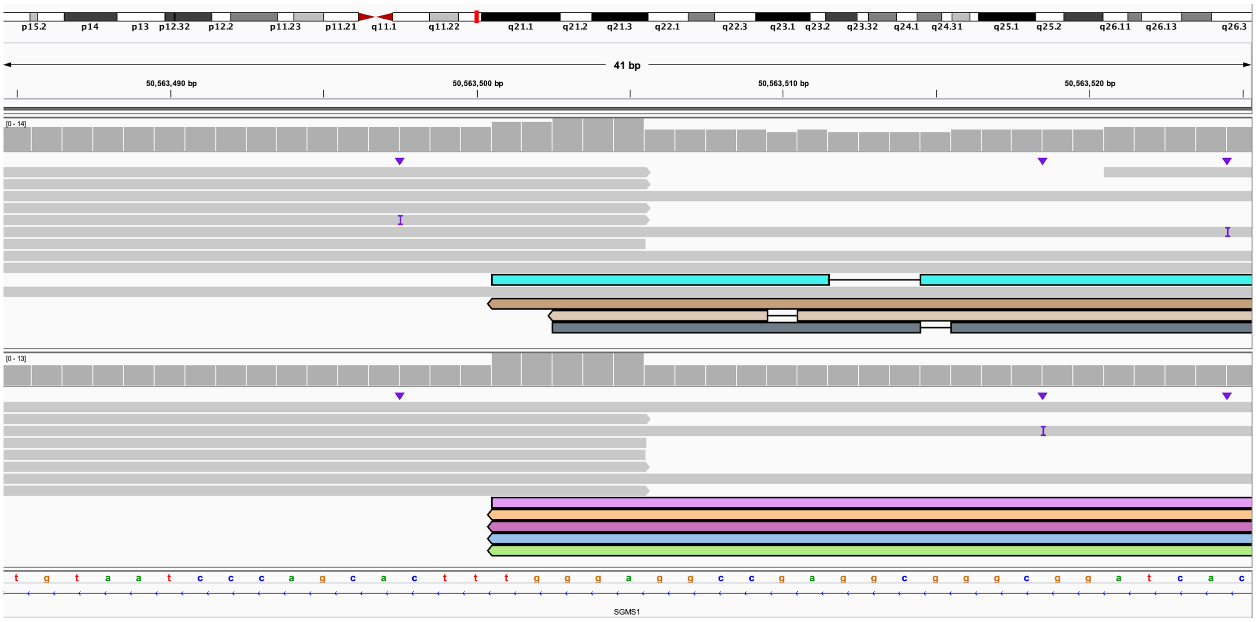

**V.** The 3' end of 10:C (IGV: chr10:50,694,757-50,694,797) is connected to the 3' end of 10:F (IGV: chr10:53,548,467-53,548,507). This is within derivative chromosome 10.

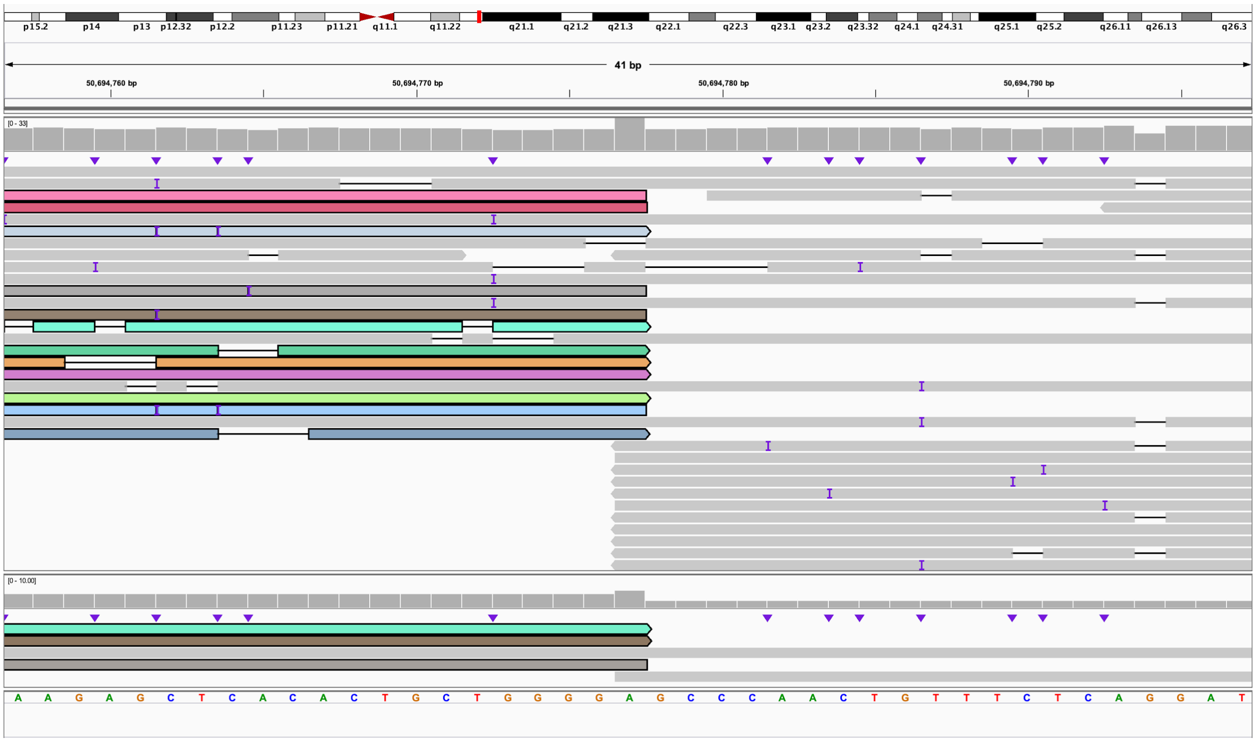

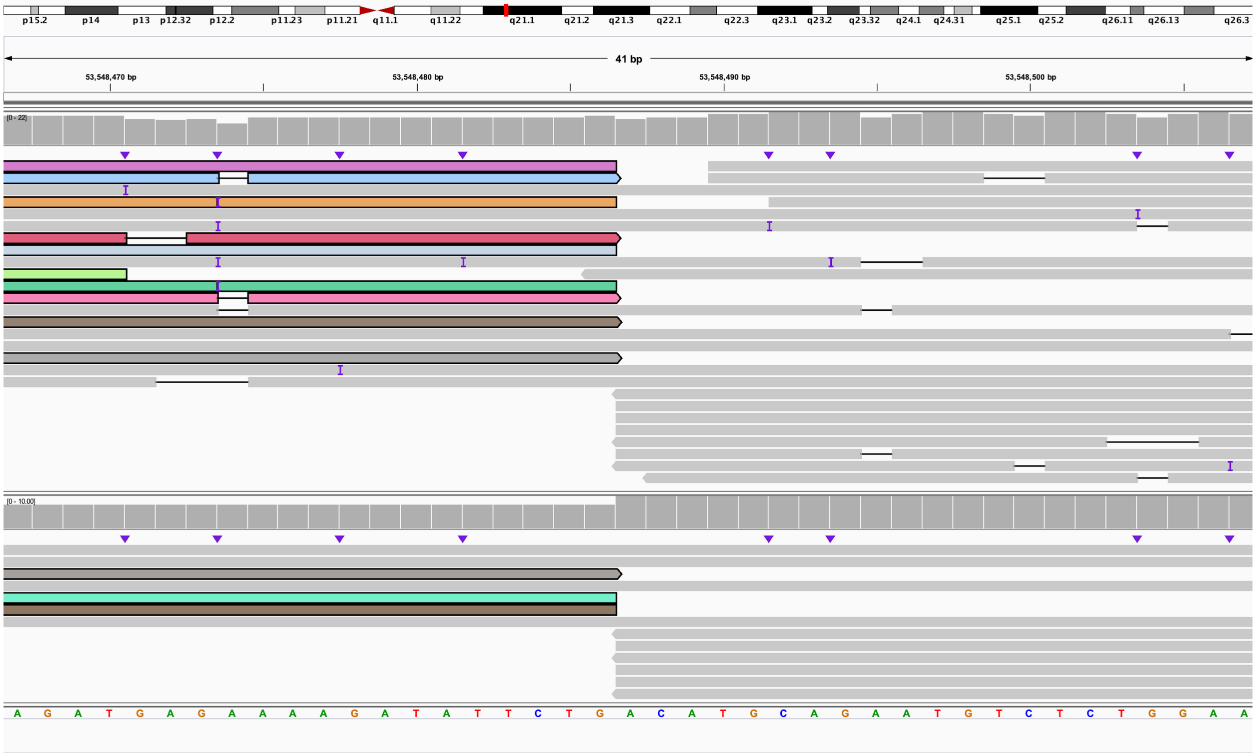

**W.** The 5' end of 10:F bisects *PRKG1* (IGV: chr10:51,840,859-51,840,899) and is connected to the 5' end of 4:F and bisects *AFP* (IGV: chr4:73,446,108-73,446,148). This is within derivative chromosome 10.

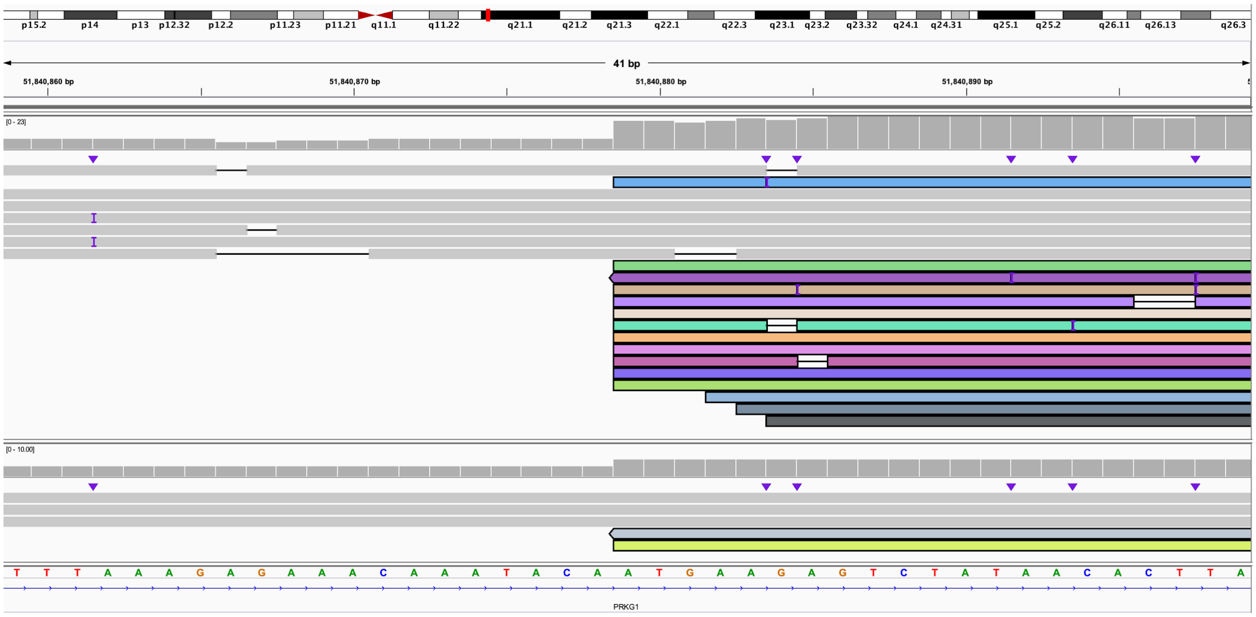

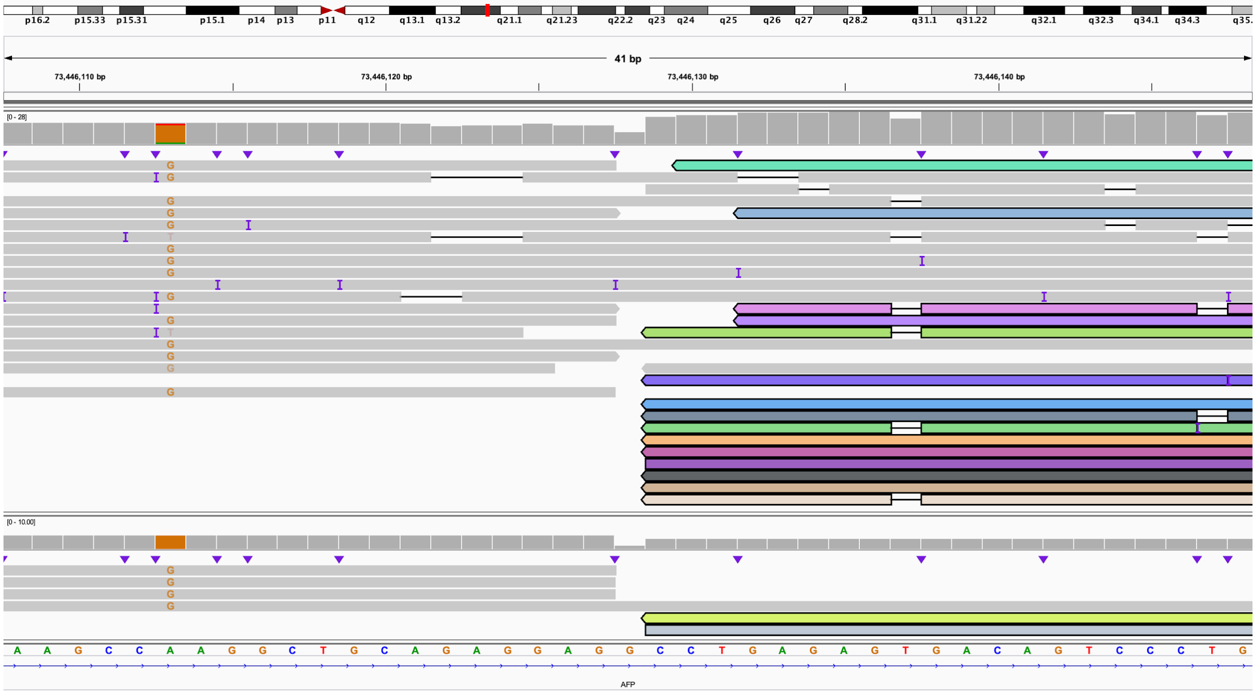

**X.** The 3' end of 4:F (IGV: chr4:73,771,810-73,771,850) is connected to the 5' end of 4:E which bisects *ANKRD17* (IGV: chr4:73,129,539-73,129,579). This is within derivative chromosome 10.

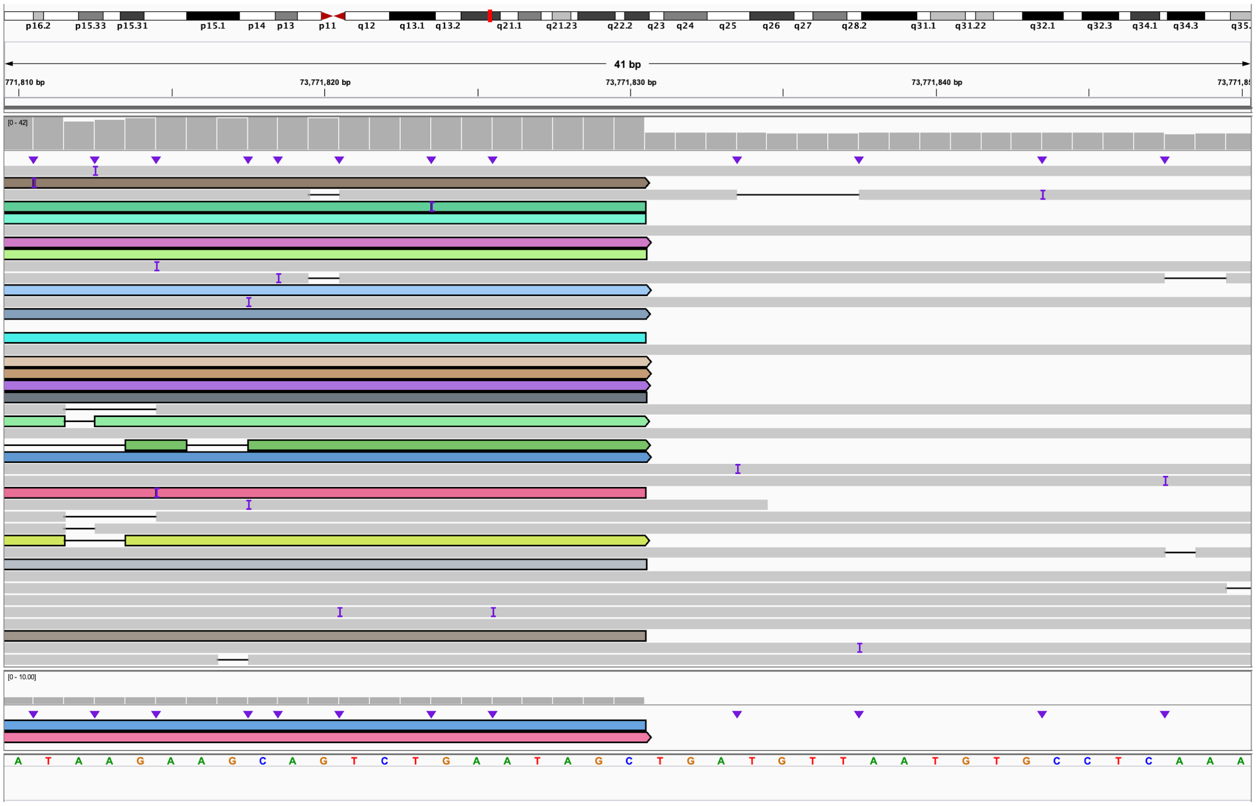

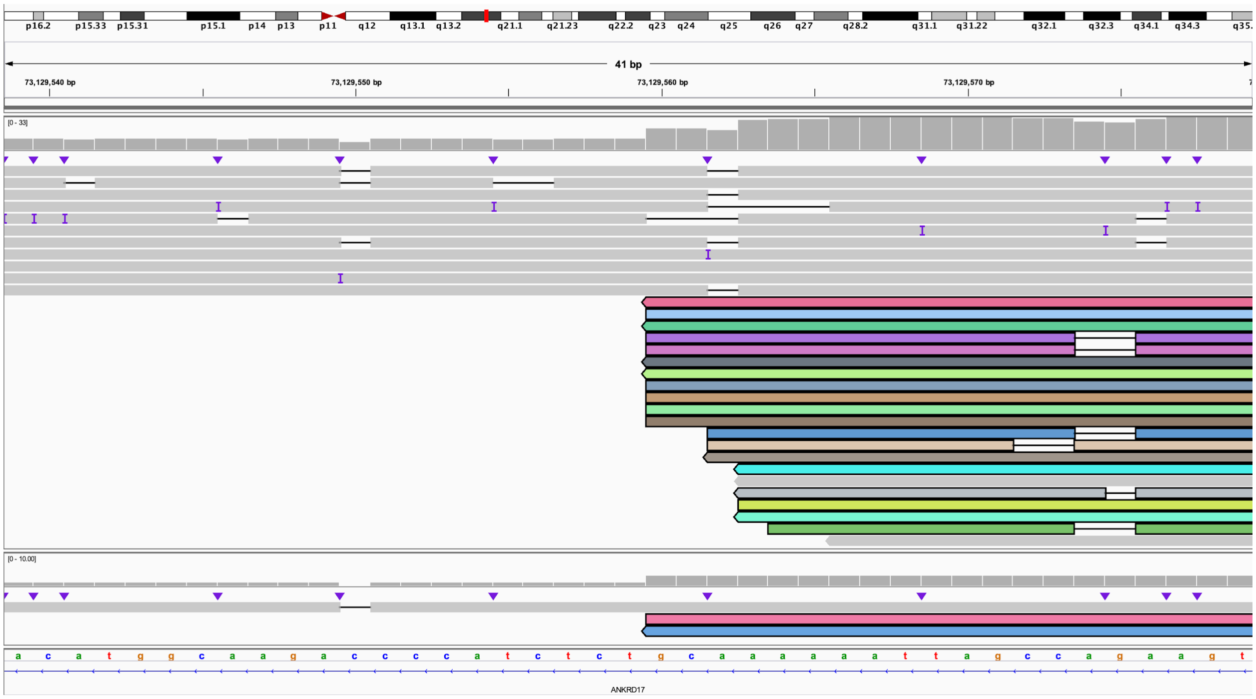

**Y.** The 3' end of 4:E bisects *AFP* (IGV: chr4:73,446,108-73,446,148) and is connected to the 5' end of 10:K (IGV: chr10:65,519,765-65,519,894). This is within derivative chromosome 10.

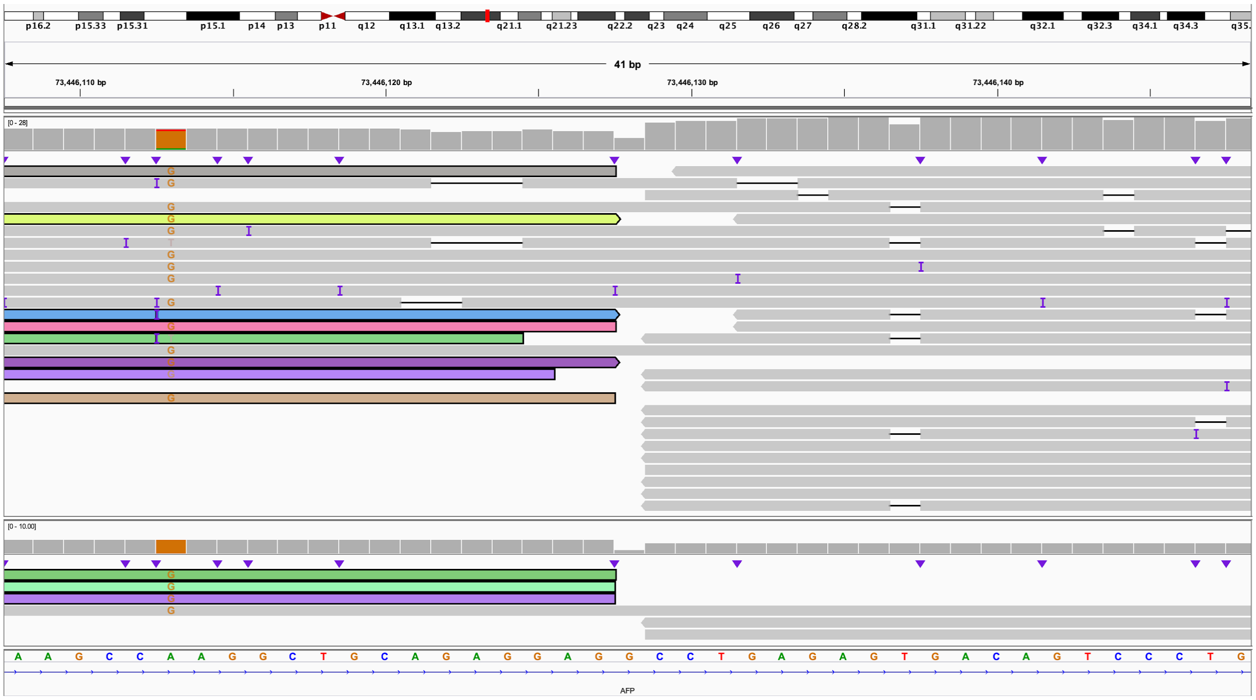

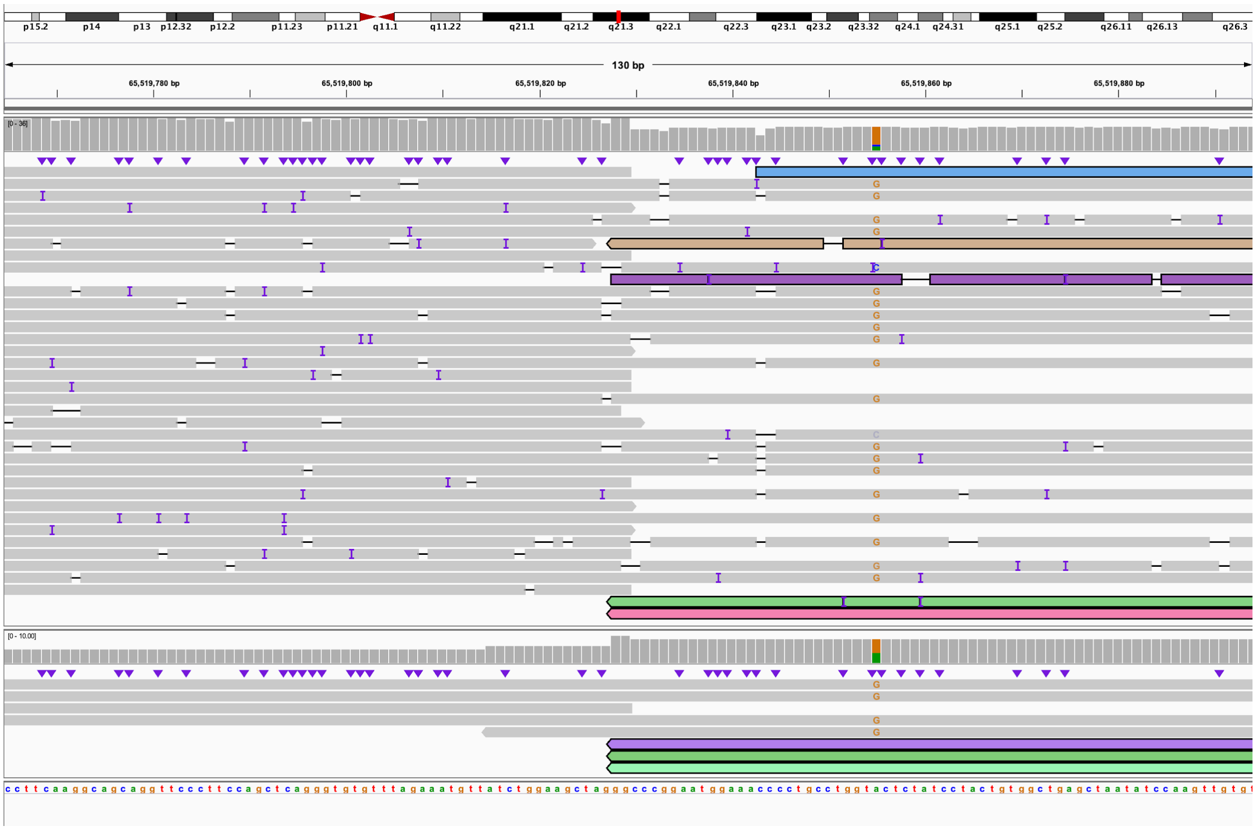

**Z.** The 3' end of 10:K bisects *CTNNA3* (IGV: chr10:67,462,177-67,462,306) and is connected to the 3' end of 10:L (IGV: chr10:74,772,252-74,772,291). This is within derivative chromosome 10.

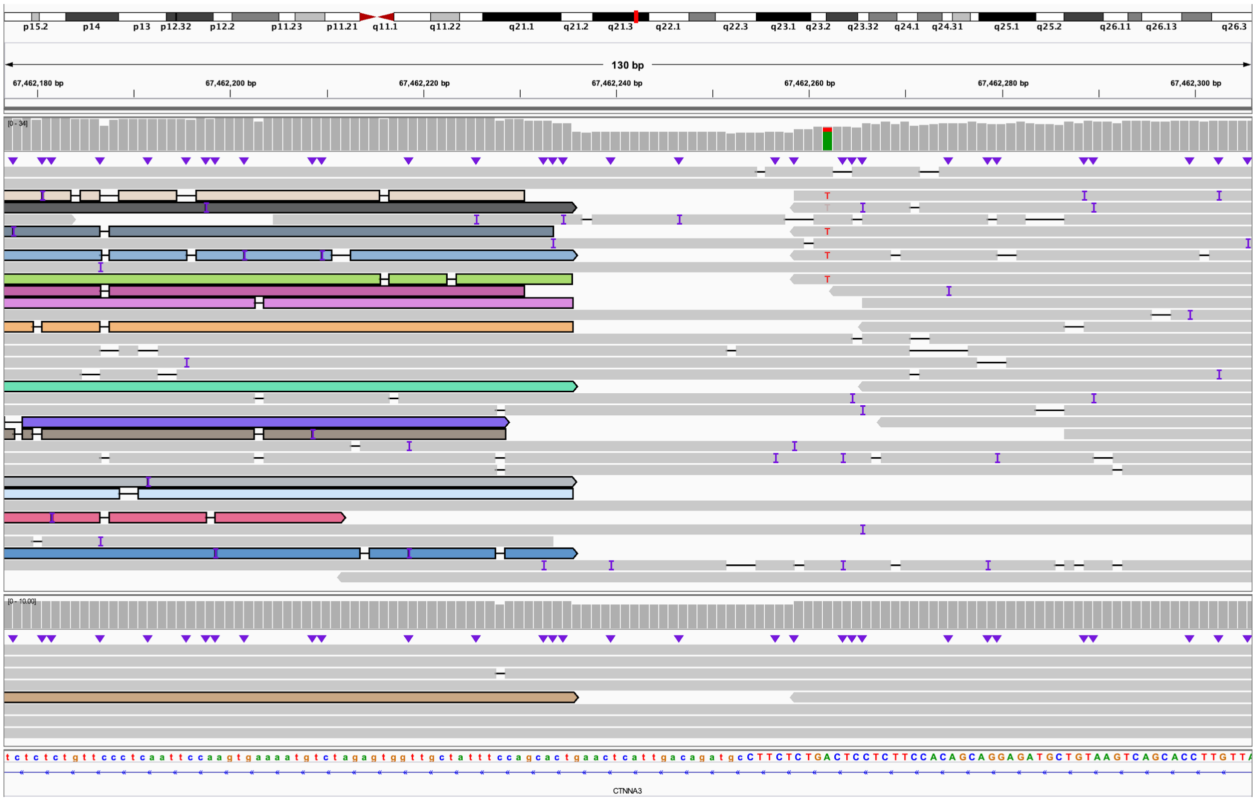

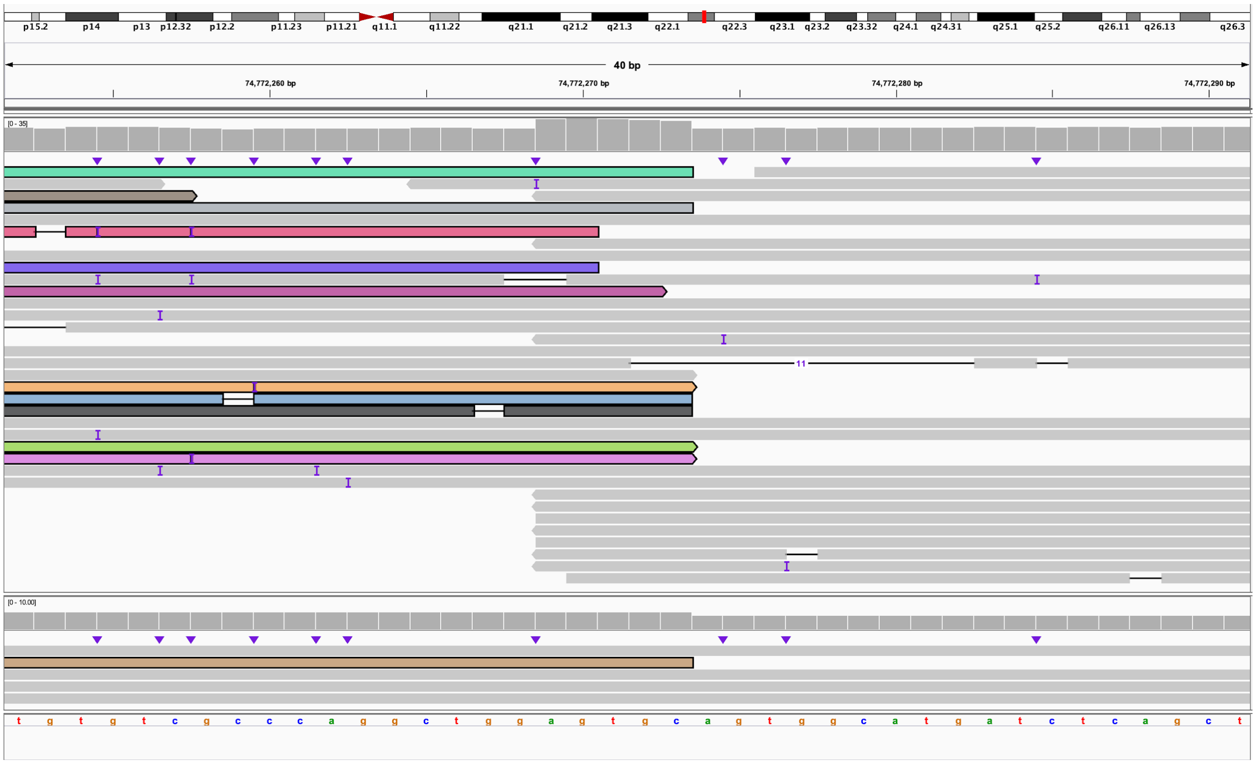

**AA.** The 5' end of 10:L bisects *CTNNA3* (IGV: chr10:67,462,190-67,462,292) and is connected to the 5' end of 14:C (IGV: chr14:22,881,911-22,881,951). This is the end of derivative chromosome 10.

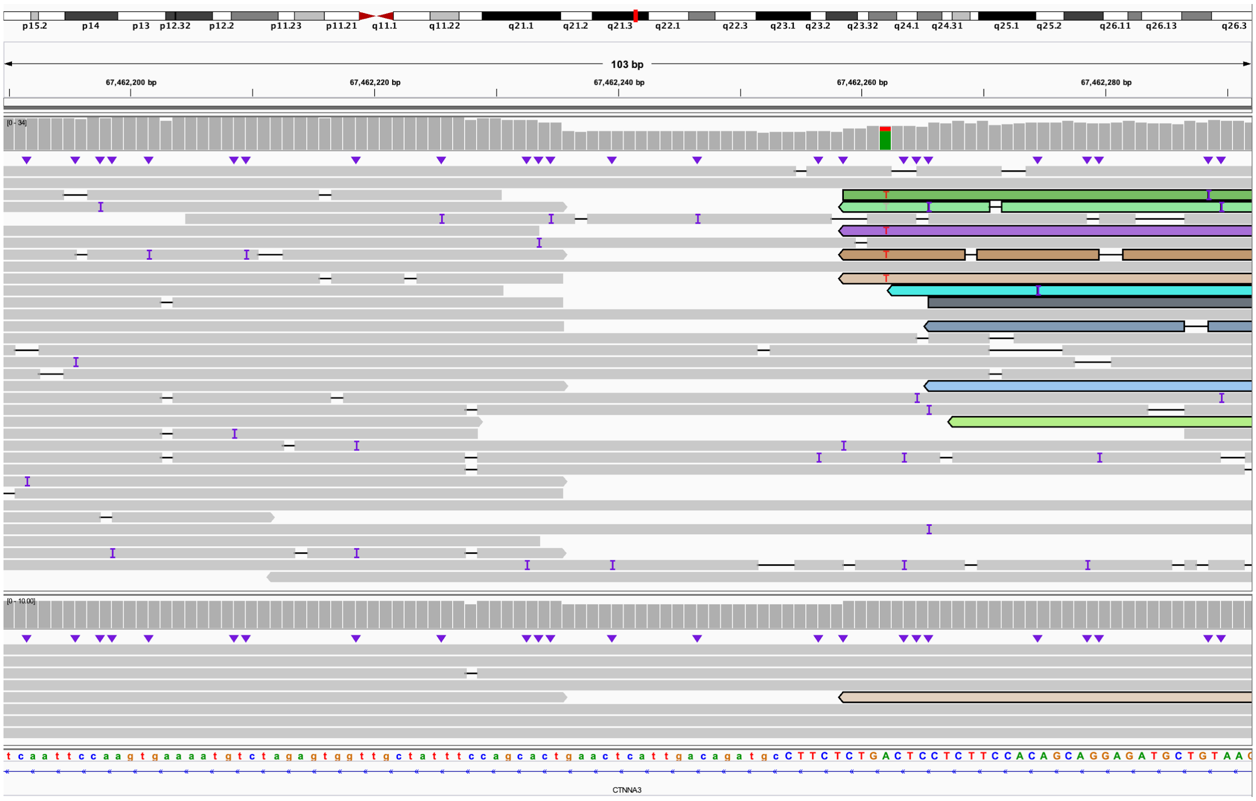

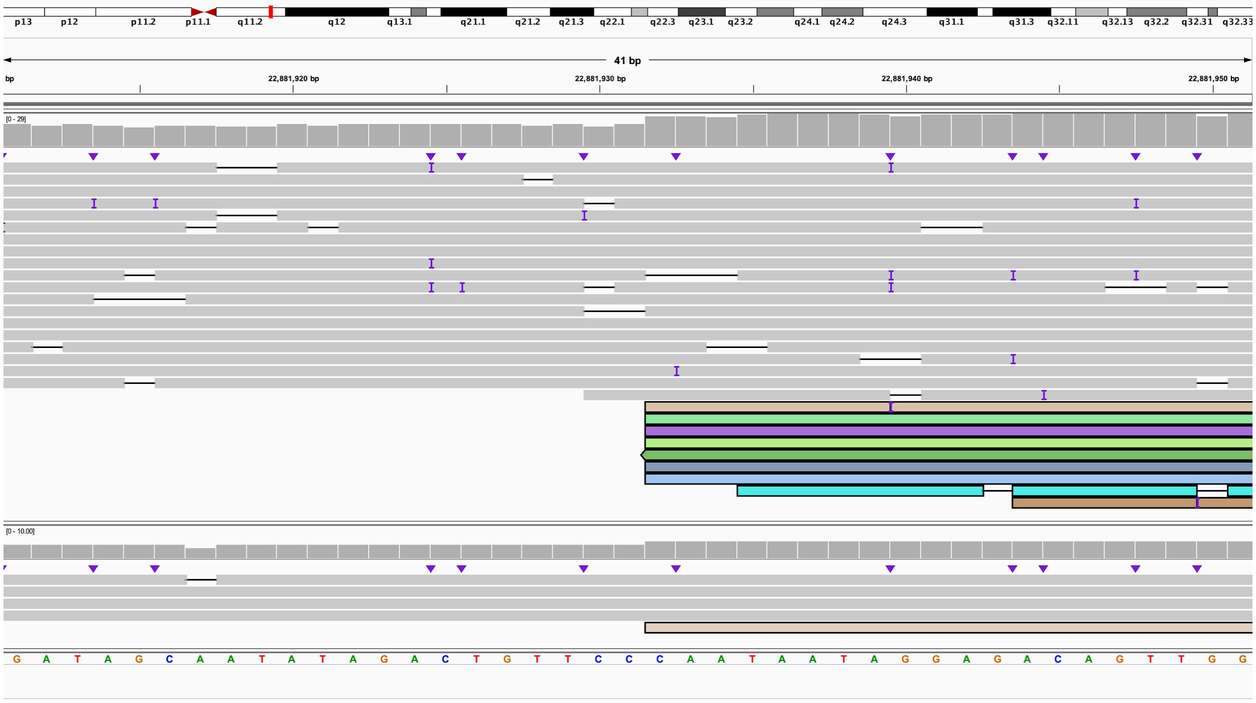

**AB.** The 3' end of 2:A (IGV chr2:7,777,973-7,778,013) is connected to the 5' end of 10:G (IGV chr10:53,548,468-53,548,507). This is the beginning of derivative chromosome 14.

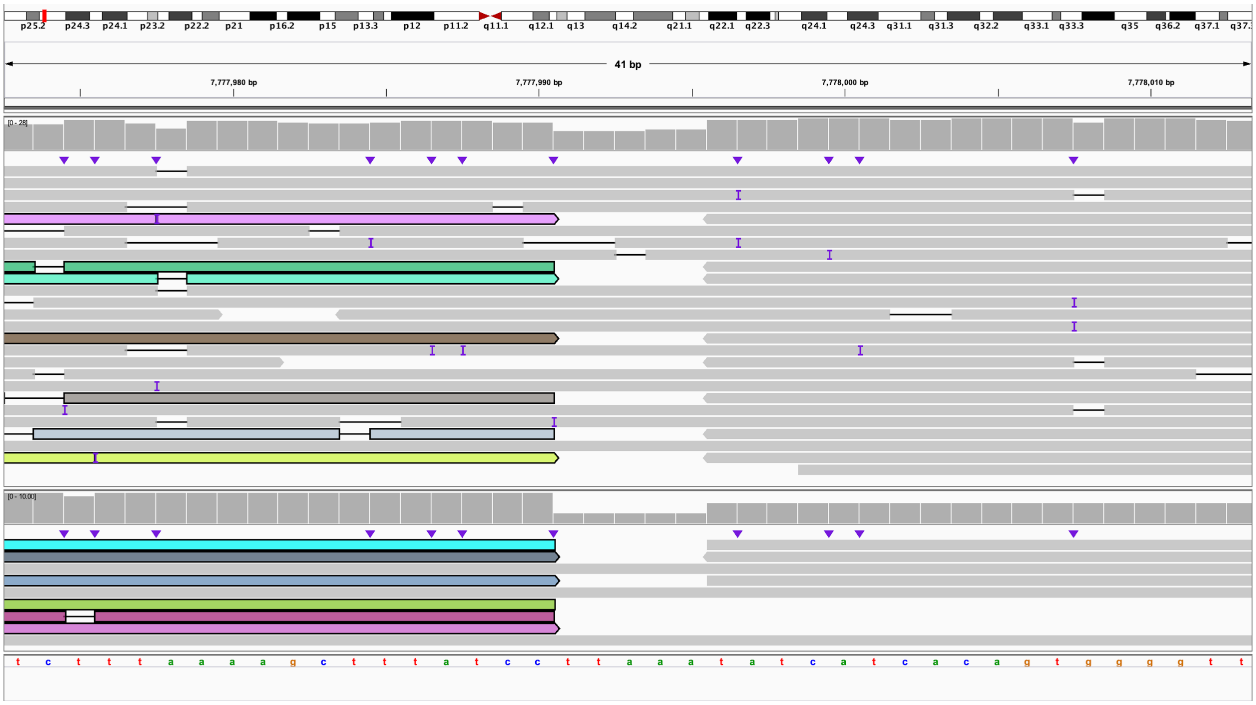

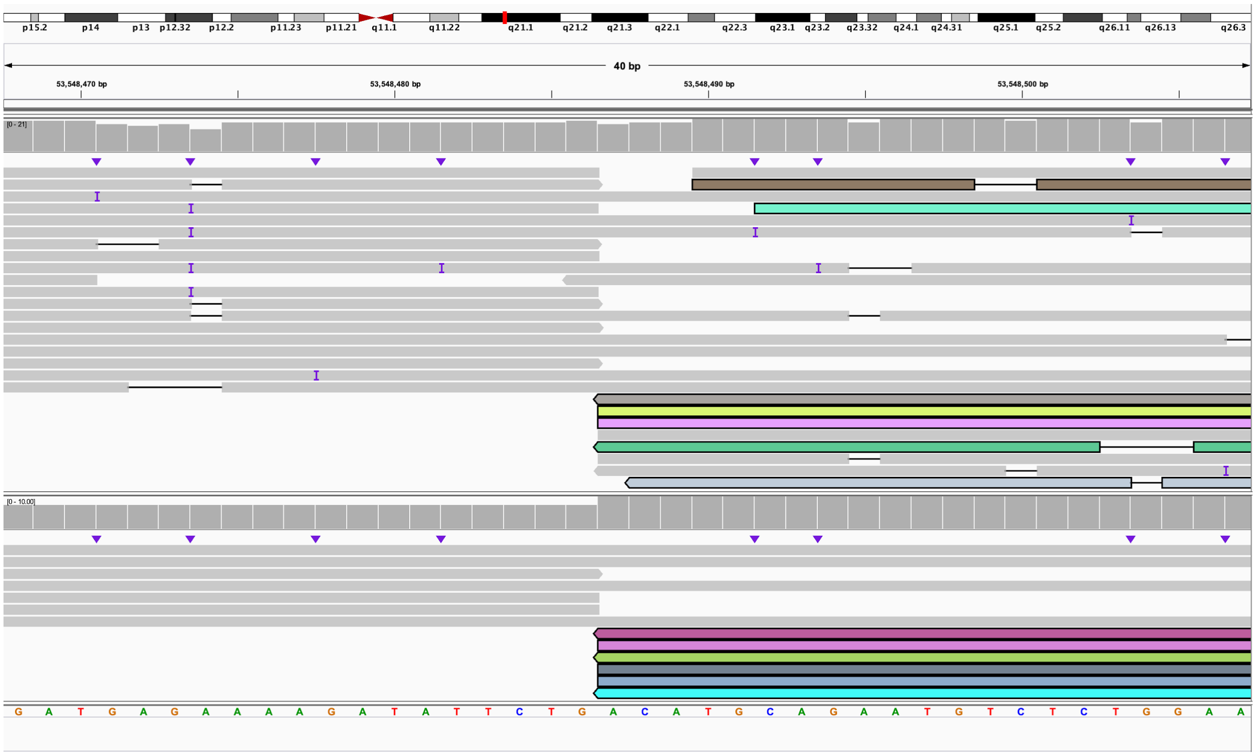

**AC.** The 3' end of 10:G (IGV: chr10:53,563,496-53,563,535) is connected to the 3' end of 4:I and bisects *XR_938881.1* (IGV: chr4:74,831,230-74,831,270). This is within derivative chromosome 14.

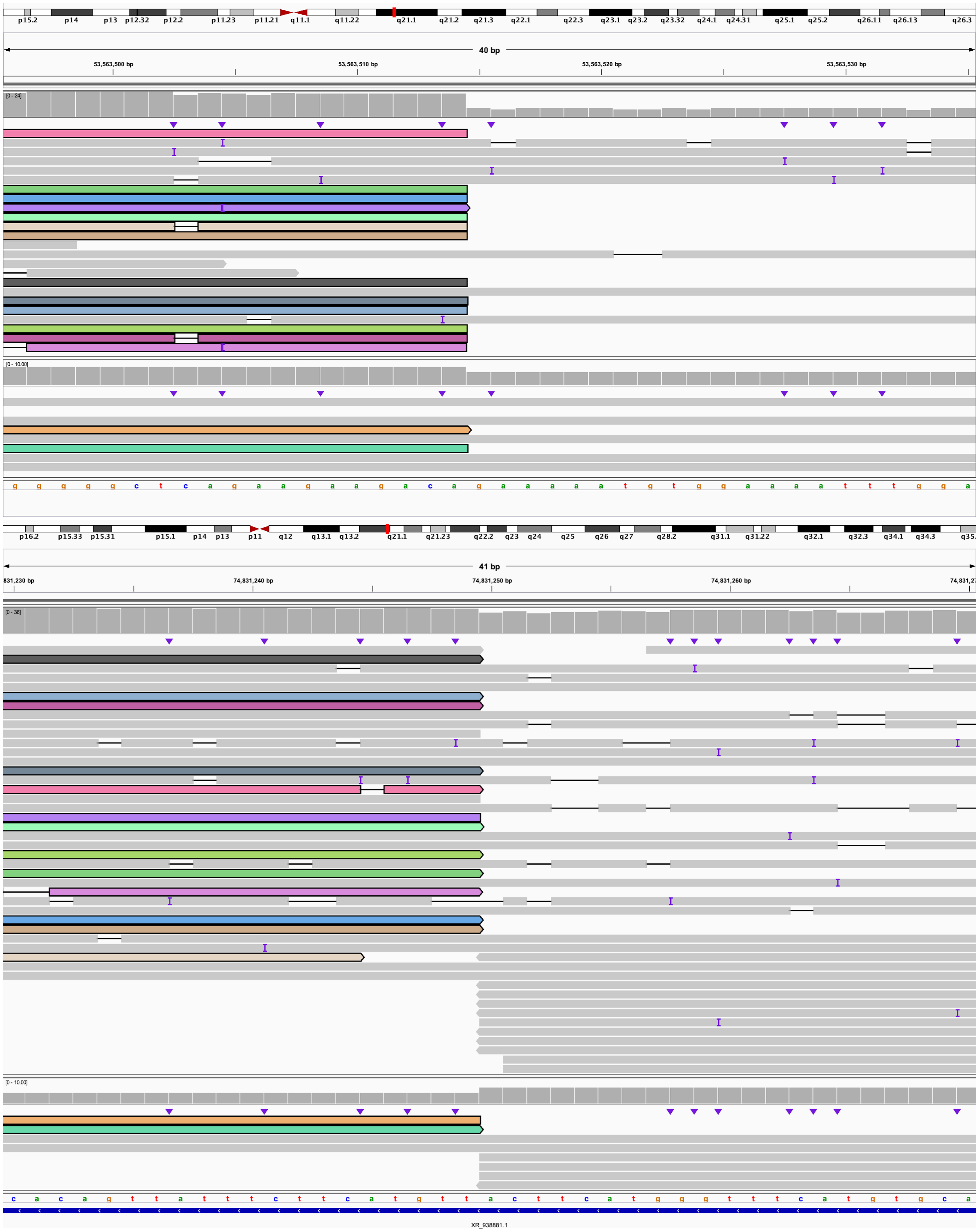

**AD.** The 5' end of 4:I bisects *BTC* (IGV: chr4:74,794,332-74,794,371) and is connected to the 5' end of 4:H and bisects *XR_001741513.1* (IGV: chr4:74,581,595-74,581,635). This is within derivative chromosome 14.

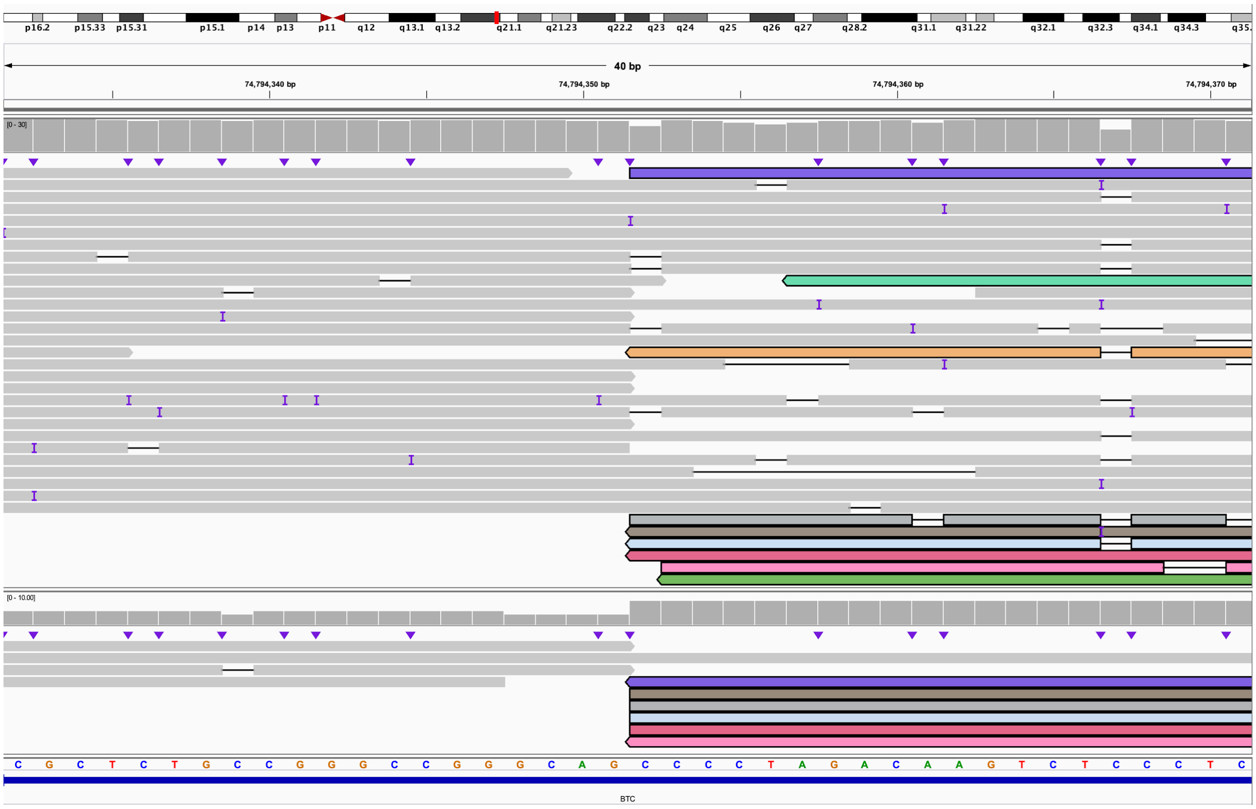

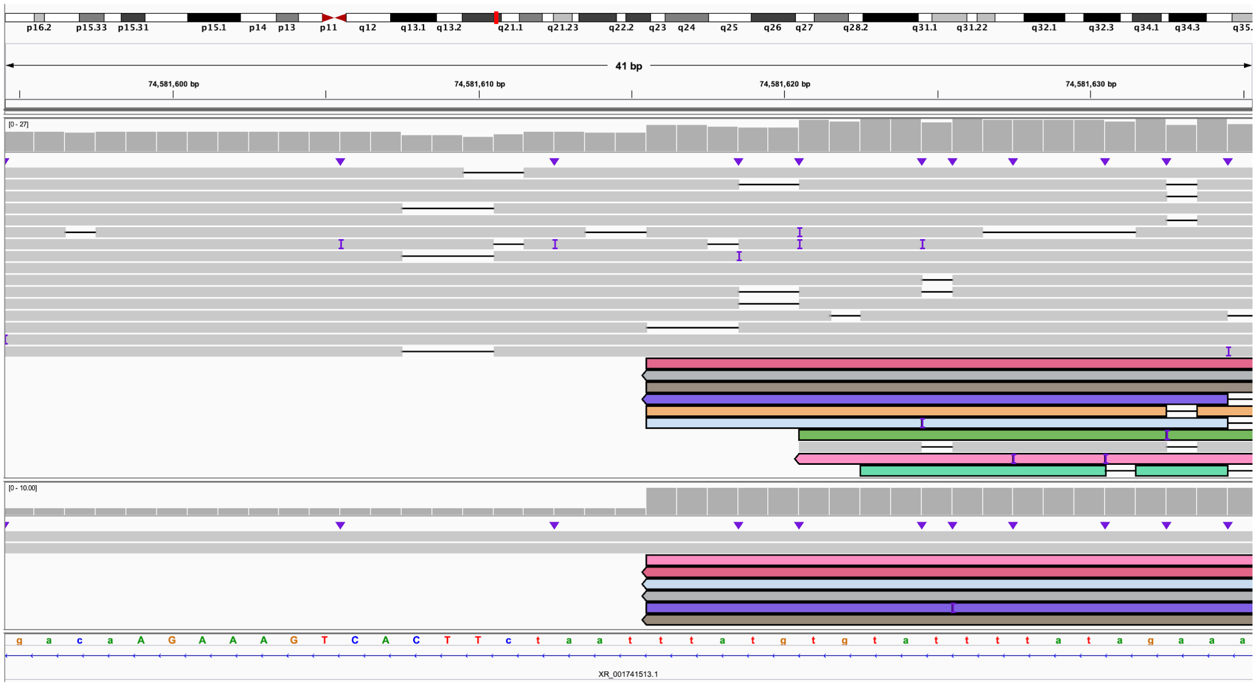

**AE.** The 3' end of 4:H bisects *BTC* (IGV: chr4:74,794,332-74,794,372) and is connected to the 3' end of 4:L (IGV: chr4:79,626,700-79,626,739). This is within derivative chromosome 14.

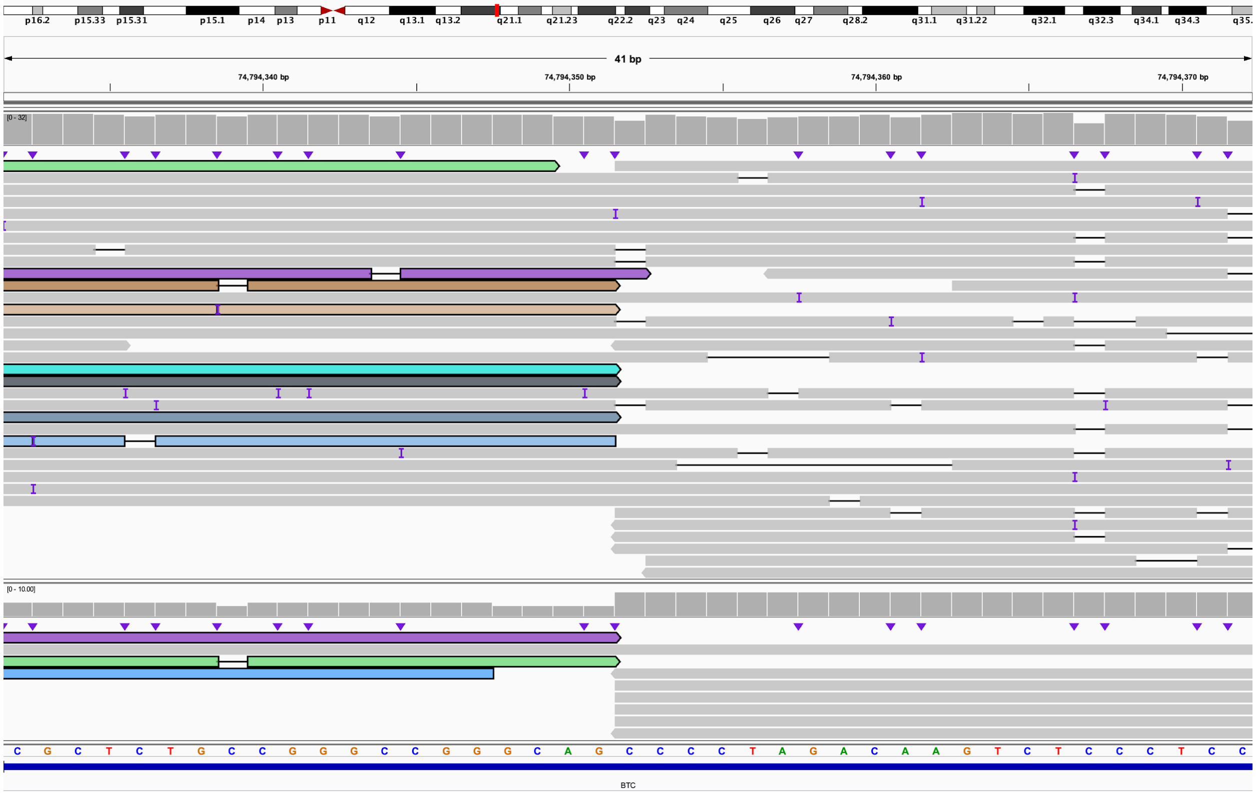

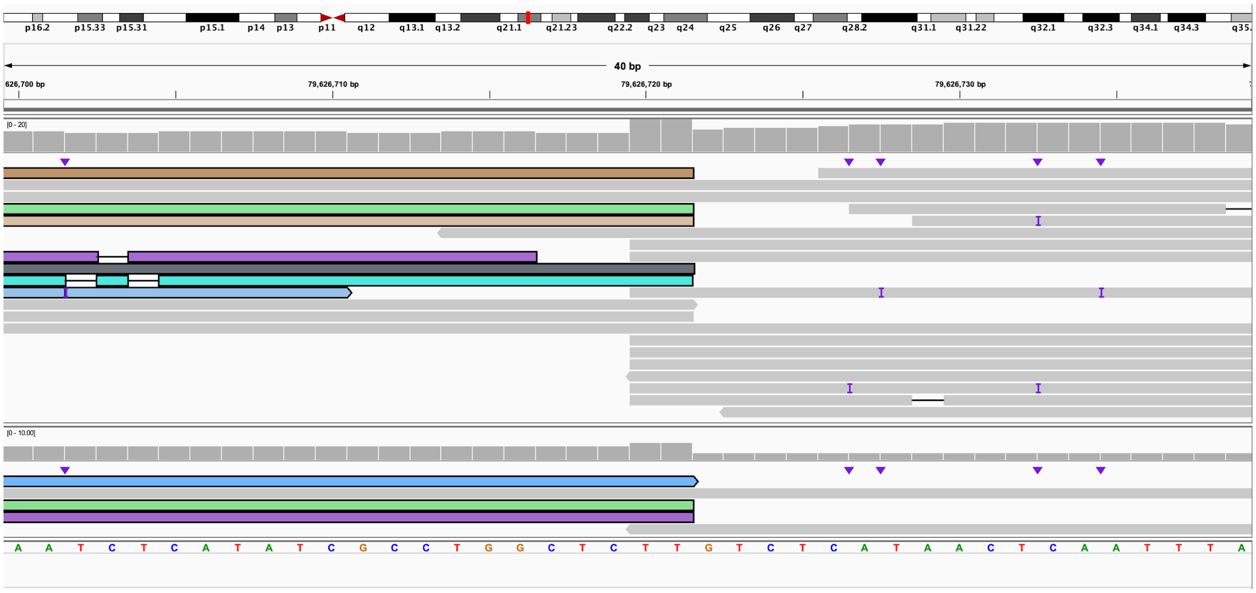

**AF.** The 5' end of 4:L bisects *SEPTIN1* (IGV: chr4:77,012,155-77,012,195) and connects to the 5' end of 4:C and bisects *CSN1S2BP* (IGV: chr4:70,140,480-70,140,520). This is within derivative chromosome 14.

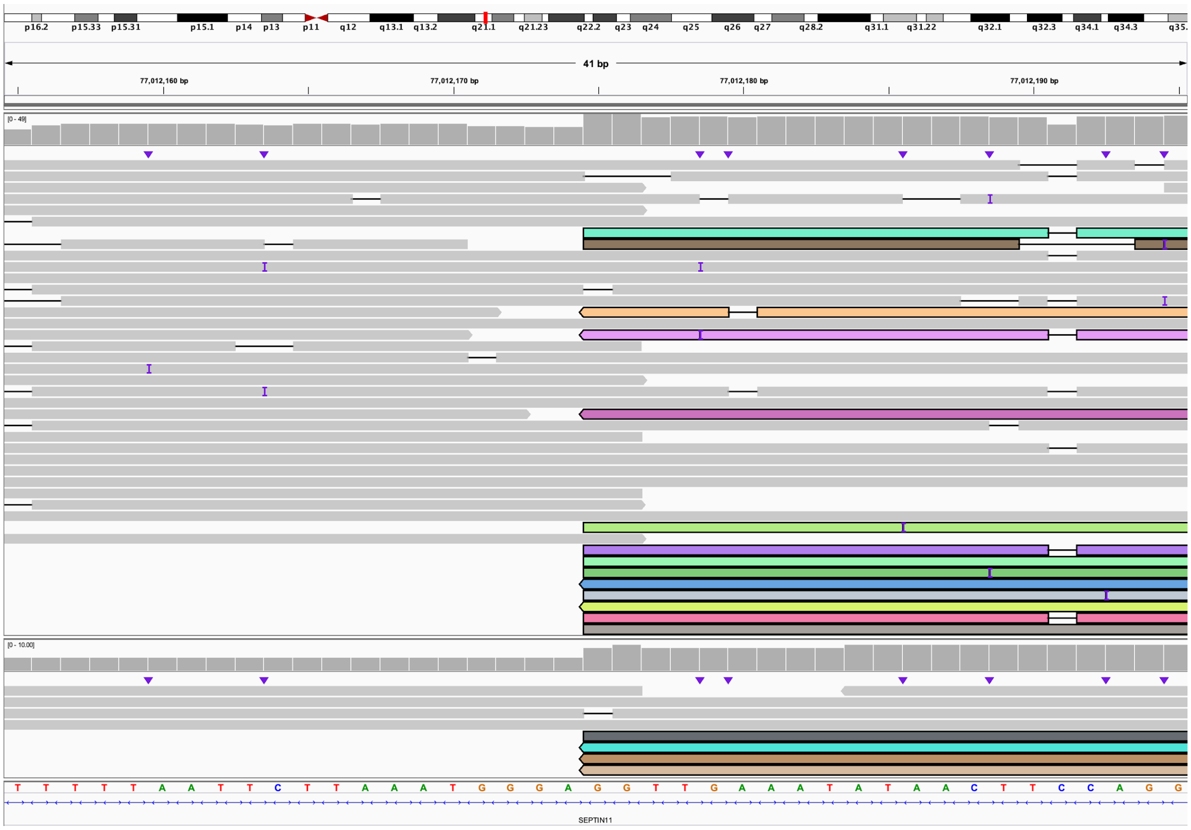

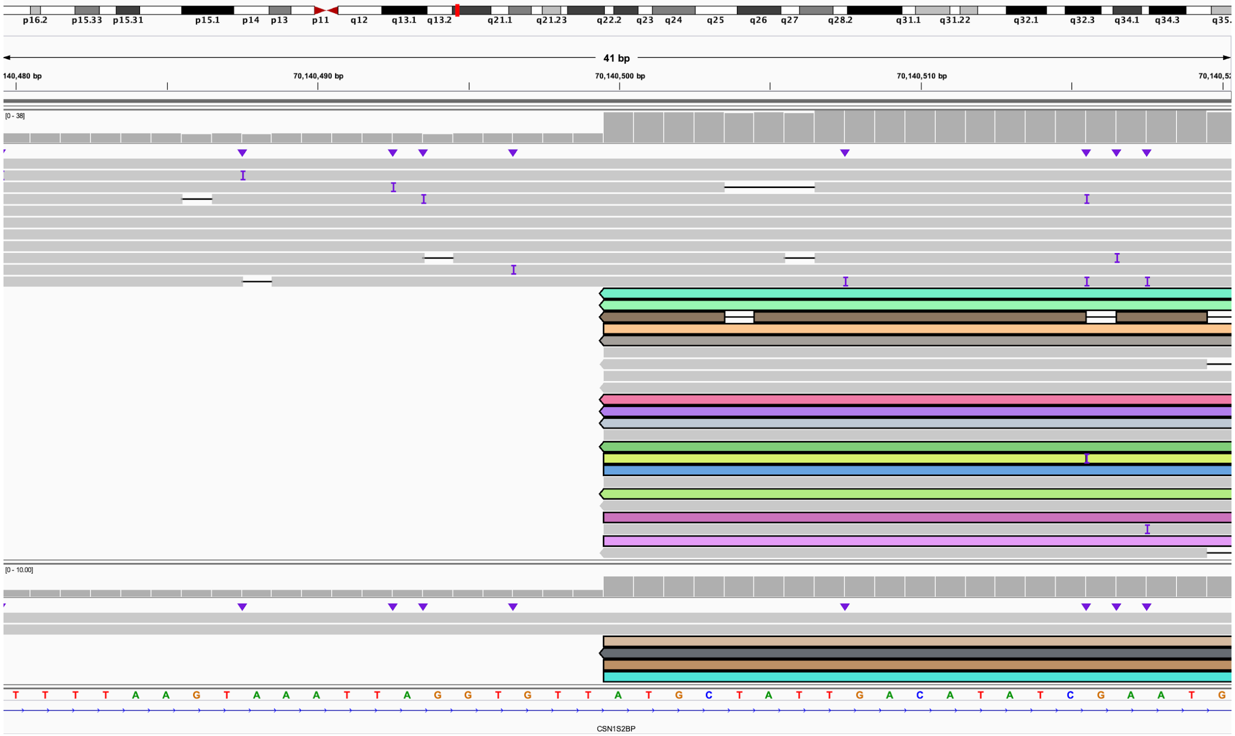

**AG.** The 3' end of 4:C bisects *COX18* (IGV: chr4:73,061,473-73,061,513) and is connected to the 3' end of 14:A and bisects *XR_110261.3* (IGV: chr14:20,934,790-20,934,830). This is the end of derivative chromosome 14.

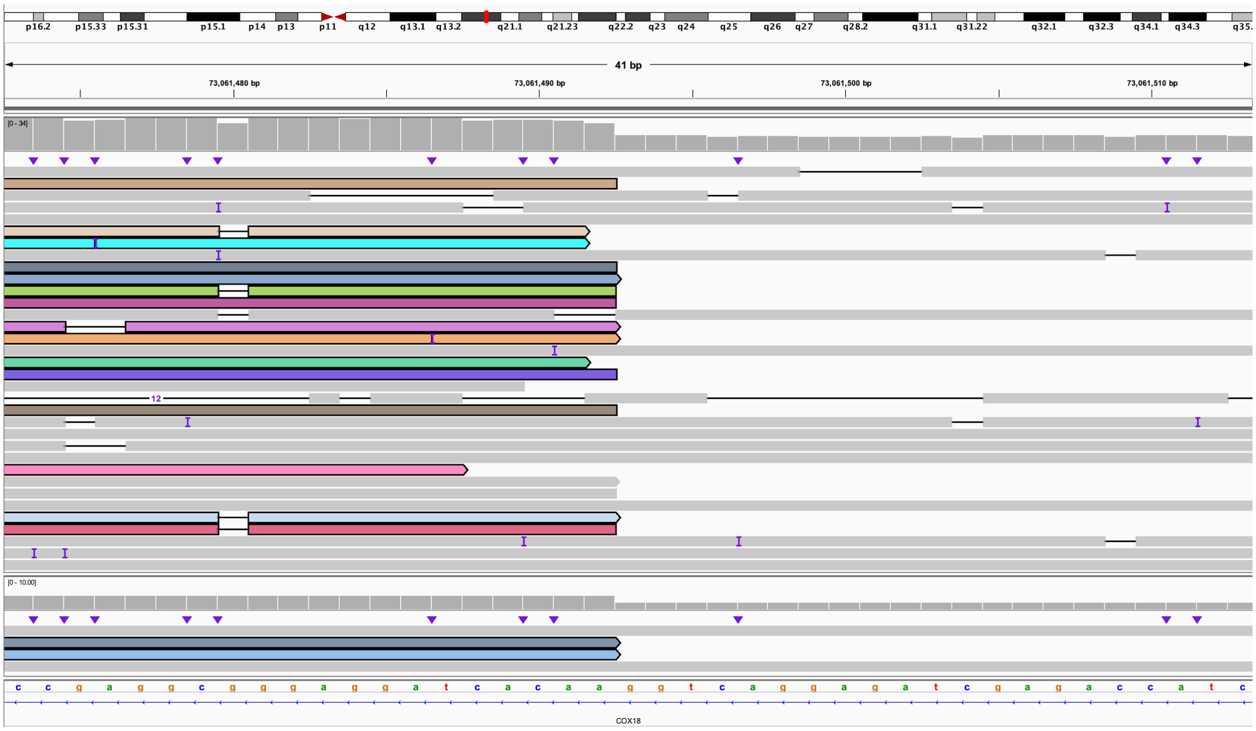

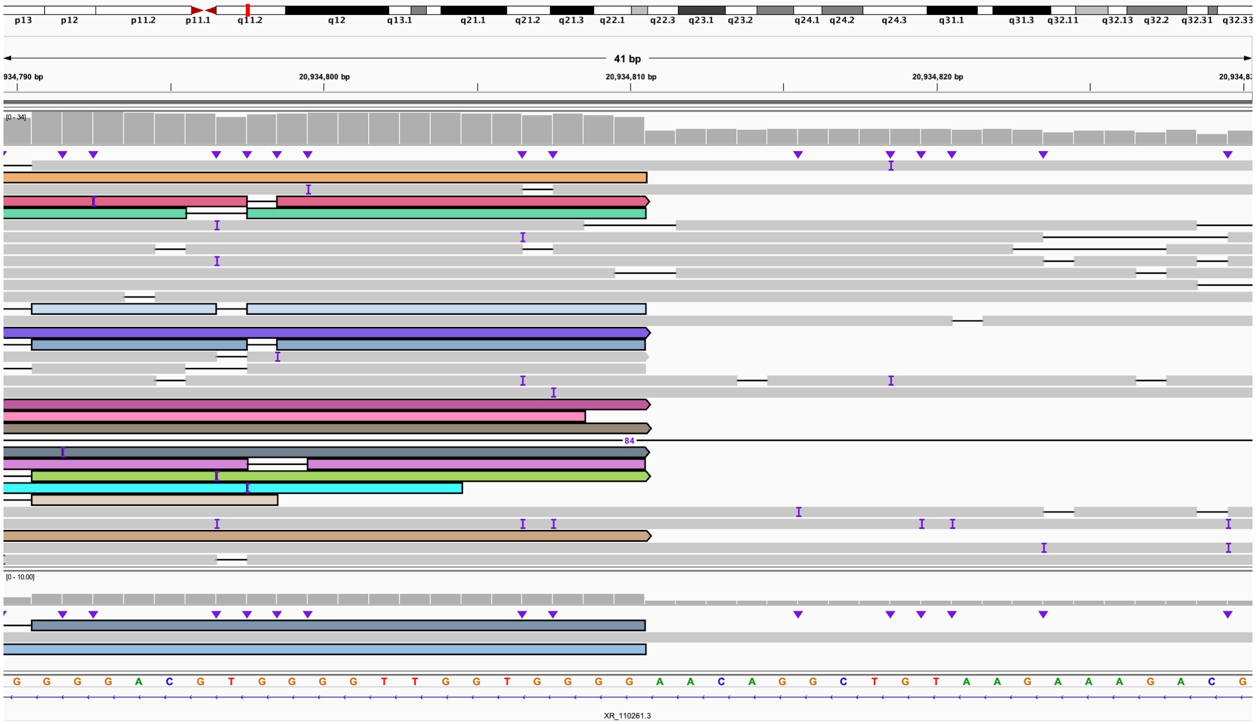

**Figure S21. S021, mosaic loss of 8p and mosaic gain of 8q.**

**A.** Coverage of chr8:1-15,000,000. This demonstrates the presence of a deletion of one chromosome from the telomere to 3,458,035. Also apparent is the mosaic region between 3,469,752 and 7,186,524, then the normal copy state after the defensin locus. The duplication around 2 Mbp is frequently observed in SNP array studies and is often not reported on the array as it is considered benign (see panels B–D for details).

**A.**

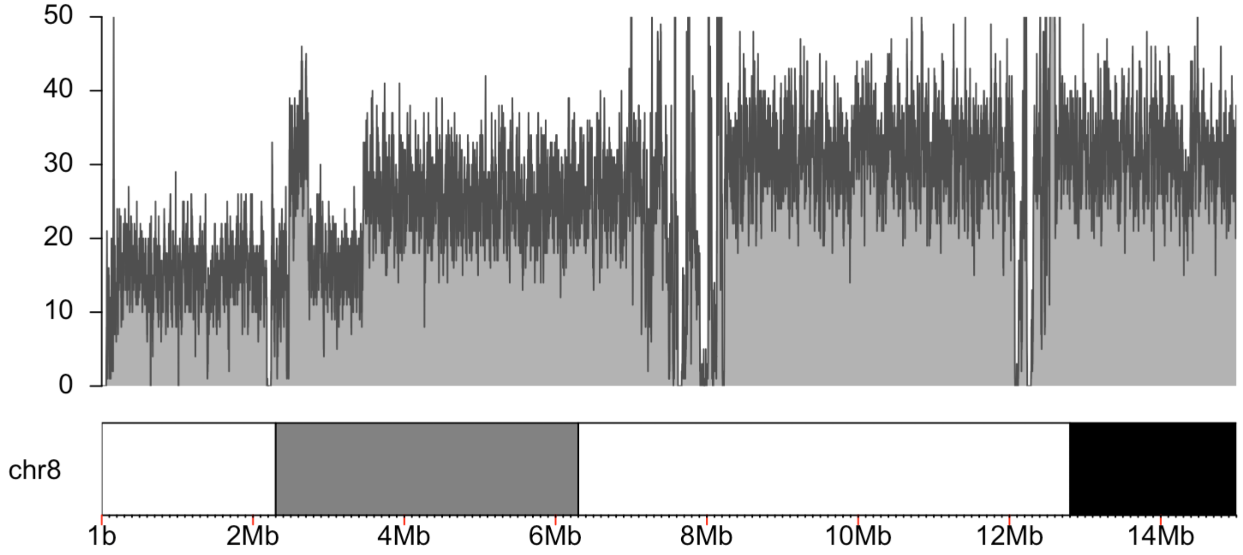

**B.** The 5' end of the duplication observed in the coverage plot in A begins at approximately chr8:2,475,083, within a VNTR. Read color shows that reads were split and re-aligned to the repetitive region. IGV view is of chr8:2,475,009-2,475,591.

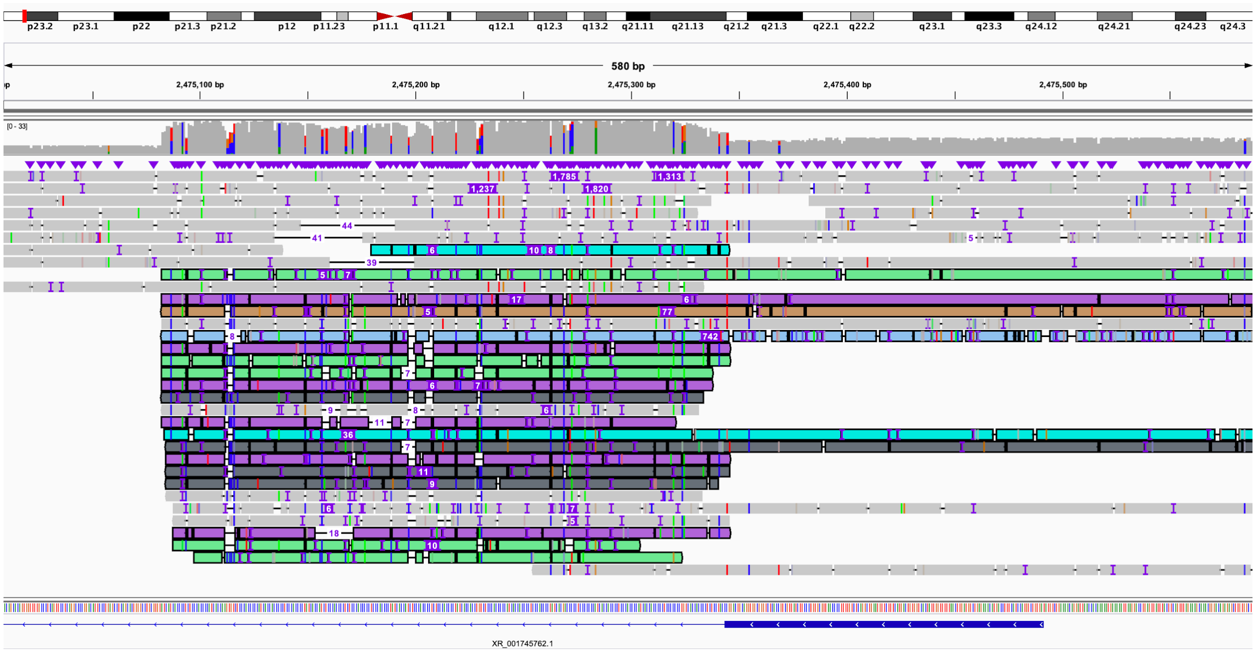

**C.** The 3' end of the duplication observed in A ends at chr8:2,729,059, meaning the duplication is approximately 254 kbp. IGV view is of chr8:2,729,039-2,729,079. Reads are linked to those shown in panel D and color is consistent between the two.

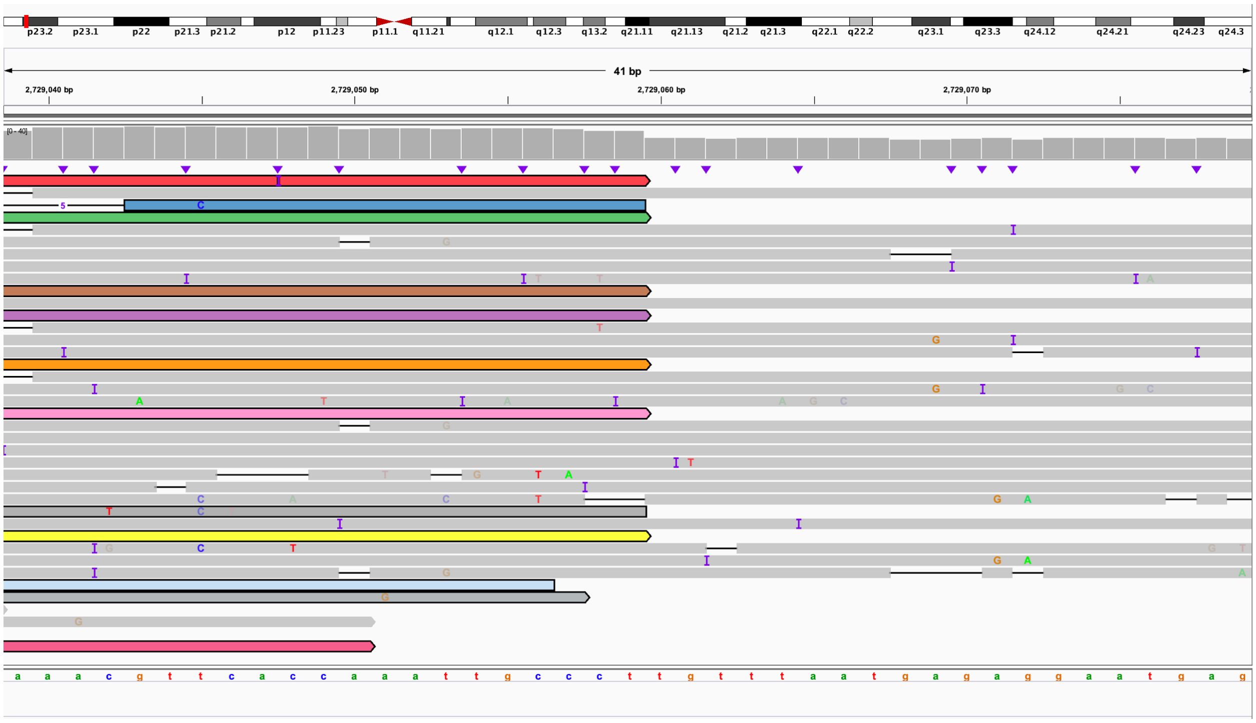

**D.** Reads from the 3' end of the duplication shown in C link to the beginning of a deletion on chr8 at chr8:2,182,175. The orientation of the reads suggests that the duplication is inverted. IGV view is of chr8:2,182,155-2,182,194.

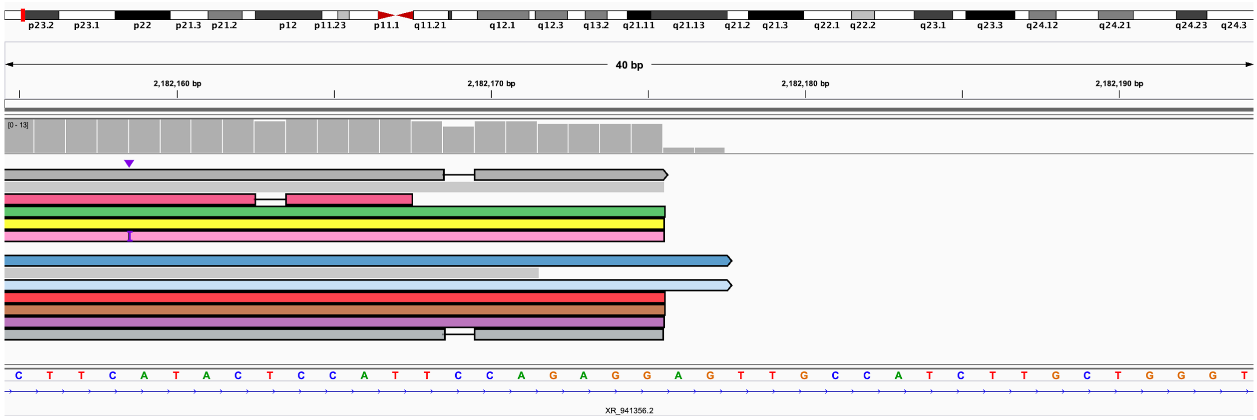

**E:** Coverage of chr8:93,000,000-98,000,000 demonstrating the increase of copy state from 2 to 3 observed on array at 95,270,423. No definitive breakpoint could be identified in the long-read data.

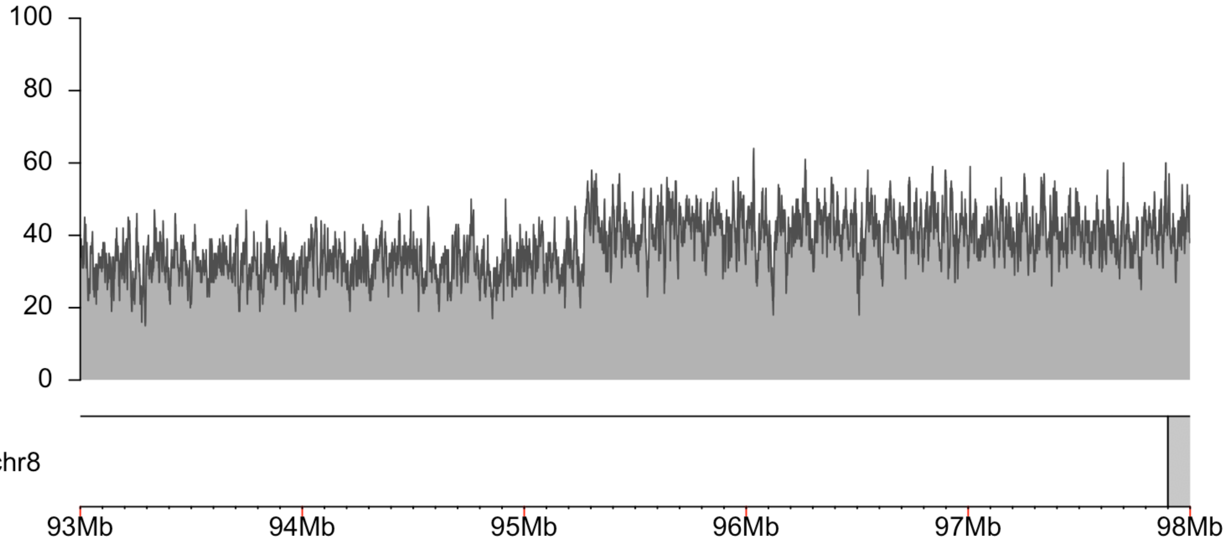

**Figure S22. S022, focal amplification of 4q with adjacent region of homozygosity, duplication of 15q11.2.**

**A.** Coverage of the 4q target region shows the presence of an amplification.

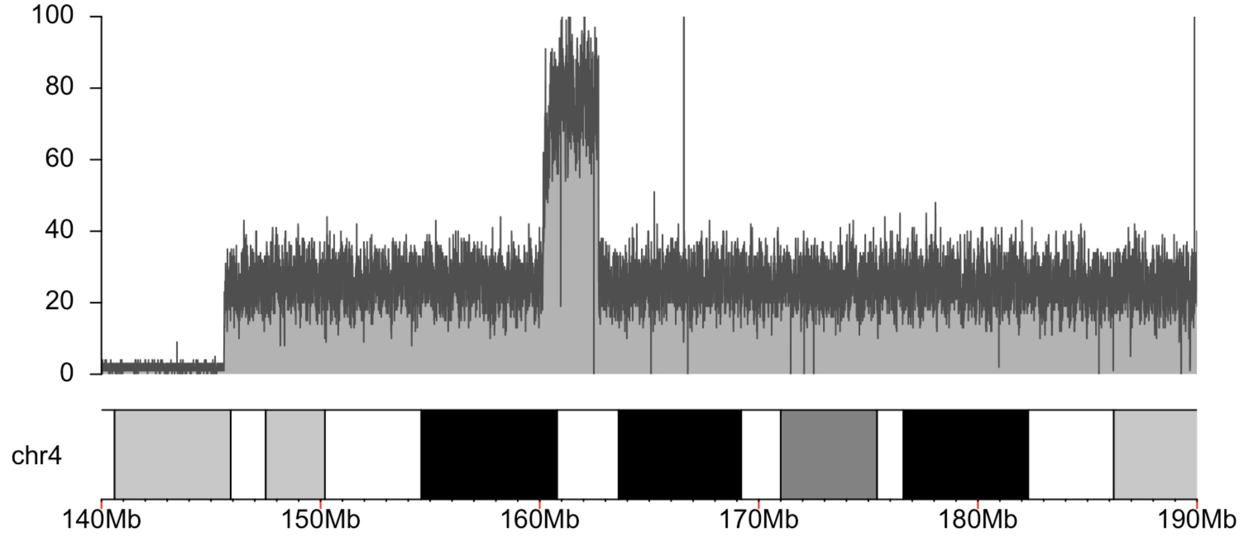

**B.** Coverage of 15q11.2 target region, regions with no coverage represent regions of the reference genome denoted as ‘N’.

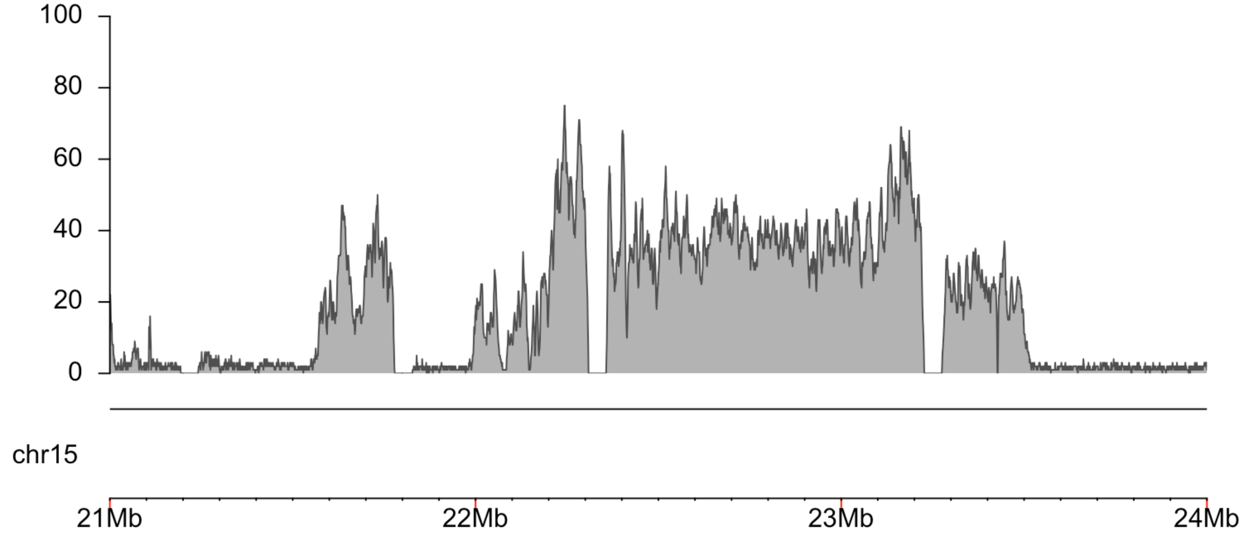

**C.** Coverage of the focal amplification of 4q.

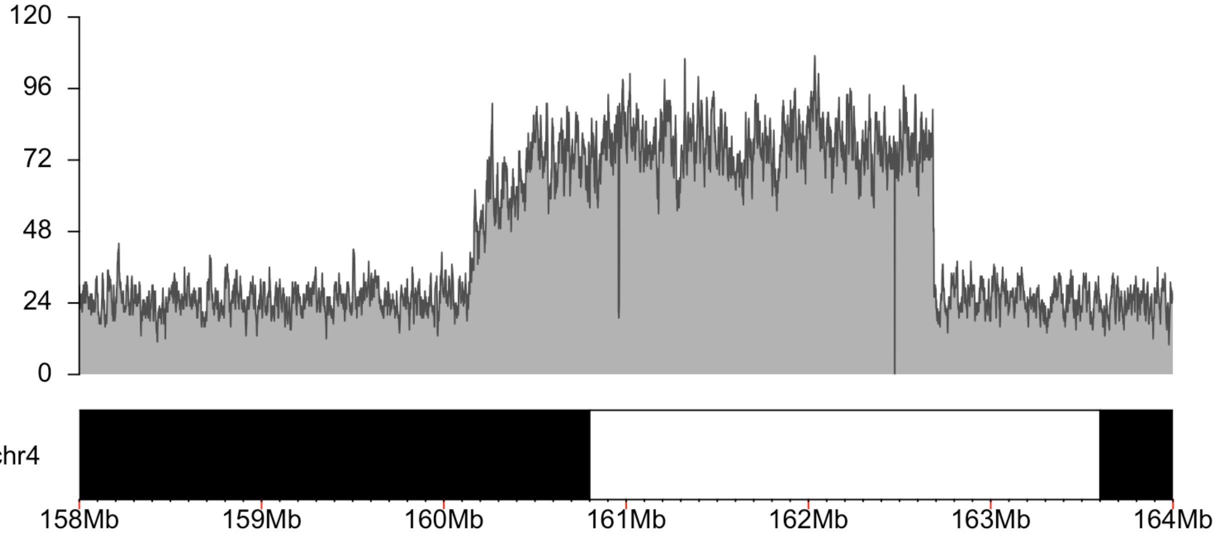

**D.** View of the beginning of the focal amplification of 4q, this is the centromere-proximal side and includes the second increase in coverage. Within this region, copy number estimate increases from 2x to 4x in a stepwise fashion (Table S9). Reads that span both breakpoints are highlighted. IGV view is of chr4:160,141,260-160,190,550.

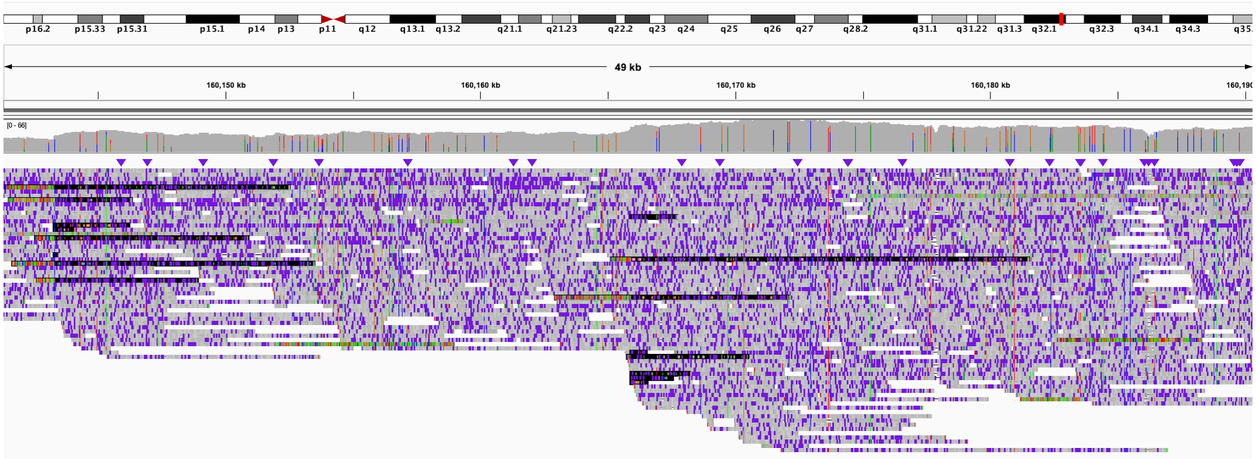

**E.** View of the third centromere-proximal breakpoint and amplification. Copy number estimate in this region increases from 4x to 5x (Table S9). IGV view is of chr4:160,248,150-160,248,189.

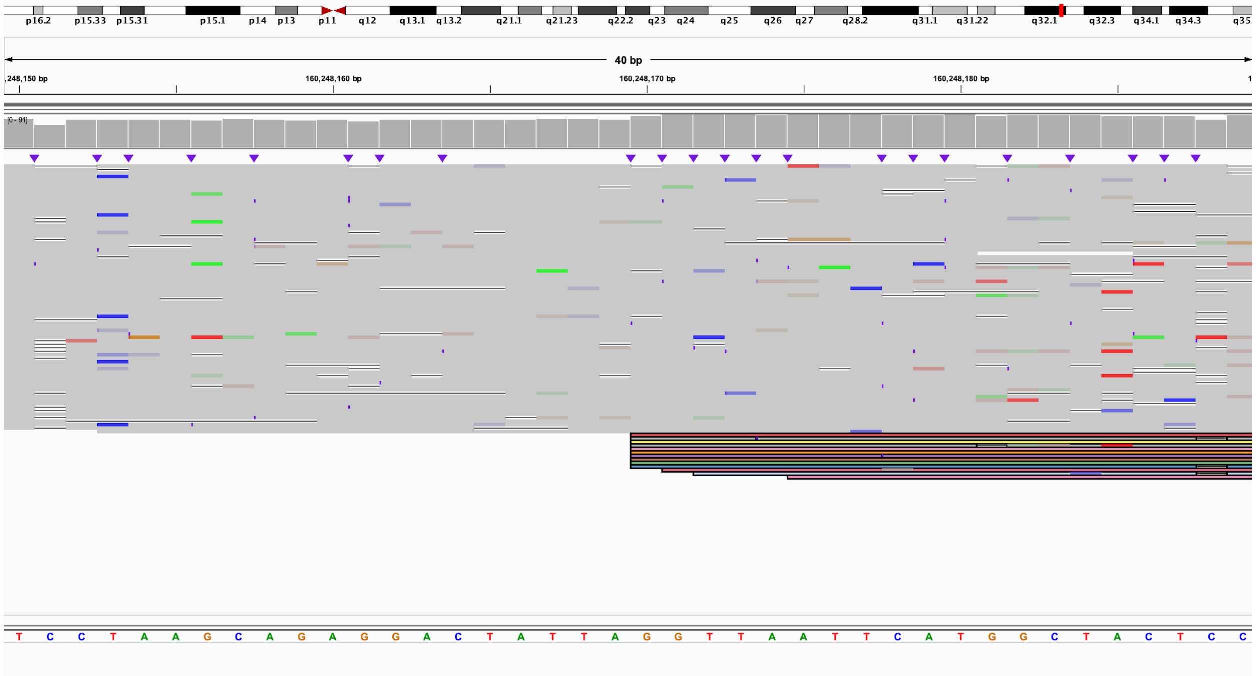

**F.** View of the fourth centromere-proximal breakpoint and amplification. Copy number estimate increases from 5x to 6x in this region (Table S9). IGV view is of chr4:160,450,935-160,450,974.

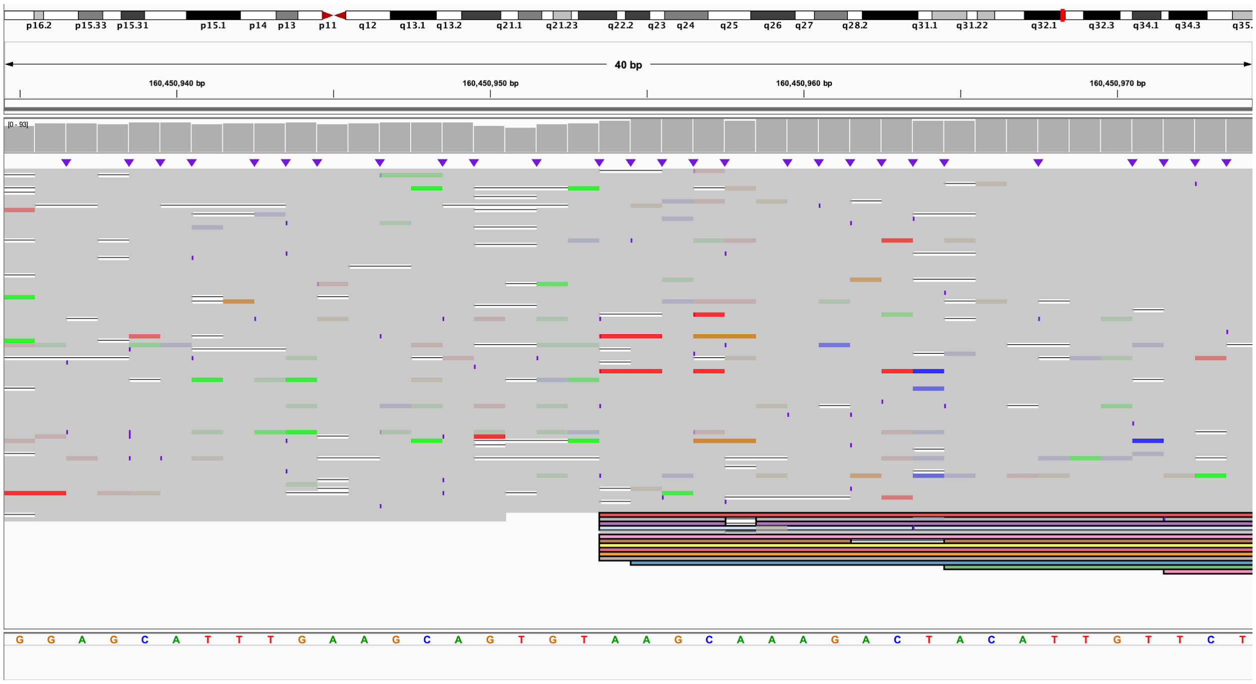

**G.** View of the telomere-proximal region of the focal amplification of 4q. Reads that span both breakpoints are highlighted. Copy number estimate decreases from 6x to 4x, then from 4x to 2x over a 3,920 bp interval (Table S9). IGV view is of chr4:162,683,982-162,690,349.

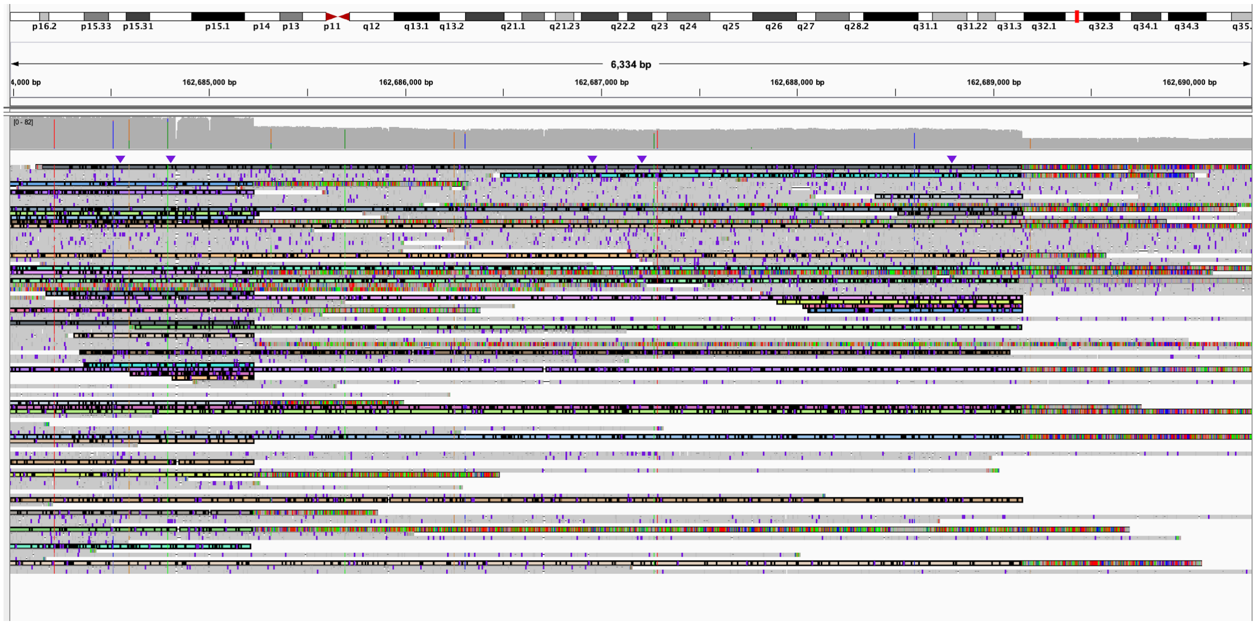

**H.** Estimate of the possible structure of the 4q amplification. We were not able to determine the exact order in which segments were duplicated. A possible arrangement of the region is shown based on this estimate.

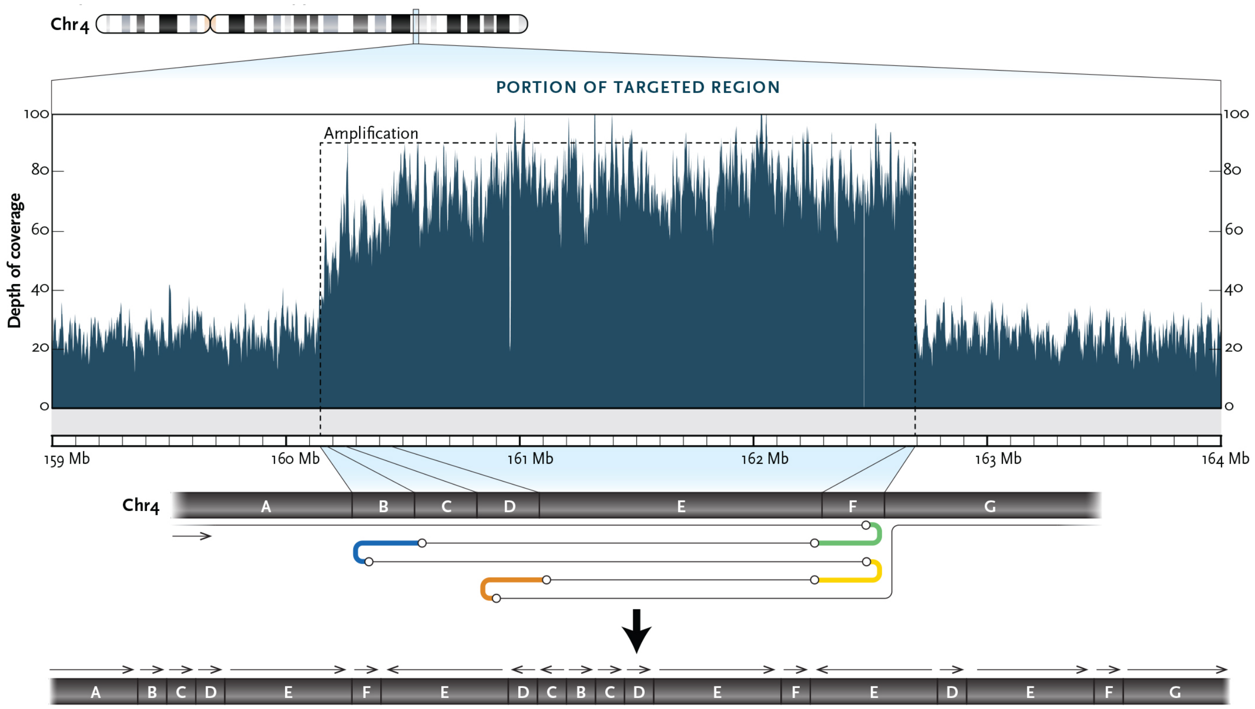

**Figure S23. S023, mosaic ring 18 present in 40% of cells.**

**A.** Coverage of chromosome 18p target region shows mosaic copy loss. **B.** Coverage of chromosome 18q target region shows mosaic copy loss.

A:

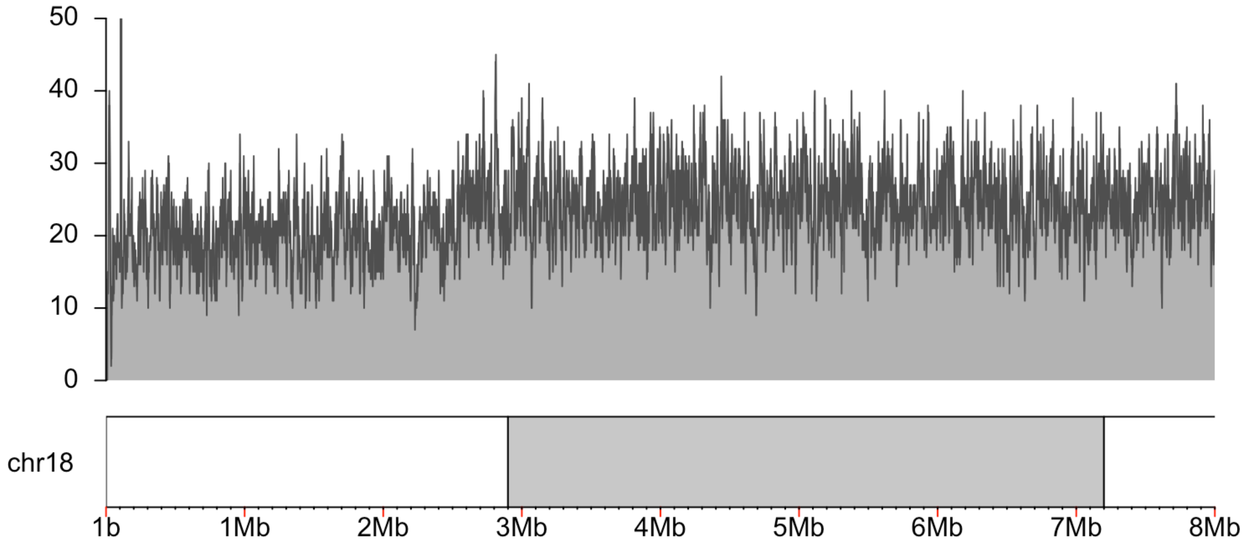

B:

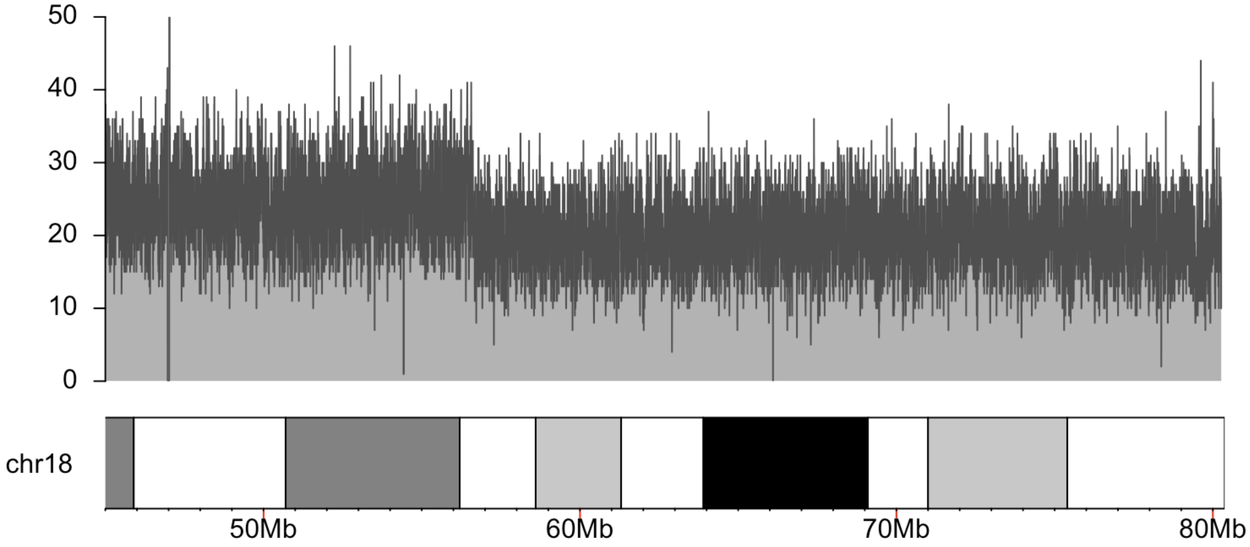

**Figure S24. S035, duplications of 8q24 and 16p13.11 identified by clinical testing.**

**A.** Coverage of chromosome 8 target region. **B.** Coverage of chromosome 16 target region.
**C.** Centromere-proximal end of the chromosome 8 duplication, reads spanning the duplication are indicated by color. IGV coordinates are chr8:141,645,041-141,645,164. **D.** Telomere-proximal end of the chromosome 8 duplication shows that the duplication occurred within a TE-dense region. Reads on the 5' end of the image span an approximately 8 kbp deletion. Reads colored as in C. IGV coordinates are chr8:143,617,985-143,620,407.

A:

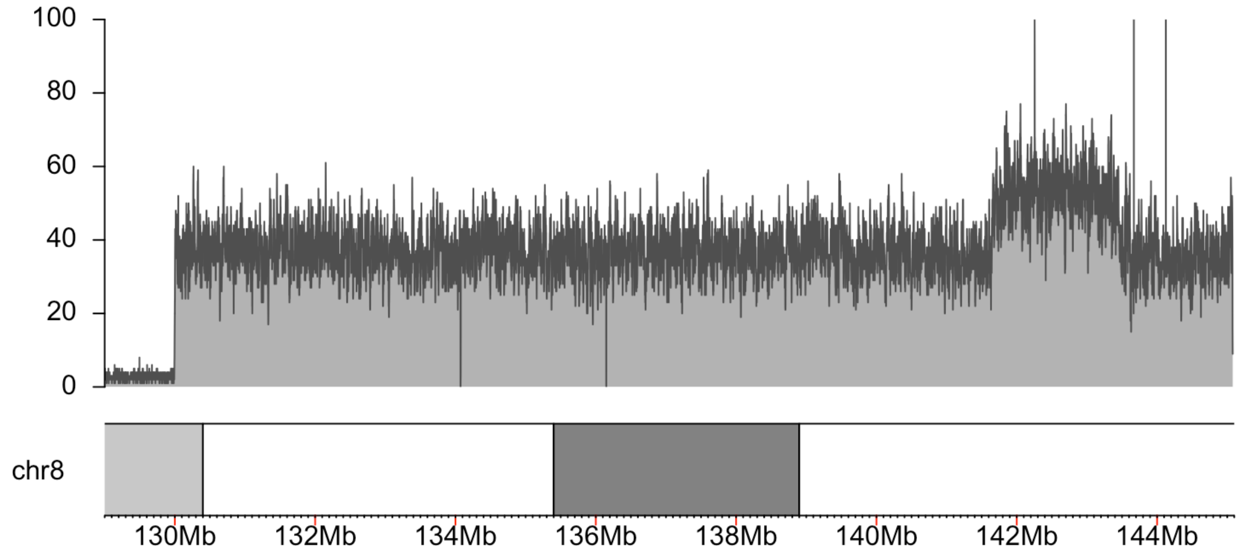

B:

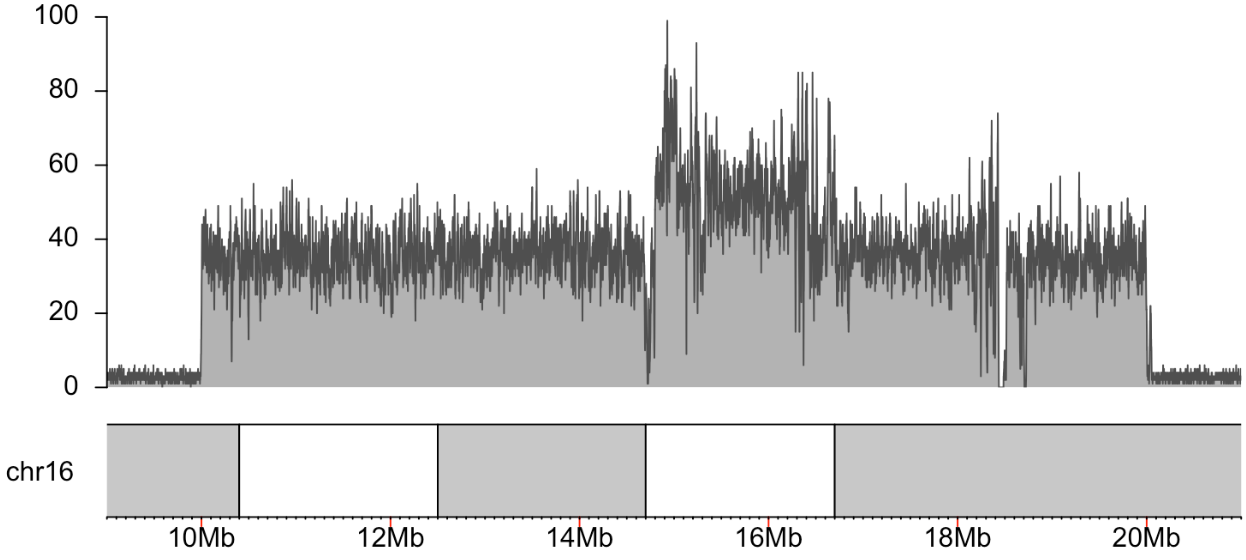

C:

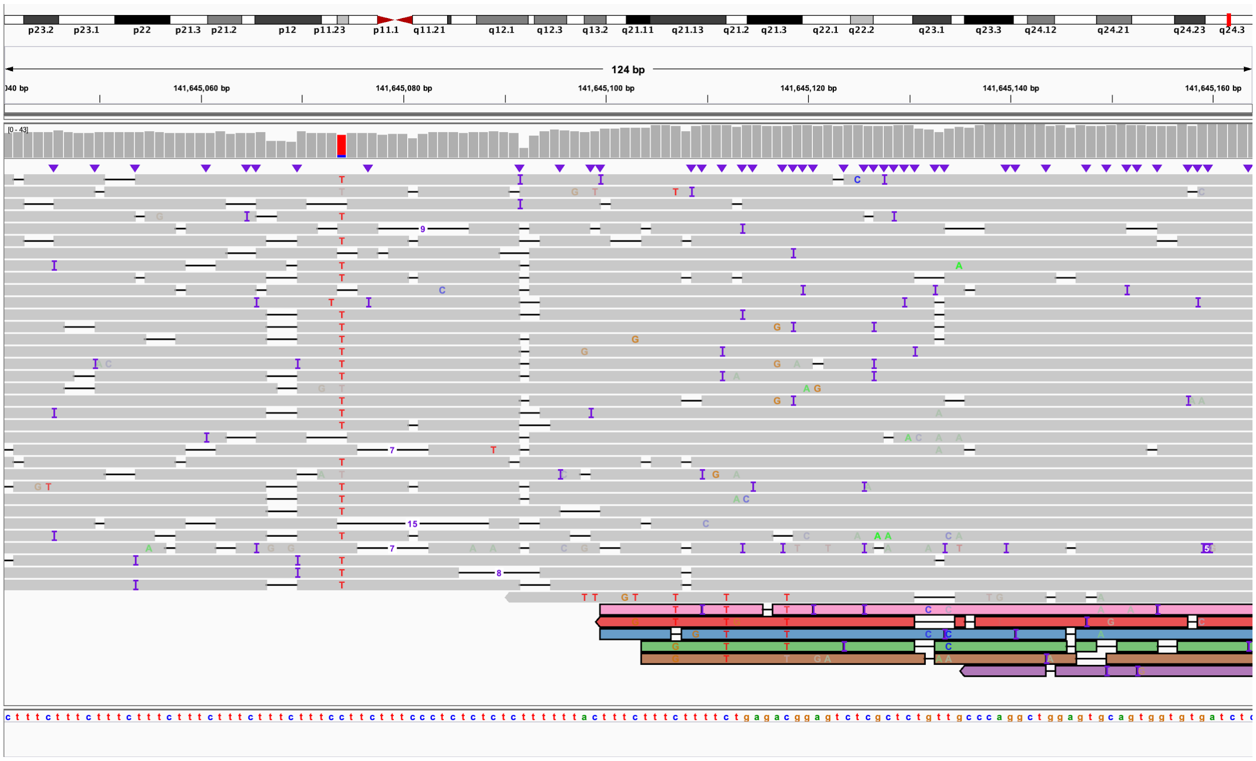
p23.2
p23.1
p22
p21.3
p21.2
p12
p11.23
p11.1
q11.21
q12.1
q12.3
q13.2
q21.11
q21.13
q21.2
q21.3
q22.1
q22.2
q23.1
q23.3
q24.12
q24.21
q24.23
q24.3
124 bp
141,645,060 bp
141,645,080 bp
141,645,100 bp
141,645,120 bp
141,645,140 bp
141,645,160 bp
[0 - 43]

D:

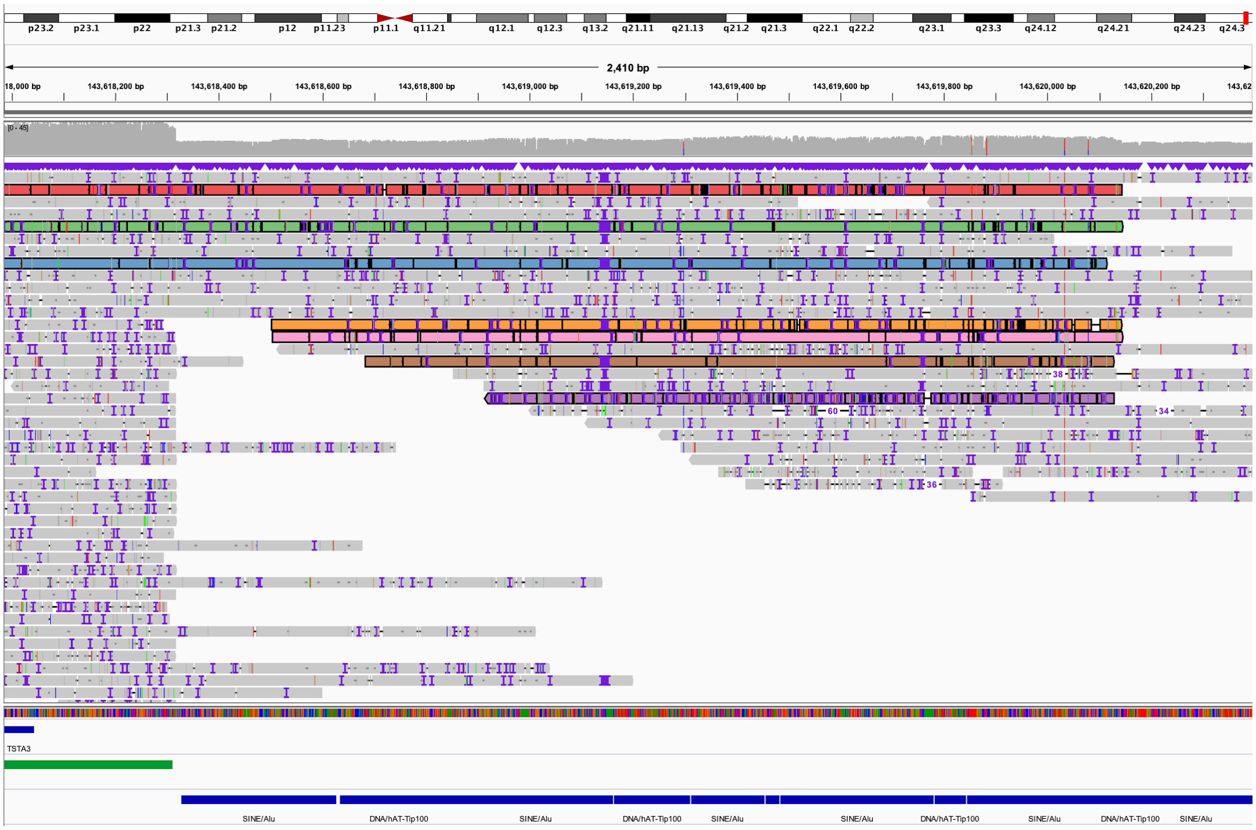
p23.2
p23.1
p22
p21.3
p21.2
p12
p11.23
p11.1
q11.21
q12.1
q12.3
q13.2
q21.11
q21.13
q21.2
q21.3
q22.1
q22.2
q23.1
q23.3
q24.12
q24.21
q24.23
q24.3
2,410 bp
143,618,200 bp
143,618,400 bp
143,618,600 bp
143,618,800 bp
143,619,000 bp
143,619,200 bp
143,619,400 bp
143,619,600 bp
143,619,800 bp
143,620,000 bp
143,620,200 bp
[0 - 45]
TSTA3
SINE/Alu
DNA/hAT-Tip100
SINE/Alu
DNA/hAT-Tip100
SINE/Alu
SINE/Alu
DNA/hAT-Tip100
SINE/Alu
DNA/hAT-Tip100
SINE/Alu

### Figure S25. S036, individual with multiple rearrangements and translocations of chromosomes 5, 6, 10, and 18.

**A.** In a child with multiple deletions of chromosome 10, karyotyping revealed translocations between chromosomes 6 and 18 along with a pericentric inversion of chromosome 10. LRS identified additional translocations involving chromosomes 10 and 5 as well as several additional intrachromosomal rearrangements. Rearrangements can be resolved by following the subway plot for each derivative chromosome. Estimates of the reconstructed chromosome sizes are shown along with where they are estimated to correspond to on the karyotype (colored boxes). **B.** Coverage of chromosome 10p target region from the first Read Until run. **C.** Coverage of chromosome 10q target region from the first Read Until run. **D.** Coverage of chromosome 6 target region from the first Read Until run. **E.** Coverage of chromosome 18 target region from the first Read Until run. **F.** Coverage of chromosome 5 target region from the second Read Until run. **G.** Coverage of chromosome 6 target region from the second Read Until run. **H.** Coverage of chromosome 18 target region from the second Read Until run. **I.** Karyotype of the patient, arrows denote derivative chromosomes. **J–W.** IGV screenshots of rearrangement, translocation, or deletion junctions. Reads are colored to indicate where they map to between both screenshots. Coordinates for each view are given. J–M represents derivative chromosome 10, N–R represents derivative chromosome 5, S and T show derivative chromosome 6, and U–W show derivative chromosome 18.

**A:**

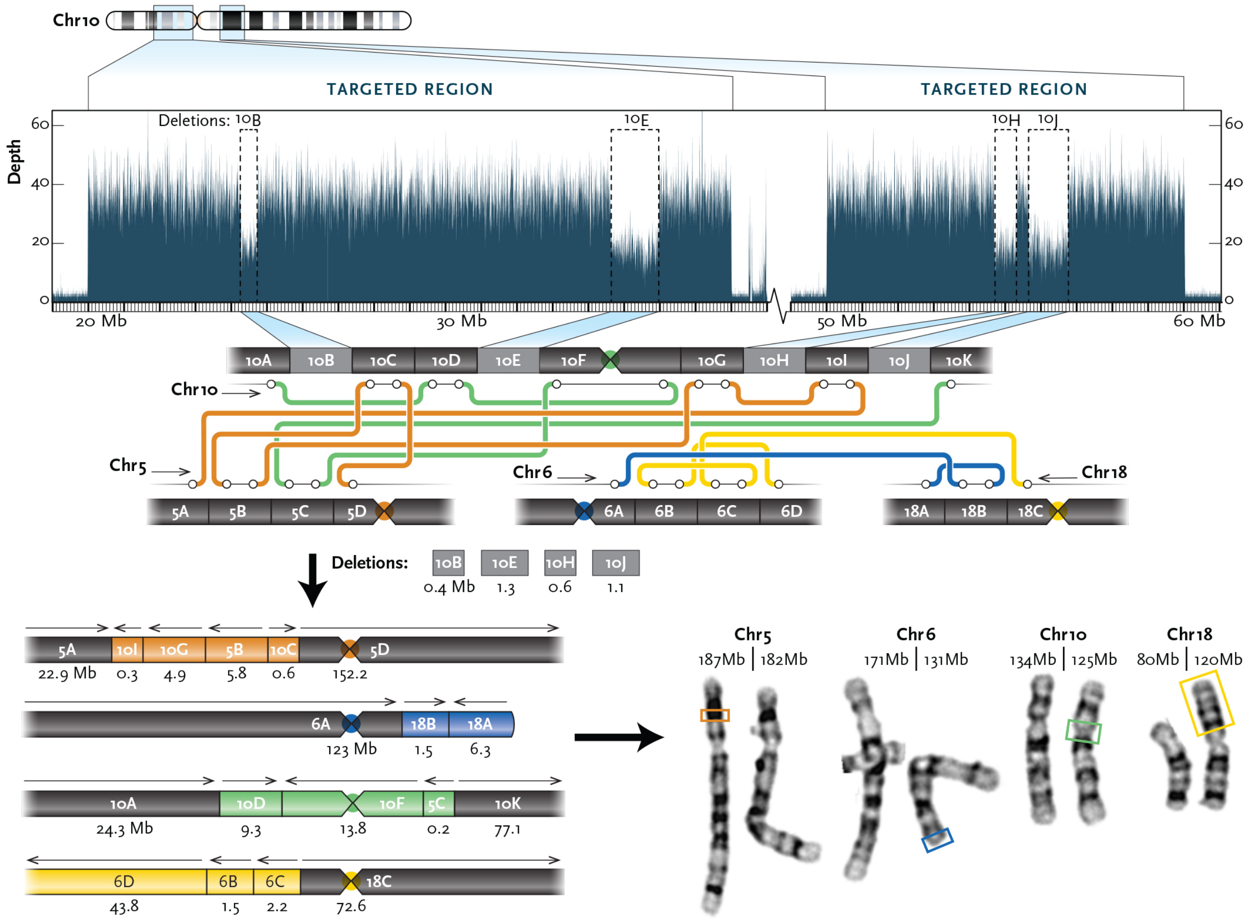

B:

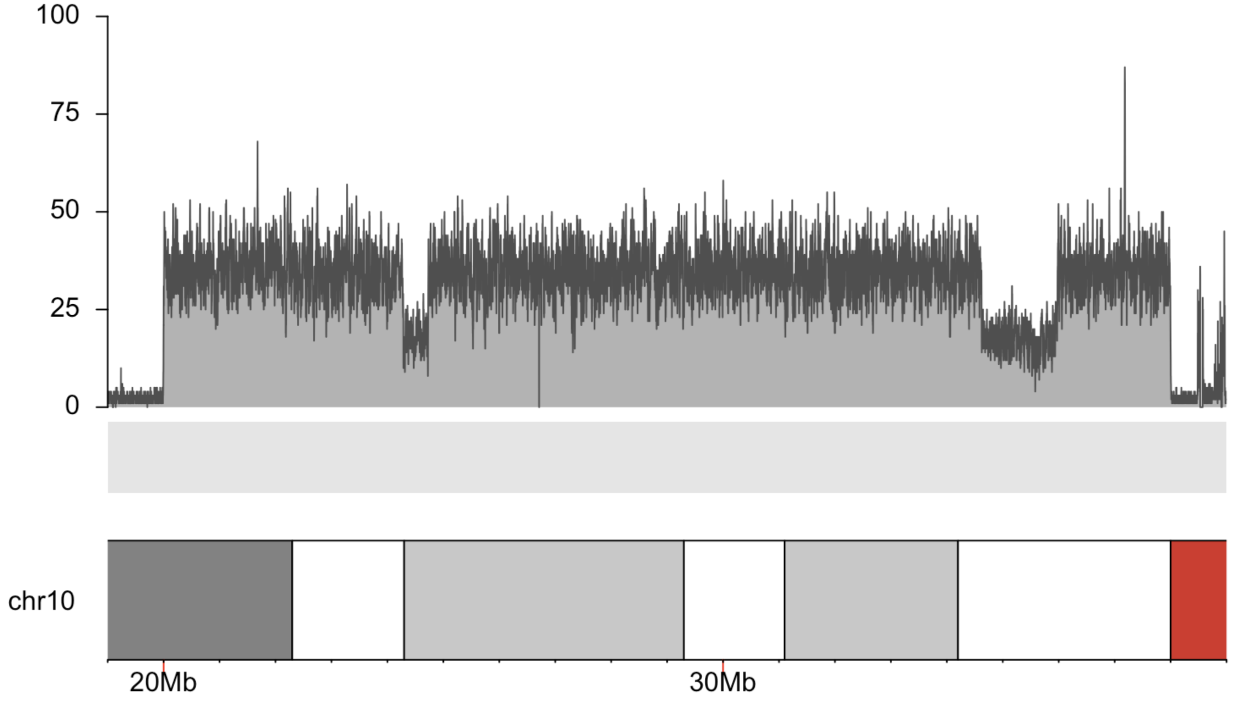

C:

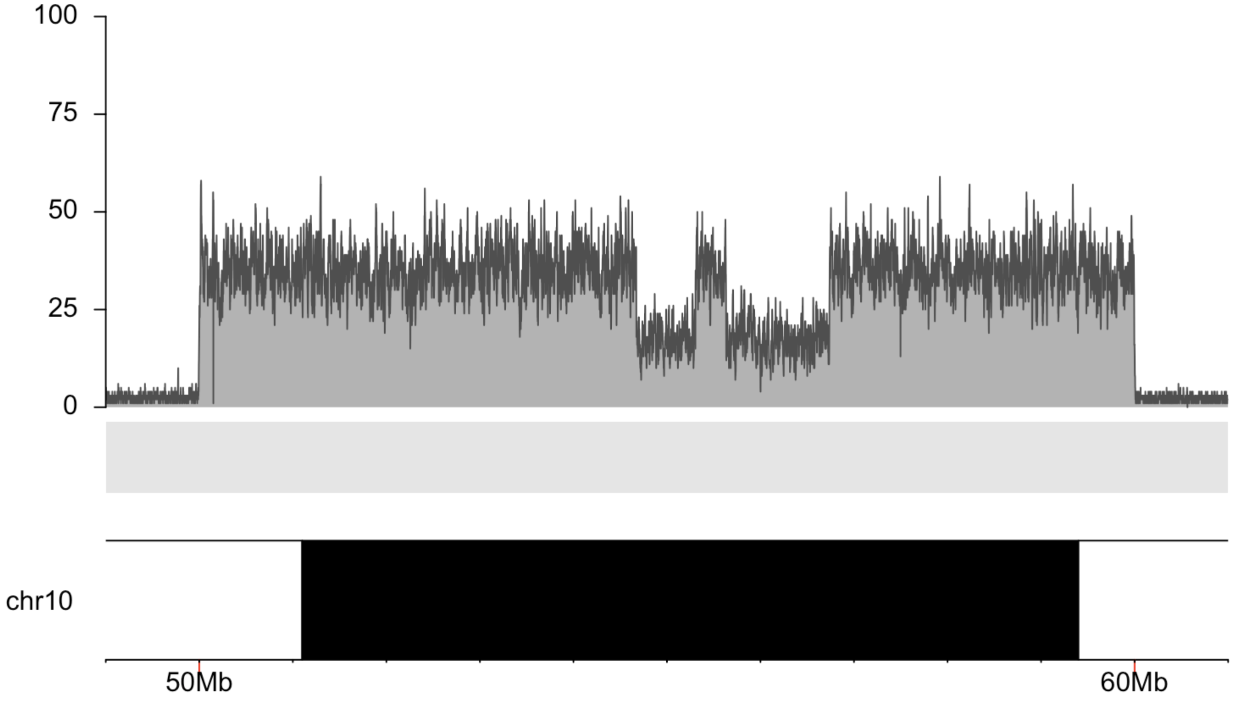

D:

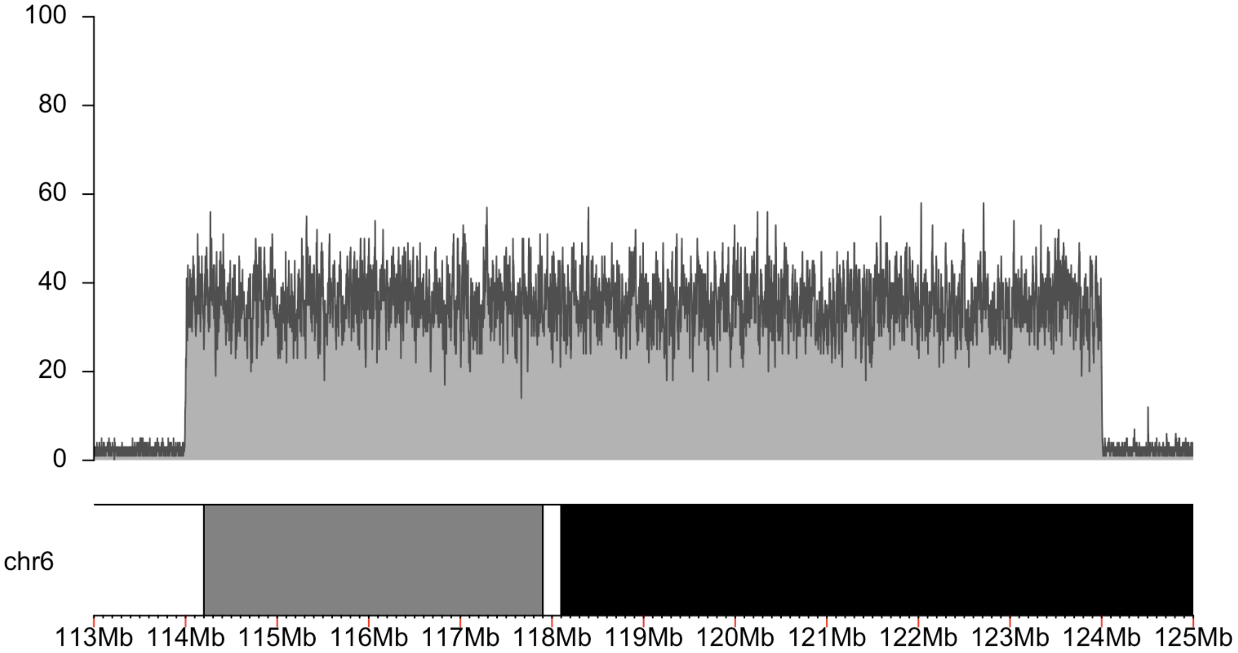
100
80
60
40
20
0
chr6
113Mb 114Mb 115Mb 116Mb 117Mb 118Mb 119Mb 120Mb 121Mb 122Mb 123Mb 124Mb 125Mb

E:

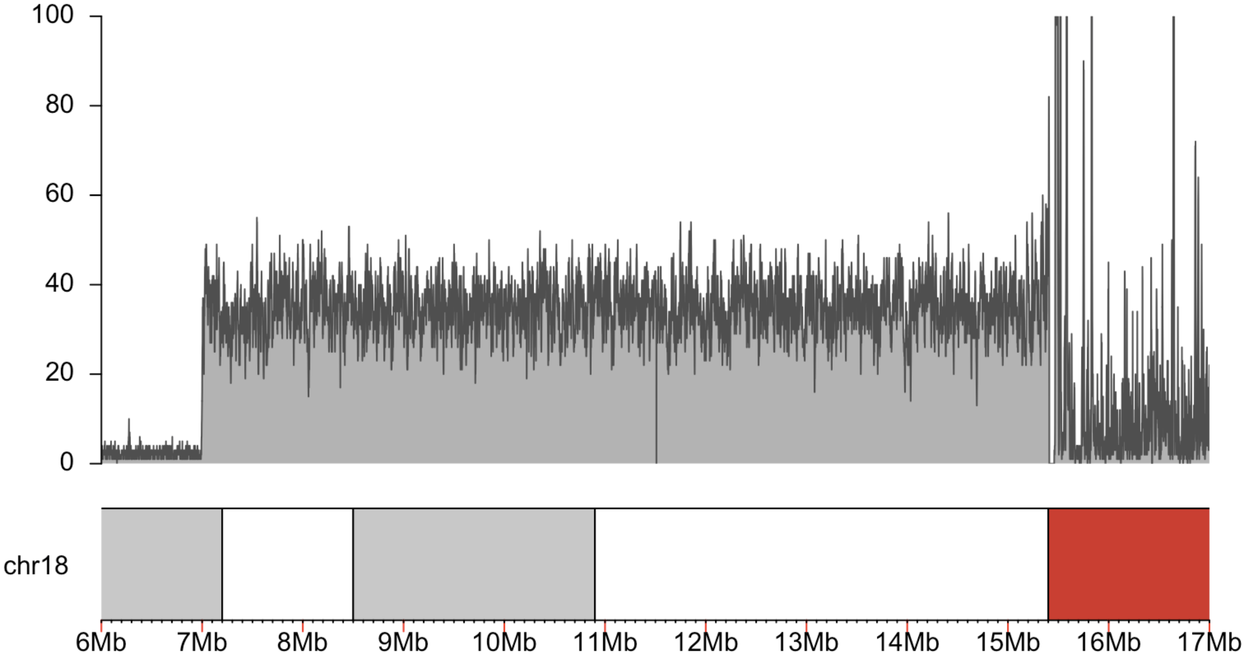
100
80
60
40
20
0
chr18
6Mb 7Mb 8Mb 9Mb 10Mb 11Mb 12Mb 13Mb 14Mb 15Mb 16Mb 17Mb

**F:**

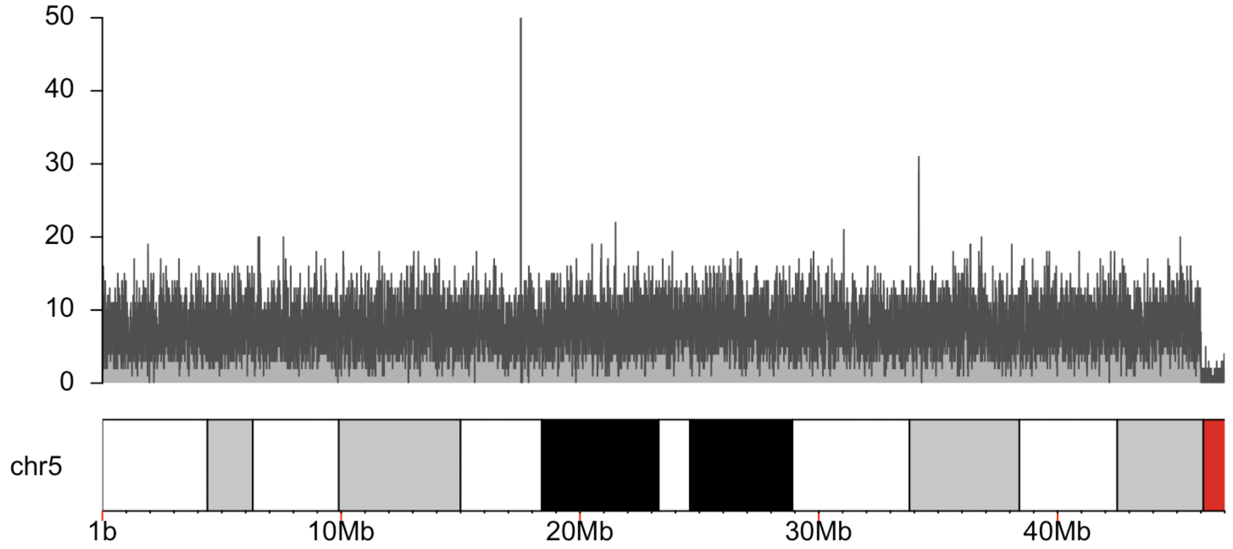

50
40
30
20
10
0
chr5
1b
10Mb
20Mb
30Mb
40Mb

**G:**

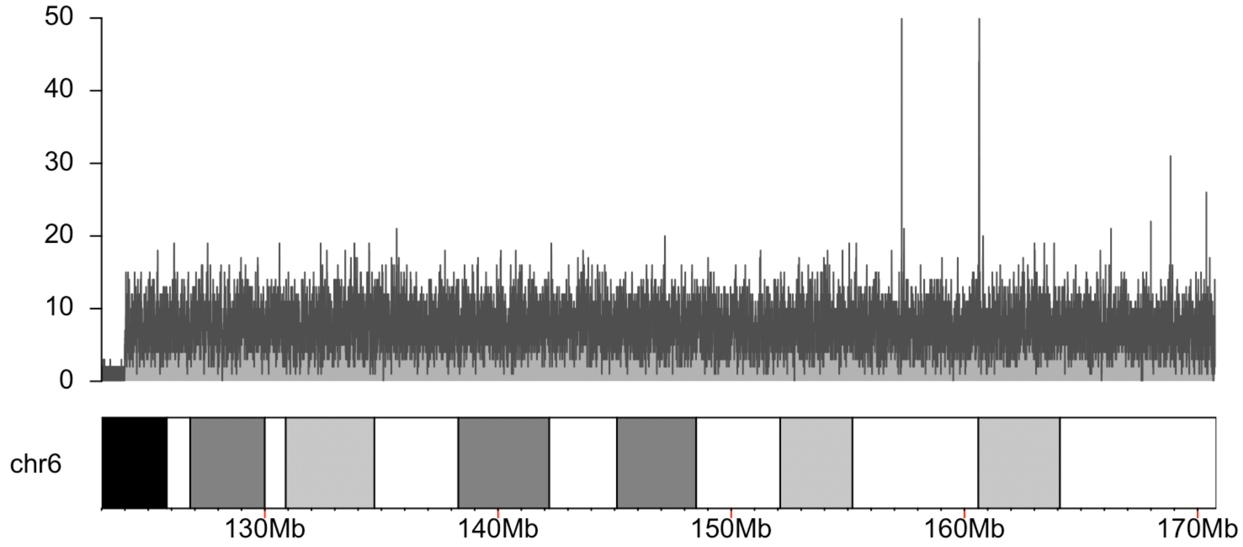

50
40
30
20
10
0
chr6
130Mb
140Mb
150Mb
160Mb
170Mb

H:

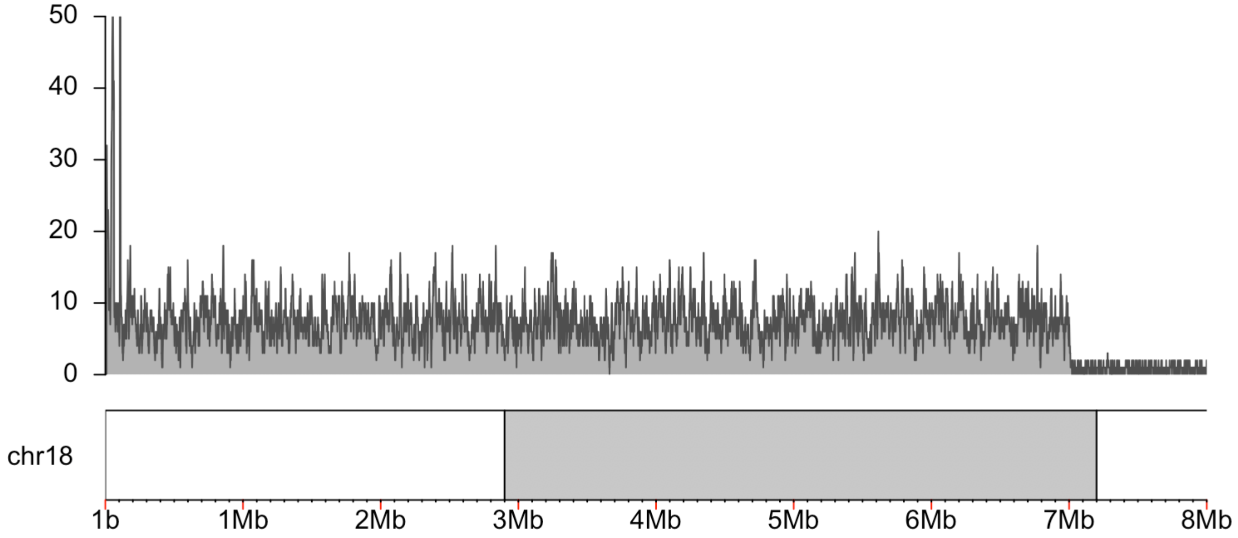
50
40
30
20
10
0
chr18
1b
1Mb
2Mb
3Mb
4Mb
5Mb
6Mb
7Mb
8Mb

I:

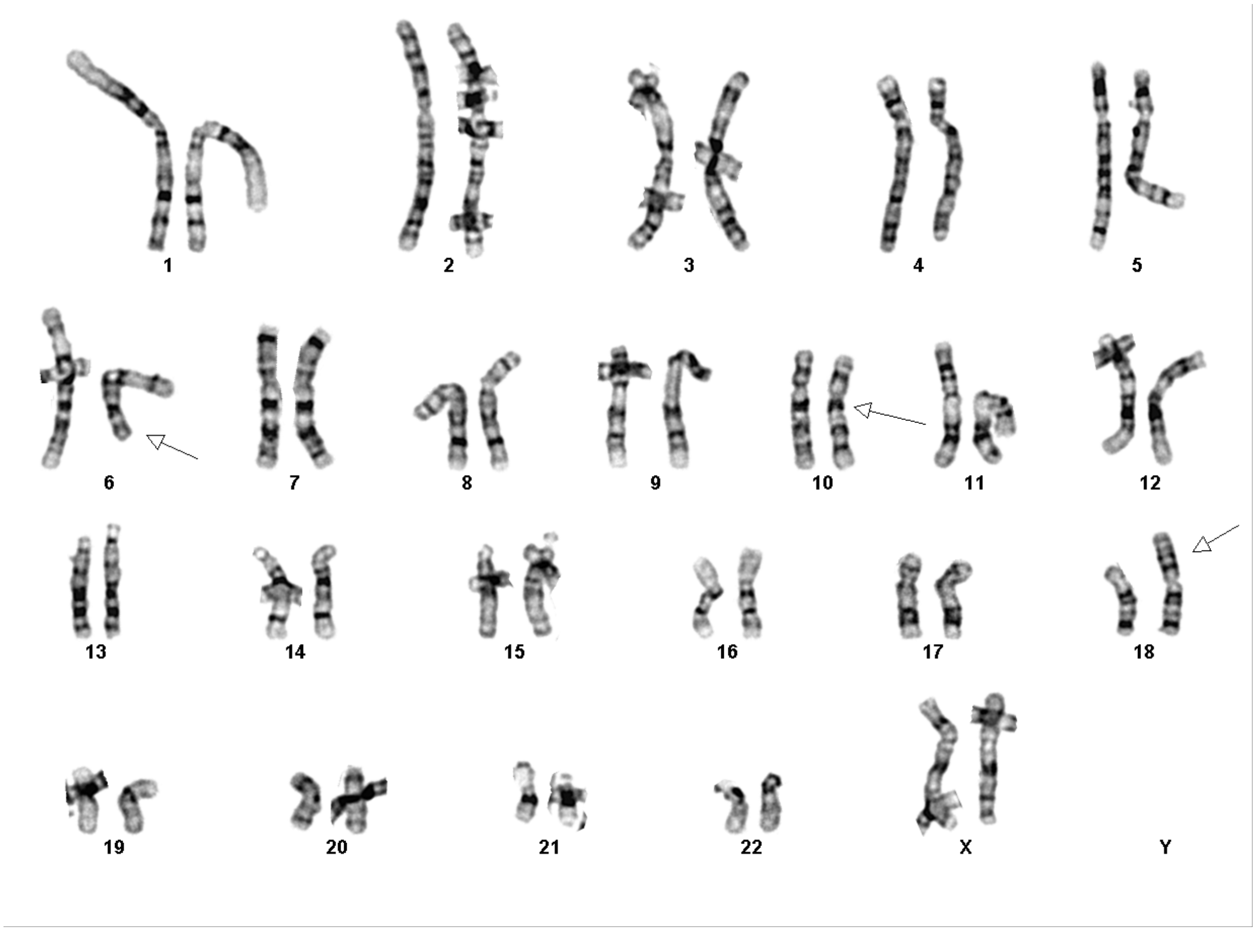
X
Y

**J:** 3' end of 10:A bisects *KIAA1217* (IGV: chr10:24,286,090-24,286,130) and is linked to the 5' end of 10:D which bisects *GPR158* (IGV: chr10:25,278,377-25,278,417). This is the beginning of derivative chromosome 10.

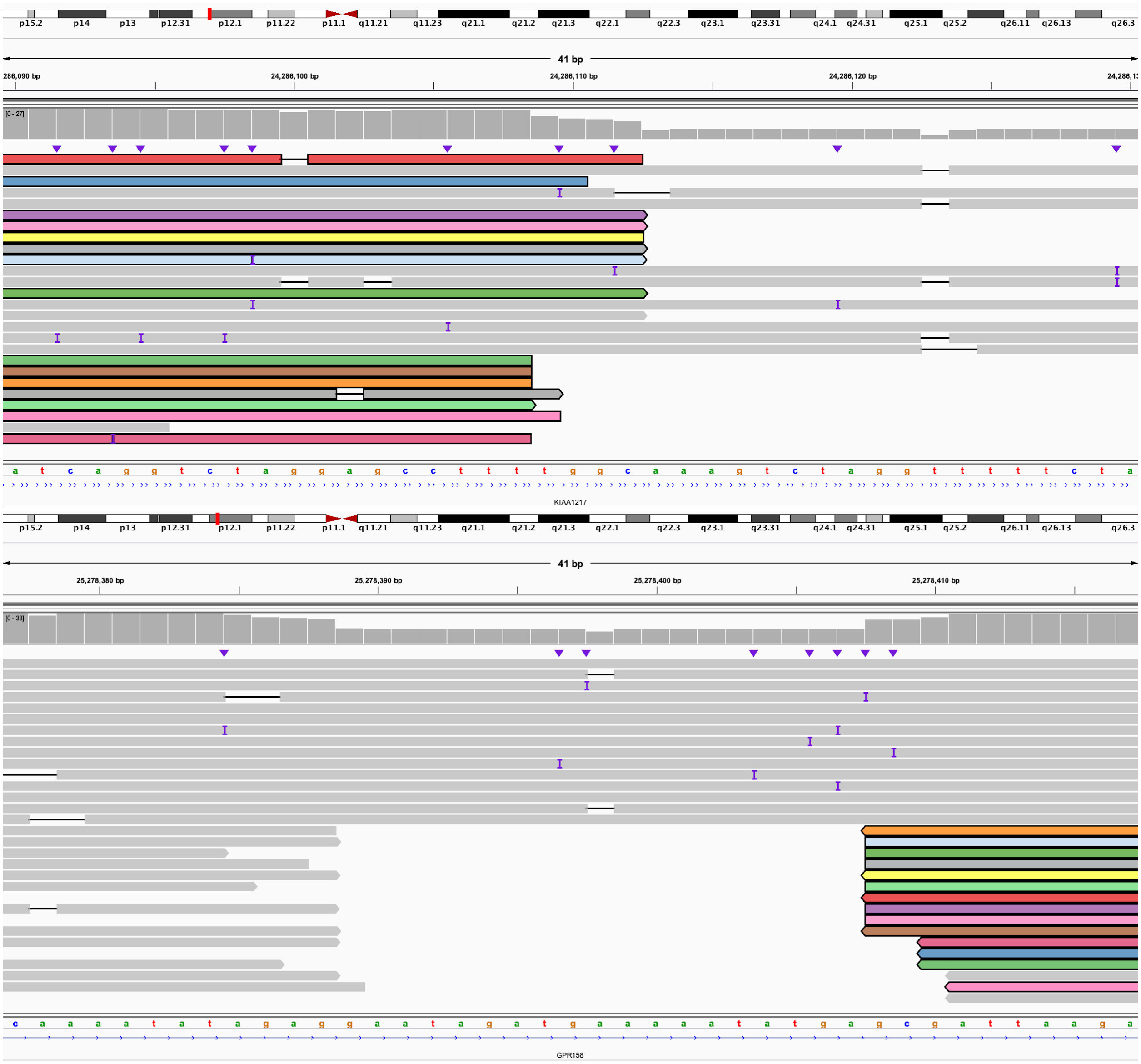

**K:** 3' end of 10:D bisects *PARD3* (IGV: chr10:34,622,779-34,622,819) and is linked to the 3' end of 10:F (IGV: chr10:49,775,879-49,775,919). This is within derivative chromosome 10.

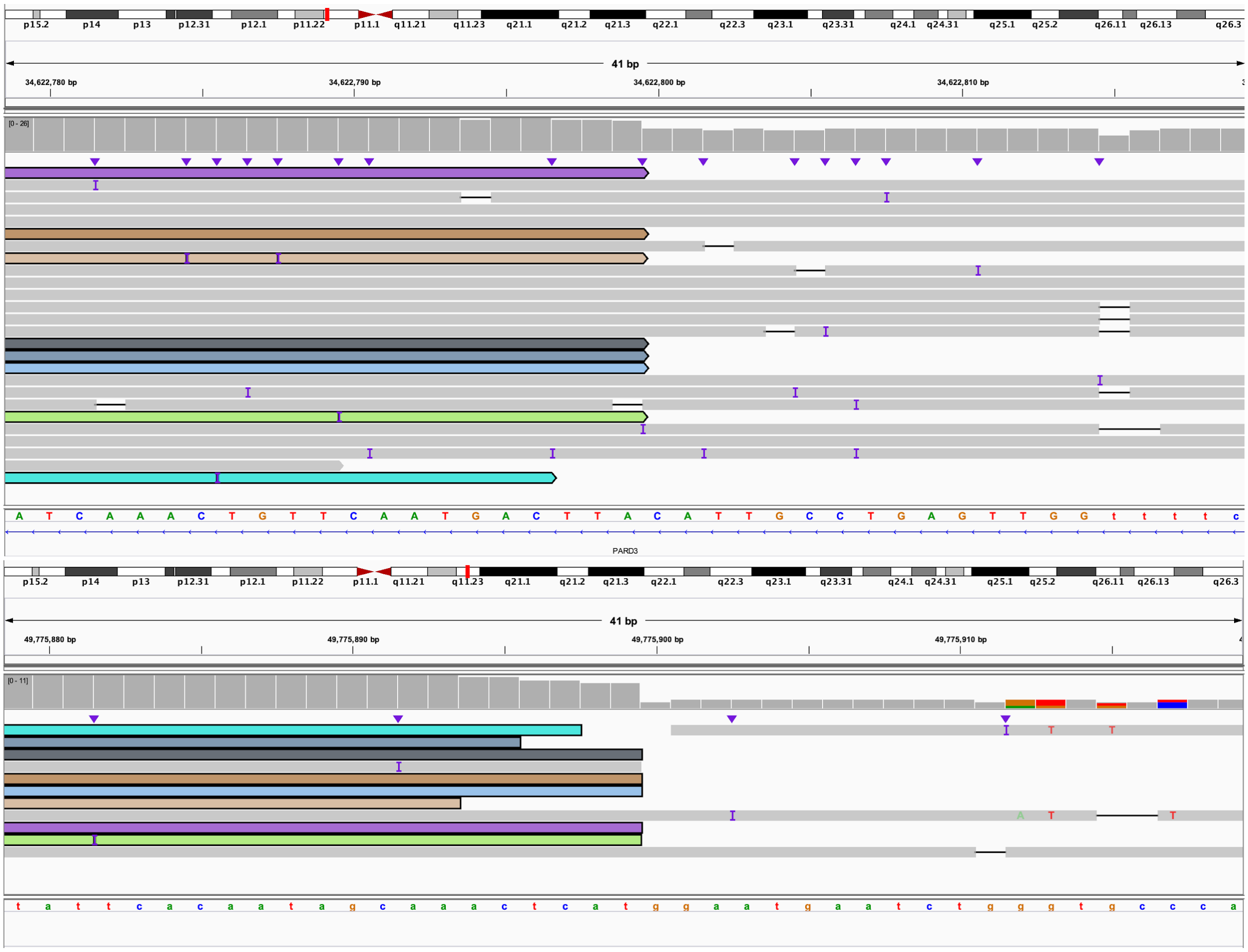

**L:** 5' end of 10:F (IGV: chr10:35,972,571-35,972,611) is linked to the 3' end of 5:C (IGV: chr5:28,826,569-28,826,609). This is within derivative chromosome 10.

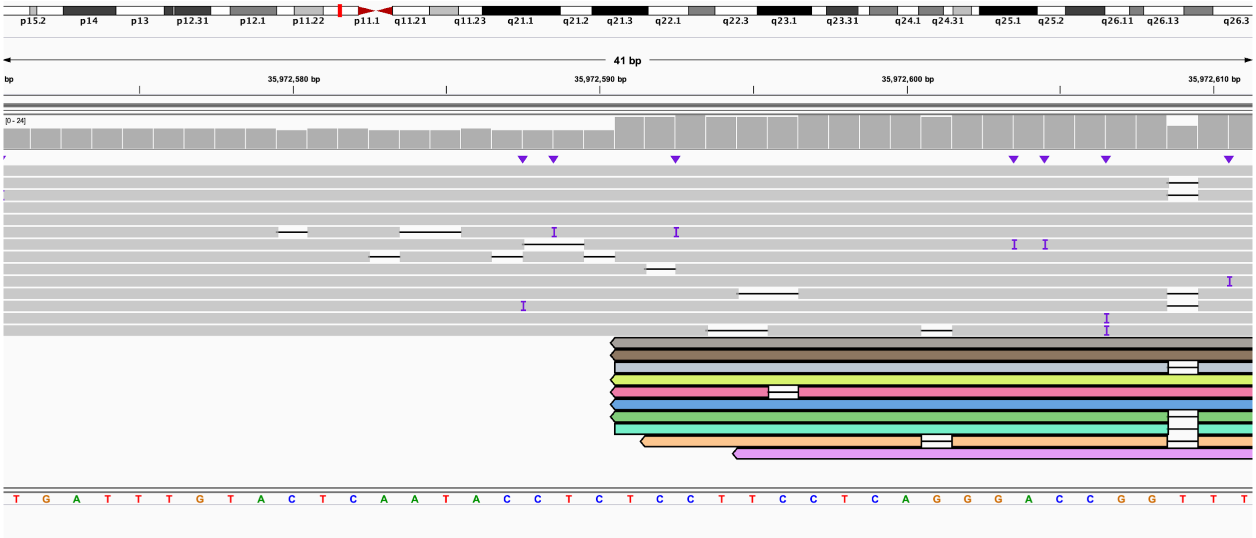

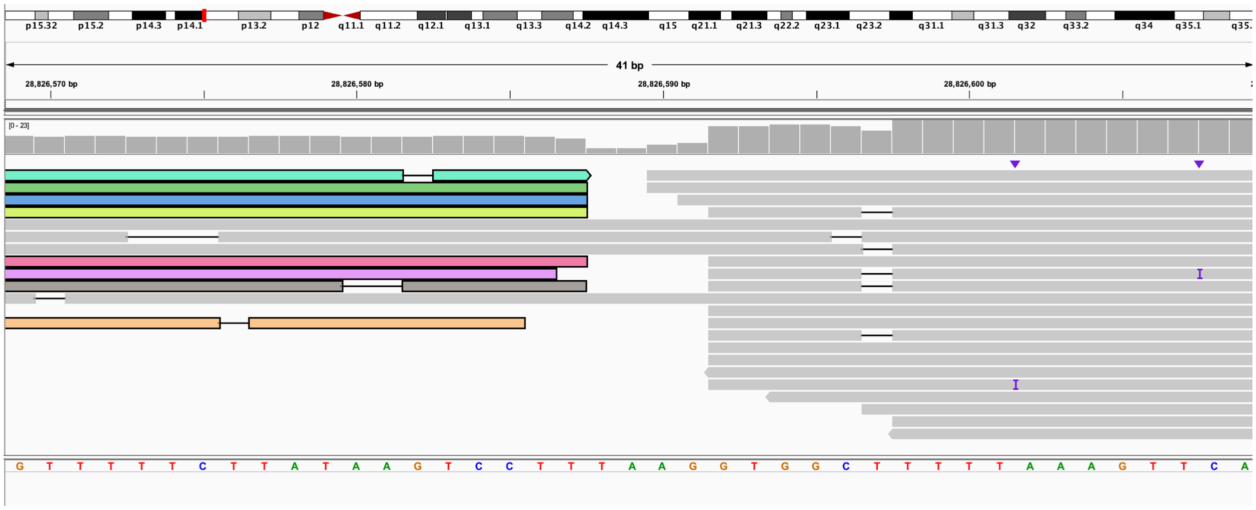

**M:** 5' end of 5:C (IGV: chr5:28,650,520-28,650,560) is linked to the 5' end of 10:K (IGV: chr10:56,736,263-56,736,303). This is the end of derivative chromosome 10.

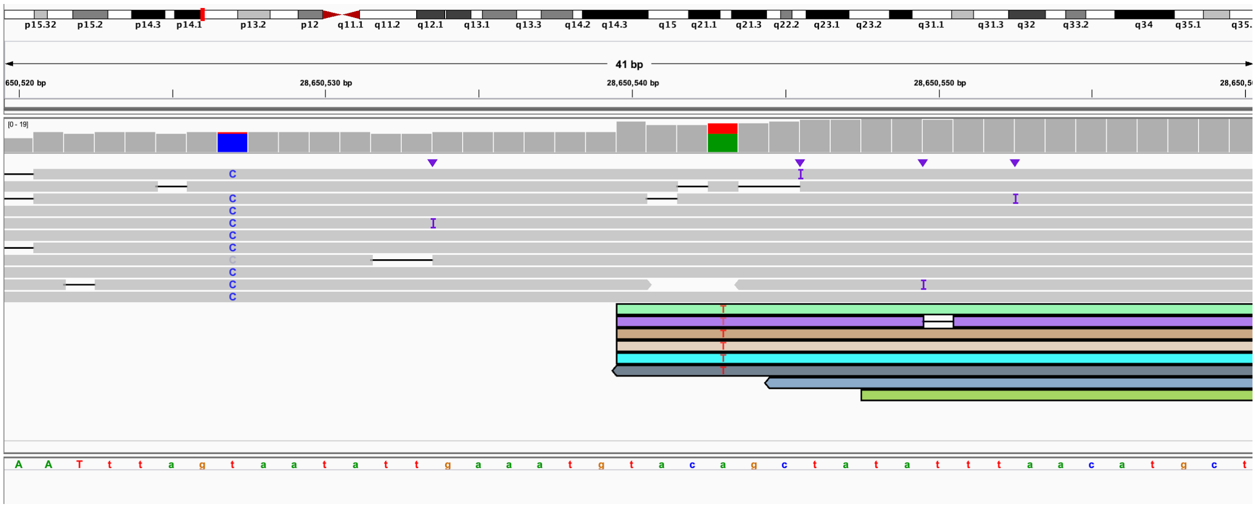

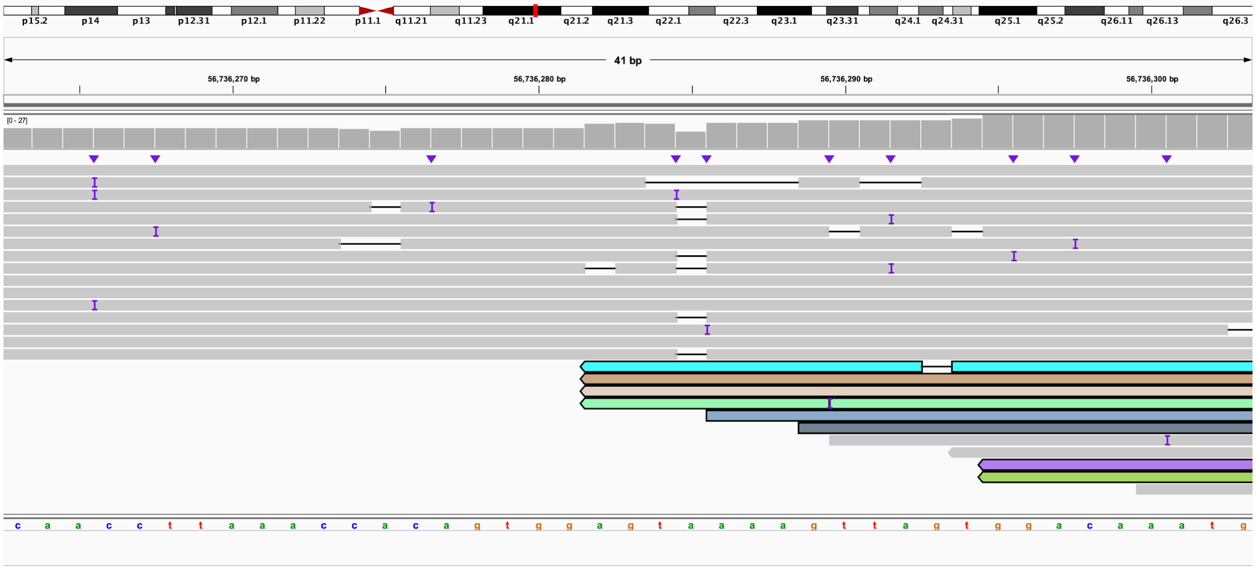

**N:** 3' end of 5:A (IGV: chr5:22,881,832-22,881,872) linked to 3' end of 10:I (IGV: chr10:55,632,859-55,632,898). This is the beginning of derivative chromosome 5.

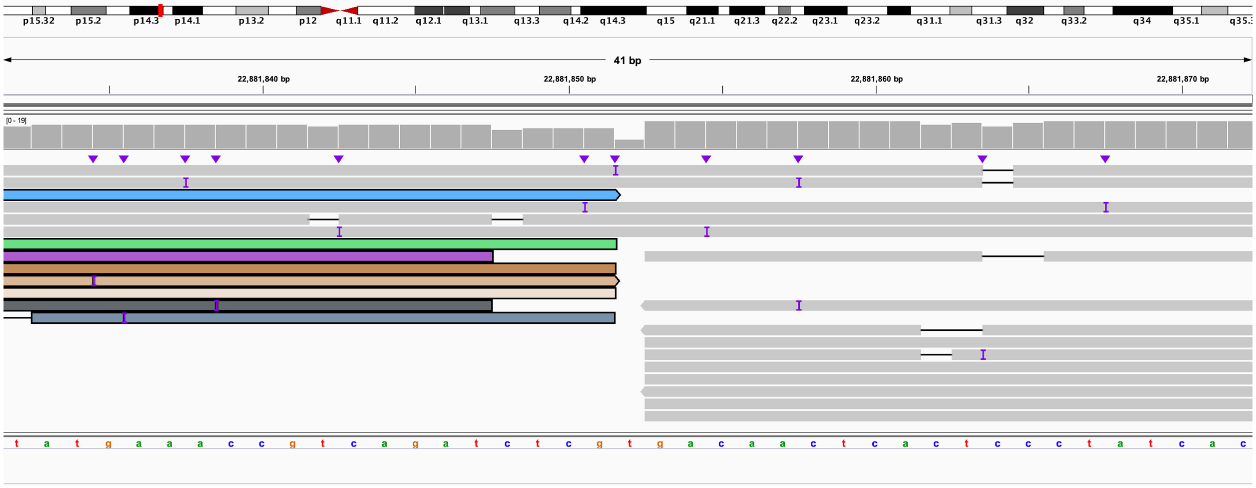

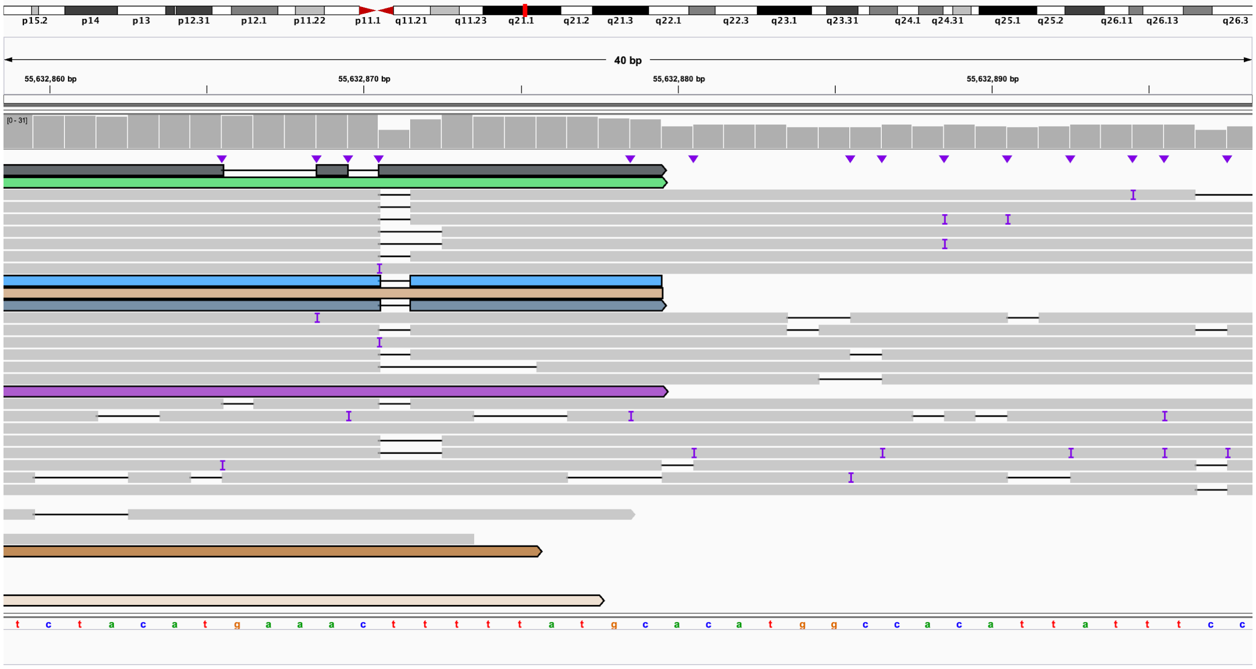

**O:** 5' end of 10:I bisects *PCDH15* (IGV: chr10:55,305,294-55,305,334) linked to 3' end of 10:G which also bisects *PCDH15* (IGV: chr10:54,672,847-54,672,886). This is within derivative chromosome 5.

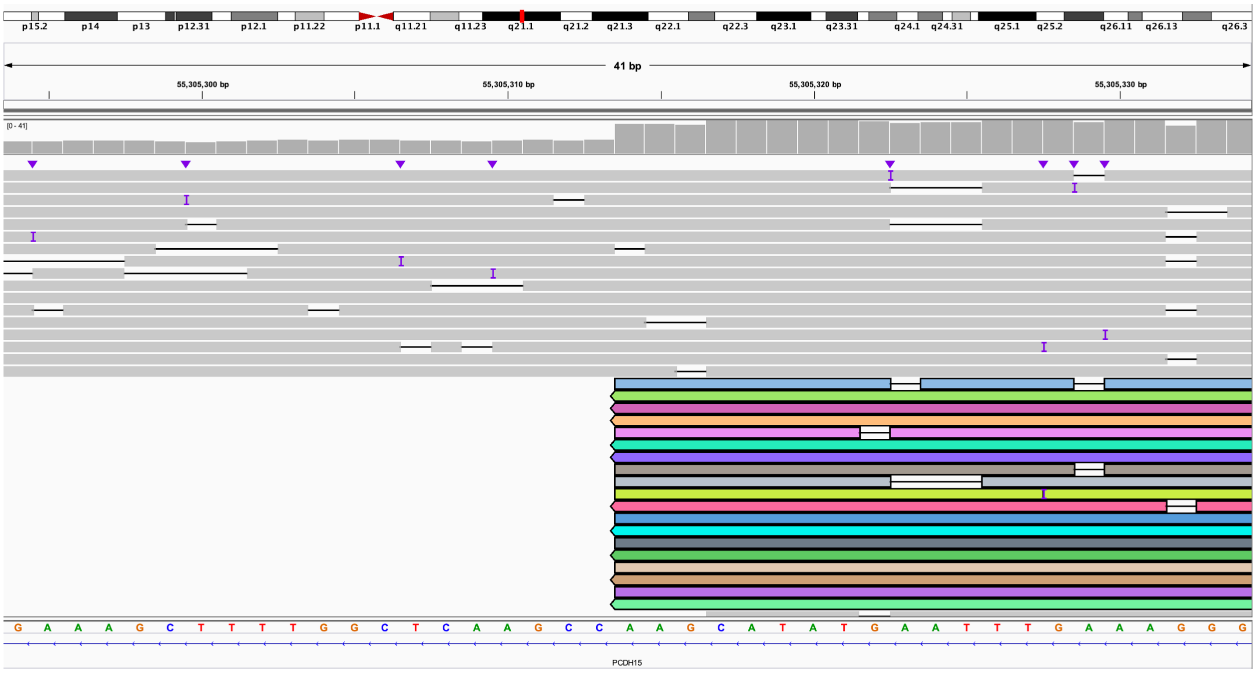

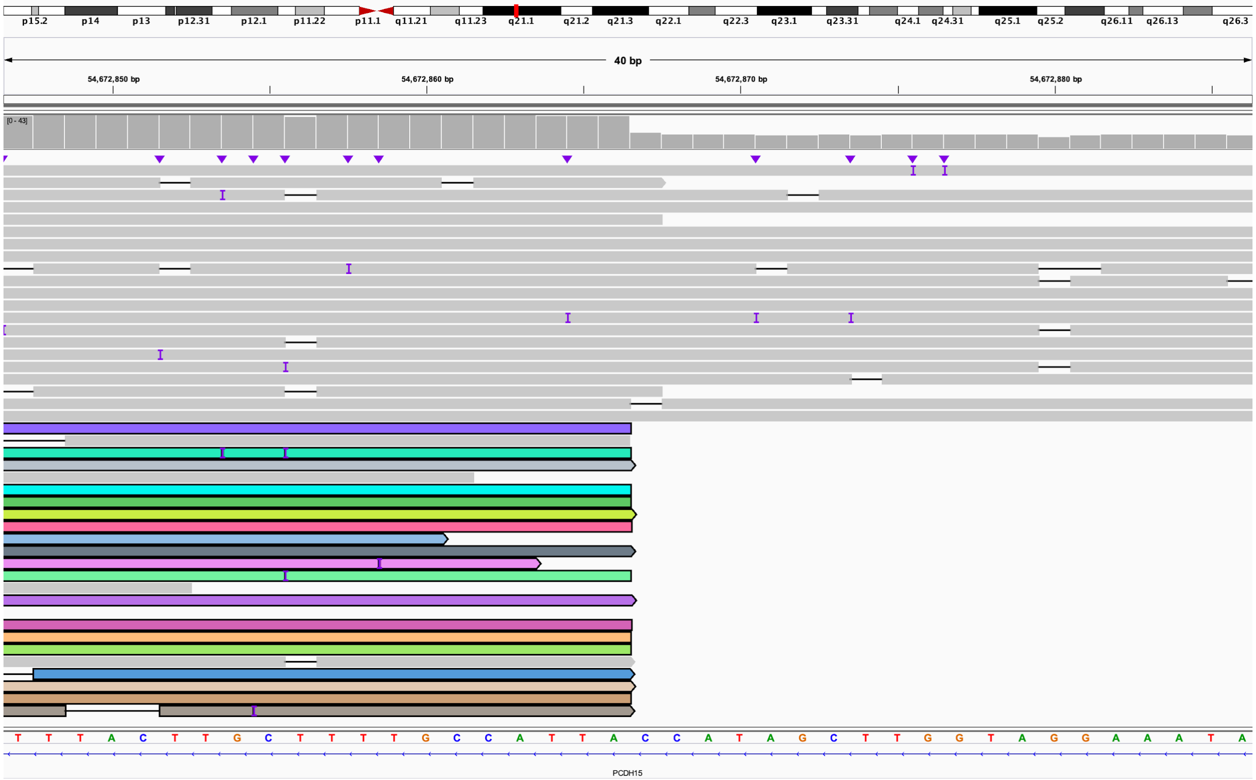

**P:** 5' end of 10:G (IGV: chr10:49,775,880-49,775,920) linked to 3' end of 5:B (IGV: chr5:28,650,519-28,650,558). This is within derivative chromosome 5.

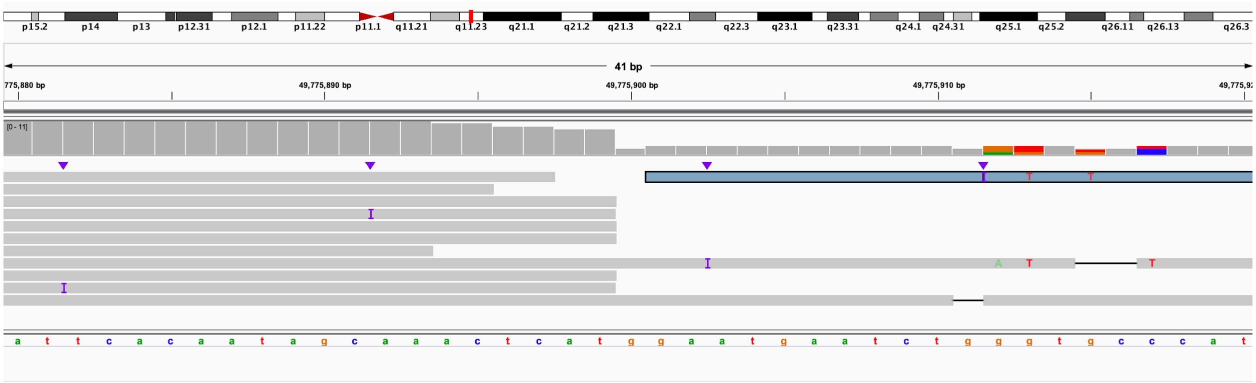

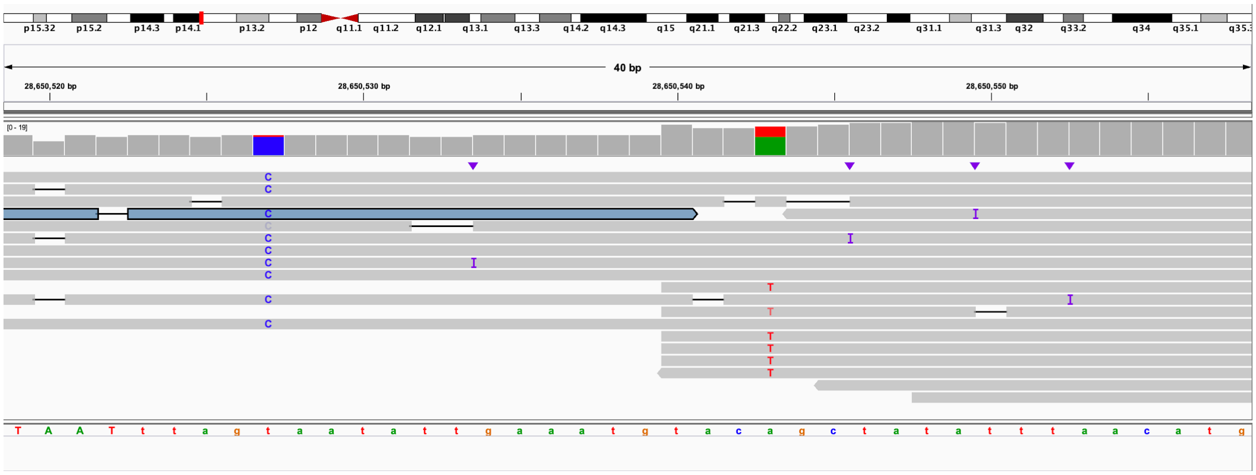

**Q:** 5' end of 5:B (IGV: chr5:22,881,832-22,881,872) linked to 5' end of 10:C (IGV: chr10:24,725,830-24,725,870). This is within derivative chromosome 5.

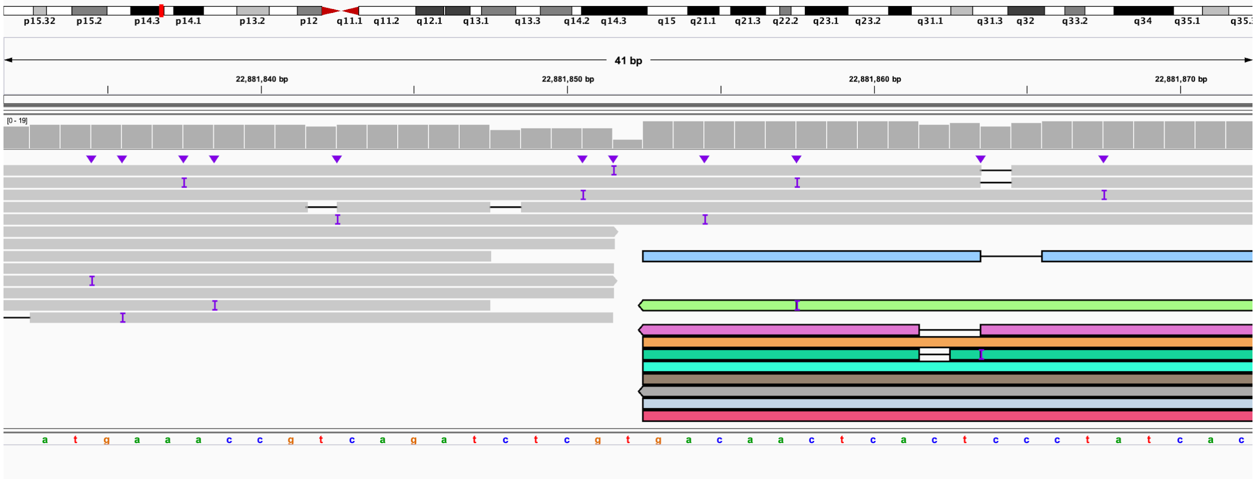

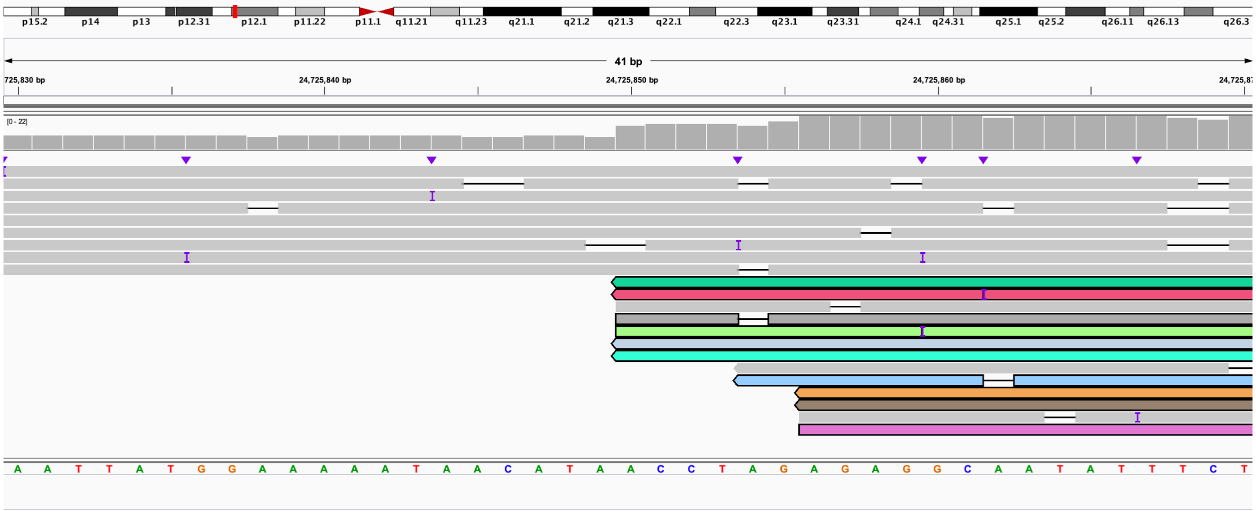

**R:** 3' end of 10:C is within *GPR158* (IGV: chr10:25,278,367-25,278,407) and is linked to 5' end of 5:D (IGV: chr5:28,826,570-28,826,609). This is the end of derivative chromosome 5.

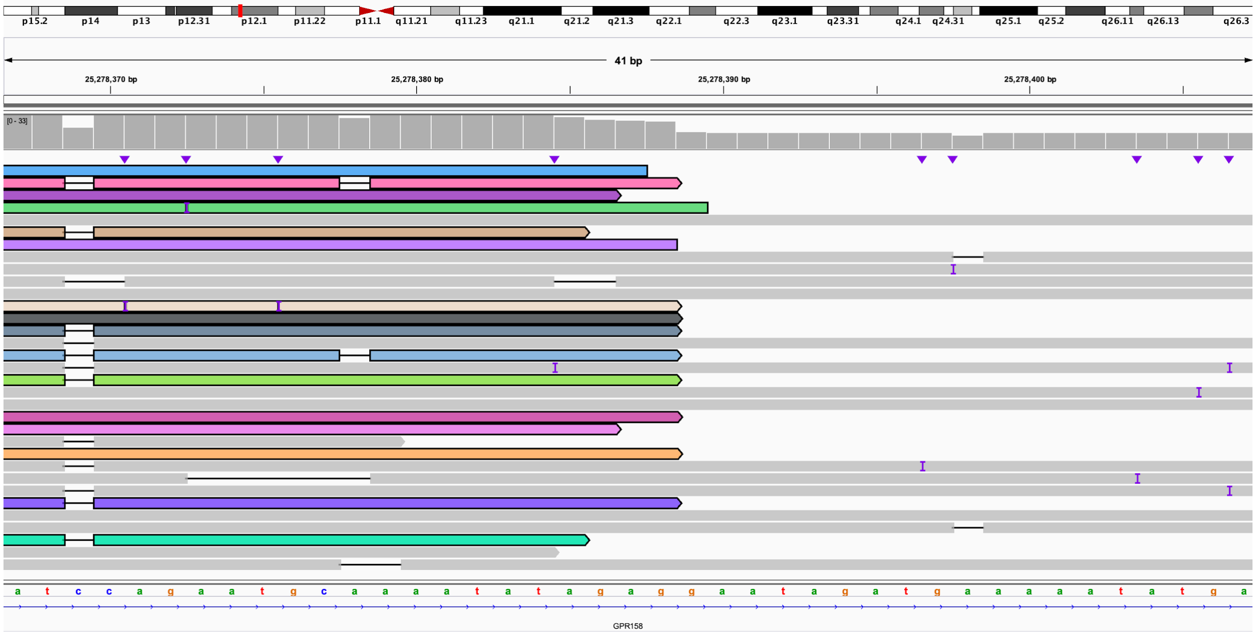

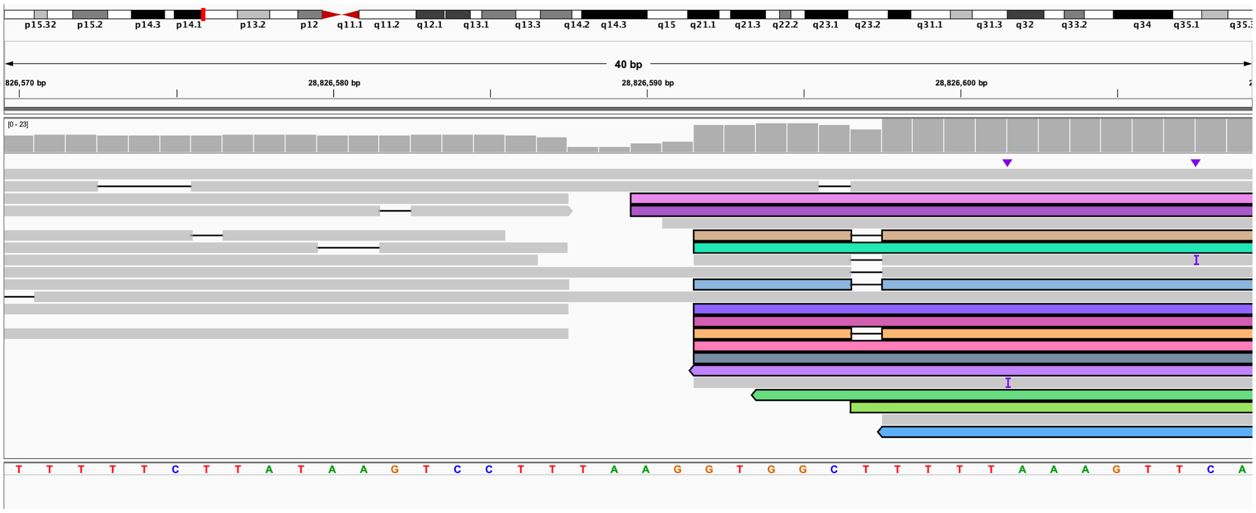

**S:** 3' end of 6:A bisects *CLVS2* (IGV: chr6:123,005,528-123,005,567) linked to 5' end of 18:C which bisects *L3MBTL4* (IGV: chr18:6,280,036-6,280,076). This is the beginning of derivative chromosome 6.

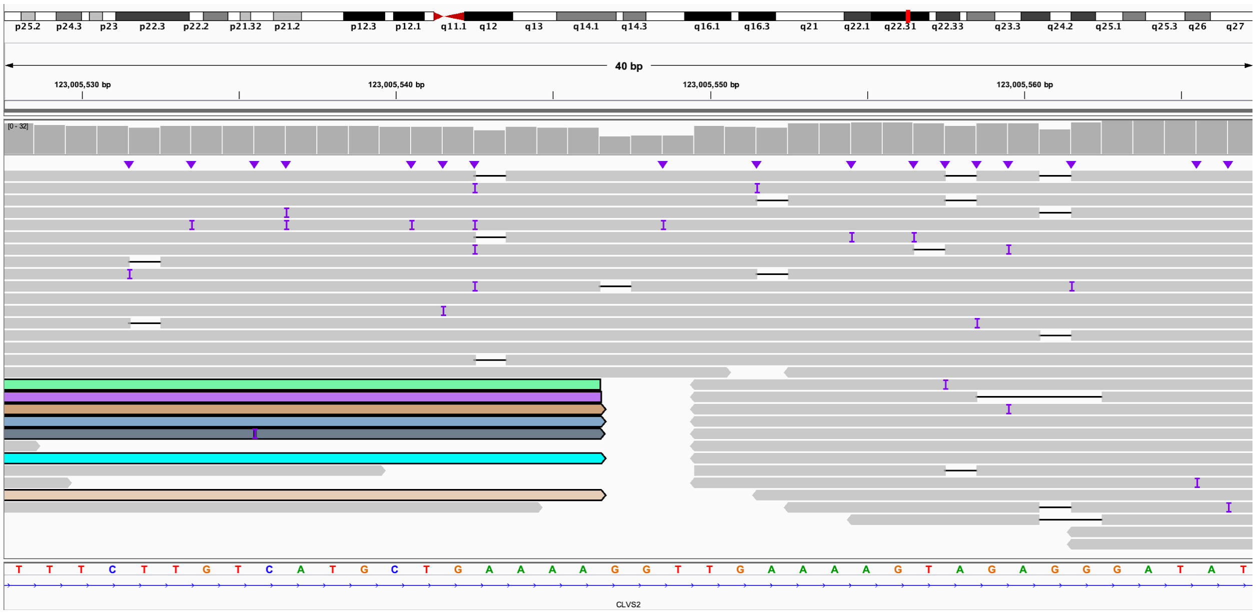

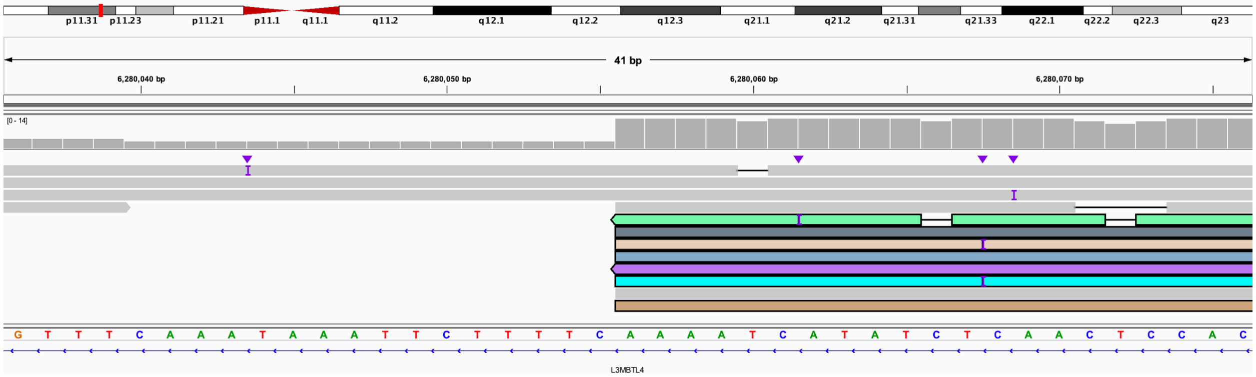

**T:** 3' end of 18:C (IGV: chr18:7,279,949-7,279,989) linked to 3' end of 18:A which bisects *L3MBTL4* (IGV: chr18:6,275,113-6,275,152). This is the end of derivative chromosome 6.

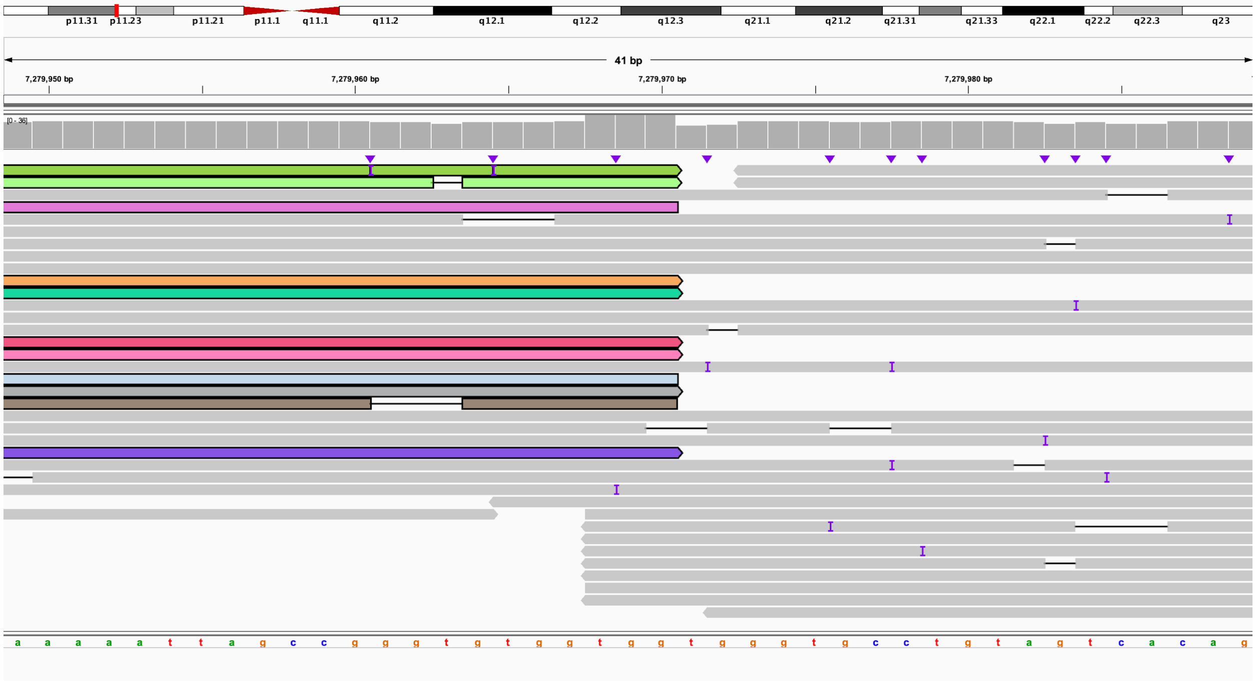

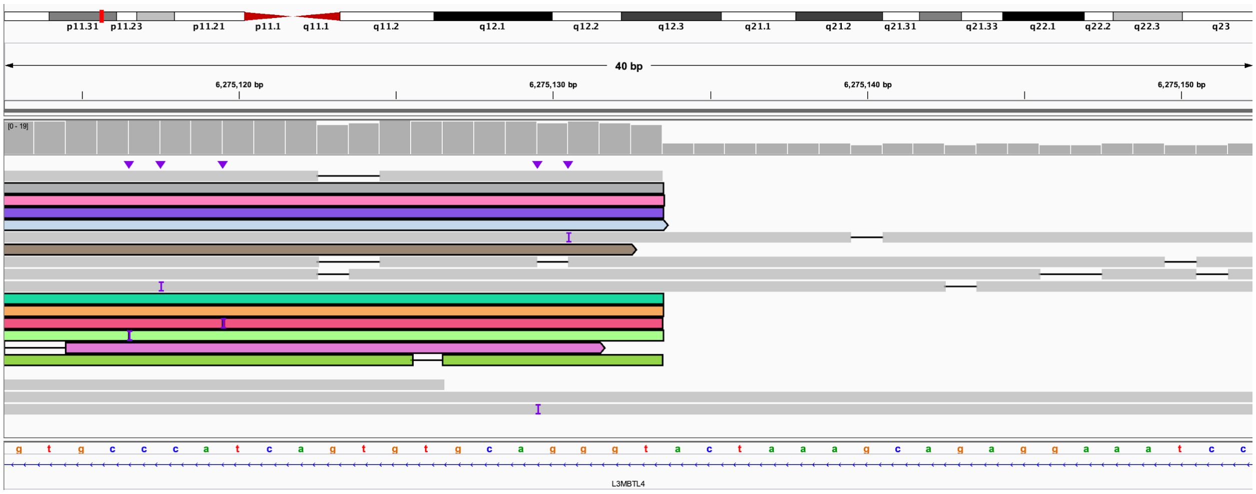

**U:** 5' end of 18:D (IGV: chr18:7,279,950-7,279,989) linked to 5' end of 6:C which bisects *NKAIN2* (IGV: chr6:124,505,710-124,505,749). This is the beginning of derivative chromosome 18.

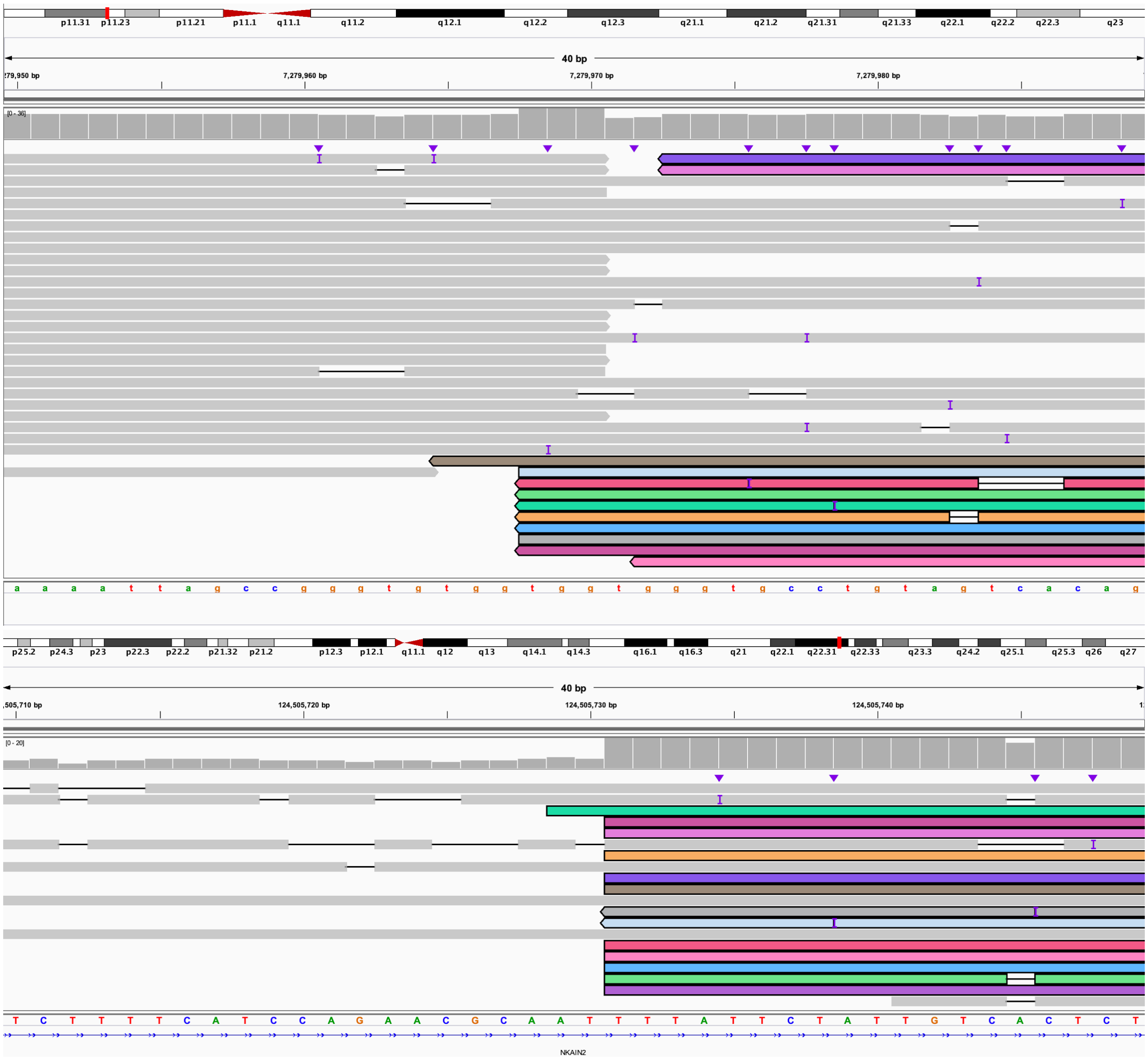

**V:** 3' end of 6:C bisects *XR_001743835.1* (IGV: chr6:126,657,900-126,657,940) linked to 5' end of 6:B which bisects *CLVS2* (IGV: chr6:123,005,528-123,005,568). This is within derivative chromosome 18.

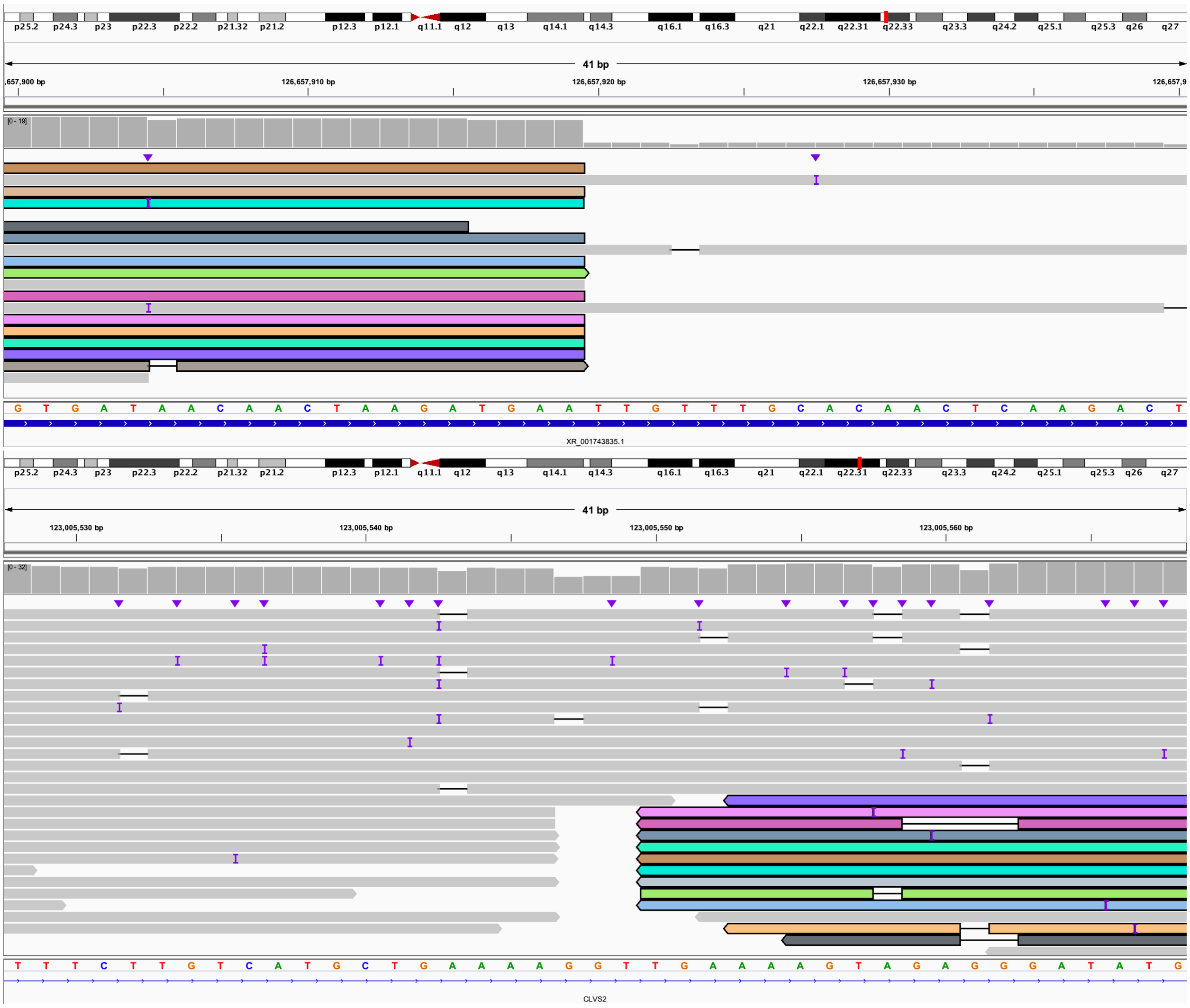

**W:** 3' end of 6:B bisects *NKAIN2* (IGV: chr6:124,505,249-124,505,393) linked to 5' end of 6:E which bisects *XR_001743835.1* (IGV: chr6:126,680,021-126,680,060). This is the end of derivative chromosome 18.

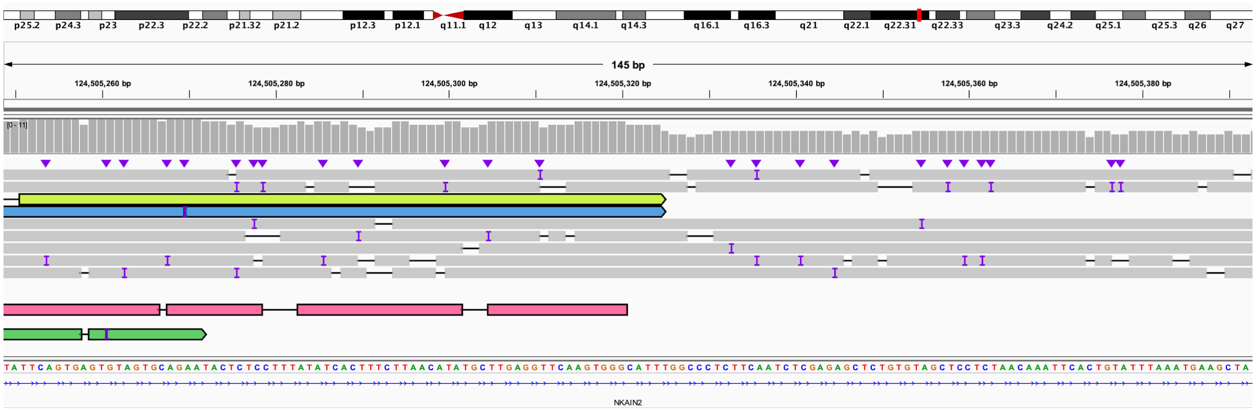

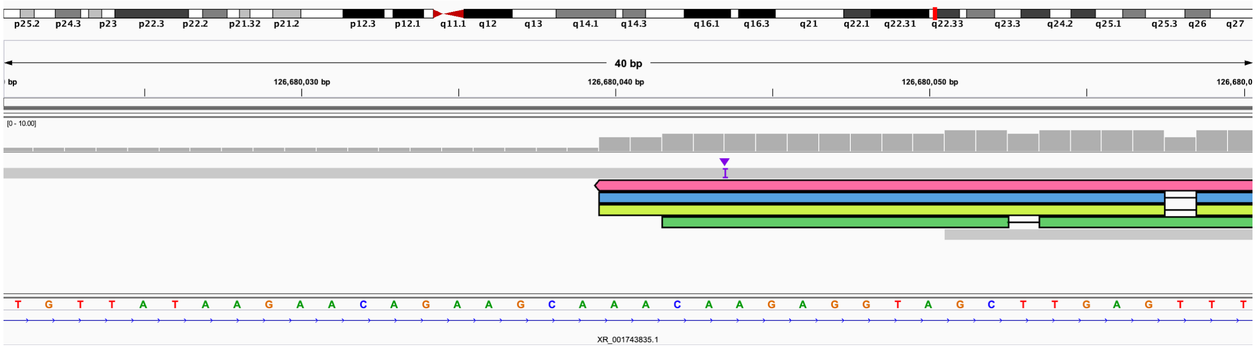

**Figure S26. S002 (*ALMS1*), IGV views of known inherited stop variant and *Alu* insertion.**

**A.** IGV view of all reads showing the known paternally inherited C>G that results in a stop codon. IGV coordinates for the screenshot are chr2:73,448,739-73,448,779. **B.** Screenshot of ~300 bp insertion in exon 20 that represents an *Alu* insertion, all reads are shown. IGV view is chr2:73,602,213-73,602,266.

A:

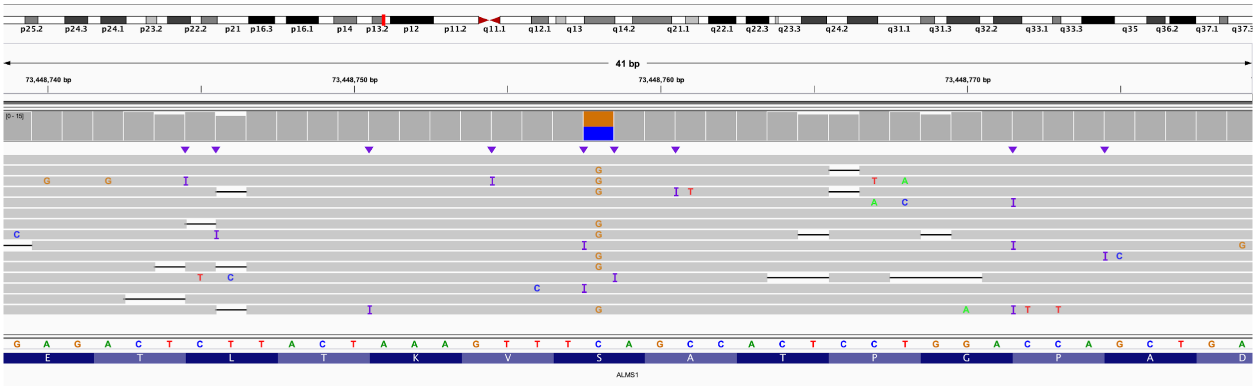

B:

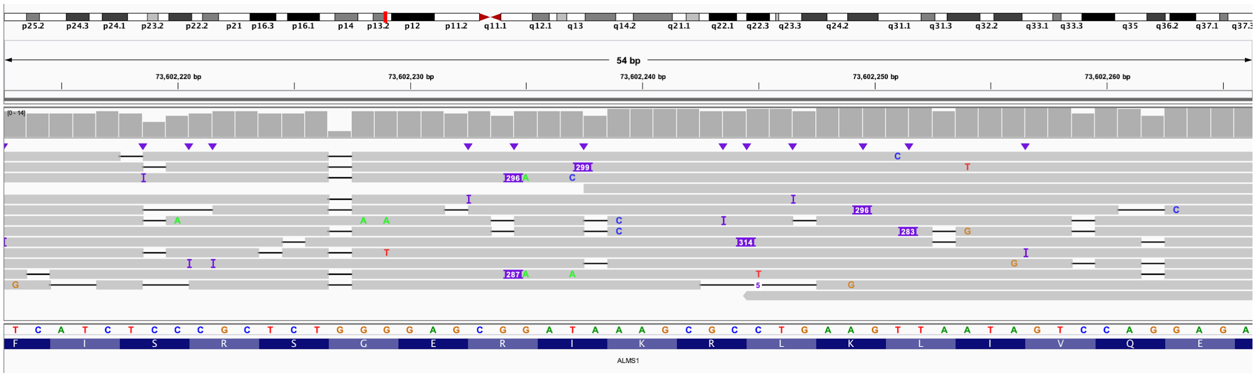

### Figure S27. S003 (*NPHP4*), IGV views of inherited stop, splice variant, and data showing variant does affect splicing.

**A.** IGV view of all reads (top track) and reads phased into two haplotypes (middle and bottom tracks). DNA sequence and gene body are located below. The known paternally inherited G>A is located in the bottom haplotype. IGV screenshot is from chr1:5,986,136-5,986,175. **B.** IGV view with reads as in A showing the G>C predicted to create a novel splice donor site on a different haplotype than the known paternally inherited G>A in A. IGV screenshot is from chr1:5,967,229-5,967,269. **C.** Analysis of SVs using both SVIM and Sniffles identified two insertions within an AG-rich repeat in an intron of *NPHP4* at approximately chr1:5,977,890. Phasing of reads separated all reads into a haplotype containing the ~800 bp insert and another containing the ~1,340 bp insert, similar to sizes observed in both nonhuman primates and human samples (Sulovari et al. 2019). IGV screenshot is from chr1:5,977,792-5,978,001. **D.** Splicing defect caused by deep intronic *NPHP4* variant in individual S004. **i.** *NPHP4* exon structure around the c.517+50C>G variant predicted to generate a new splice donor site (gtgag). **ii.** Normal splice isoform indicating primer pair 1 spanning the Exon 5-6 junction and corresponding Sanger sequencing of the PCR product. **iii.** Predicted aberrant transcript with inclusion of 49 intronic base pairs indicating predicted PCR product size for primer pair 1. **iv.** Predicted aberrant transcript indicating predicted PCR product size for primer pair 2, which should only amplify the aberrant transcript and corresponding Sanger sequencing of the PCR product. **v.** PCR products from S004 and unaffected fibroblast cDNA. Bands match the sizes predicted in B-D. NPHP4 reference sequence: NM_015102.4. bp=base pairs; NTC=No Template Control PCR; WT=wild-type, unaffected fibroblast cDNA.

A:

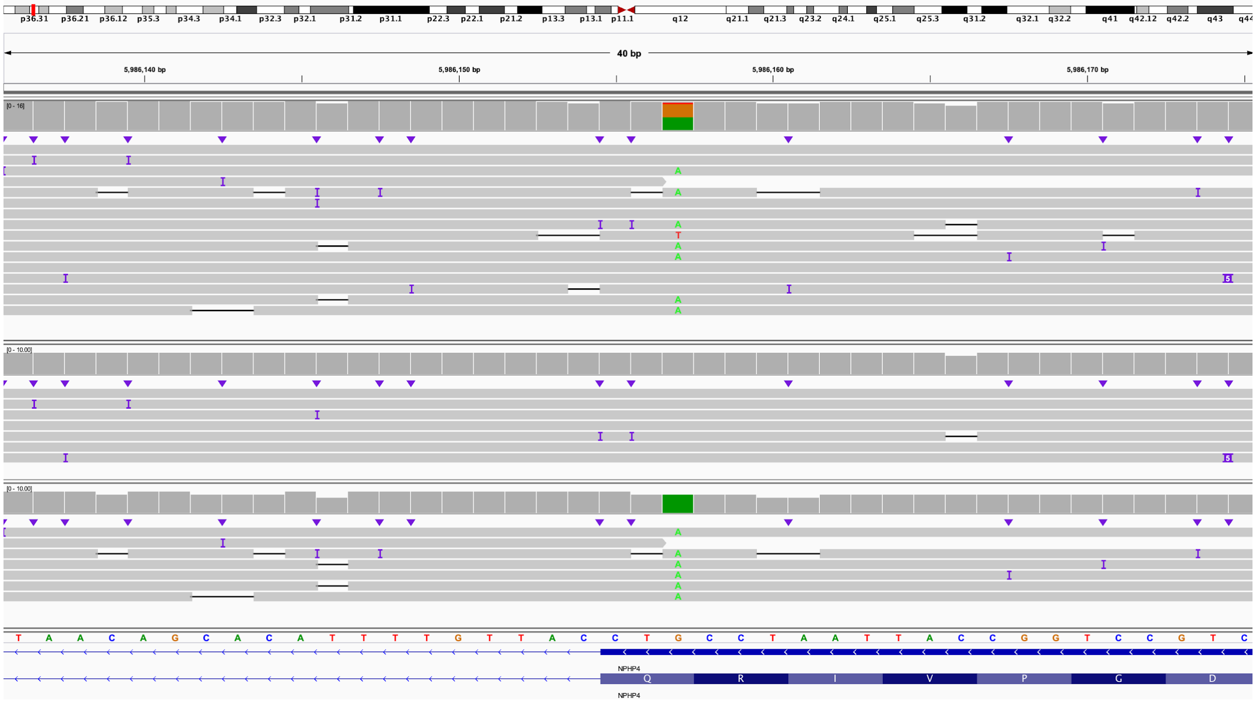

B:

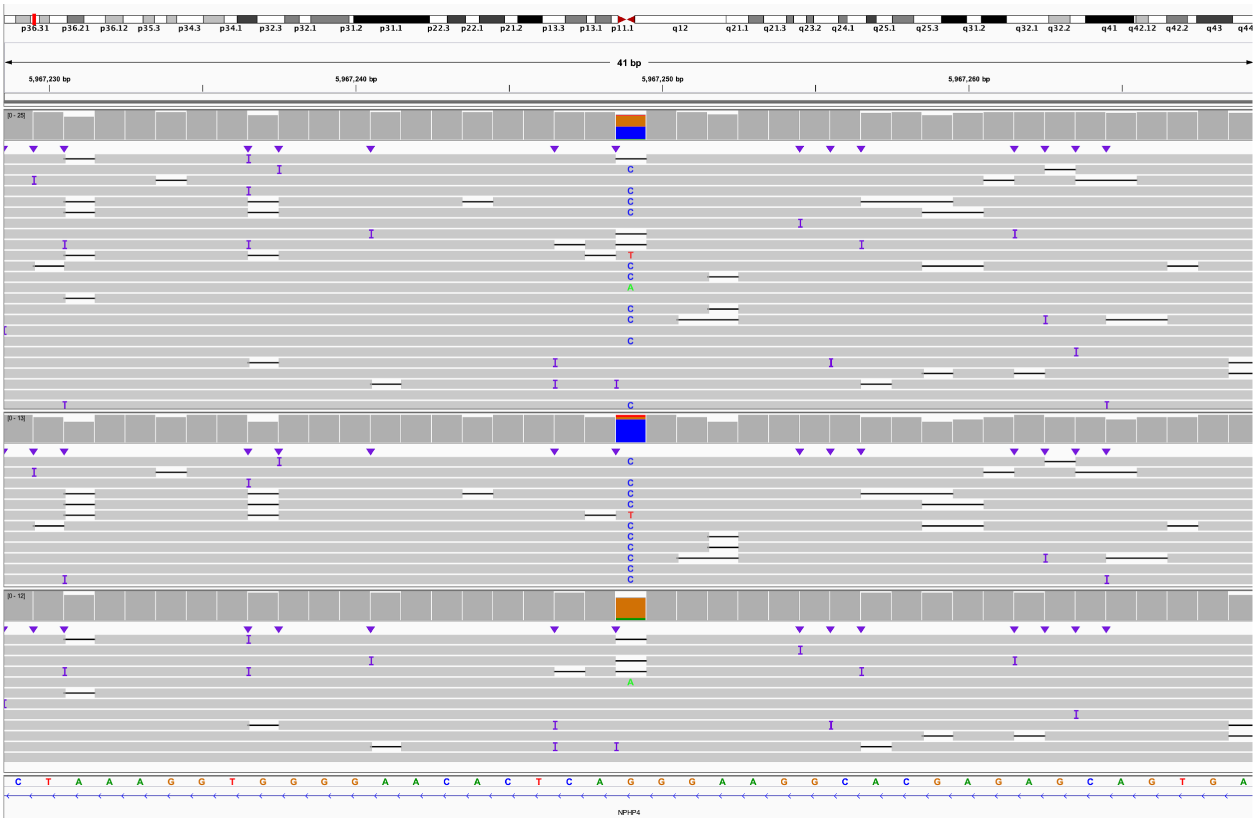
p36.31 p36.21 p36.12 p35.3 p34.3 p34.1 p32.3 p32.1 p31.2 p31.1 p22.3 p22.1 p21.2 p13.3 p13.1 p11.1 q12 q21.1 q21.3 q23.2 q24.1 q25.1 q25.3 q31.2 q32.1 q32.2 q41 q42.12 q42.2 q43 q44
41 bp
5,967,230 bp
5,967,240 bp
5,967,250 bp
5,967,260 bp
[0 - 25]
[0 - 13]
[0 - 12]
C T A A A G G T G G G G A A C A C T C A G G G A A G G C A C G A G A G C A G T G A
NPHP4

C:

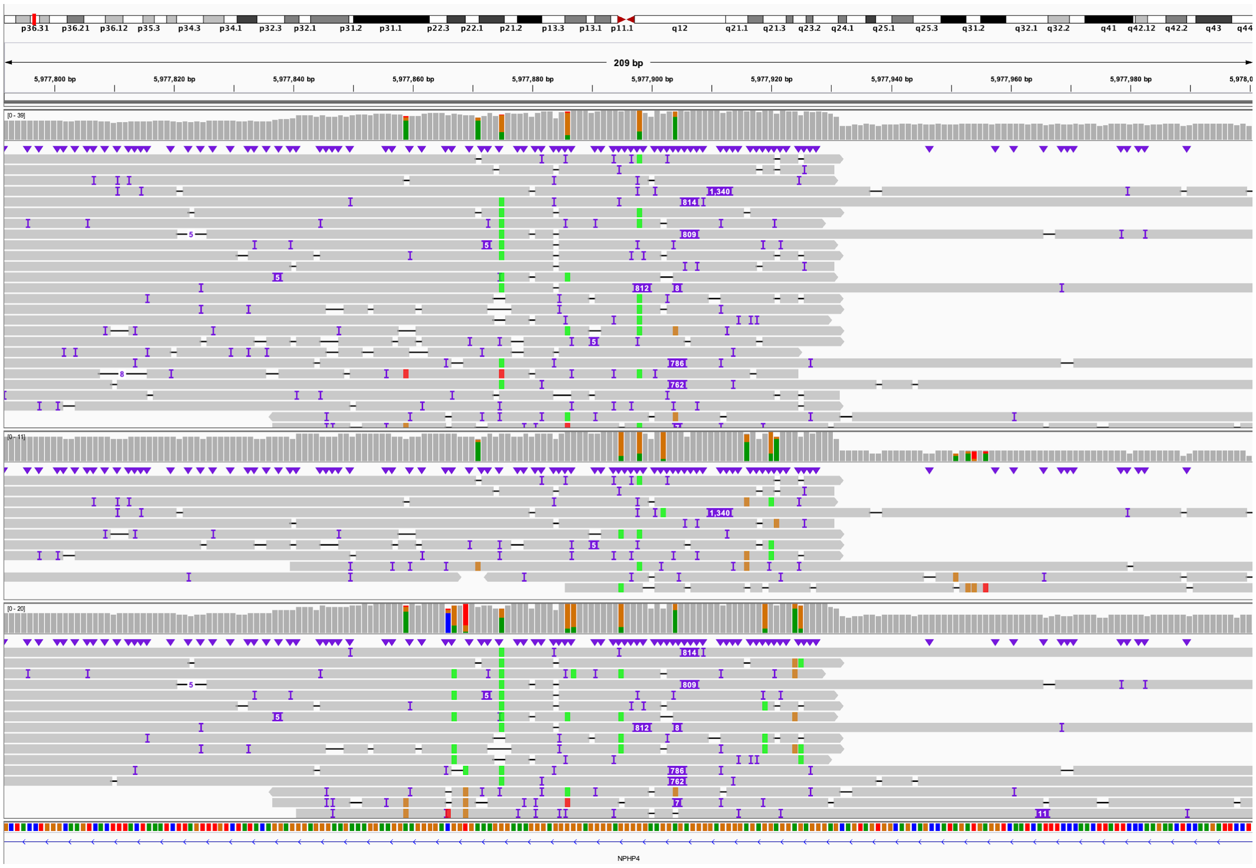
p36.31
p36.21
p36.12
p35.3
p34.3
p34.1
p32.3
p32.1
p31.2
p31.1
p22.3
p22.1
p21.2
p13.3
p13.1
p11.1
q12
q21.1
q21.3
q23.2
q24.1
q25.1
q25.3
q31.2
q32.1
q32.2
q41
q42.12
q42.2
q43
q44
209 bp
5,977,800 bp
5,977,820 bp
5,977,840 bp
5,977,860 bp
5,977,880 bp
5,977,900 bp
5,977,920 bp
5,977,940 bp
5,977,960 bp
5,977,980 bp
NPHP4

D:

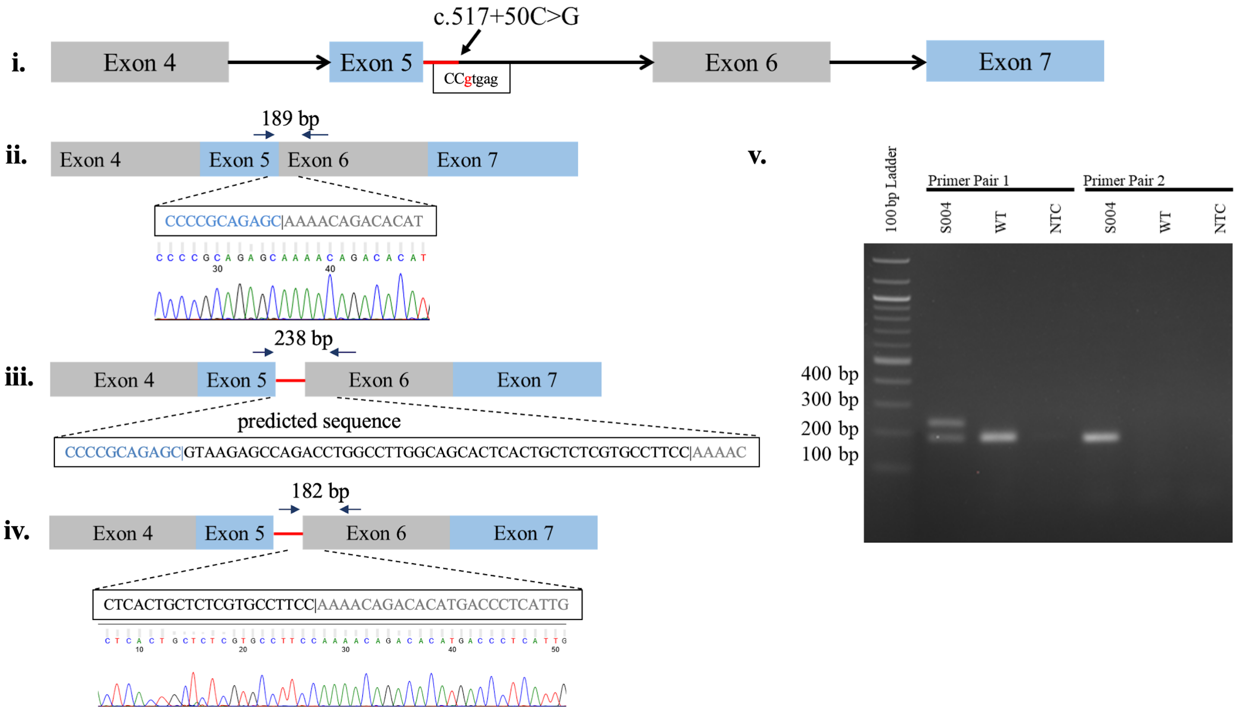

c.517+50C>G
i.
Exon 4
Exon 5
CCgtgag
Exon 6
Exon 7
189 bp
ii.
Exon 4
Exon 5
Exon 6
Exon 7
CCCCGCAGAGC|AAAACAGACACAT
238 bp
iii.
Exon 4
Exon 5
Exon 6
Exon 7
predicted sequence
CCCCGCAGAGC|GTAAGAGCCAGACCTGGCCTTGGCAGCACTCACTGCTCTCGTGCCTTCC|AAAAC
182 bp
iv.
Exon 4
Exon 5
Exon 6
Exon 7
CTCACTGCTCTCGTGCCTTCC|AAAACAGACACATGACCCTCATTG
v.
100 bp Ladder
Primer Pair 1
Primer Pair 2
S004
WT
NTC
S004
WT
NTC
400 bp
300 bp
200 bp
100 bp

**Figure S28. S004 (*VARS2*), IGV view of known inherited variant.**

**A.** IGV view of known pathogenic paternally inherited G>A. No second variant was found in this case. IGV view is of chr6:30,920,072-30,920,111. **B.** Analysis of reads reveals no change in methylation at the 5' UTR of *VARS2*. In this view, blue represents a hypomethylated region, suggesting an open 5' region and promoter. The green box under the promoter indicates the position of the CpG island. IGV view is chr6:30,912,208-30,928,459.

A:

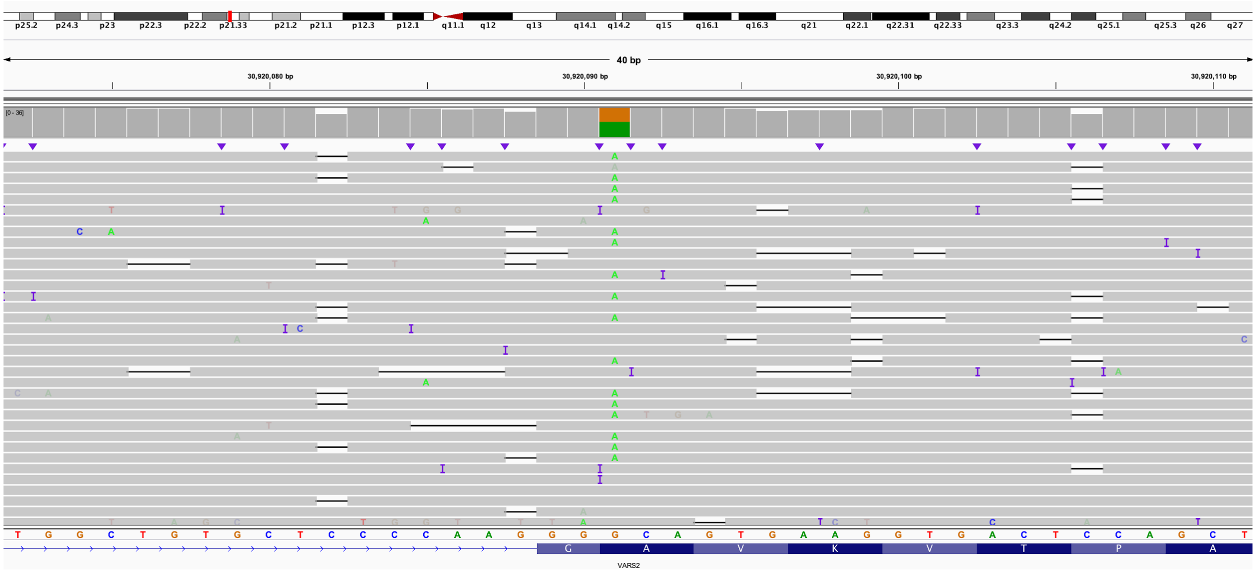

B:

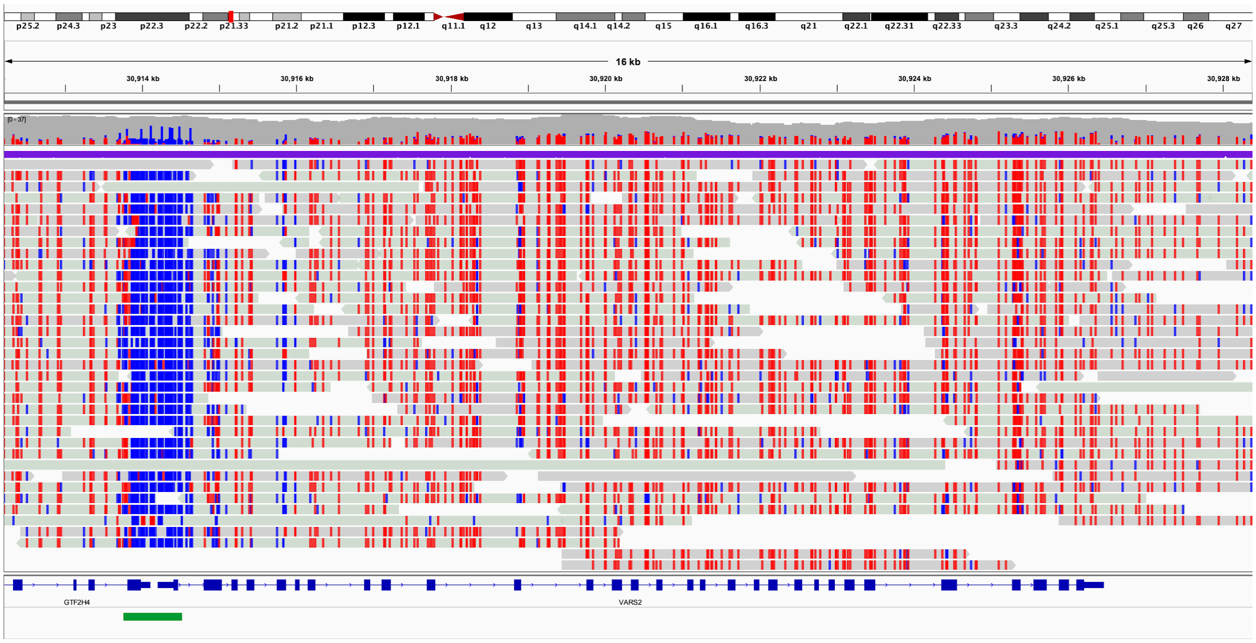

**Figure S29. S008 (*HPRT1*), view of inversion and FISH results.**

**A.** T-LRS suggested an approximately 17 Mbp inversion bisected *HPRT1*. Reads on the left partially mapped to approximately chrX:117,359,013 and reads on the right partially mapped to chrX:117,488,245. IGV view is chrX:134,482,505-134,484,004. **B.** FISH probes were designed to confirm the presence of an inversion. **C.** Image from control case, dashed circle highlights X chromosome. **D.** Image from patient confirmed the presence of a 17 Mbp inversion reported as inversion (X)(q24q26.3), dashed circle highlights the X chromosome.

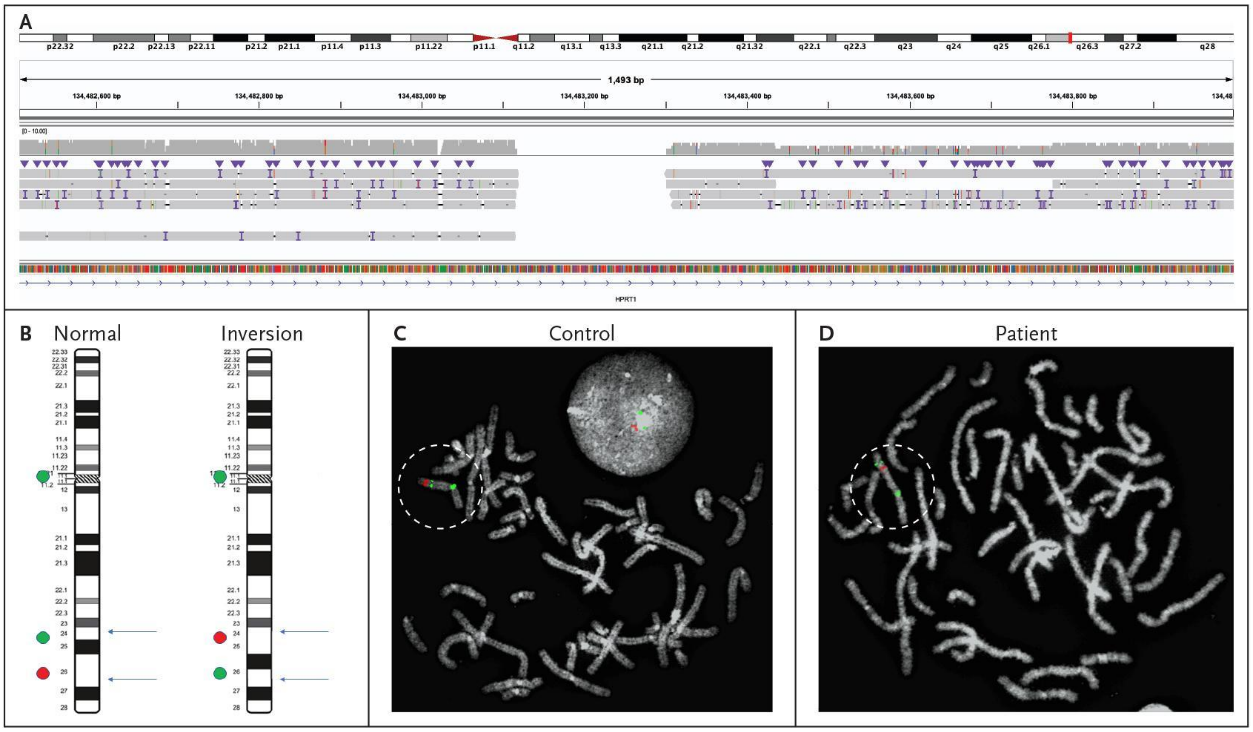

### Figure S30. S009 (*DMD*), IGV view of AGAA expansion and frequency in SSC samples.

**A.** Expansion of an AGAA repeat represents a variant of uncertain significance in a child with a clinical diagnosis of Duchenne muscular dystrophy, but no molecular diagnosis. This view shows the insertion in intron 16 of the gene that is homozygous in the proband (top panel) and heterozygous in the bottom panel (the insert is divided into a 72 bp insert and a 236 bp insert). Bottom panel is from the proband's unaffected full brother who does not have an insertion at this position. IGV view is of chrX:32,554,713-32,555,087. **B.** Histogram showing the number of individuals predicted to have an expansion of the AGAA repeat at the same position as the proband within the SSC collection. Position of the repeat expansion in the proband is shown in red, the length of the two haplotypes from the proband's mother is shown in red.

A:

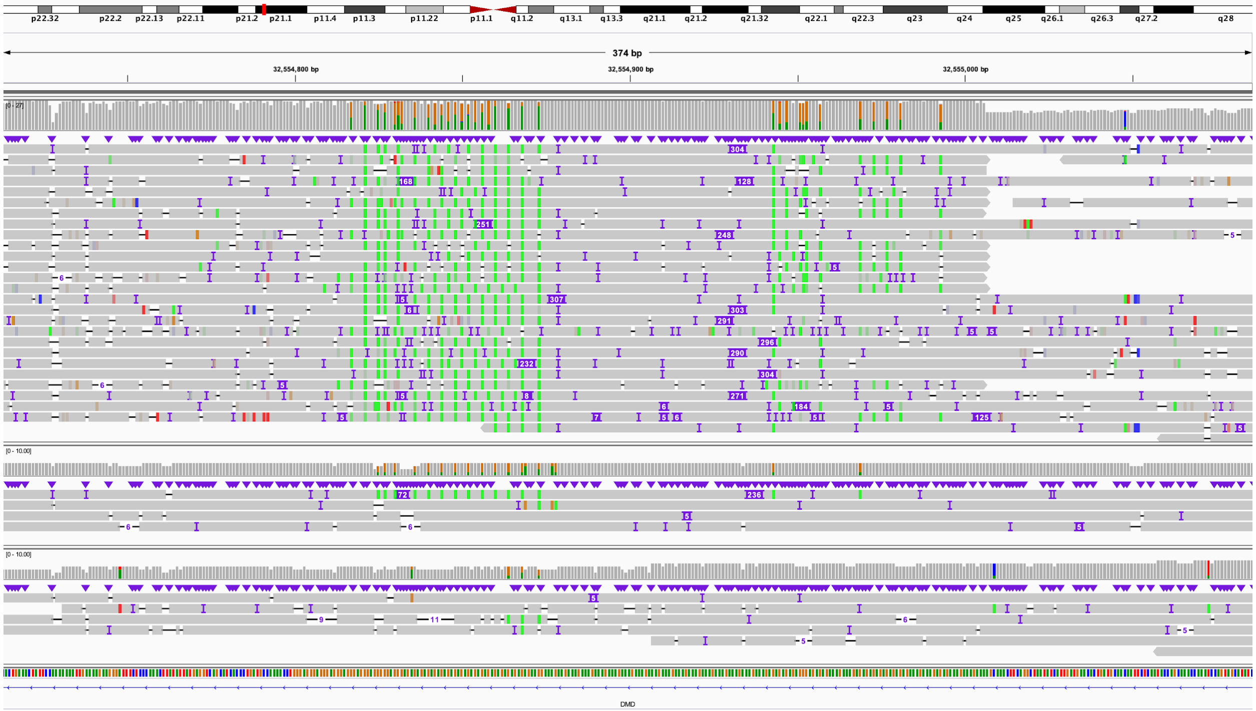

B:

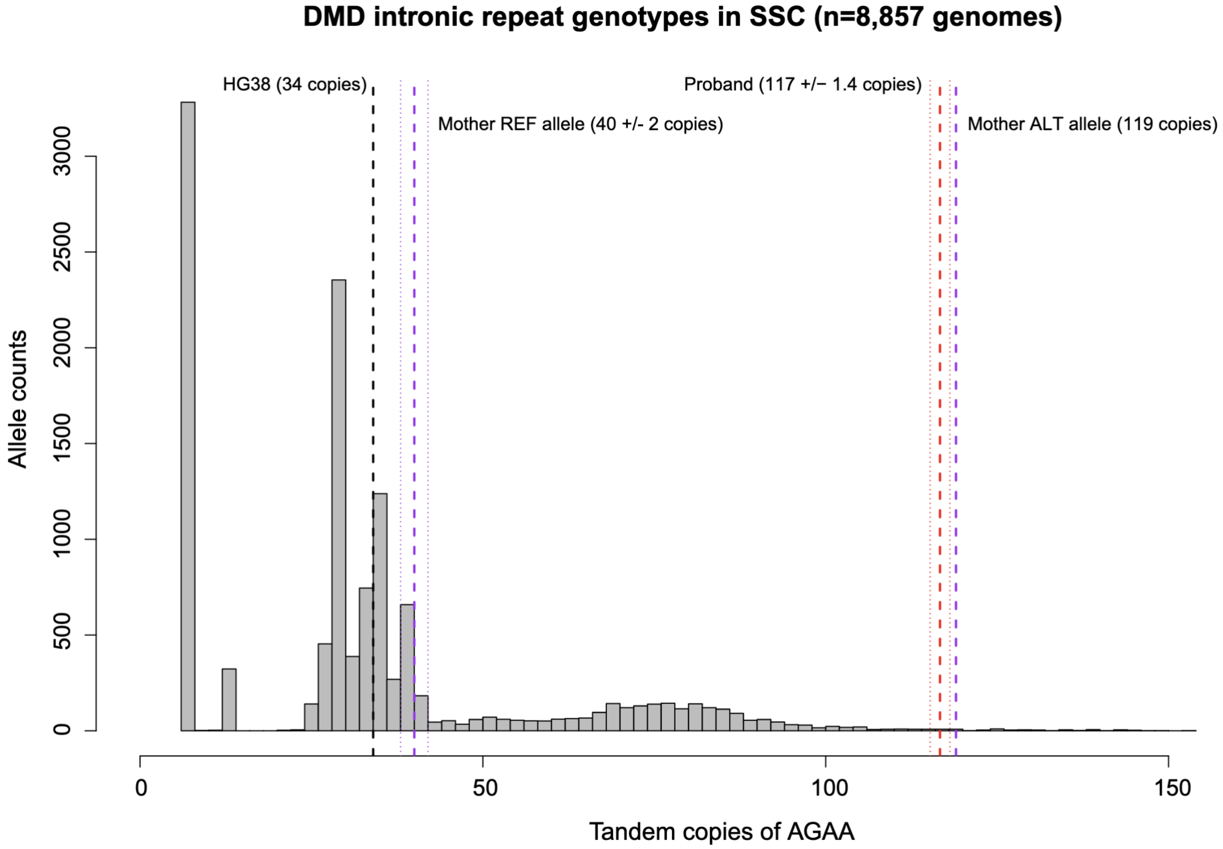
DMD intronic repeat genotypes in SSC (n=8,857 genomes)
HG38 (34 copies)
Mother REF allele (40 +/- 2 copies)
Proband (117 +/− 1.4 copies)
Mother ALT allele (119 copies)
Allele counts
0
500
1000
1500
2000
2500
3000
0
50
100
150
Tandem copies of AGAA

### Figure S31. S013 (*HPS1*), IGV view of inherited variant and deletion identified by LRS.

**A.** Screenshot of known paternally inherited pathogenic G>A variant. IGV view is of chr10:98,425,534-98,425,573. All reads are shown in the top track, haplotype 1 is the middle, and haplotype 2 is the bottom. Haplotype 2 is assumed to be the paternal track as the known paternally inherited G>A is in that track.
**B.** SV calling identified an approximately 1,900 bp deletion that included all of exon 3 on a different haplotype than the known G>A. Tracks are the same as in A. Screenshot from chr10:98,442,300-98,445,184.

A:

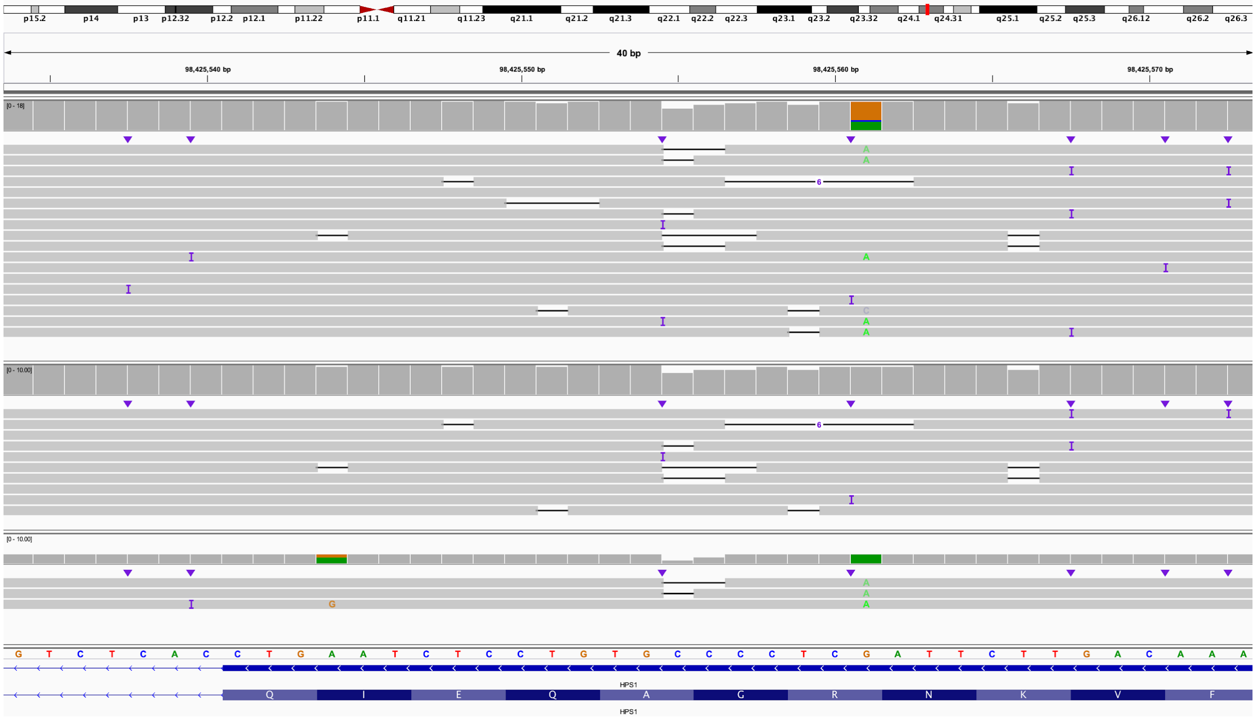

B:

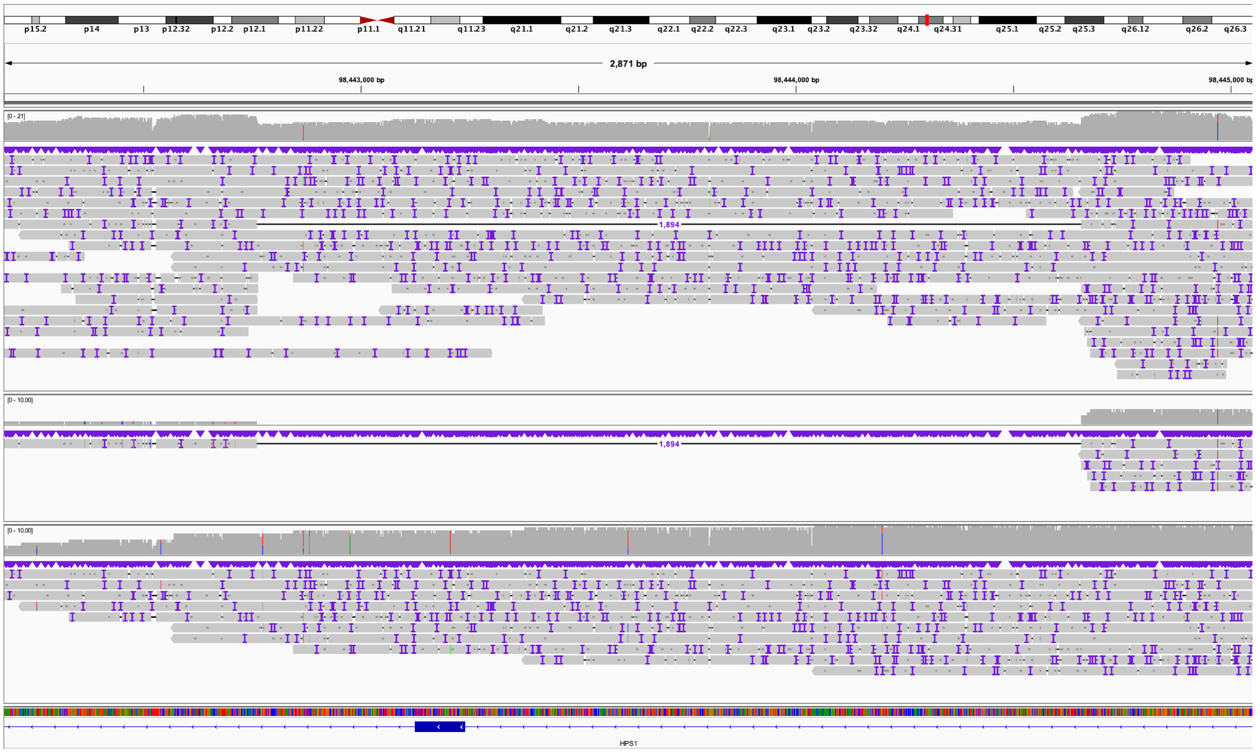
p15.2
p14
p13
p12.32
p12.2
p12.1
p11.22
p11.1
q11.21
q11.23
q21.1
q21.2
q21.3
q22.1
q22.2
q22.3
q23.1
q23.2
q23.32
q24.1
q24.31
q25.1
q25.2
q25.3
q26.12
q26.2
q26.3
2,871 bp
98,443,000 bp
98,444,000 bp
98,445,000 bp
[0 - 21]
1,894
[0 - 10.00]
1,894
[0 - 10.00]
HPS1

**Figure S32. S018 (*PAH*), known inherited splice variant identified, no second variant found.**

**A.** IGV view of previously known inherited pathogenic C>T splice variant. No second variant was found in this case. IGV view is of chr12:102,843,769-102,843,809. **B.** Bisulfite view of the entire *PAH* gene showing no hypermethylation near the 5' end of the gene (blue). Green blocks at the bottom of the image represent CpG islands. In this view, blue represents a CpG that is not methylated while red represents a methylated CpG.

A:

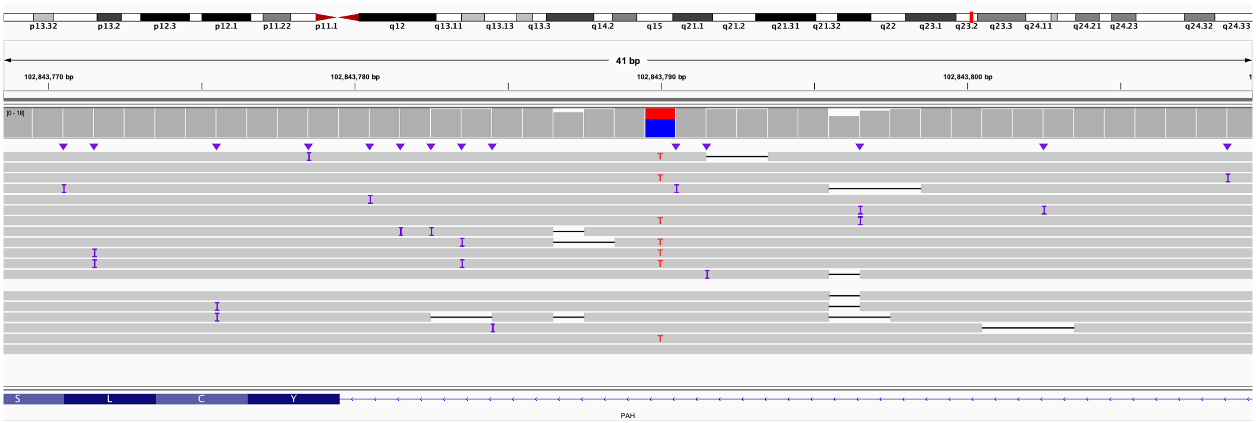

B:

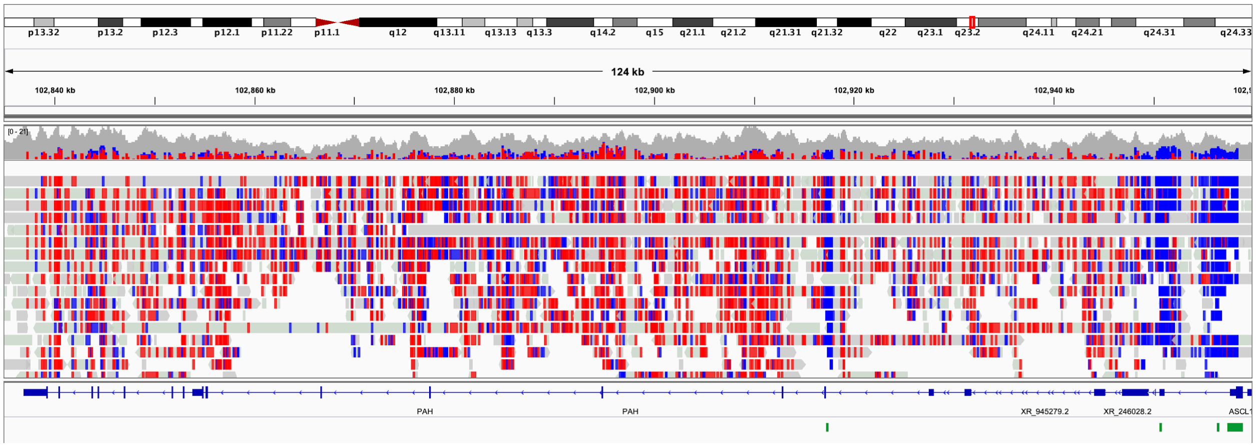

**Figure S33. S025 (*ABCA4*), the previously known variant and a 1,500 bp insertion can be phased into different haplotypes.**

**A.** IGV screenshot of long reads showing the previously identified inherited pathogenic variant. IGV View is of chr1:94,042,747-94,042,787.

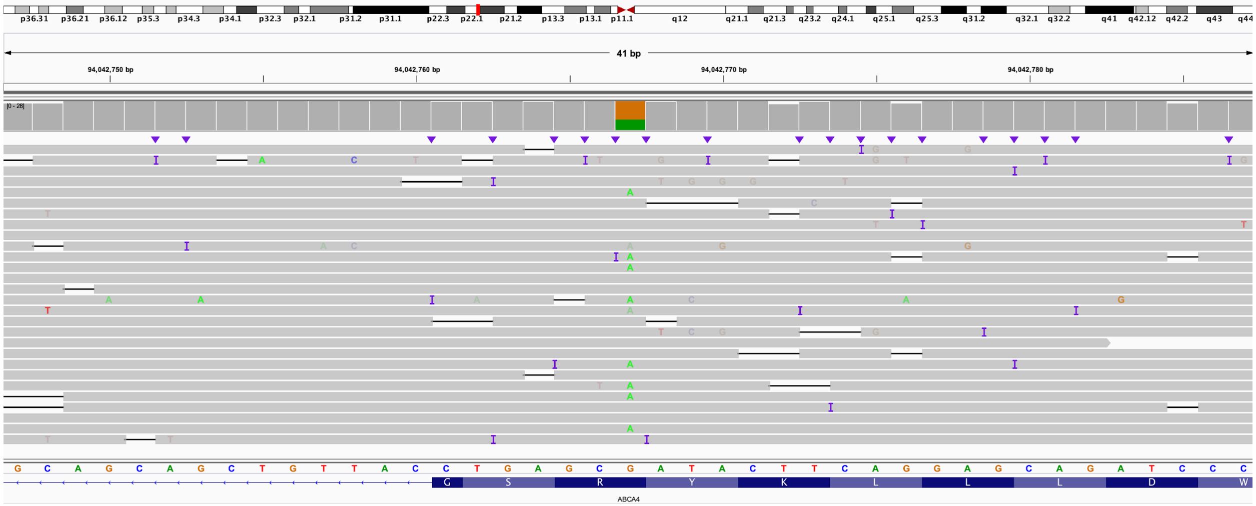

**B.** IGV screenshot of long reads showing two reads that include the 1,500 bp insertion in intron 1 of *ABCA4*. Several reads on either side of the region include the insertion and are soft clipped. IGV view is of chr1:94,120,209-94,120,274.

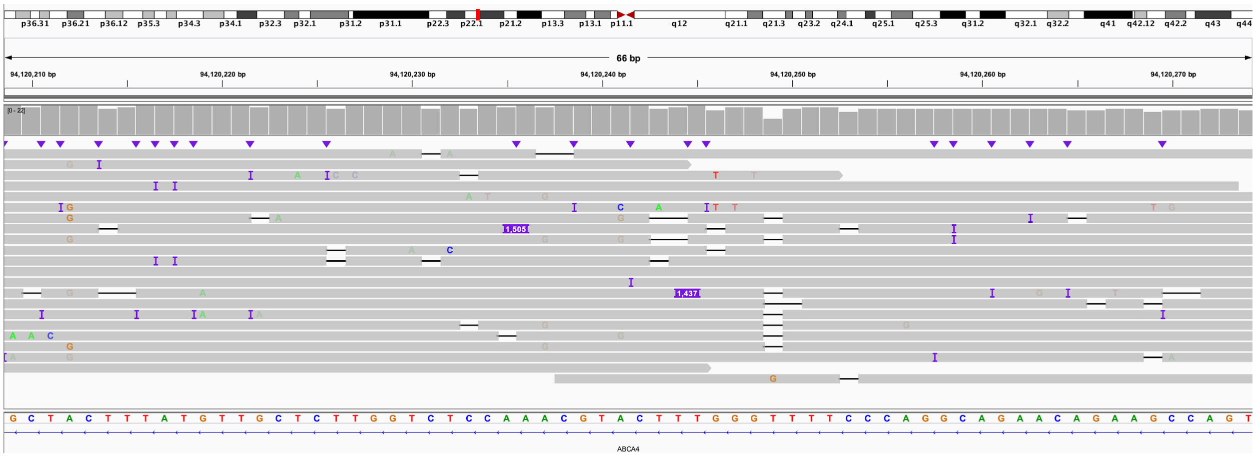

**C.** The linkage disequilibrium matrix represents R^2 and D' values in the lower and upper diagonal of the heatmap, respectively. Higher intensity colors represent higher linkage disequilibrium. Using orthogonal Illumina WGS data from S025, we identified all SNVs overlapping *ABCA4* using GATK, followed by calculating all pairwise R^2 and D' values using the 1000 Genomes Project Phase III genotypes.

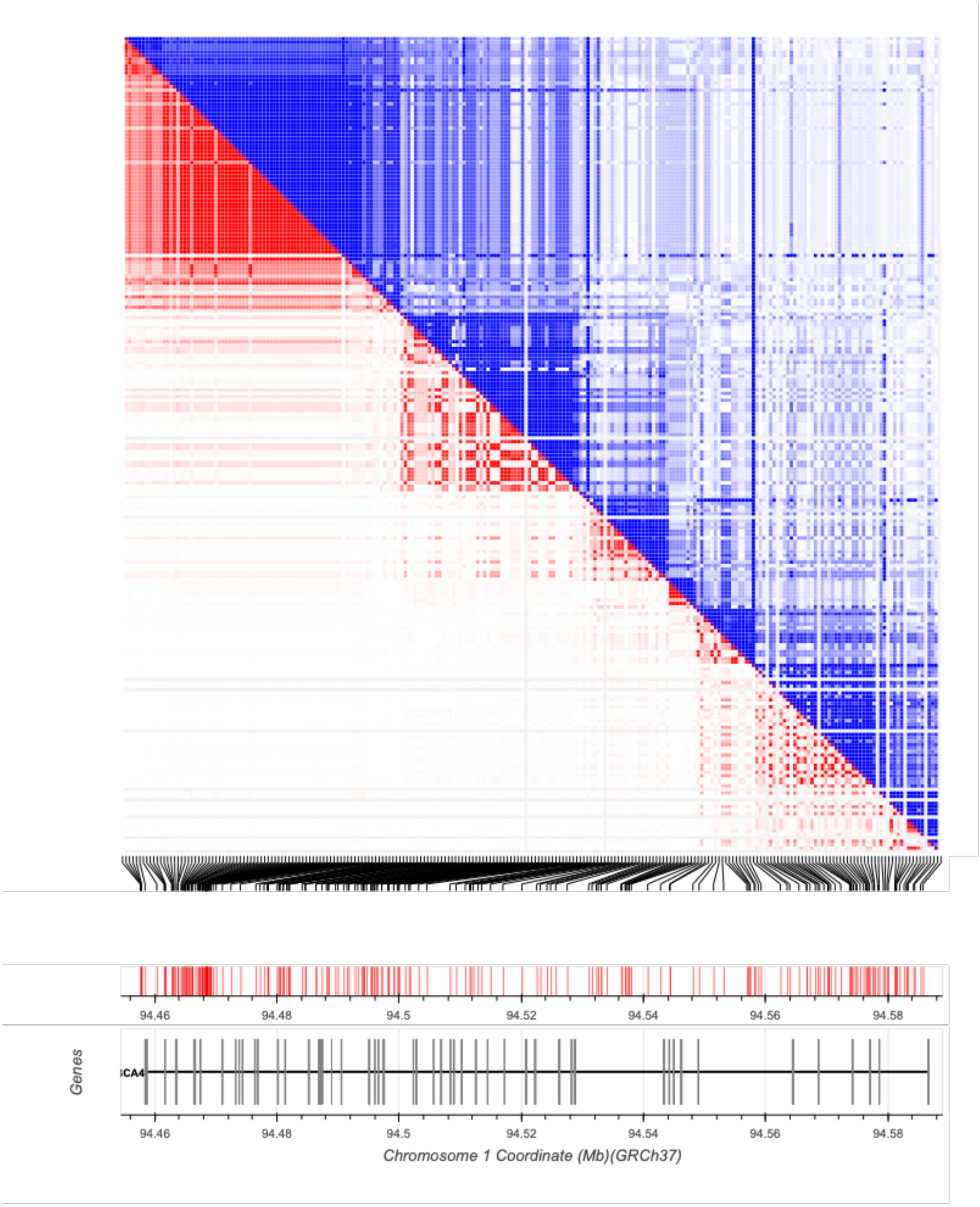
94.46
94.48
94.5
94.52
94.54
94.56
94.58
Genes
Chromosome 1 Coordinate (Mb)(GRCh37)

**D.** Phasing detail. The missense mutation on exon 22 corresponds to rs61750120 (G>A), while rs2184339 (T>C) has its alternative allele on the same allele as the 1.5 kbp insertion. Using the 1000 Genomes Project variation data (Machiela and Chanock 2015), we confirmed that the A allele of rs61750120 and the C allele of rs2184339 were never observed on the same haplotype (D'=1 and R^2=0.0001).

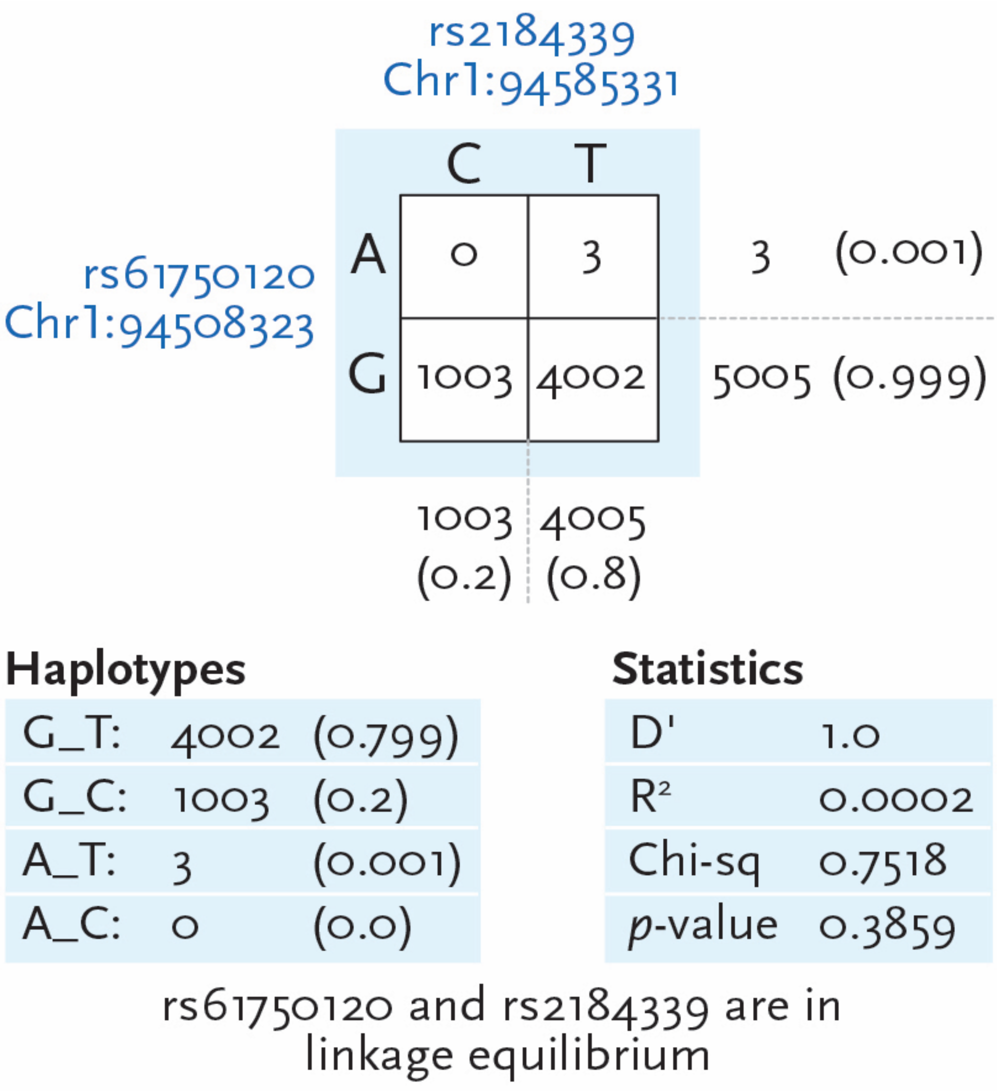

**E.** Phasing of long reads shows that rs2184339 (T>C) is present on reads with the 1,500 bp insertion. Most reads terminating at the insertion site are soft clipped. IGV view is of chr1:94,119,700-94,120,304.

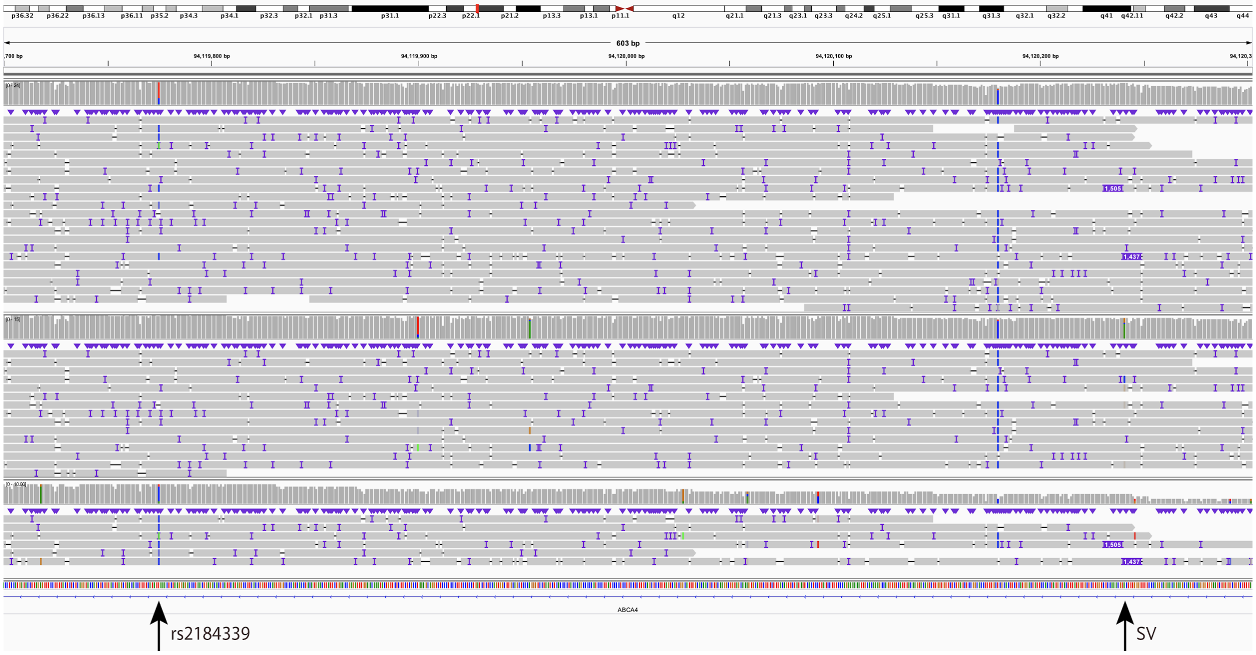

**F.** Analysis of short-read sequencing data reveals a 9 bp target site duplication at the position of the suspected insertion. IGV view is of chr1:94,120,221-94,120,260.

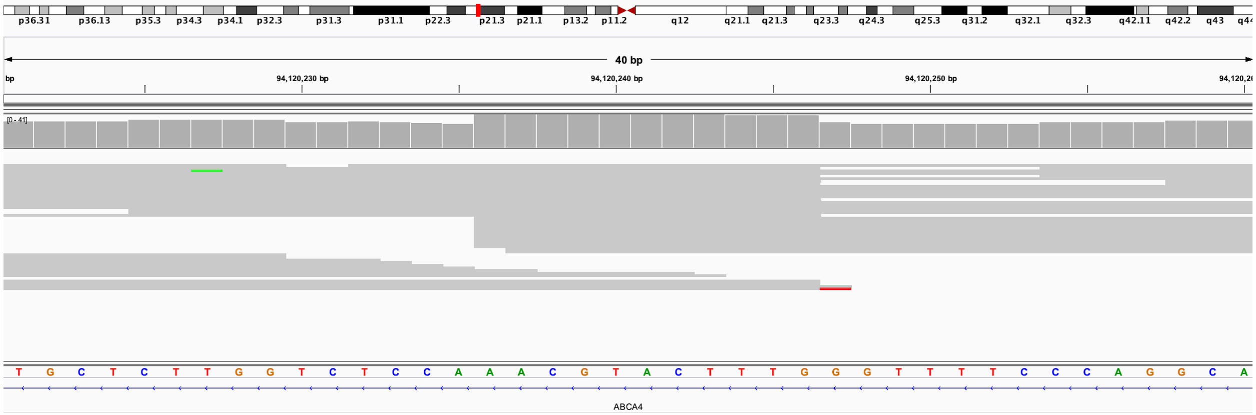

**Figure S34. S056 (*WDR19*), IGV views of inherited and splice variants identified by LRS.**

**A.** IGV view of known pathogenic inherited G>A. Top panel is all reads, middle panel is haplotype 1, and bottom panel is haplotype 2. IGV view is of chr4:39,273,009-39,273,048. **B.** An intronic C>A variant in haplotype 2 is predicted to increase the likelihood this position acts as both a splice acceptor and donor. IGV view is of chr4:39,216,917-39,216,957.

A:

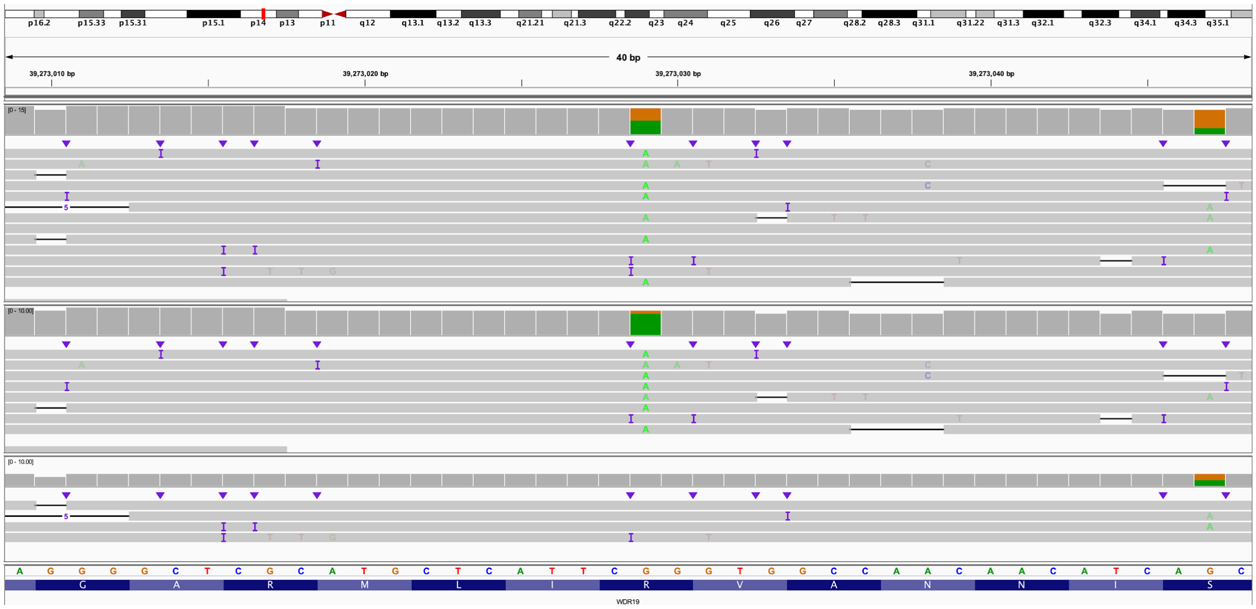

B:

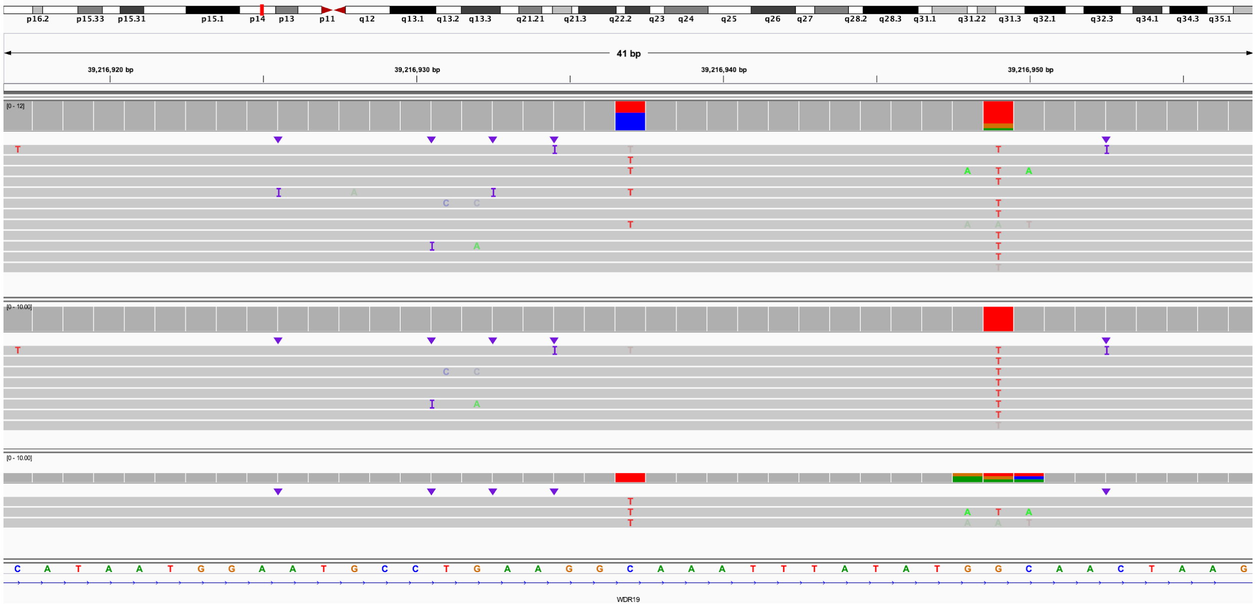

### Supplementary Tables

#### Table S1: Sample summary, DNA source, flow cells and libraries used per sample.

| Type | Patient | DNA Source | Flow cells used | Libraries used | Summary of sample |
|---|---|---|---|---|---|
| Missing Variant Case | S002 | Blood | 1 | 1 | Single variant in *ALMS1* |
| | S003 | Fibroblast | 2 | 2 | Single variant in *NPHP4* |
| | S004 | Blood | 3 | 3 | Single variant in *VARS2* |
| | S008 | Blood | 2 | 3 | No variant in *HPRT1* |
| | S009 | Blood | 3 | 4 | No variant in *DMD* |
| | S013 | Blood | 1 | 2 | Single variant in *HPS1* |
| | S018 | Blood | 1 | 1 | Single variant in *PAH* |
| | S025 | Blood | 1 | 1 | Single variant in *ABCA4* |
| | S056 | Blood | 1 | 1 | Single variant in *WDR19* |
| Repeat Expansion | S011 | Saliva | 1 | 1 | Expansions of both *ATXN3* and *ATXN8OS* |
| | S039 | Cell line | 1 | 1 | Expansion of *FMR1* |
| | S040 | Cell line | 1 | 1 | Expansion of *FXN* |
| | S041 | Cell line | 1 | 1 | Expansion of *FXN* |
| | 04-01 | Blood | 1 | 1 | Expansion and methylation of *XYLT1* |
| | 06-01 | Saliva | 2 | 2 | Expansion and methylation of *XYLT1* |
| | 04-02 | Fibroblasts | 1 | 1 | Mother of 04-01 |
| | 04-03 | Saliva | 1 | 1 | Father of 04-01 |
| | 06-02 | Saliva | 1 | 1 | Mother of 06-01 |
| | 06-03 | Saliva | 1 | 1 | Father of 06-01 |
| SV Case - Complex | S014 | Blood | 1 | 1 | Three noncontiguous deletions of chromosome 6 |
| | S020 | Blood | 2 | 3 | Multiple deletions of chromosomes 4 and 14, multiple translocations |
| | S021 | Blood | 2 | 2 | Multiple mosaic deletions of chromosome 8 |
| | S022 | Blood | 1 | 1 | Amplification of 4q with adjacent region of homozygosity |
| | S023 | Blood | 1 | 1 | Mosaic ring 18 |
| | S035 | Blood | 1 | 1 | Two duplications |
| | S036 | Blood | 2 | 2 | Four deletions on chromosome 10, multiple translocations |
| SV Case - Simple | BK144-03 | Blood | 1 | 1 | 22q13.3 deletion |
| | BK180-03 | Blood | 1 | 1 | 15q11-13 duplication |
| | BK294-03 | Blood | 1 | 1 | 22q11.2 duplication |
| | BK364-03 | Blood | 1 | 1 | 1p36.11 duplication |
| | BK397-101 | Blood | 1 | 1 | 16p11.2 deletion |
| | BK430-103 | Blood | 1 | 1 | 16p11.2 duplication |
| | BK482-101 | Blood | 1 | 1 | 1q21.1 duplication |
| | BK487-101 | Blood | 1 | 1 | 1q21 deletion |
| | BK506-03 | Blood | 1 | 1 | 5p15.33 deletion |
| | S016 | Blood | 1 | 1 | Tandem duplication in *CTNND2* |
| | S046 | Blood | 1 | 1 | Unbalanced translocation between chromosomes 4 and 15 |

### Table S2: Per sample sequencing targets, coverage, and average read length.

| Patient | Target chr | Target start | Target end | Size of target region (bp) | Target, gene, or target | Average coverage of whole genome (x) | Average coverage of target region (x) | Average length of all reads (bp) | Average length of reads in target region (bp) |
|---|---|---|---|---|---|---|---|---|---|
| **Simple SV cases** | | | | | | | | | |
| BK144-03 | 22 | 17,500,000 | 50,818,000 | 33,318,000 | 22q13.3 del | 3.5 | 14.7 | 539 | 2,133 |
| BK180-03 | 15 | 20,000,000 | 35,000,000 | 15,000,000 | 15q11-13 dup | 3.6 | 61.5 | 524 | 4,289 |
| BK294-03 | 22 | 17,500,000 | 50,818,000 | 33,318,000 | 22q11.2 dup | 3.5 | 33.7 | 524 | 4,617 |
| BK364-03 | 1 | 24,000,000 | 30,000,000 | 6,000,000 | 1p36.11 dup | 2.0 | 21.2 | 427 | 3,692 |
| BK397-101 | 16 | 26,500,000 | 35,300,000 | 8,800,000 | 16p11.2 del | 0.7 | 12.5 | 865 | 6,230 |
| BK430-103 | 16 | 25,000,000 | 35,000,000 | 10,000,000 | 16p11.2 dup | 1.5 | 17.0 | 612 | 4,057 |
| BK482-101 | 1 | 140,000,000 | 155,000,000 | 15,000,000 | 1q21.1 dup | 2.2 | 22.8 | 417 | 2,258 |
| BK487-101 | 1 | 140,000,000 | 155,000,000 | 15,000,000 | 1q21 del | 1.6 | 36.9 | 515 | 6,883 |
| BK506-03 | 5 | 1 | 8,000,000 | 7,999,999 | 5p15.33 del | 2.7 | 15.8 | 443 | 3,814 |
| S016 | 5 | 11,026,788 | 12,124,078 | 1,097,290 | *CTNND2* dup | 1.8 | 25.3 | 440 | 4,626 |
| S046 | 4 | 180,000,000 | 190,214,555 | 10,214,555 | translocation | 2.0 | 9.8 | 436 | 2,757 |
| | 15 | 90,000,000 | 101,991,189 | 11,991,189 | translocation | 2.0 | 17.7 | 436 | 3,068 |
| **Repeat expansion cases** | | | | | | | | | |
| S011 | 14 | 92,041,011 | 92,101,011 | 60,000 | *ATXN3* | 2.7 | 8.3 | 481 | 1,099 |
| | 13 | 70,109,384 | 70,169,384 | 60,000 | *ATXN8OS* | 2.7 | 9.2 | 481 | 1,197 |
| S039 | X | 147,862,037 | 147,962,037 | 100,000 | *FMR1* | 2.1 | 10.0 | 450 | 3,772 |
| S040 | 9 | 69,007,285 | 69,067,285 | 60,000 | *FXN* | 1.6 | 21.2 | 483 | 5,649 |
| S041 | 9 | 69,007,285 | 69,067,285 | 60,000 | *FXN* | 0.9 | 15.5 | 484 | 8,134 |
| 04-01 | 16 | 16,500,000 | 18,000,000 | 1,500,000 | *XYLT1* | 2.5 | 18.2 | 474 | 5,170 |
| 06-01 | 16 | 16,500,000 | 18,000,000 | 1,500,000 | *XYLT1* | 2.3 | 8.0 | 415 | 2,121 |
| 04-02 | 16 | 16,500,000 | 18,000,000 | 1,500,000 | *XYLT1* | 0.9 | 11.5 | 511 | 5,815 |
| 04-03 | 16 | 16,500,000 | 18,000,000 | 1,500,000 | *XYLT1* | 2.3 | 4.7 | 894 | 2,308 |
| 06-02 | 16 | 16,500,000 | 18,000,000 | 1,500,000 | *XYLT1* | 0.4 | 3.0 | 645 | 3,276 |
| 06-03 | 16 | 16,500,000 | 18,000,000 | 1,500,000 | *XYLT1* | 1.1 | 2.2 | 470 | 2,036 |
| **Cases with complex copy number changes** | | | | | | | | | |
| S014 | 6 | 150,000,000 | 165,000,000 | 15,000,000 | - | 2.7 | 13.3 | 420 | 2,105 |
| S020 | 2 | 1 | 36,300,000 | 36,299,999 | 1st run | 1.9 | 16.3 | 872 | 6,639 |
| | 4 | 65,000,000 | 78,000,000 | 13,000,000 | 1st run | 1.9 | 14.4 | 872 | 6,556 |
| | 10 | 59,474,150 | 80,320,091 | 20,845,941 | 1st run | 1.9 | 16.3 | 872 | 6,569 |
| | 14 | 20,881,587 | 24,829,792 | 3,948,205 | 1st run | 1.9 | 12.1 | 872 | 6.549 |
| | 10 | 41700000 | 59,475,000 | 17,775,000 | 2nd run | 0.6 | 9.2 | 785 | 7,158 |
| | 4 | 51800000 | 65,000,000 | 13,200,000 | 2nd run | 0.6 | 7.3 | 785 | 8,057 |
| | 4 | 78000000 | 100,000,000 | 22,000,000 | 2nd run | 0.6 | 7.2 | 785 | 8,072 |
| | 14 | 25900000 | 50,000,000 | 24,100,000 | 2nd run | 0.6 | 7.4 | 785 | 8,239 |
| S021 | 8 | 1 | 145,138,636 | 145,138,635 | - | 4.9 | 34.7 | 642 | 3,603 |
| S022 | 4 | 162611564 | 190000000 | 27,388,436 | - | 2.0 | 31.2 | 634 | 8,158 |
| | 15 | 22,000,000 | 23,500,000 | 1,500,000 | - | 2.0 | 31.8 | 634 | 6,003 |
| S023 | 18 | 1 | 5,000,000 | 4,999,999 | - | 2.2 | 23.1 | 561 | 5,106 |
| | 18 | 45,000,000 | 80,373,285 | 35,373,285 | - | 2.2 | 21.4 | 561 | 5,947 |
| S035 | 8 | 130,000,000 | 145,138,636 | 15,138,636 | - | 2.7 | 38.7 | 538 | 5,712 |
| | 16 | 10,000,000 | 20,000,000 | 10,000,000 | - | 2.7 | 38.5 | 538 | 5,671 |
| S036 | 6 | 114,000,000 | 124,000,000 | 10,000,000 | 1st run | 2.6 | 35.7 | 602 | 6,443 |
| | 18 | 7,000,000 | 16,000,000 | 9,000,000 | 1st run | 2.6 | 34.9 | 602 | 6,010 |
| | 10 | 20,000,000 | 38,000,000 | 18,000,000 | 1st run | 2.6 | 33.4 | 602 | 6,487 |
| | 10 | 50,000,000 | 60,000,000 | 10,000,000 | 1st run | 2.6 | 32.4 | 602 | 6,446 |
| | 5 | 1 | 46,000,000 | 45,999,999 | 2nd run | 0.7 | 7.8 | 828 | 8,413 |
| | 6 | 124,000,000 | 170,805,979 | 46,805,979 | 2nd run | 0.7 | 7.6 | 828 | 8,511 |
| | 18 | 1 | 7,000,000 | 6,999,999 | 2nd run | 0.7 | 7.8 | 828 | 7,535 |
| **Missing variant cases** | | | | | | | | | |
| S002 | 2 | 73,200,000 | 73,800,000 | 600,000 | *ALMS1* | 1.4 | 17.0 | 608 | 6,780 |
| S003 | 1 | 5,762,810 | 6,092,425 | 329,615 | *NPHP4* | 2.9 | 21.6 | 792 | 6,232 |
| S004 | 6 | 30814208 | 31026459 | 212,251 | *VARS2* | 3.5 | 41.9 | 640 | 6,443 |
| S008 | X | 134,360,165 | 134,600,668 | 240,503 | *HPRT1* | 2.5 | 7.2 | 1,151 | 7,160 |
| S009 | X | 31,019,219 | 33,439,460 | 2,420,241 | *DMD* | 6.9 | 29.5 | 501 | 3,344 |
| S013 | 10 | 98,316,193 | 98,546,963 | 230,770 | *HPS1* | 3.7 | 18.7 | 751 | 3,484 |
| S018 | 12 | 102,736,889 | 103,058,441 | 321,552 | *PAH* | 2.1 | 11.4 | 459 | 2,102 |
| S025 | 1 | 93,000,000 | 95,000,000 | 2,000,000 | *ABCA4* | 3.2 | 23.1 | 574 | 3,392 |
| S056 | 4 | 35,000,000 | 43,000,000 | 8,000,000 | *WDR19* | 1.4 | 22.1 | 537 | 8,282 |

### Table S3: Overview of individuals with a single structural variant.

| Patient | Previously known result | Confirmation of known result | Additional information gained with T-LRS | Previously reported event | Events seen by LRS |
|---|---|---|---|---|---|
| BK144-03 | 22q13.3 deletion | Identified the known deletion. | Identified exact position of deletion breakpoints. | 1 deletion | 1 deletion |
| BK180-03 | 15q11-13 duplication | Identified the known duplication. | None | 1 duplication | 1 duplication |
| BK294-03 | 22q11.2 duplication | Identified the known duplication. | None | 1 duplication | 1 duplication |
| BK364-03 | 1p36.11 duplication | Identified the known duplication. | Identified the exact position of the duplication breakpoints and found that the duplication is tandem. | 1 duplication | 1 duplication |
| BK397-101 | 16p11.2 deletion | Identified the known deletion. | None | 1 deletion | 1 deletion |
| BK430-103 | 16p11.2 duplication | Identified the known duplication. | None | 1 duplication | 1 duplication |
| BK482-101 | 1q21.1 duplication | Identified the known duplication. | None | 1 duplication | 1 duplication |
| BK487-101 | 1q21 deletion | Identified the known deletion. | None | 1 deletion | 1 deletion |
| BK506-03 | 5p15.33 deletion | Identified the known deletion. | Appears to be an unbalanced translocation between chr5 and chr12. Short-read data from patient agrees and the translocation is not found in parental short-read data. | 1 deletion | Translocation between chr5 and chr12 |
| S016 | Tandem duplication in *CTNND2* | Identified the known duplication and confirmed it is tandem. | None | 1 duplication | 1 duplication |
| S046 | Unbalanced translocation between chr4 and chr15. | Identified the translocation via coverage. | Identified exact position of translocation breakpoints. | Translocation | Translocation |

**Table S4: Overview of individuals with repeat expansions, including expected and observed repeat expansion sizes.**

| Patient | Gene(s) | Repeat | Previously reported repeat size | Repeat position | BP added to window for repeat assay | Number of reads spanning window | Average repeat lengths (% difference from expected if known) |
|---|---|---|---|---|---|---|---|
| S011 | *ATXN3* | CAG | Clinically reported as 74 and 28 repeats | chr14:92071012-92071052 | 100 | 10 | 25 (11%), 77 (4%) |
| S011 | *ATXN8 OS* | CAG | Clinically reported as 80 and 25 repeats | chr13:70139385-70139429 | 25 | 8 | 15 (40%), 73 (9%) |
| S039 | *FMR1* | CGG | Coriell sample 06897, reported as 477 repeats. | chrX:147911980-147912111 | 200 | 3 | 386 (19%) |
| S040 | *FXN* | GAA | Coriell sample 16789, reported as 750 and 1000 repeats. | chr9:69,037,285-69,037,302 | 50 | 4 | 333 (56%), 1049 (5%) |
| S041 | *FXN* | GAA | Coriell sample 15850, reported as 650 and 1030 repeats. | chr9:69,037,285-69,037,302 | 50 | 9 | 647 (1%), 958 (7%) |
| 04-01 | *XYLT1* | GGC | Previously reported expansion. | chr16:17470921-17470922 | 500 | 4 | 758 |
| 04-02 | *XYLT1* | GGC | Mother of 04-01, reported as one wild-type and one premutation allele. | chr16:17470921-17470922 | 50 | 16 | 97, 221 |
| 04-03 | *XYLT1* | GGC | Father of 04-01, reported as two wild-type alleles. | chr16:17470921-17470922 | 50 | 2 | 71 |
| 06-01 | *XYLT1* | GGC | Previously reported expansion. | chr16:17470921-17470922 | 500 | 5 | 224 |
| 06-02 | *XYLT1* | GGC | Mother of 06-01, reported as one wild-type and one premutation allele | chr16:17470921-17470922 | 50 | 1 | 81 |
| 06-03 | *XYLT1* | GGC | Father of 06-01, reported as two wild-type alleles | chr16:17470921-17470922 | 50 | 1 | 78 |

### Table S5: Per-read details of individuals with *ATXN3*, *ATXN8OS*, *FMR1*, and *FXN* repeat expansions.

| Read ID | Repeat Length (bp) | Repeats (Estimate) | Repeat Group | Average Number of Repeats |
|---|---|---|---|---|
| **S011 - ATXN3** | | | | |
| c4b0552d-118e-4d6b-9411-09b0861262fe | 55 | 18 | 1 | 25 |
| 616cf7f8-a13f-429a-b0c5-825f1a15da3d | 72 | 24 | 1 | |
| e133bc59-4c37-4c5c-8ce9-b206180058eb | 77 | 26 | 1 | |
| eadb0392-728a-4437-8e5e-c2814638894f | 79 | 26 | 1 | |
| 2f107739-f4a6-40f0-b52b-252132f992f3 | 85 | 28 | 1 | |
| 137c7831-3cb4-4f7c-99b6-196a969702dc | 218 | 73 | 2 | 77 |
| 72ed9321-3c6b-4cab-b508-83d6efc1efc0 | 230 | 77 | 2 | |
| b6a6b164-c767-4514-b723-4a6643a0f3f5 | 231 | 77 | 2 | |
| e6c40588-04c8-4a17-b740-fc1990995267 | 232 | 77 | 2 | |
| 0fa6df78-d10e-42f8-9e1e-6db5a907d858 | 250 | 83 | 2 | |
| **S011 - ATXN8OS** | | | | |
| 9712bb2f-b1fb-40db-9e05-6f708b6443e4 | 41 | 14 | 1 | 15 |
| 3e0a7cc1-ee7f-47ff-97f4-9516cbebb21e | 43 | 14 | 1 | |
| d5eea8a6-f1d5-459f-b865-bff3c35a26b3 | 44 | 15 | 1 | |
| c03fb6ab-a4a1-4518-b946-ad650f578e25 | 46 | 15 | 1 | |
| 70fec058-32ab-4270-aaa6-1984af7b1402 | 197 | 66 | 2 | 73 |
| 4f49b32d-139c-4beb-b831-9c4a5b7a19be | 206 | 69 | 2 | |
| a7304699-9d25-4843-b587-cb84de848b84 | 226 | 75 | 2 | |
| 489bde02-eae6-4270-a7db-1e488b61ccfd | 245 | 82 | 2 | |
| **S039 - FMR1** | | | | |
| 5fac5744-2c5c-4211-88c8-5b9ee3a20752 | 759 | 253 | 1 | 386 |
| b215ab66-d863-46d3-878e-69d3d855a302 | 1448 | 483 | 1 | |
| ef287abe-e2a4-4b0e-8121-1de041a53f77 | 1268 | 423 | 1 | |
| **S040 - FXN** | | | | |
| 44970178-6c34-48e8-afd9-f529d6bda0e2 | 985 | 328 | 1 | 333 |
| 1e915a81-c955-4c88-8946-b3e73d371b64 | 1004 | 335 | 1 | |
| afddceb3-0802-404d-a61c-24ecdba887db | 1007 | 336 | 1 | |
| a66018cb-a390-4cb6-89f4-b9c0073e68f8 | 3147 | 1049 | 2 | 1049 |
| **S041 - FXN** | | | | |
| a93466ef-353b-403d-a2ec-d5439867b826 | 1788 | 596 | 1 | 647 |
| b1ac947d-e98d-41f4-b5ca-d464e3668388 | 1939 | 646 | 1 | |
| 8164bc3d-6cae-4928-a975-e695c48bc757 | 1963 | 654 | 1 | |
| 0bfda482-97a2-42c8-ad67-798ebf2bd5b8 | 2079 | 693 | 1 | |
| bf8f85f8-f878-4f38-aefc-b7aa0009b8ae | 2415 | 805 | 2 | 958 |
| 89100931-0b2b-4938-a882-f7c81652858a | 2895 | 965 | 2 | |
| 20ad64ba-a1b7-4371-93f8-e3c425c2c008 | 2996 | 999 | 2 | |
| d13da846-f4f3-45ba-806a-0f77f50bb71a | 3000 | 1000 | 2 | |
| c0a0f7ad-c30a-4e40-ab51-fe00b1ab699a | 3058 | 1019 | 2 | |

### Table S6: Per-read details of *XYLT1* repeat expansions.

| Read ID | Repeat Length (bp) | Insert between *KpnI* cut sites (bp) | Repeats | Repeat Group | Average Number of Repeats |
|---|---|---|---|---|---|
| **04-01 - XYLT1** | | | | | |
| 71ed6a24-3280-465a-8114-338c0eb6308d | 1846 | 4435 | 615 | 1 | 758 |
| c522d590-4165-47ac-9d4e-fb98f391bc13 | 2861 | 5450 | 954 | 1 | |
| beb4b5c7-293c-4f69-b275-d393c6a3ae34 | 1591 | 4180 | 530 | 1 | |
| a09207e2-a9cc-4cb6-9e12-f9fb56214399 | 2800 | 5389 | 933 | 1 | |
| **04-02 - XYLT1** | | | | | |
| b2bebdb7-8e03-4b86-833f-445c1c2cd1d0 | 167 | 2756 | 56 | 1 | 97 |
| 7c629eb4-81f6-4d27-988b-b827b81419a2 | 282 | 2871 | 94 | 1 | |
| 775ca9d2-16b7-43f2-8855-6ef25cfd26a0 | 283 | 2872 | 94 | 1 | |
| 0cce5876-2f35-44f8-835f-867f08e4e571 | 288 | 2877 | 96 | 1 | |
| 879df89f-3293-4bec-93ec-073c985c3a18 | 288 | 2877 | 96 | 1 | |
| 83d4020e-16f1-4b94-95d8-1bf48fca6de6 | 295 | 2884 | 98 | 1 | |
| 970f3e26-1807-42d3-b15e-4c85a25b6ab4 | 433 | 3022 | 144 | 1 | |
| 0edbaf14-3d7e-46a2-a03d-268c784560f3 | 603 | 3192 | 201 | 2 | 221 |
| b09a91bf-aee4-4fc0-a7bf-51d7cb1ad126 | 615 | 3204 | 205 | 2 | |
| 40767c26-5d0c-467d-bdcf-c1a6346d4229 | 641 | 3230 | 214 | 2 | |
| f85efe4f-2d32-4a6d-a217-165c6dbdc661 | 645 | 3234 | 215 | 2 | |
| b0a508ce-069d-43ac-865e-7b7cd900eb70 | 653 | 3242 | 218 | 2 | |
| 5c6ac009-694d-4f86-9511-8708f952a81c | 688 | 3277 | 229 | 2 | |
| 4e315aca-60a8-4e61-8cd6-c08f129d701a | 693 | 3282 | 231 | 2 | |
| 227e44ce-2051-4169-9b25-17cfdb50a4c5 | 716 | 3305 | 239 | 2 | |
| 011e133a-ea64-43fd-a4f4-0c2cfc8fe56e | 725 | 3314 | 242 | 2 | |
| **04-03 - XYLT1** | | | | | |
| bba6d56d-5293-4c5b-bd03-327dc12f7ece | 218 | 2807 | 73 | 1 | 71 |
| 370138b9-090a-4fb0-80bc-01723888a41b | 209 | 2798 | 70 | 1 | |
| **06-01 - XYLT1** | | | | | |
| c66d7ab6-2ec4-417d-9445-d1d68ef3cf8e | 668 | 3257 | 223 | 1 | 224 |
| 7db4e75c-c2f7-4029-9c6e-a8e06ff269a6 | 720 | 3309 | 240 | 1 | |
| 210471f7-0e15-436e-862f-85557ad3d4ff | 734 | 3323 | 245 | 1 | |
| 8f3c0400-f5dc-47e2-bc69-611a477181bf | 622 | 3211 | 207 | 1 | |
| 5e39f080-90c9-4f3d-ba8b-bed01bef0caa | 616 | 3205 | 205 | 1 | |
| **06-02 - XYLT1** | | | | | |
| 236a7c09-2c88-4ee8-b2b3-fda20bc625c1 | 243 | 2832 | 81 | 1 | 81 |
| **06-03 - XYLT1** | | | | | |
| 9e7e1ed4-ebc2-4b30-a8ea-4c561d87e793 | 235 | 2824 | 78 | 1 | 78 |

### Table S7: Summary of individuals with complex SVs, including previously known events and new events identified by T-LRS.

| Patient | Previously known result | Confirmation of known result | Additional information gained with T-LRS | Total previously known events | Total new events |
|---|---|---|---|---|---|
| S014 | Three noncontiguous deletions of chromosome 6. | Identified the three deletions seen on array | Identified two additional deletions and one rearrangement, and found no pathogenic or likely pathogenic variants in deleted regions. | 3 | 3 |
| S020 | Two deletions of 4q and one of 14q identified on the array. Karyotype revealed translocation between chromosomes 2, 4, 10, and 14. | Identified the three deletions seen on array and several translocations between chromosomes 2, 4, 10, and 14. | Identified one additional deletion on chromosome 4 and two on chromosome 10. Found the exact position of all translocation breakpoints, revealing two that bisected genes with AD phenotypes. Identified additional rearrangement breakpoints within chromosome 10 and chromosome 14 not involved in a deletion or translocation. | 7 | 22 |
| S021 | Terminal 3.2 Mbp loss of 8p23.3 to p23.2 with a copy state of 1. Adjacent interstitial 3.7 Mbp mosaic loss of 8p23.2 to p23.1 with a copy state between 1 and 2. Terminal 50 Mbp mosaic gain of 8q22.1 to q24.3 with a copy state between 2 and 3. | Confirmed mosaic loss and gain seen on array. | Find that a common 400 kbp duplication on 8p not reported on the array appears to be inverted and attached to the chromosome 8 not carrying a deletion. | 3 | 1 |
| S022 | Focal amplification of 4q (copy state 4 or greater) with adjacent region of homozygosity and 15q11.2 duplication. | See the amplification of 4q32, homozygosity of 4q, and duplication in 15q11.2. | Determined the structure of the tandem amplification on chr 4 and confirmed 5 copies of the amplification (6 with the wild-type chromosome). Do not see breakpoints of the 15q11.2 duplication, but do see the duplication. No methylation differences or pathogenic variants in the homozygous region. | 2 | 0 |
| S023 | Mosaic ring 18 in 40% of cells. | See decreased coverage in regions reported on the array. | None. | 2 | 0 |
| S035 | Duplications of 8q24 and 16p13.11 | Both duplications observed. | Found that 8q duplication is tandem and identified exact breakpoints. Could not identify the exact position of the 16p13.11 duplication that is flanked by segmental duplications. | 2 | 0 |
| S036 | Four deletions on chr 10, karyotype found translocation between chr 18 and 6 and pericentric inversion of chr10. | Identified four deletions and found the t(6;18). | Found additional translocations between chromosome 10 and chromosome 5 as well as additional rearrangements on chromosomes 6, 10, and 18. | 7 | 13 |

### Table S8: Known and new events observed for individuals with known complex SVs.

| | Known | | | | Observed | | | | New Events | | | |
|---|---|---|---|---|---|---|---|---|---|---|---|---|
| Patient | Deletion | Duplication | Translocation | Rearrangement | Deletion | Duplication | Translocation | Rearrangement | Deletion | Duplication | Translocation | Rearrangement |
| S014 | 3 | 0 | 0 | 0 | 5 | 0 | 0 | 1 | 2 | 0 | 0 | 1 |
| S020 | 3 | 0 | 4 | 0 | 5 | 0 | 11 | 13 | 2 | 0 | 7 | 13 |
| S021 | 2 | 1 | 0 | 0 | 2 | 1 | 0 | 1 | 0 | 0 | 0 | 1 |
| S022 | 0 | 2 | 0 | 0 | 0 | 2 | 0 | 0 | 0 | 0 | 0 | 0 |
| S023 | 2 | 0 | 0 | 0 | 2 | 0 | 0 | 0 | 0 | 0 | 0 | 0 |
| S035 | 0 | 2 | 0 | 0 | 0 | 2 | 0 | 0 | 0 | 0 | 0 | 0 |
| S036 | 4 | 0 | 2 | 1 | 6 | 0 | 8 | 6 | 2 | 0 | 6 | 5 |

#### Table S9: Details of focal amplification of 4q in patient S022.

| Region | Size (bp) | Copy number estimate | Coverage | Increase or decrease in coverage from prior |
|---|---|---|---|---|
| chr4:160,000,000-160,143,267 | 143,267 | 2 | 31.3 | |
| chr4:160,143,268-160,165,900 | 22,632 | 3 | 43.9 | 12.6 |
| chr4:160,165,901-160,248,169 | 82,268 | 4 | 62.7 | 18.8 |
| chr4:160,248,170-160,450,953 | 202,783 | 5 | 76.0 | 13.3 |
| chr4:160,450,954-162,685,228 | 2,234,274 | 6 | 93.8 | 17.8 |
| chr4:162,685,229-162,689,149 | 3,920 | 4 | 57.6 | -36.2 |
| chr4:162,689,150-162,700,000 | 110,850 | 2 | 30.6 | -27.0 |

#### Table S10: Rearrangement numbers and sizes for patient S014.

| Chr | Segment | Start | End | Size (bp) | Source/Type |
|---|---|---|---|---|---|
| 6 | A | 1 | 153,854,905 | 153,854,904 | - |
| 6 | B | 153,854,906 | 154,095,842 | 240,936 | Deletion |
| 6 | C | 154,095,843 | 154,587,147 | 491,304 | Inversion |
| 6 | D | 154,587,148 | 154,588,799 | 1,651 | Deletion |
| 6 | E | 154,588,800 | 154,661,283 | 72,483 | - |
| 6 | F | 154,661,284 | 156,456,517 | 1,795,233 | Deletion |
| 6 | G | 156,456,518 | 156,460,113 | 3,595 | Inversion |
| 6 | H | 156,460,118 | 156,464,495 | 4,377 | - |
| 6 | I | 156,464,502 | 157,015,124 | 550,622 | - |
| 6 | J | 157,015,125 | 158,766,820 | 1,751,695 | Deletion |
| 6 | K | 158,766,821 | 164,500,069 | 5,733,248 | - |
| 6 | L | 164,500,070 | 164,654,921 | 154,851 | Deletion |
| 6 | M | 164,654,922 | 170,805,979 | 6,151,057 | - |

#### Table S11: Genes impacted by S014 breakpoints.

| Chr | Breakpoint | Gene affected by breakpoint | Previously known based on clinical testing? | OMIM phenotype |
|---|---|---|---|---|
| 6 | B/C | *OPRM1* | Y | none |
| 6 | I/J | *ARID1B* | Y | Autosomal dominant: Coffin-Siris syndrome I |
| 6 | K/L | *EZR* | Y | none |

### Table S12: Rearrangement numbers and sizes for patient S020.

| Chr | Segment | Start | End | Size (bp) | Source/Type | Validated with HiFi? |
|---|---|---|---|---|---|---|
| 2 | A | 1 | 7,777,990 | 7,777,989 | Derivative 14 | Yes |
| 2 | B | 7,777,996 | 16,235,712 | 8,457,716 | Derivative 4 | Yes |
| 2 | C | 16,235,711 | 242,193,529 | 225,957,818 | Derivative 2 | Yes |
| 4 | A | 1 | 67,572,970 | 67,572,969 | Derivative 4 | Yes |
| 4 | B | 67,572,971 | 70,140,499 | 2,567,528 | Deletion | Yes |
| 4 | C | 70,140,500 | 73,061,492 | 2,920,992 | Derivative 14 | Yes |
| 4 | D | 73,061,493 | 73,129,559 | 68,066 | Deletion | Yes |
| 4 | E | 73,129,560 | 73,446,127 | 316,567 | Derivative 10 | Yes |
| 4 | F | 73,446,129 | 73,771,830 | 325,701 | Derivative 10 | Yes |
| 4 | G | 73,771,831 | 74,581,615 | 809,784 | Deletion | Yes |
| 4 | H | 74,581,616 | 74,794,351 | 212,735 | Derivative 14 | Yes |
| 4 | I | 74,794,352 | 74,831,249 | 36,897 | Derivative 14 | Yes |
| 4 | J | 74,831,250 | 75,039,118 | 207,868 | Derivative 4 | Yes |
| 4 | K | 75,039,137 | 77,012,176 | 1,973,039 | Derivative 4 | Yes |
| 4 | L | 77,012,175 | 79,626,721 | 2,614,546 | Derivative 14 | Yes |
| 4 | M | 79,626,720 | 81,632,570 | 2,005,850 | Derivative 4 | Yes |
| 4 | N | 81,632,570 | 190,214,555 | 108,581,985 | Derivative 2 | Yes |
| 10 | A | 1 | 50,382,094 | 50,382,093 | Derivative 10 | Yes |
| 10 | B | 50,382,092 | 50,563,505 | 181,413 | Derivative 10 | Yes |
| 10 | C | 50,563,501 | 50,649,777 | 86,276 | Derivative 10 | Yes |
| 10 | D | 50,694,777 | 51,260,021 | 565,244 | Derivative 10 | Yes |
| 10 | E | 51,260,022 | 51,840,878 | 580,856 | Deletion | Yes |
| 10 | F | 51,840,879 | 53,548,486 | 1,707,607 | Derivative 10 | Yes |
| 10 | G | 53,548,487 | 53,563,514 | 15,027 | Derivative 14 | Yes |
| 10 | H | 53,563,515 | 53,584,434 | 20,919 | Deletion | Yes |
| 10 | I | 53,584,435 | 56,035,820 | 2,451,385 | Derivative 4 | Yes |
| 10 | J | 56,035,823 | 65,519,829 | 9,484,006 | Derivative 4 | Yes |
| 10 | K | 65,519,828 | 67,462,235 | 1,942,407 | Derivative 10 | Yes |
| 10 | L | 67,462,259 | 74,772,273 | 7,310,014 | Derivative 10 | Yes |
| 10 | M | 74,772,269 | 76,402,382 | 1,630,113 | Derivative 4 | Yes |
| 10 | N | 76,402,383 | 133,797,422 | 57,395,039 | Derivative 4 | Yes |
| 14 | A | 1 | 20,934,810 | 20,934,809 | Derivative 14 | Yes |
| 14 | B | 20,934,811 | 22,881,931 | 1,947,120 | Deletion | Yes |
| 14 | C | 22,881,932 | 107,043,718 | 84,161,786 | Derivative 10 | Yes |
| | | | Total Size | 334,539,803 | Derivative 2 | |
| | | | Total Size | 151,177,985 | Derivative 4 | |
| | | | Total Size | 146,979,108 | Derivative 10 | |
| | | | Total Size | 34,512,995 | Derivative 14 | |

### Table S13: Genes impacted by S020 breakpoints.

| Chr | Breakpoint | Gene affected by breakpoint | Previously known based on clinical testing? | OMIM phenotype |
|---|---|---|---|---|
| 4 | A/B | *STAP1* | Y | none |
| 4 | B/C | *CSN1S2BP* | N | none |
| 4 | C/D | *COX18* | N | none |
| 4 | D/E | *ANKRD17* | N | none |
| 4 | E/F | *AFP* | N | Autosomal dominant: hereditary persistence of alpha-fetoprotein; autosomal recessive: alpha-fetoprotein deficiency |
| 4 | H/I | *BTC* | N | none |
| 4 | J/K | *PARM1* | N | none |
| 4 | K/L | *SEPTIN11* | N | none |
| 10 | A/B, B/C | *SGMS1* | N | none |
| 10 | D/E, E/F | *PRKG1* | N | Autosomal dominant: aortic aneurysm, familial thoracic |
| 10 | K/L | *CTNNA3* | N | Autosomal dominant: arrhythmogenic right ventricular dysplasia, familial |
| 10 | M/N | *LRMDA* | N | Autosomal recessive: Albinism, oculocutaneous, type VII |

### Table S14: Rearrangement numbers and sizes for patient S036.

| Chr | Segment | Start | End | Size (bp) | Source/Type |
|---|---|---|---|---|---|
| 5 | A | 1 | 22,881,851 | 22,881,850 | Derivative 5 |
| 5 | B | 22,881,853 | 28,650,540 | 5,768,687 | Derivative 5 |
| 5 | C | 28,650,540 | 28,826,587 | 176,047 | Derivative 10 |
| 5 | D | 28,826,592 | 181,538,259 | 152,711,667 | Derivative 5 |
| 6 | A | 1 | 123,005,546 | 123,005,545 | Derivative 6 |
| 6 | B | 123,005,550 | 124,505,324 | 1,499,774 | Derivative 18 |
| 6 | C | 124,505,731 | 126,657,919 | 2,152,188 | Derivative 18 |
| 6 | D | 126,657,920 | 126,680,039 | 22,119 | Deletion |
| 6 | E | 126,680,040 | 170,805,979 | 44,125,939 | Derivative 18 |
| 10 | A | 1 | 24,286,112 | 24,286,111 | Derivative 10 |
| 10 | B | 24,286,113 | 24,725,849 | 439,736 | Deletion |
| 10 | C | 24,725,850 | 25,278,388 | 552,538 | Derivative 5 |
| 10 | D | 25,278,408 | 34,622,799 | 9,344,391 | Derivative 10 |
| 10 | E | 34,622,800 | 35,972,590 | 1,349,790 | Deletion |
| 10 | F | 35,972,591 | 49,775,899 | 13,803,308 | Derivative 10 |
| 10 | G | 49,775,901 | 54,672,866 | 4,896,965 | Derivative 5 |
| 10 | H | 54,672,867 | 55,305,313 | 632,446 | Deletion |
| 10 | I | 55,305,314 | 55,632,879 | 327,565 | Derivative 5 |
| 10 | J | 55,632,880 | 56,736,281 | 1,103,401 | Deletion |
| 10 | K | 56,736,282 | 133,797,422 | 77,061,140 | Derivative 10 |
| 18 | A | 1 | 6,275,133 | 6,275,132 | Derivative 6 |
| 18 | B | 6,275,134 | 6,280,055 | 4,921 | Deletion |
| 18 | C | 6,280,056 | 7,279,970 | 999,914 | Derivative 6 |
| 18 | D | 7,279,968 | 80,373,285 | 73,093,317 | Derivative 18 |
| | | | Total Length | 187,139,272 | Derivative 5 |
| | | | Total Length | 130,280,591 | Derivative 6 |
| | | | Total Length | 124,670,997 | Derivative 10 |
| | | | Total Length | 120,871,218 | Derivative 18 |

### Table S15: Genes impacted by S036 breakpoints.

| Chr | Breakpoint | Gene affected by breakpoint | Previously known based on clinical testing? | OMIM phenotype |
|---|---|---|---|---|
| 6 | A/B | *CLVS2* | N | none |
| 6 | B/C | *NKAIN2* | N | none |
| 10 | A/B | *KIAA1217* | N | none |
| 10 | C/D | *GPR158* | N | none |
| 10 | D/E | *PARD3* | Y | none |
| 10 | F/G, G/H, H/I | *PCDH15* | Y | Autosomal dominant and autosomal recessive: Usher syndrome, type 1D/F digenic |
| 18 | A/B, C/D | *L3MBTL4* | N | none |

### Table S16: Summary of individuals with missing variants.

| Patient | Clinical workup | Confirmation of inherited variant | Variant found by T-LRS | Confirmation or supporting findings |
|---|---|---|---|---|
| S002 | CMA normal, exome with paternally inherited stop in *ALMS1*, consistent with suspected diagnosis of Alstrom syndrome. Deletion/duplication analysis of *ALMS1* negative. | Confirmed the known paternally inherited variant. | *Alu* insertion in exon 20 | Clinically confirmed |
| S003 | CMA normal, exome with paternally inherited stop in *NPHP4*, consistent with suspected diagnosis of NPH. Deletion/duplication analysis of *NPHP4* negative. | Confirmed the known paternally inherited variant. | Intronic splice variant | Confirmed by qPCR |
| S004 | CMA normal, exome identified single paternally inherited pathogenic variant in *VARS2*, deletion/duplication analysis of *VARS2* negative. | Confirmed the known paternally inherited pathogenic p.A420T. | No second hit found. | n/a |
| S008 | Biochemical diagnosis of Lesch-Nyhan based on enzyme analysis. CMA normal, sequencing and deletion/duplication analysis of *HPRT1* negative. | No pathogenic SNVs or copy number changes observed, consistent w/clinical testing. | Identified an inversion within *HPRT1* and confirmed with PCR. | Clinically confirmed |
| S009 | Suspected diagnosis of Duchenne muscular dystrophy. Sequencing and deletion/duplication analysis of *DMD* negative. Muscle biopsy consistent with diagnosis. The proband's maternal uncle died of muscular dystrophy. | No pathogenic SNVs or copy number changes observed, consistent w/clinical testing. | Identified a candidate AGAA repeat expansion in intron 16. | Mom is heterozygous, unaffected brother has wild-type allele |
| S013 | CMA normal, exome with paternally inherited stop in *HPS1*, consistent with suspected diagnosis of Hermansky-Pudlak syndrome. No deletion/duplication analysis done. | Confirmed the known paternally inherited variant. | Identified 1,900 bp deletion that removes exon 3 and confirmed with PCR. | PCR confirmed, pending exon-level array |
| S018 | Elevated phenylalanine, exome with single pathogenic variant in *PAH*. | Confirmed the known inherited variant. | No second hit found. | No second hit found with short-read sequencing |
| S025 | Diagnosed with Stargardt disease by clinical retinal exam. Exome with single pathogenic variant in *ABCA4*. No second hit seen on research WGS. | Confirmed the known pathogenic variant. | ~1,500 bp transposable element insertion in the first intron. | Insertion is present in Illumina short-read data. |
| S056 | CMA normal, exome with single pathogenic variant in *WDR19*, consistent with presumed diagnosis of Sensenbrenner syndrome. Deletion/duplication analysis negative. | Confirmed the known inherited variant. | Intronic splice variant | None |

#### Table S17: Reads used to calculate length of AGAA motif in patient S009.

| Read ID/Contig | Start Pos | End Pos | Period Size | Copy Number | Percent Matches | Percent Indels | Score | Motif |
|---|---|---|---|---|---|---|---|---|
| 268a7f22-1c5e-40a6-b50a-764cc962ecd4:20612-21234 | 84 | 562 | 4 | 118.5 | 94 | 2 | 753 | AAGA |
| 33430113-f292-4c8a-abaa-ef24685b0275:9126-9747 | 90 | 540 | 4 | 117 | 79 | 15 | 393 | AAGA |
| ab709b56-560e-442b-b415-6538973c978d:4402-4955 | 66 | 485 | 4 | 107.5 | 86 | 10 | 505 | AAGA |
| 157b7b75-7a60-4e4e-8717-6bfa1fc8424f:3823-4453 | 81 | 569 | 4 | 121.8 | 92 | 5 | 720 | AAGA |
| 4683a48e-1bd4-430f-a361-712f000d8ff1:14088-14711 | 84 | 562 | 4 | 119.8 | 93 | 4 | 727 | AAGA |
| 622462ec-1093-4532-91dc-a8366bd9f712:1867-2467 | 83 | 537 | 4 | 116.8 | 91 | 6 | 620 | AAAG |
| aee7ddf8-5473-4c67-9e45-65ddb6b3512b:1667-2278 | 78 | 556 | 4 | 120.2 | 91 | 5 | 695 | AGAA |
| 4885dba3-b149-4870-b903-eb240a6f34e0:1488-2110 | 85 | 562 | 4 | 119.5 | 96 | 1 | 762 | AAGA |
| ffc51d09-c8a1-4308-a843-9540a3c812ea:1304-1906 | 83 | 546 | 4 | 113.8 | 78 | 13 | 297 | AAGA |
| **Average** | | | | 117.2 | | | | |
| **Excluded read IDs containing TGTT motif** | | | | | | | | |
| 1fd6ae08-37dc-47e1-8804-28117b821352 | | | | | | | | |
| 8f3ea358-76e1-41a7-88a7-ef0e287b4c77 | | | | | | | | |
| 37a6233f-91bb-4a5f-909d-e3951cc855be | | | | | | | | |

**Table S18: Predicted strength of the canonical splice donor site at the Exon 1–Intron 1 boundary of *ABCA4* (NM_000350) and alternative sites introduced by the ~1,500 bp insertion in patient S025.**

While the SV is absent from all accessible population genetic databases (gnomAD, BRAVO), assessment by multiple *in silico* prediction tools suggest a strong likelihood of pathogenicity. Specifically, the deep residual neural network SpliceAI predicts a splice-altering consequence of the pre-mRNA through the introduction of a *de novo* donor site by the insertion event (Δ score = 0.33, high recall). The ~1,500 bp insertion sequence itself contains two strong alternative splice donors and six alternative splice acceptors. The relative strength of these alternative donors and acceptors indicates a high probability of competition with canonical sites. One alternative donor site in particular exhibits a stronger splice signal than the canonical donor site at the Exon 1–Intron 1 boundary is the most probable site of alternative splicing leading to the inclusion of the 5' portion of the intron in the final mRNA transcript.

| Splice donor | SpliceSiteFinder-like | MaxEntScan | NNSPLICE | GeneSplicer | Human Splice Finder |
|---|---|---|---|---|---|
| Canonical | 81.70 | 8.90 | 1.00 | 5.20 | 84.70 |
| Alternative | 90.10 | 8.00 | 0.90 | - | 95.00 |
| Alternative | 81.50 | 7.20 | 0.80 | 0.60 | 90.80 |
| **Splice acceptor** | **SpliceSiteFinder-like** | **MaxEntScan** | **NNSPLICE** | **GeneSplicer** | **Human Splice Finder** |
| Canonical | 95.25 | 10.87 | 0.99 | 12.03 | 92.11 |
| Alternative | 93.50 | 10.00 | 1.00 | 3.20 | 90.60 |
| Alternative | 88.54 | 7.30 | 1.00 | 1.50 | 79.83 |
| Alternative | 77.80 | 6.10 | 0.60 | 2.30 | 78.50 |
| Alternative | 83.20 | 5.90 | 0.70 | - | 85.80 |
| Alternative | 71.70 | 5.70 | - | 1.60 | 79.50 |
| Alternative | 74.60 | 6.90 | 0.60 | - | 78.30 |

Length of color bars (blue = donor, green = acceptor) are proportioned to the respective scales of each algorithm: SpliceSiteFinder-Like (0-100), MaxEntScan (0-16), NNSPLICE (0-1), Gene Splicer (0-15), Human Splice Finder v.3.1 (0-100).

### Supplementary Appendix References

1000 Genomes Project Consortium, Adam Auton, Lisa D. Brooks, Richard M. Durbin, Erik P. Garrison, Hyun Min Kang, Jan O. Korbel, et al. 2015. “A Global Reference for Human Genetic Variation.” *Nature* 526 (7571): 68–74.

Audano, Peter A., Arvis Sulovari, Tina A. Graves-Lindsay, Stuart Cantsilieris, Melanie Sorensen, Annemarie E. Welch, Max L. Dougherty, et al. 2019. “Characterizing the Major Structural Variant Alleles of the Human Genome.” *Cell* 176 (3): 663–75.e19.

Benson, G. 1999. “Tandem Repeats Finder: A Program to Analyze DNA Sequences.” *Nucleic Acids Research* 27 (2): 573–80.

Chaisson, Mark J. P., Ashley D. Sanders, Xuefang Zhao, Ankit Malhotra, David Porubsky, Tobias Rausch, Eugene J. Gardner, et al. 2019. “Multi-Platform Discovery of Haplotype-Resolved Structural Variation in Human Genomes.” *Nature Communications* 10 (1): 1–16.

Edge, Peter, and Vikas Bansal. 2019. “Longshot Enables Accurate Variant Calling in Diploid Genomes from Single-Molecule Long Read Sequencing.” *Nature Communications* 10 (1): 4660.

Gel, Bernat, and Eduard Serra. 2017. “karyoploteR: An R/Bioconductor Package to Plot Customizable Genomes Displaying Arbitrary Data.” *Bioinformatics* 33 (19): 3088–90.

Guo, Hui, Michael H. Duyzend, Bradley P. Coe, Carl Baker, Kendra Hoekzema, Jennifer Gerdts, Tychele N. Turner, et al. 2019. “Genome Sequencing Identifies Multiple Deleterious Variants in Autism Patients with More Severe Phenotypes.” *Genetics in Medicine: Official Journal of the American College of Medical Genetics* 21 (7): 1611–20.

Heller, David, and Martin Vingron. 2019. “SVIM: Structural Variant Identification Using Mapped Long Reads.” *Bioinformatics* 35 (17): 2907–15.

Huizing, Marjan, May Christine V. Malicdan, Bernadette R. Gochuico, and William A. Gahl. 2000. “Hermansky-Pudlak Syndrome.” In *GeneReviews®*, edited by Margaret P. Adam, Holly H. Ardinger, Roberta A. Pagon, Stephanie E. Wallace, Lora J. H. Bean, Karen Stephens, and Anne Amemiya. Seattle (WA): University of Washington, Seattle.

Jaganathan, Kishore, Sofia Kyriazopoulou Panagiotopoulou, Jeremy F. McRae, Siavash Fazel Darbandi, David Knowles, Yang I. Li, Jack A. Kosmicki, et al. 2019. “Predicting Splicing from Primary Sequence with Deep Learning.” *Cell* 176 (3): 535–48.e24.

Karczewski, Konrad J., Laurent C. Francioli, Grace Tiao, Beryl B. Cummings, Jessica Alföldi, Qingbo Wang, Ryan L. Collins, et al. 2020. “The Mutational Constraint Spectrum Quantified from Variation in 141,456 Humans.” *Nature* 581 (7809): 434–43.

Killick, Rebecca, and Idris Eckley. 2014. “Changepoint:an R Package for Changepoint Analysis.” *Journal of Statistical Software* 58 (3): 19.

LaCroix, Amy J., Deborah Stabley, Rebecca Sahraoui, Margaret P. Adam, Michele Mehaffey, Kelly Kernan, Candace T. Myers, et al. 2019. “GGC Repeat Expansion and Exon 1 Methylation of XYLT1 Is a Common Pathogenic Variant in Baratela-Scott Syndrome.” *American Journal of Human Genetics* 104 (1): 35–44.

Lee, Isac, Roham Razaghi, Timothy Gilpatrick, Norah Sadowski, Fritz J. Sedlazeck, and Winston Timp. 2018. “Simultaneous Profiling of Chromatin Accessibility and Methylation on Human Cell Lines with Nanopore Sequencing.” https://doi.org/10.1101/504993.

Li, Heng. 2018. “Minimap2: Pairwise Alignment for Nucleotide Sequences.” *Bioinformatics* 34 (18): 3094–3100.

Li, Heng, Bob Handsaker, Alec Wysoker, Tim Fennell, Jue Ruan, Nils Homer, Gabor Marth, Goncalo Abecasis, Richard Durbin, and 1000 Genome Project Data Processing Subgroup. 2009. “The Sequence Alignment/Map Format and SAMtools.” *Bioinformatics* 25 (16): 2078–79.

Loman, Nicholas J., Joshua Quick, and Jared T. Simpson. 2015. “A Complete Bacterial Genome Assembled de Novo Using Only Nanopore Sequencing Data.” *Nature Methods* 12

(8): 733–35.
Luo, Ruibang, Chak-Lim Wong, Yat-Sing Wong, Chi-Ian Tang, Chi-Man Liu, Chi-Ming Leung, and Tak-Wah Lam. 2020. “Exploring the Limit of Using a Deep Neural Network on Pileup Data for Germline Variant Calling.” *Nature Machine Intelligence* 2 (4): 220–27.
Machiela, Mitchell J., and Stephen J. Chanock. 2015. “LDlink: A Web-Based Application for Exploring Population-Specific Haplotype Structure and Linking Correlated Alleles of Possible Functional Variants.” *Bioinformatics* 31 (21): 3555–57.
McLaren, William, Laurent Gil, Sarah E. Hunt, Harpreet Singh Riat, Graham R. S. Ritchie, Anja Thormann, Paul Flicek, and Fiona Cunningham. 2016. “The Ensembl Variant Effect Predictor.” *Genome Biology* 17 (1): 122.
Miller, Danny E., Audrey Squire, and James T. Bennett. 2020. “A Child with Autism, Behavioral Issues, and Dysmorphic Features Found to Have a Tandem Duplication within CTNND2 by Mate-Pair Sequencing.” *American Journal of Medical Genetics. Part A* 182 (3): 543–47.
Paisey, Richard B., Rick Steeds, Tim Barrett, Denise Williams, Tarekegn Geberhiwot, and Meral Gunay-Aygun. 2003. “Alström Syndrome.” In *GeneReviews®*, edited by Margaret P. Adam, Holly H. Ardinger, Roberta A. Pagon, Stephanie E. Wallace, Lora J. H. Bean, Karen Stephens, and Anne Amemiya. Seattle (WA): University of Washington, Seattle.
Payne, Alexander, Nadine Holmes, Thomas Clarke, Rory Munro, Bisrat Debebe, and Matthew Loose. 2020. “Nanopore Adaptive Sequencing for Mixed Samples, Whole Exome Capture and Targeted Panels.” https://doi.org/10.1101/2020.02.03.926956.
Pedersen, Brent S., Ryan M. Layer, and Aaron R. Quinlan. 2016. “Vcfanno: Fast, Flexible Annotation of Genetic Variants.” *Genome Biology* 17 (1): 118.
Rentzsch, Philipp, Daniela Witten, Gregory M. Cooper, Jay Shendure, and Martin Kircher. 2019. “CADD: Predicting the Deleteriousness of Variants throughout the Human Genome.” *Nucleic Acids Research* 47 (D1): D886–94.
Sedlazeck, Fritz J., Philipp Rescheneder, Moritz Smolka, Han Fang, Maria Nattestad, Arndt von Haeseler, and Michael C. Schatz. 2018. “Accurate Detection of Complex Structural Variations Using Single-Molecule Sequencing.” *Nature Methods* 15 (6): 461–68.
Sulovari, Arvis, Ruiyang Li, Peter A. Audano, David Porubsky, Mitchell R. Vollger, Glennis A. Logsdon, Human Genome Structural Variation Consortium, et al. 2019. “Human-Specific Tandem Repeat Expansion and Differential Gene Expression during Primate Evolution.” *Proceedings of the National Academy of Sciences of the United States of America* 116 (46): 23243–53.
Wenger, Aaron M., Paul Peluso, William J. Rowell, Pi-Chuan Chang, Richard J. Hall, Gregory T. Concepcion, Jana Ebler, et al. 2019. “Accurate Circular Consensus Long-Read Sequencing Improves Variant Detection and Assembly of a Human Genome.” *Nature Biotechnology* 37 (10): 1155–62.
